# Comparison of Human Melanoma Single Cell Profiles to Evolutionary Medicine Model *Xiphophorus* Provides Insights in Disease Control

**DOI:** 10.1101/2025.09.22.677876

**Authors:** Yanting Xing, Rachel Carrol, X Maggs, Wes C. Warren, Manfred Schartl, Yuan Lu

## Abstract

Melanoma remains one of the deadliest forms of cancer. Despite recent therapeutic advances including immune checkpoint inhibitors and small-molecule kinase inhibitors, patients frequently develop treatment resistance. Novel models are needed to devise strategies that overcome resistance and further reduce melanoma-related mortality. Interspecies hybrid fish from the Xiphophorus lineage develop mutant *Epidermal Growth Factor Receptor* (EGFR)-driven melanomas that display morphology, bulk gene expression, disease initiation and progression processes mimicking those of human melanomas. These similarities have enabled their comparative use in evaluating why human melanomas exhibit cancer cell plasticity, including dynamic transitions between proliferative and invasive states. However, it remains unclear whether *Xiphophorus* melanomas recapitulate some or all of these features. To address this, we performed single-nucleus RNA sequencing (snRNAseq) analysis of *Xiphophorus* melanomas. Multiple cancer cell types mirroring the human melanoma cell populations were identified, including proliferative cancer cells, dedifferentiated neural crest-like cells, mesenchymal-like cancer cells in addition to fibroblast, endothelial, and immune cells. Employing comparative analyses with results from human melanoma studies, it is demonstrated that *Xiphophorus* melanomas faithfully mimic the cellular heterogeneity observed in human melanoma.

## Introduction

Despite recent advancements in melanoma therapies and a corresponding decline in melanoma-related mortality, melanoma still remains one of the most lethal cancers, accounting for approximately 75% of skin cancer deaths (1-4). The five-year survival rate for patients with advanced melanoma stays low, ranging from 34% to 52% (5). Currently available treatments include immunotherapies (e.g., PD-1 inhibitor, CTLA-4 inhibitor, LAG-3 inhibitor), small-molecule kinase inhibitors targeting BRAF and MEK and recently approved tumor-infiltrating lymphocyte (TIL) therapy (2, 6-9).

Melanoma cells exhibit profound plasticity at the molecular, cellular, and morphological levels, allowing for reversible phenotypic transitions between a locally restricted, proliferative state and an invasive, non-proliferative state. Single cell studies of human melanoma underscore this diversity of malignant cell states within a single tumor, including proliferative, invasive, and drug-tolerant phenotypes (10, 11). One of the most comprehensive evaluations of the melanoma tumor microenvironment (TME) showed the malignant cells exhibited two distinct transcriptional cell states, high and low levels of the MITF transcription factor, the later cells also having elevated levels of the AXL kinase (11). The proliferative phenotype is generally more sensitive to both kinase inhibitors and immunotherapy, whereas the invasive phenotype demonstrates greater therapeutic tolerance (12-19). In addition, the tumor microenvironment is composed of diverse stromal and immune populations, such as cancer-associated fibroblasts, endothelial cells, tumor-associated macrophages, exhausted T cells, and B-cell aggregates, each contributing to tumor growth and immune evasion (20, 21). These findings highlight both the phenotypic plasticity of melanoma cells and the complex cellular ecosystem that underlies disease progression and therapeutic response.

However, c therapies often lead to the development of resistance (7, 22, 23). Resistance arises from both intrinsic and extrinsic mechanisms, including innate and acquired resistance to kinase inhibitors, changes in melanoma immunogenicity, impaired immune cell trafficking, inefficient T-cell priming, evasion of cell death, neovascularization, modulation of the tumor microenvironment, extracellular matrix remodeling, metabolic competition, T-cell exhaustion, immune checkpoint upregulation, and alterations in the gut microbiota (2, 24).

Given these challenges, there is an urgent need for evaluating new models of melanoma progression that can either target key steps in the phenotype-switching process or bypass resistance altogether. Interspecies hybrids between select *Xiphophorus* species can spontaneously develop genetically programmed melanoma (25). The *Xiphophorus* cancer model has been instrumental in elucidating critical steps in tumor initiation, progression, and invasion. It has also led to the identification of several clinically relevant melanoma biomarkers, including SLC45A2 for early diagnosis, Osteopontin for malignancy, and Cystathionase (for senescence escape). Notably, discovery of the oncogene *xmrk* in *Xiphophorus* preceded and helped inform the discovery of the RAS/RAF/ERK pathway as a key driver in human melanomagenesis (for a comprehensive review see (25)).

Melanomagenesis in these hybrids follows Mendelian segregation patterns, involving inheritance of a well characterized oncogene and variant alleles of modulator genes. Well-characterized hybrid combinations that develop melanoma include *X. maculatus × X. hellerii, X. maculatus × X. couchianus*, and *X. birchmanni × X. malinche* (25). In each case, the melanoma-bearing hybrids inherit a mutant *Epidermal Growth Factor Receptor* gene (*xmrk*) from one parent and lack functional modulators (such as Cdkn2ab, Rab3d, Myrip, Adgre5/CD97, or Long Neurotoxin OH-57-like) from the other (26-31). The *xmrk* oncogene activates multiple downstream signaling pathways that promote melanophore dedifferentiation, tumor proliferation, survival, angiogenesis, invasion, and escape from senescence (25). However, melanoma does not develop in the parental species that naturally carry the oncogene *xmrk*, because the oncogenic activity is completely suppressed, and melanocytes remain in a nevus-like state. This clearly points to the presence of co-evolved suppressor mechanisms that restrain xmrk-driven tumorigenesis, warranting further investigation into these evolutionarily developed defenses for novel anti-melanoma strategies.

Despite these important discoveries, tumor heterogeneity in the *Xiphophorus* model remains understudied. Prior work has described tumor morphology (32) and bulk transcriptomic changes (33), but these approaches cannot resolve how diverse stromal, immune, and pigment cell populations interact to shape tumor ecosystems. Recent advances in single-cell and single-nucleus RNA sequencing (sc/snRNA-seq) have transformed our understanding of melanoma biology by revealing lineage hierarchies, dynamic immune states, and signaling networks that regulate tumor growth and immune evasion (11, 34, 35). Applying these approaches to *Xiphophorus* is essential not only to uncover the organization of its tumor microenvironment but also to provide a comparative framework for understanding conserved versus model-specific features of melanoma biology.

Accordingly, we performed snRNA-seq on spontaneous melanomas from *Xiphophorus* hybrids and resolved the cellular and transcriptional heterogeneity encompassed in *Xiphophorus* melanoma. By comparing to human melanoma, we identify conserved gene programs between human and *Xiphophorus* melanoma cancer cells and similar cellular composition.

## Results

### Global Heterogeneity in *Xiphophorus* Melanoma

To characterize the cellular heterogeneity of *Xiphophorus* melanoma, we performed single-nucleus RNA sequencing (snRNA-seq) on melanoma dissected from interspecies *Xiphophorus* hybrid exhibiting nodular growth of melanoma in the dorsal fin and penetration into musculatures. The dataset includes quality-filtered 5,839 sequenced nuclei. Clustering with Seurat identified eight distinct cell populations (Fig. 1).

**Figure 1.**
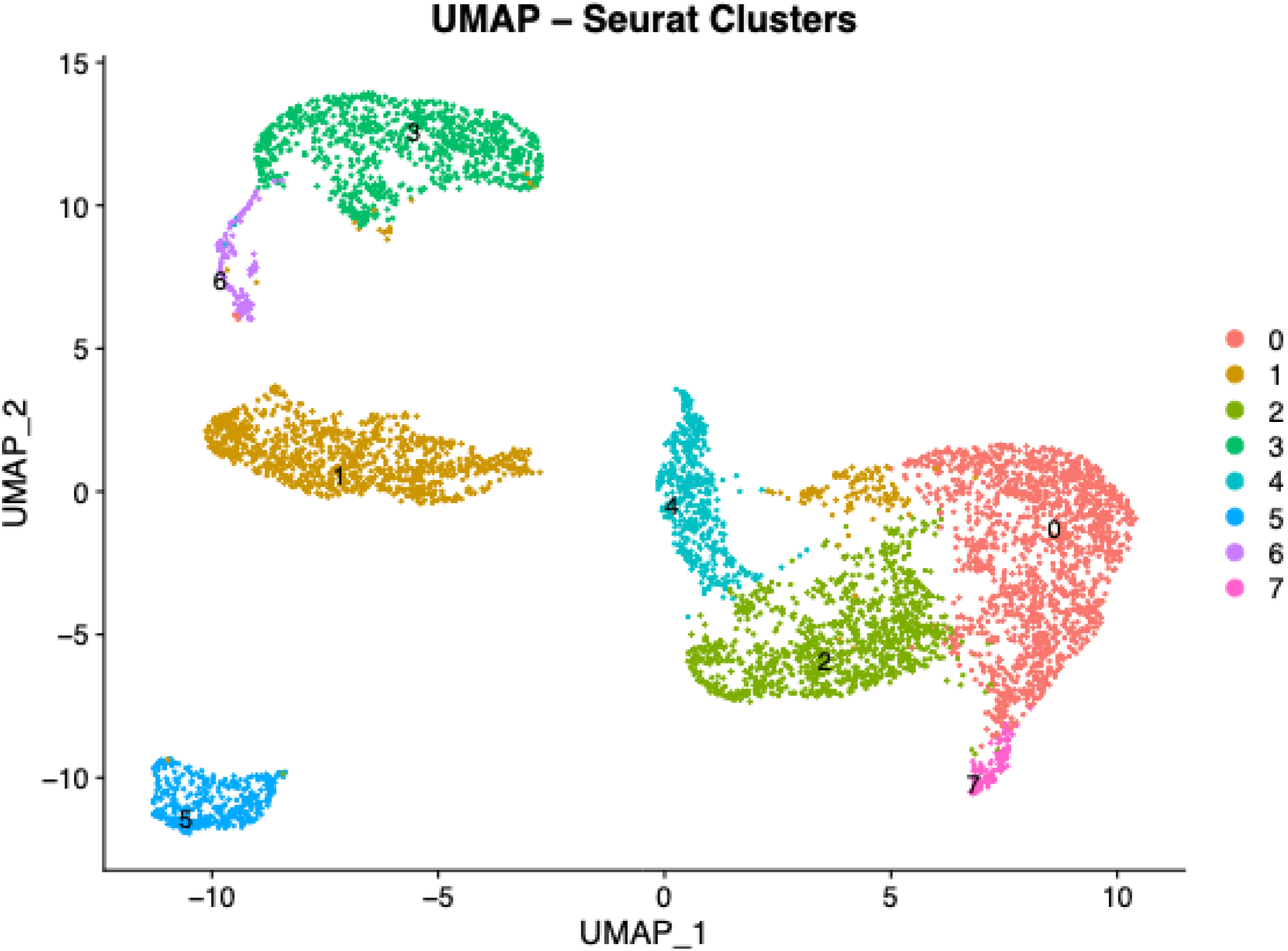
UMAP plot of *Xiphophorus* melanoma tumors by snRNA-seq. UMAP visualization of nuclei from *Xiphophorus* melanoma tumors after graph-based clustering, revealing eight transcriptionally distinct clusters. Each dot represents a single nucleus, colored by its assigned Seurat cluster.

Using a reference-based annotation approach, it is found that clusters 0 and 7 are enriched for epithelial populations; Clusters 1 contains vascular and mesenchymal cells; Clusters 2 and 4 represent immune compartments with differing lymphoid and myeloid compositions; Clusters 3 and 6 consist primarily of melanoma cancer cells, and Cluster 5 represent muscles cells (Supplemental Figure S4-S15, Supplemental Table S1-S3).

### Subtyping of cells in each cell population

Clusters 0 and 7 (Fig. 2A & 2H) collectively accounted for 28.7% of cells encompassed within the melanoma. Cells in cluster 0 are annotated as epidermal basal cells, periderm cells, and fin basal epidermal cells, and cluster 7 annotated as periderm cells. Although the term ‘periderm’ is linked to early zebrafish development in reference dataset, both clusters expressed hallmark epithelial markers (e.g., *grhl3, esrp1, tjp3* in Fig. S5), supporting their classification as localized epithelial lineages adjacent to the tumor.

**Figure 2.**
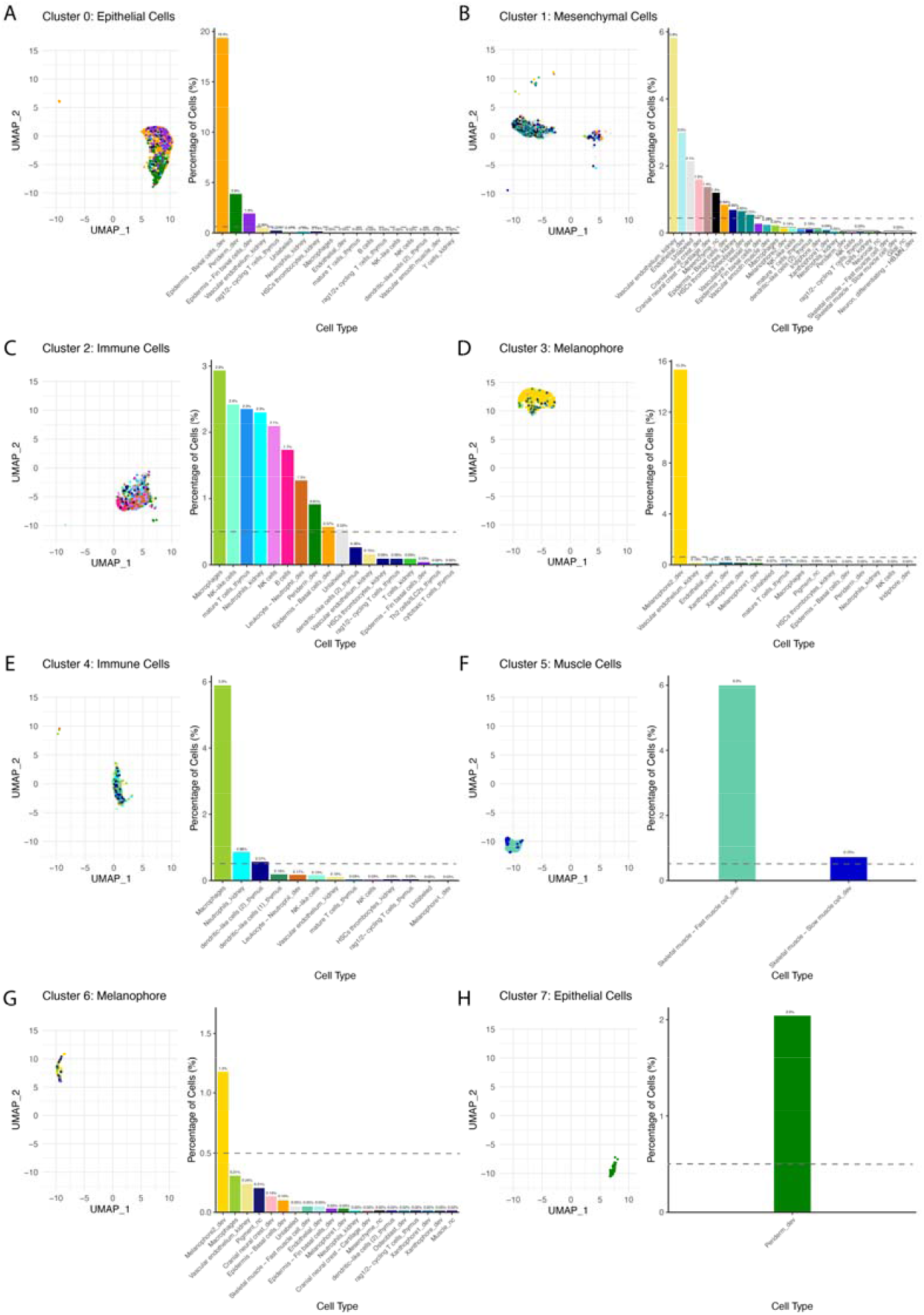
Cell type composition of each cluster based on lineage-specific marker enrichment. (A–H) UMAP projections (left) and bar plots (right) show the proportional representation of annotated cell types within each cluster. Bar plots display the percentage of all nuclei in the dataset, with a horizontal dashed line indicating the applied 0.5% cutoff for annotation. X-axis labels represent the annotated cell type followed by the reference dataset abbreviation (formatted as cell type_reference dataset, e.g., Melanophore2_dev), while the y-axis indicates percentage of nuclei.

Dominated by endothelial and mesenchymal lineages, Cluster 1 (Fig. 2B) represents stromal-rich compositions accounted for 19.5% of the total cell population. The most prominent cell types include vascular endothelium, developmental endothelial cells, and cranial neural crest-derived mesenchyme, such as cartilage precursors and osteoblast-like stromal cells, suggesting the presence of progenitor-like or undifferentiated cell populations. The presence of epidermal basal cells and hematopoietic progenitors (e.g., HSCs/thrombocyte cells) suggests a heterogeneous niche. These features define a perivascular stromal niche within the tumor microenvironment.

Clusters 2 and 4 (Fig. 2C & 2E) are enriched in immune related cells. Cluster 4 exhibited a relatively mono-faceted immune microenvironment, dominated by macrophages with minor contributions from neutrophils and thymus-derived dendritic-like cells. In contrast, Cluster 2 displayed a more complex immune microenvironment, encompassing not only myeloid elements (macrophage and neutrophils) but also a broader array of lymphoid populations, including NK and NK-like cells, B cells, and mature thymic T cells.

Clusters 3 and 6 (Fig. 2D & 2G) defined a cancer cell-dominant component, together accounting for 16.5% of the total cells.

Cluster 5 (Fig. 2F) contributed 6.7% of the total cell population and was composed exclusively of skeletal muscle cells. The majority were annotated as fast muscle cells (6.0%), accompanied by a smaller fraction of slow muscle cells (0.7%).

### Conservation of Human Melanoma Gene Programs in *Xiphophorus*

To evaluate gene expression programs (i.e., a suite of genes showing cell type-specific co-expression) relevant to cell types between human and *Xiphophorus* melanoma, we projected canonical human melanoma and tumor– microenvironment gene sets (11) onto the *Xiphophorus* melanoma dataset (Fig. 3). The melanoma-cell expression program is enriched in melanoma-dominant clusters (3 and 6), consistent with expression of melanoma genes *xmrk, mitfa, sox10, tyrp1b*, and *pmela* (Fig. 4A; Fig. S8). Stromal cluster 1 showed significant enrichment for endothelial and cancer associated fibroblast (CAF) gene signatures, reflecting a perivascular, mesenchymal niche like fibroblast- and endothelium-rich stromal compartments described in human melanoma. Immune programs localized to discrete clusters. Macrophage signatures are concentrated in cluster 4, matching its enrichment for *mrc1b, csf1rb (ENSXMAG00000006342), cd74a (ENSXMAG00000008646)*, and *card9 (ENSXMAG00000015148)* (Fig. S11). T cell programs are confined primarily to cluster 2, where markers such as *zap70* and *ikzf1* were expressed (Fig. S14), and B cell programs (*cd79a, pax5, pou2f2a, swap70a*) are detected in a restricted subset within the same cluster (Fig. S13). These immune programs mirror the structured lymphoid and myeloid organization reported in human melanoma single-cell datasets.

**Figure 3.**
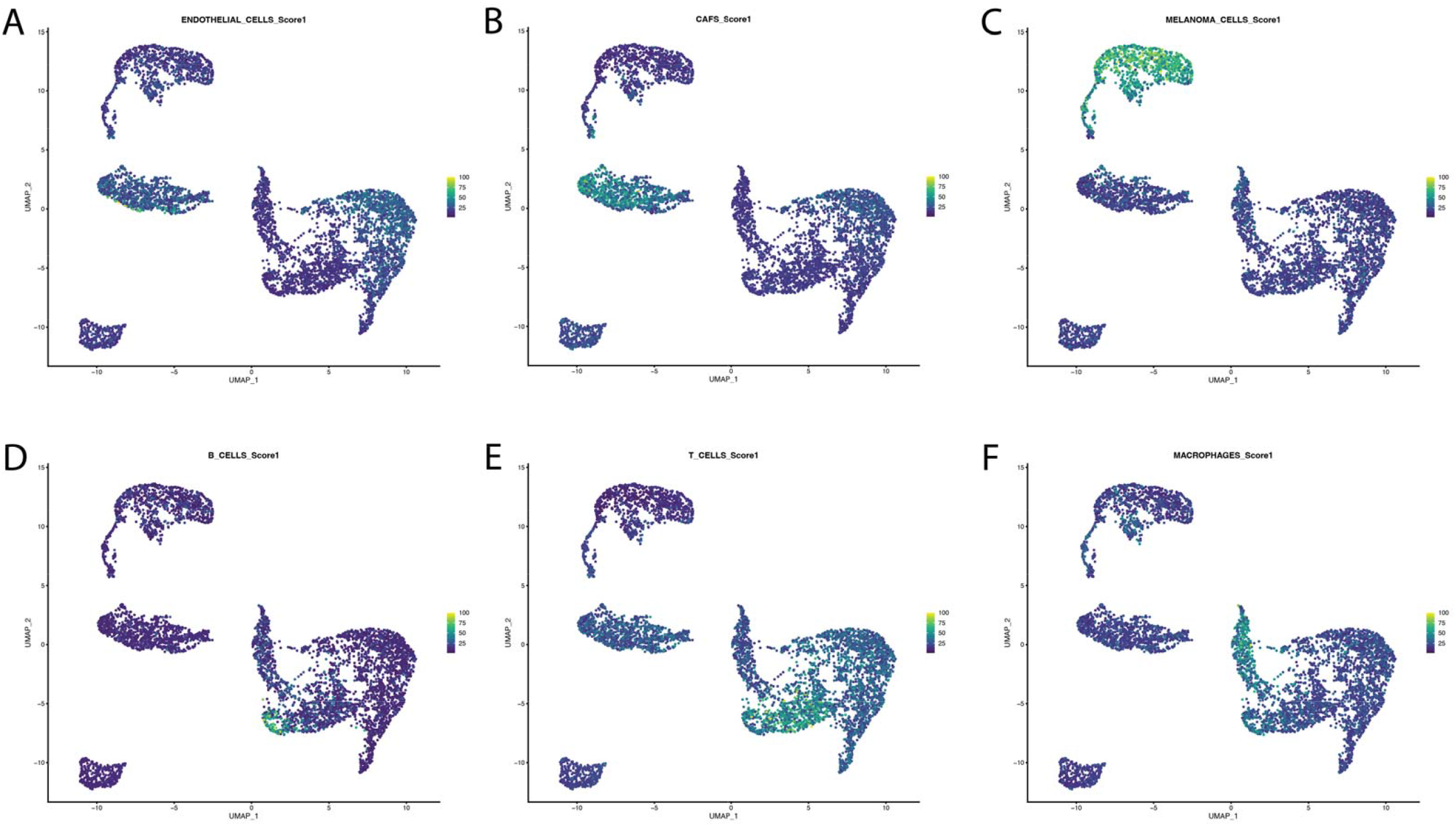
UMAP plots showing enrichment scores for canonical human cell-type programs projected onto *Xiphophorus* melanoma nuclei. (A) Endothelial cell program. (B) Cancer-associated fibroblast (CAF) program. (C) Melanoma cell program. (D) B-cell program. (E) T-cell program. (F) Macrophage program. Color scale represents projection scores (purple = low, yellow = high), indicating the relative degree of enrichment for each human program across *Xiphophorus* clusters.

**Figure 4.**
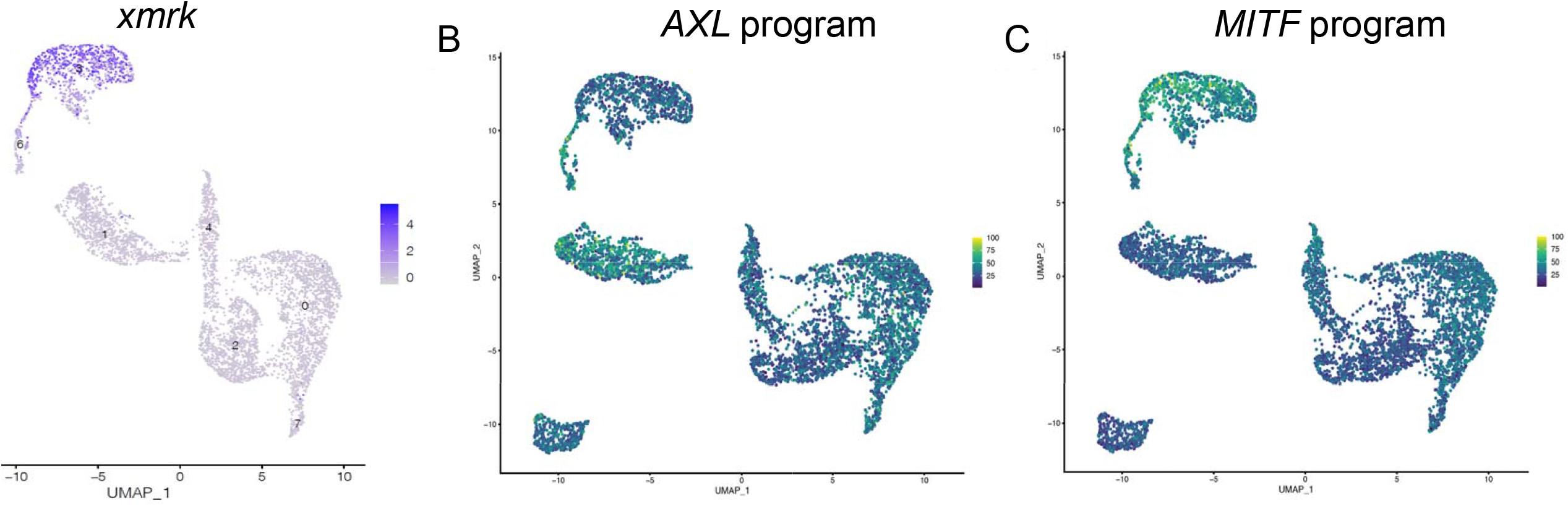
(A) Expression of *xmrk* across cell populations. (B) AXL- and (B) MITF-program module scores across the tumor UMAP. (B) UMAP colored by AXL program module score. AXL-high cells, indicating of a dedifferentiated, invasive/EMT-like state, are most enriched in cluster 1 (fibroblast) and are also detected in cluster 6 (pigment cells). (C) UMAP colored by MITF program module score. MITF-high cells, reflecting a differentiated, melanocytic/proliferative state, localize to pigment clusters 3 and 6. Module scores were computed in Seurat (AddModuleScore, RNA assay) from mapped human AXL/MITF gene sets.

Together, the cross-species cell mappings provide independent validation of the cluster annotations and demonstrate that key human melanoma and microenvironmental gene programs are conserved in *Xiphophorus* melanoma.

### Cancer cell heterogeneity

Within the two cancer cell clusters (cluster 3 and 6; Fig. 4), cells in both clusters show high module scores for MITF program genes but cluster 6 shows lower *mitfa* expression and lower MITF program gene module scores (**Fig. 4C**). In comparison, cluster 6 cells highly express AXL program genes (**Fig. 4B;** for each gene’s expression pattern, see [Supplemental Data; (11, 20)]. This matches the situation in human melanoma, namely that high AXL and high MITF cells co-exist within the same melanoma lesion.

Earlier studies have shown that *xmrk* expression drives melanoma initiation and disease progression by activating multiple signaling pathways that promote proliferation, migration, angiogenesis, and resistance to apoptosis (25). We identified a total of 720 *xmrk*-expressing cells (10.8% of all sequenced cells). These cells were predominantly found in melanoma clusters (706 cells; clusters 3 and 6), with smaller numbers distributed among mesenchymal (7 cells; cluster 1), epithelial (6 cells; cluster 0), and muscle (1 cell; cluster 5) clusters (Fig. 4A; Supplemental Fig. S16).

To evaluate the cancer cell dedifferentiation status, expression patterns of neural crest cell (NCC) lineage markers among the *xmrk*-expressing cells were assessed (Fig. 5; Supplemental Fig. S17). At the cluster level, both clusters 3 and 6 expressed melanophore marker genes and pigment cell progenitor markers including *mitfa, trpm1b, tfap2e, sox10*, and *mlpha* (Fig. 5A&B). In addition, both clusters express NCC markers and mesenchymal markers, with more mesenchymal markers expressed in cluster 6. A small fraction (<10%) of cluster 6 cells expressed Schwann cell markers such as *gldn, mbph*, and *egr2b*, which were absent in cluster 3 (Supplemental Fig. S17).

**Figure 5.**
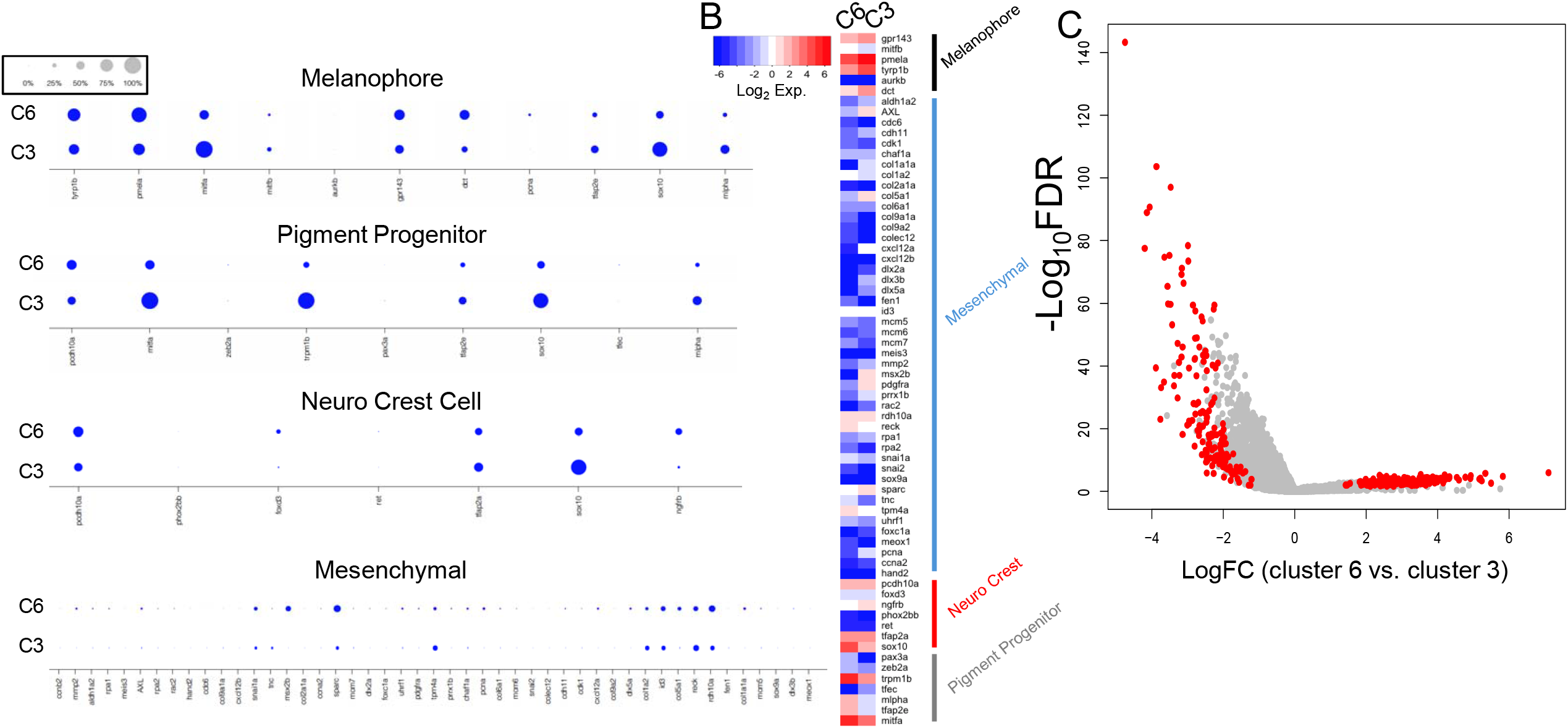
Comparative analysis of transcriptional programs between cluster 6 and cluster 3. (A) Dot plots showing expression of canonical melanophore, pigment progenitor, neural crest, and mesenchymal marker genes in clusters 6 (C6) and 3 (C3); dot size indicates the fraction of expressing cells and color intensity reflects expression level. (B) Heatmap of differentially expressed genes between clusters 6 and 3, annotated with representative lineage-associated markers. (C) Volcano plot highlighting genes significantly upregulated in cluster 3 relative to cluster 6 (red), with log fold-change (LogFC) on the x-axis and significance (–log_10_FDR) on the y-axis.

The analyses using NCC lineage markers do not indicate major differences in cellular states among the *xmrk*-expressing cells in cluster 3 and 6, suggesting their differences are beyond NCC lineages. Therefore, global gene expression differences were analyzed next to uncover cluster 3 and 6 variations.

Differential gene expression analysis identified 156 genes lowly expressed and 187 genes highly expressed in cluster 6 compared to cluster 3 (log_2_FC ≤ –1 or ≥ 1; FDR < 0.05; AUC ≥ 0.75; Fig. 5C). Pigment cell progenitor markers including *sox10, mitfa*, and *trpm1b* exhibited lower expression in cluster 6. Similarly, *xmrk* and its downstream signaling genes *fyn* and *stat5a* were also expressed at reduced levels in cluster 6 (see Supplemental Table S1 for the full list of DEGs). In contrast, cluster 6 showed elevated expression of the melanophore lineage regulator *ednrba*, several translation-associated genes such as *eif1, eif4aebp2, ENSXMAG00000023682*, and *eef1a2*. Importantly, 50 out of 87 ribosomal protein genes (*rpl* and *rps* family members) are also highly expressed in cluster 6 than cluster 3. Gene Ontology (GO) enrichment analysis confirmed a significant overrepresentation of terms related to ribosome biogenesis and translation among the differentially expressed genes between clusters 3 and 6. In addition, apoptosis-related genes are also enriched in the DEG list (Supplemental Fig. S18).

### Intercellular Communication in the Tumor Microenvironment

To characterize intercellular communication across the *Xiphophorus* melanoma tumor microenvironment, we applied CellChat (36) using a zebrafish ligand-receptor reference to infer statistically significant interactions between cell clusters. The resulting network revealed communication among all clusters (Fig. 5A, Supplemental Table S4). Inter-cluster communication with exclusively expressed ligand and receptor is observed between cancer cells (Cluster 3) and fibroblast cells (Cluster 1). The PDGF signaling pathway ligand *pdgfaa* (37) is notably expressed in cancer cell (Cluster 3) and epithelial (Cluster 0), and receptor *pdgfra* in fibroblast (Cluster 1) populations at the cluster level (Fig. 5B).

Additional signaling involved thbs1a-sdc4, with thbs1a expressed in melanophores and sdc4 in mesenchymal cells (38). Melanophores also communicated with epithelial and immune compartments, including nrg2b-erbb3a signaling to epithelial cells and NOTCH (39) signaling from epithelial and muscle clusters to melanophores. Intra-lineage signaling within pigment populations included ptn-sdc4 (40, 41) and col4a1-sdc4 (38) between Clusters 3 and 6. An EPHA signal (42) from muscle (Cluster 5) to mesenchymal cells (Cluster 1) was also detected (Supplemental Table S4).

The epithelial composition exhibited the most extensive signaling, with Clusters 0 and 7 engaging in 24 desmosome-mediated interactions. Additional epithelial-epithelial communication occurred through TIGIT, NECTIN, PVR, and JAM pathways (38, 43, 44). Epithelial-mesenchymal interactions involved similar pathways, including TIGIT, PVR, and NECTIN. Epithelial-immune communication included neuregulin (NRG) signaling (38), with nrg2a and nrg2b ligands expressed in epithelial and immune Clusters (0 and 2) and targeting erbb3a in epithelial Cluster 7 (Supplemental Table S4).

### Immune populations dynamics

To investigate dynamic changes in the immune compartment, pseudotime analysis was performed on Clusters 2 and 4 (Fig. 6A). The reconstructed trajectory revealed a continuous progression of immune states without clear branching, reflecting gradual transcriptional shifts within lymphoid and myeloid populations. Visualization of gene expression trends along pseudotime highlighted temporally ordered programs, with distinct sets of genes showing early, intermediate, or late expression patterns (Fig. 6B). These data provide a global view of immune transcriptional dynamics in Xiphophorus melanoma and serve as a framework for interpreting lineage-specific immune states in the tumor microenvironment.

**Figure 6.**
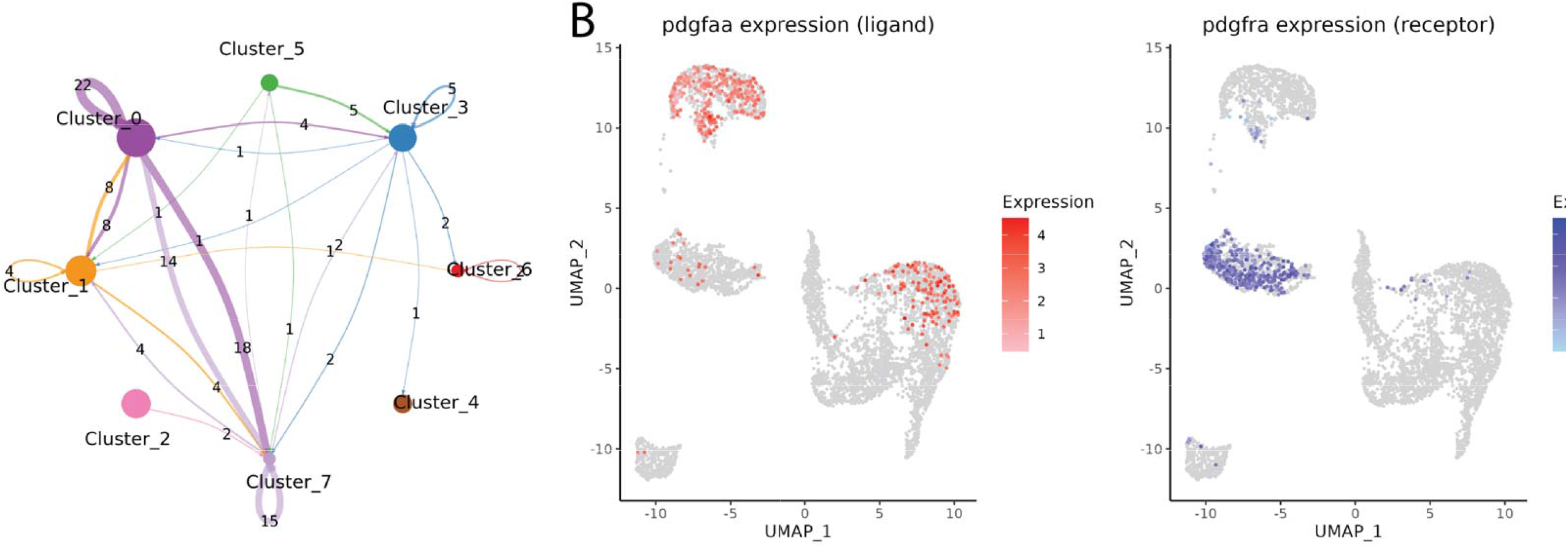
Ligand-receptor interactions between mesenchymal and melanophore clusters in the tumor microenvironment. (A) Network plot of predicted intra- and inter-cluster communication among the eight transcriptionally defined clusters using a zebrafish ligand-receptor reference. Nodes represent clusters, and edge width reflects the number of statistically significant ligand-receptor interactions. (B) UMAP feature plots showing expression of *pdgfaa* (ligand, left) and pdgfra (receptor, right). *pdgfaa* is expressed in Clusters 0 (epithelial), 1 (mesenchymal), and 3 (melanophore), while *pdgfra* is enriched in Cluster 1. These patterns suggest a signaling network involving both autocrine and paracrine PDGF signaling among epithelial, melanophore, and mesenchymal compartments.

**Figure 7.**
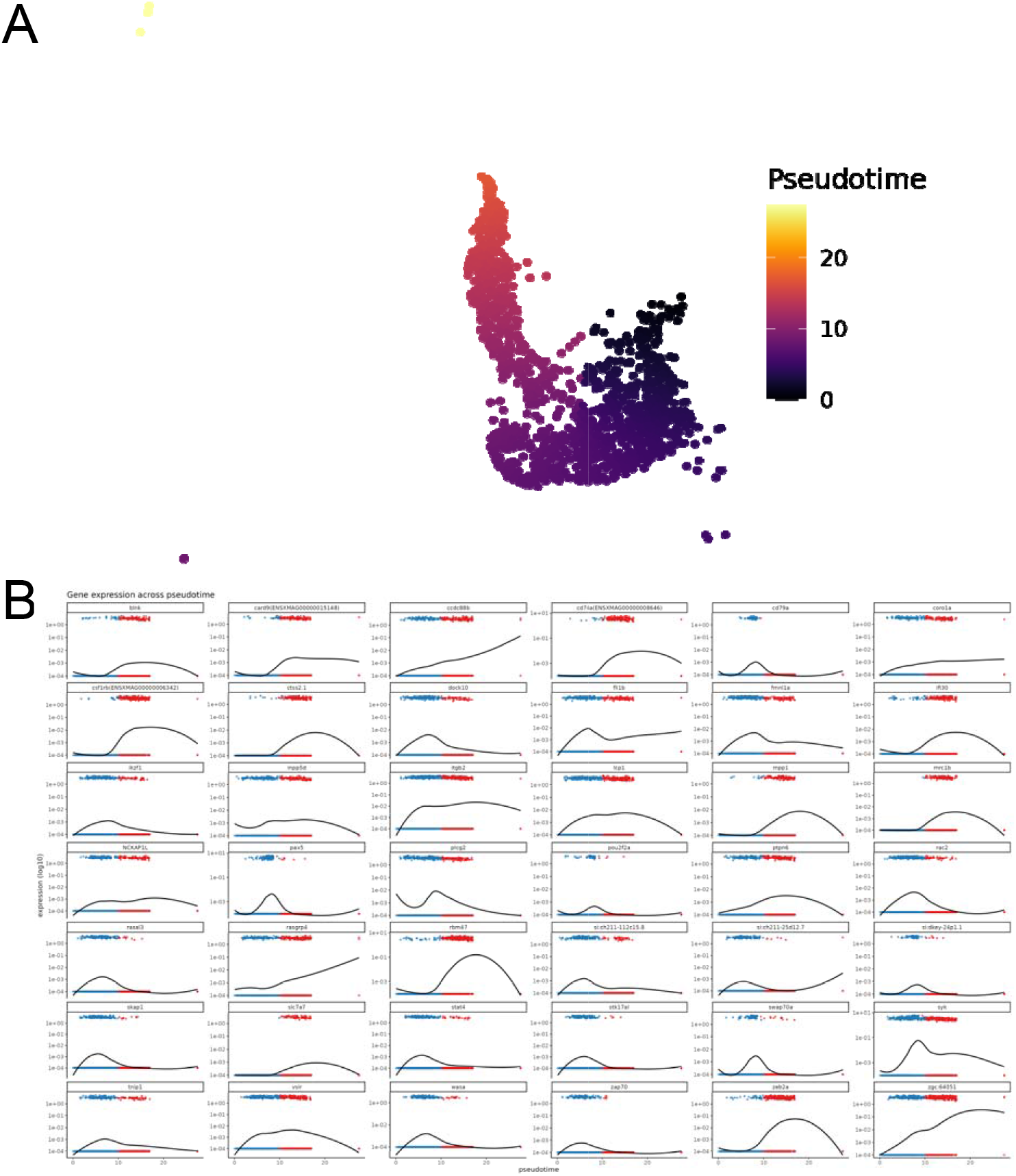
Pseudotime trajectory and dynamic gene expression of immune populations in *Xiphophorus* melanoma. (A) Pseudotime trajectory analysis of immune clusters (Clusters 2 and 4) reveals a continuous progression of cellular states. Cells are ordered along pseudotime (color gradient, purple to yellow), reflecting transcriptional transitions across immune subsets. (B) Expression dynamics of representative immune-related genes plotted across pseudotime. Each panel shows smoothed expression trends (black line) with single-cell values overlaid (blue and red dots corresponding to distinct branches of the trajectory).

## Discussion

This study presents a *Xiphophorus* melanoma cell atlas, and offers insights into the cellular heterogeneity, immune landscape, and intercellular communication networks of a type of naturally occurred melanoma. By implementing cross-species projection of gene programs hallmarking diverged cells types identified in human melanoma to *Xiphophorus* melanoma, we found that *Xiphophorus* melanoma include all key cell types of human melanoma, with gene expression closely resembled human advance melanoma.

The dominant cell populations expressing *xmrk* are derived from pigment cell lineage. This is consistent with previous studies relying on bulk RNAseq (Fig. 4) (33, 45). Co-expression of *xmrk* with neural crest cell (NCC) markers supports the dedifferentiated status of these cells (Fig. 5). While the expression of NCC markers such as *ngfrb*’s human ortholog *NGFR* (Fig. 5) hallmarks more aggressive melanoma (29358574). However, the two melanoma cell types (i.e., clusters 3 and 6) exhibit similar dedifferentiation status based on similar NCC lineage markers, suggesting they are similar in terms of dedifferentiation status.

One most notable finding of this study is the finding of cluster 6 melanoma cells are featured with elevated expression of genes involved in protein translation, including translation initiation and elongation factors, as well as a wide array of ribosomal subunits. This expression pattern suggests a global upregulation of ribosome biogenesis and translational activity. Concurrently, *xmrk* and several of its downstream effectors are expressed at lower levels in these cells. Although melanoma does not originate from epithelial cells, its phenotypic plasticity, i.e., switching between a proliferative state and an invasive mesenchymal-like state, resembles the epithelial-to-mesenchymal transition (EMT). Increased ribosomal biogenesis has been reported to fuel EMT (46, 47), and targeting ribosomal biogenesis has been shown to be a promising therapeutic strategy to impair phenotype switching in cancers such as breast cancer (47) and prostate cancer (48), such transitory cells have not been reported in melanoma (49, 50). Thus, cluster 6 cells may represent a newly identified state en route to an invasive phenotype.

The presence of developmental and progenitor-like mesenchymal and endothelial populations in Cluster 1 suggests neo-angiogenesis, a hallmark of tumor-associated stroma also reported in human melanoma (Fig. 3A-B) (11). Notably, melanophores in Cluster 3 exhibit significant interactions with mesenchymal stromal cells, mediated predominantly through PDGFR signaling (Fig. 4B). Such interactions closely parallel human melanoma, where PDGF-driven signaling facilitates tumor-promoting processes such as fibroblast activation, extracellular matrix (ECM) remodeling, and angiogenesis (51, 52). In human studies, PDGFRA and its ligand PDGFA have been implicated in tumor progression, therapeutic resistance, and the formation of desmoplastic stroma, highlighting their potential as therapeutic targets (53). In this context, the mostly exclusive distribution of *pdgfaa* in melanophores and *pdgfra* in stromal cells shows a conserved paracrine signaling in *Xiphophorus* melanoma, potentially fostering a microenvironment supporting tumor growth and structural remodeling.

Immune populations in Clusters 2 and 4 mirrored the compartmentalization of lymphoid and myeloid programs described in human melanoma (Fig. 3D-F) (11). Cluster 4 macrophages expressed M2-like, tumor-associated macrophage (TAM) markers *mrc1b (54), csf1rb (55), cd74a (56)*, and *card9 (57)*. Such macrophages are known to promote angiogenesis, extracellular matrix remodeling, and immunosuppression in patients (54-57).

The expression of *arhgap4b* was enriched in immune populations (Fig. S1). As a Rho GTPase-activating protein implicated in cytoskeletal remodeling, arhgap4b likely facilitates immune cell migration and positioning within the tumor microenvironment. Recent pan-cancer analyses have also linked *ARHGAP4* (58) to differential immune infiltration and poor prognosis in several human cancers, including pancreatic, liver, colorectal, lung and prostate cancer (58-62). However, it hasn’t been reported to be involved in melanoma progression. Underscoring its potential role in modulating immune-tumor interactions and contributing to immune surveillance or evasion dynamics.

In summary, the *Xiphophorus* melanoma model recapitulates key features of human melanoma, including structured cellular heterogeneity, immune polarization, and stromal remodeling. These findings highlight conserved mechanisms of tumor–host interaction and position *Xiphophorus* as a powerful comparative model for dissecting melanoma biology and for exploring translational strategies to modulate the tumor microenvironment.

## Conclusion

We deconvoluted the cellular landscape of *Xiphophorus* melanoma, identifying a spectrum of melanoma cell states ranging from proliferative cells to transient, non-proliferative intermediates, to potentially invasive mesenchymal-like and niche-adapted cells. These findings deepen our understanding of melanoma cell plasticity and underscore the utility of this evolutionary model for informing translational cancer biology.

## Materials and Methods

### Animal Model

*X. maculatus* (Jp163A), *X. hellerii* (*Sarabia*), and first-generation backcross (BC1) interspecies hybrids used in this study were supplied by the Institute for Molecular Life Sciences. *X. maculatus* female fish were artificially inseminated with sperm from male *X. hellerii* to produce F1 interspecies hybrids. F1 hybrid males were then backcrossed to *X. hellerii* females to generate the BC1 animals with malignant melanoma formation. At dissection, all fish were euthanized by decapitation. Upon loss of gill movement animals were decapitated. Tumors were snap-frozen using ethanol dry ice bath. All BC_1_ fish were kept, and samples taken in accordance with protocols approved by Texas State University Institutional Animal Care and Use Committee (IACUC 9048).

### Nuclei isolation

Nuclei were isolated from melanoma using the Singulator2 instrument per manufacturers protocols (S2 Genomics; Livermore, CA). After mechanical disruption, cell filter straining steps, and washes, the cell nuclei were suspended in PBS/0.1% BSA or for thymus a nuclei isolation buffer (S2 Genomics; Livermore, CA), with all buffers containing 0.4U/ul RNAase inhibitor, prior to microfluidic encapsulation on the 10X Chromium instrument (10x Genomics®, Pleasanton, CA) to nanoliter-scale.

### Single-nuclei library preparation

Single-nuclei libraries were generated using the GemCode Single-Cell Instrument and Single Cell 3′ Library and Gel Bead Kit v3 and Chip Kit (10x Genomics®, Pleasanton, CA) according to the manufacturer’s protocol. Before sequencing, every library was analyzed on a Bioanalyzer high sensitivity chip to ensure the expected cDNA fragment size distribution was achieved. The appropriate number of individually barcoded GEM libraries were pooled and sequenced on a NovaSeq 6000 instrument (Illumina) with 2×150bp length using these sequencing parameters: 26 bp read 1 – 8 bp index 1 (i7) – 98 bp read 2 with 200 cycles.

The Cell Ranger software pipeline (version 3.1.0) was used to demultiplex cellular barcodes, map reads to the *X. maculatus* reference genome (GCA_002775205.2) and transcriptome using the STAR aligner, and down-sample reads required to generate normalized aggregate data across samples, producing a matrix of gene counts versus cells. The individual tissue-specific sequenced Gel Bead-In Emulsion (GEM) libraries were each processed with the CellRanger v7.0.1 pipeline (10X Genomics) to create a cellular barcode by genomic feature matrix.

### Cell clustering

The Cell Ranger software pipeline (version 3.1.0) was used to demultiplex cellular barcodes, map reads to the *X. maculatus* reference genome (GCA_002775205.2) and transcriptome using the STAR aligner, and down-sample reads required to generate normalized aggregate data across samples, producing a matrix of gene counts versus cells. The individual tissue-specific sequenced Gel Bead-In Emulsion (GEM) libraries were each processed with the CellRanger v7.0.1 pipeline (10X Genomics) to create a cellular barcode by genomic feature matrix. The resulting matrices were imported into R for downstream analysis using the Seurat package (v4.4.0) (63). Expression data were normalized using the NormalizeData method, followed by identification of highly variable genes with the FindVariableFeatures function. The data were scaled and subjected to linear dimensional reduction through principal component analysis (PCA) (Fig. S2), selecting significant principal components (PCs) based on the elbow plot visualization (Fig. S3). Subsequent clustering was performed using the Louvain algorithm implemented by the FindNeighbors and FindClusters functions, with cluster resolution optimized to balance granularity and biological interpretability. Visualization and assessment of clusters were performed using Uniform Manifold Approximation and Projection (UMAP).

### Cell type annotation

For cell-type annotation, *Xiphophorus* genes were first mapped to their zebrafish orthologs using Ensembl BioMart (Ensembl Release 109, October 2022 archive), followed by a hypergeometric enrichment test against curated gene marker sets compiled from four zebrafish resources: developmental atlas (64), neural crest (65), thymus (66), and kidney (67) (Supplemental Table S2). This workflow is similar to BLUEPRINT annotation framework

(68) but without gene expression ranking. Multiple testing correction was applied using the Benjamini-Hochberg false discovery rate (FDR). Cells were assigned to a cell type only if they exhibited statistically significant enrichment with an FDR-adjusted p-value ≤ 0.05; among significant hits, the cell type with the lowest adjusted p-value was selected. Cells not meeting this threshold were classified as Unlabeled. Marker genes were filtered under multiple parameters sets using combinations of expression specificity thresholds. Among these, the threshold combination of pct.1 ≥ 0.2, pct.2 ≤ 0.2, and logFC ≥ 0.5 was selected for final cell-type annotation (Supplemental Table S3). To minimize potential noise and focus on biologically meaningful populations, we excluded annotated cell types representing less than 0.5% of total nuclei from downstream analyses. Cell type labels were subsequently aggregated at cluster level to define the characteristic cellular composition of each cluster (Fig. 2).

### Xmrk-expressing cell identification and cellular status analyses

Nuclei expressing feature ENSMAG00000024953 (*xmrk*), determined by normalized reads > 0, are identified for analyses. We collected neural crest cell (NCC) lineage markers from zebrafish reference dataset (64) and manually curated reported gene markers representing different cell types and sub-cell types into reference gene markers hallmarking major lineages, i.e., NCC, neural, mesenchyme, muscle, pigment progenitors and melanophores. For each *xmrk-*expressing single nucleus, binary expression data (i.e., expressed = 1, not expressed = 0) was made from normalized expression matrix of NCC lineage marker genes. PCA and pseudotime analyses were conducted using R package SCORPIUS. NCC marker genes co-expression analyses was conducted, with hierarchical clustering performed and dendrograms for cells and genes plotted using R package ggplots. Genes contributing to the cell separation along PC1 were identified using rotation function of R prcomp.

### Differential gene expression analyses between pigment cell sub-clusters

The *xmrk-*expressing cells belonging to melanophore cell clusters 3 and 6 were extracted from normalized gene expression matrix and were compared for differential gene expression using t-test. The p-values were corrected for multiple test correction using False Discovery Rate (FDR). Differentially expressed genes were determined by |log2 [Cluster 3 expression/Cluster 6 expression]|>1, FDR<0.05, and Area Under the Receiver Operating Characteristic Curve (AUC) > 0.75.

### Comparative analyses between *Xiphophorus* and human melanoma

To evaluate cross-species conservation of tumor programs, we projected canonical human melanoma and tumor-microenvironment gene sets (11) onto the Xiphophorus snRNA-seq dataset. Human marker genes were mapped to *Xiphophorus* orthologs using Ensembl BioMart (October 2022 archive), and only markers with successful orthology were retained. For each program, Seurat’s AddModuleScore was applied to compute per-cell module scores, requiring at least five mapped genes per set. Module scores were then visualized on the UMAP embedding to assess how human melanoma and microenvironmental programs aligned with annotated Xiphophorus clusters (Fig. 3).

The human MITF-program and AXL-program genes were identified in literatures (11). The *Xiphophorus* orthologs for these genes are identified using R package Biomart in Ensembl genome database. Scores are computed in Seurat (AddModuleScore, RNA assay) from mapped human AXL and MITF program genes in the *Xiphophorus* snRNAseq data.

### Inter-cluster communication analysis

Intercellular communication analysis was conducted using the CellChat package (v1.6.0), employing a zebrafish ligand-receptor interaction reference database. Statistical inference of ligand-receptor interactions between cell types was performed using a bootstrapping approach (10,000 iterations), and interactions were retained only if both the sending and receiving cell groups contained at least 10 cells. These criteria defined the confidence thresholds used to infer robust communication events.

### Gene Ontology

Differentially expressed genes between the two melanoma clusters (cluster 3 and 6) were first mapped to human orthologs using R package Biomart in Ensembl genome database. Gene ontology (GO) term over-enrichment analysis is conducted using web tool g:Profiler (69). Adjusted p-value < 0.05 was used as statistical threshold to determine GO term over-enrichment.

## Supplemental Figures

**Supplemental Figure S1. UMAP visualization of single-nucleus RNA-seq data showing the distribution of arhgap4b expression across all clusters**.

**Supplemental Figure S2. Principal component analysis (PCA) of *Xiphophorus* melanoma snRNA-seq dataset**. PCA was performed on the scaled and normalized dataset to reduce dimensionality. The scatter plot shows the distribution of cells along the top principal components, which capture the major sources of transcriptional variance across the dataset.

**Supplemental Figure S3. Elbow plot for principal component selection**. Variance explained by each principal component (PC) is plotted. The “elbow” at which additional PCs contribute only marginally to variance was used to guide the selection of significant PCs for downstream clustering and UMAP visualization.

**Supplemental Figure S4-S15. Gene markers for Basal Cells, Periderm Cells, Vascular Endothelium, Endothelial Cells, Melanophore, Skeletal Muscle, Macrophages, Neutrophils, B Cells, T Cells, Mesenchyme**.

**Supplemental Figure S16. Normalized *xmrk* expression across clusters**. Scatter plot showing normalized xmrk expression values in individual cells grouped by cluster identity. Each dot represents a single cell. The y-axis shows normalized xmrk expression, and the x-axis denotes cluster assignments (0–6). Colors correspond to cluster identity, highlighting that xmrk expression is strongly enriched in cluster 3, with moderate expression detected in clusters 0, 1, 5, and 6.

**Supplemental Figure S17. Dot plot of cell-type marker gene expression in clusters C3 and C6**. Dot plots show the scaled expression and percentage of cells expressing marker genes for selected cell types across clusters C3 and C6. Each panel represents a different reference cell type, including otic epithelium, fin bud, muscle, Schwann cells, iridophore, xanthophore, pigmented muscle, and neural lineages. The size of each dot corresponds to the fraction of cells within the cluster expressing the marker gene (0–100%).

**Supplemental Figure S18. Gene Ontology analyses of inter-cancer clusters differentially expressed genes**. Gene Ontology (GO) analyses are conducted using human homolog of differentially expressed *Xiphophorus* genes. Statistically significant over-represented GO terms are shown, with bargraph height represented - log10(adjusted_p_value).

## Supplemental Tables

Supplemental Table S1 Cluster Marker List

Supplemental Table S2 Reference Marker Table

Supplemental Table S3 Cell Annotation

Supplemental Table S4 Ligand receptor interactions

## Supplemental Data

UMAP of AXL program orthologs expression (UMAP_AXL_orthologs_expressed.pdf) UMAP of MITF program orthologs expression (UMAP_MITF_orthologs_expressed.pdf)

## Acknowledgements

This work was supported by the National Institutes of Health, National Cancer Institute, R15 CA-223964 to Y. Lu, Office of Director R24 OD-031467 to Y. Lu, W. Warren and M. Schartl, R2R1 accelerator award from Texas State University to Y. Lu and M. Schartl.

