## Supplementary material for "Comparison of Human Melanoma Single Cell Profiles to Evolutionary Medicine Model *Xiphophorus* Provides Insights in Disease Control": Supp. files: Supp.Figure.pptx

### Slide 1
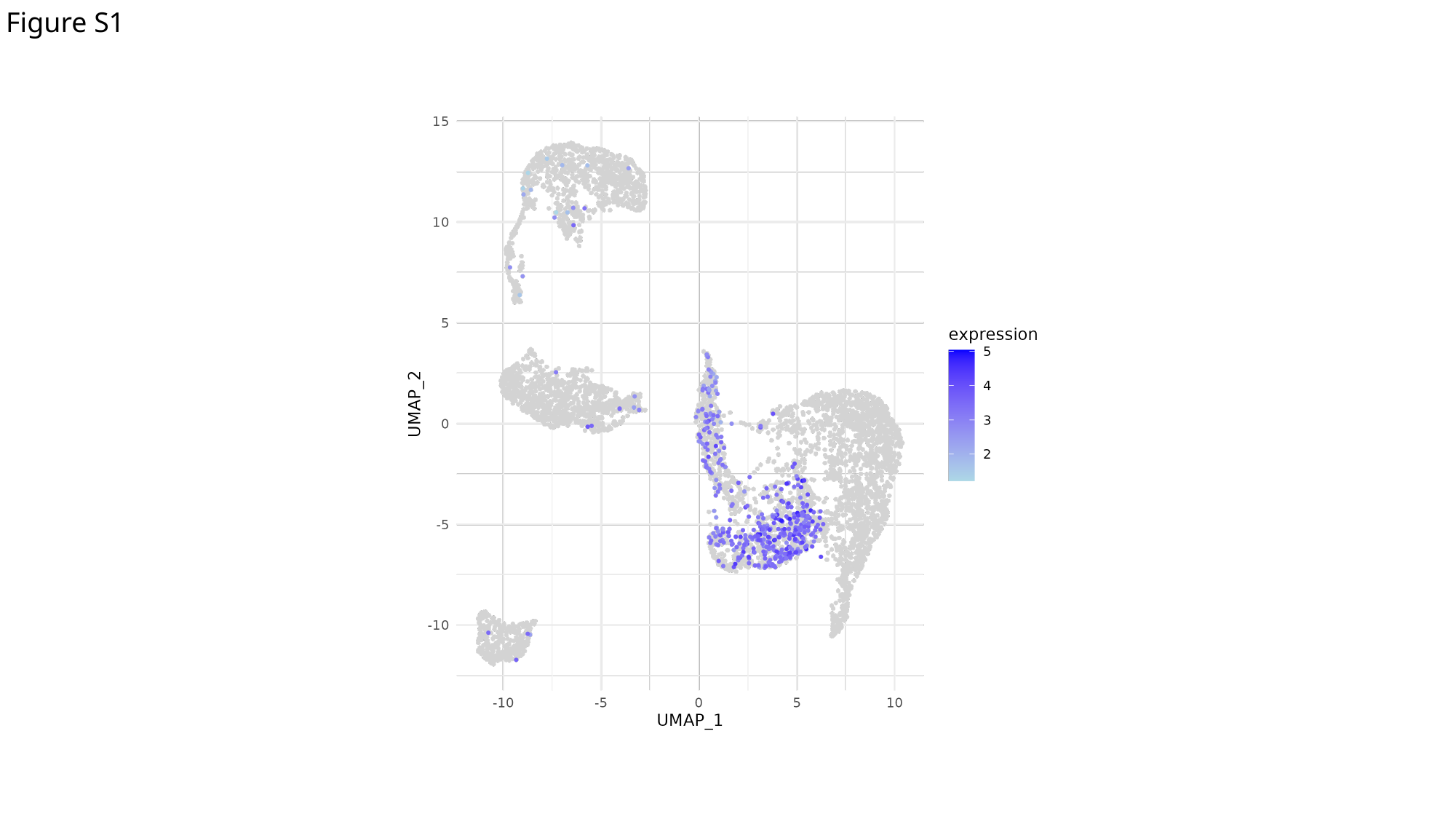

Figure S1

### Slide 2
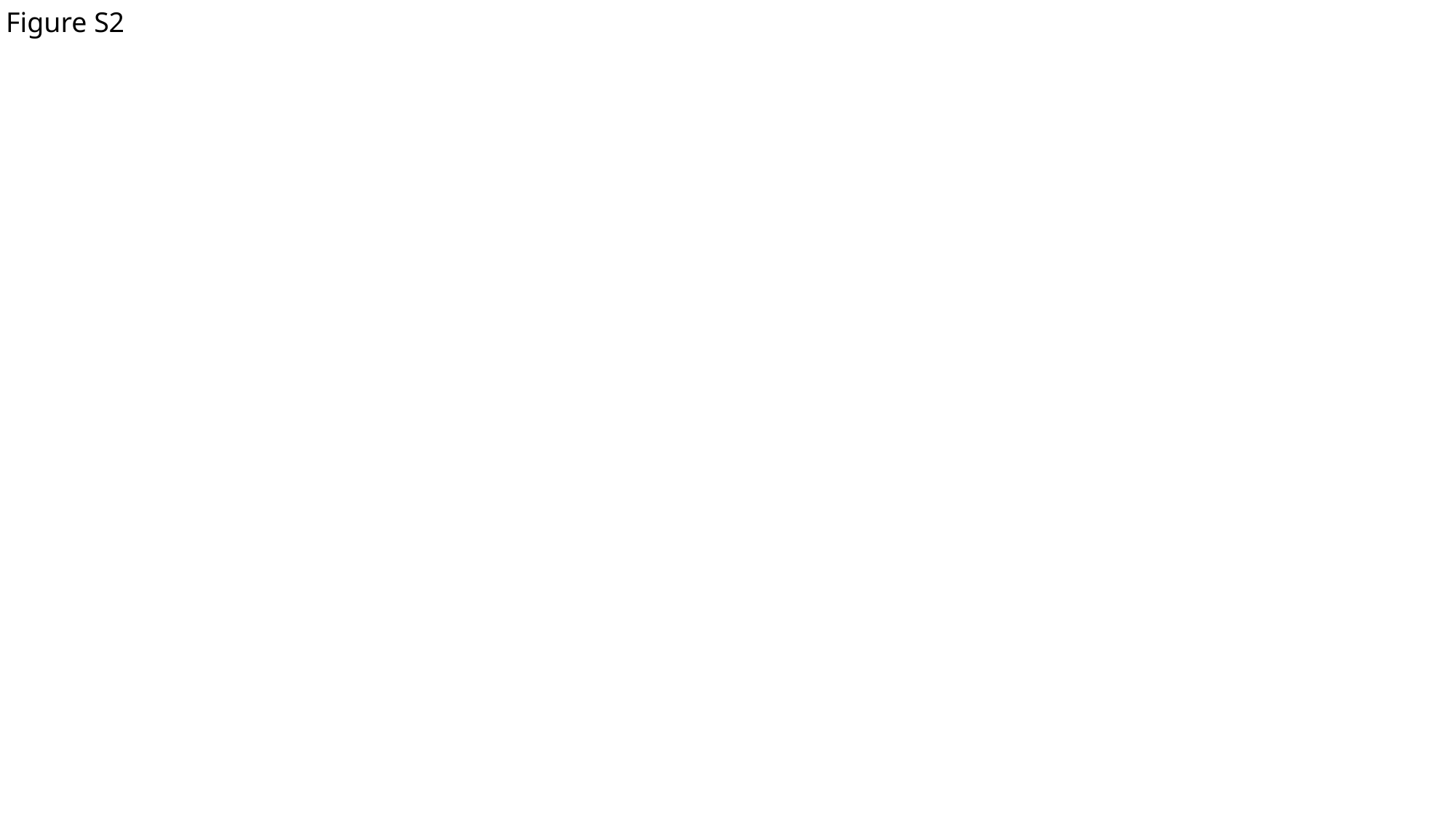

Figure S2

### Slide 3
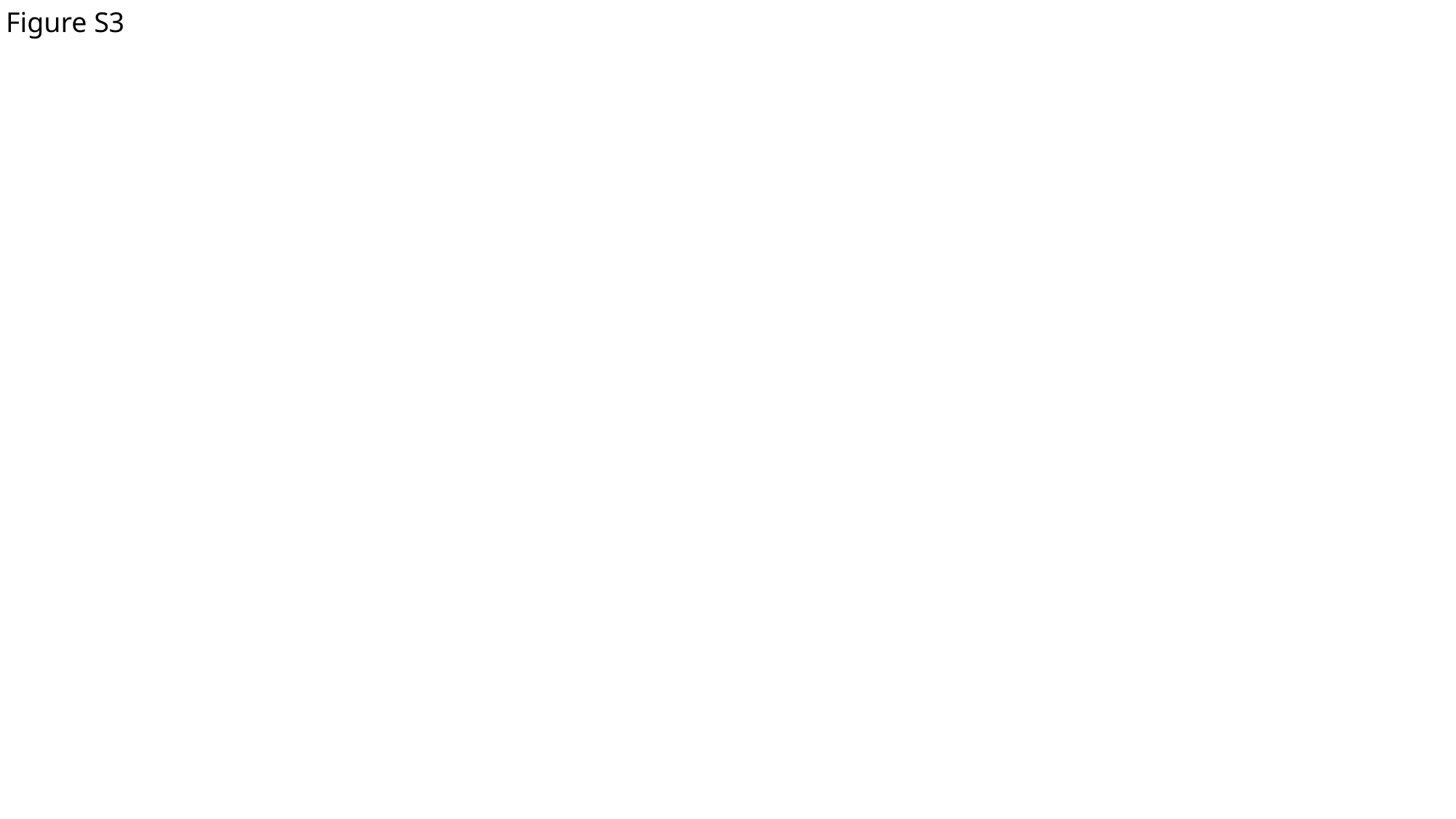

Figure S3

### Slide 4
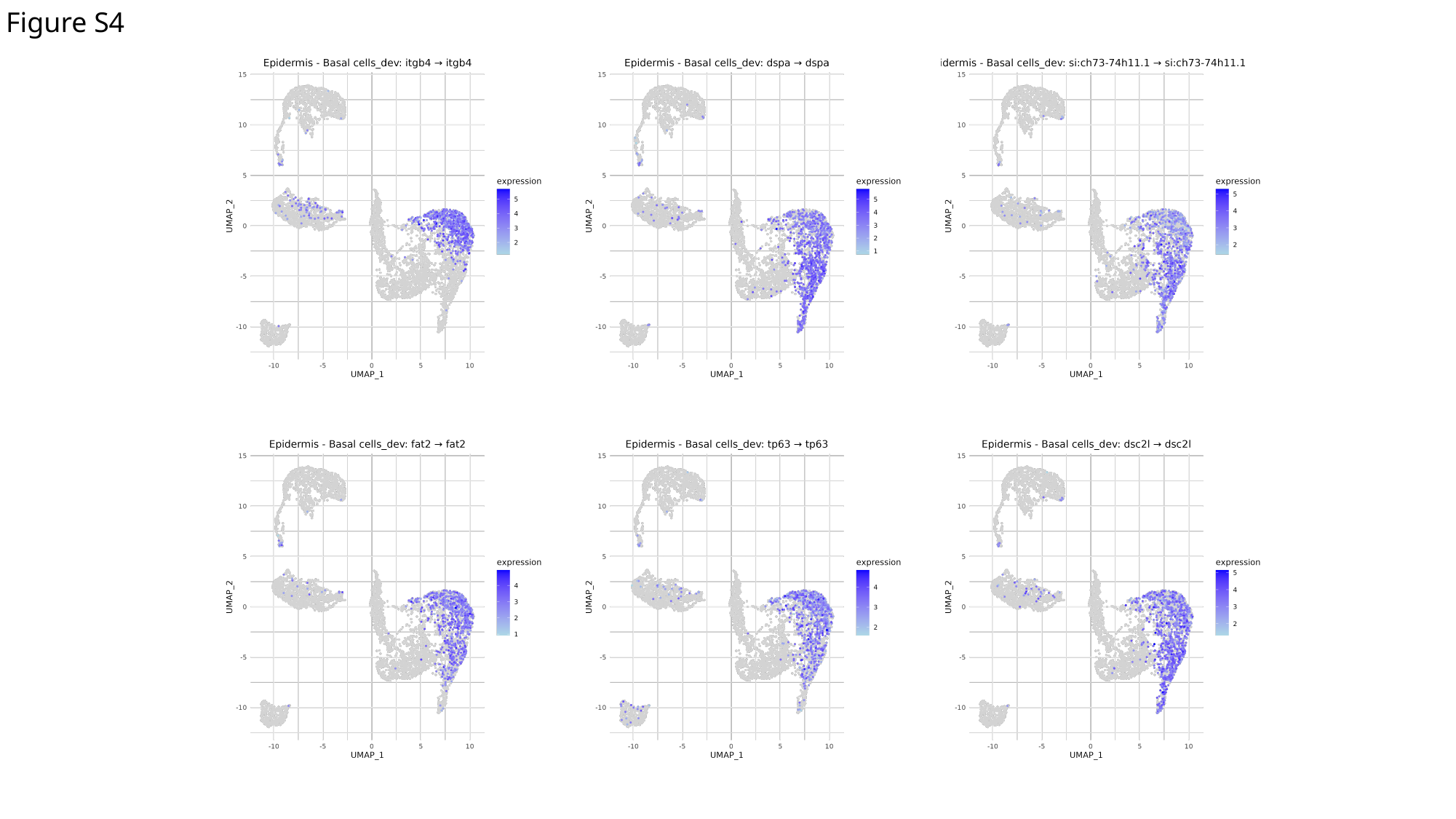

Figure S4

### Slide 5
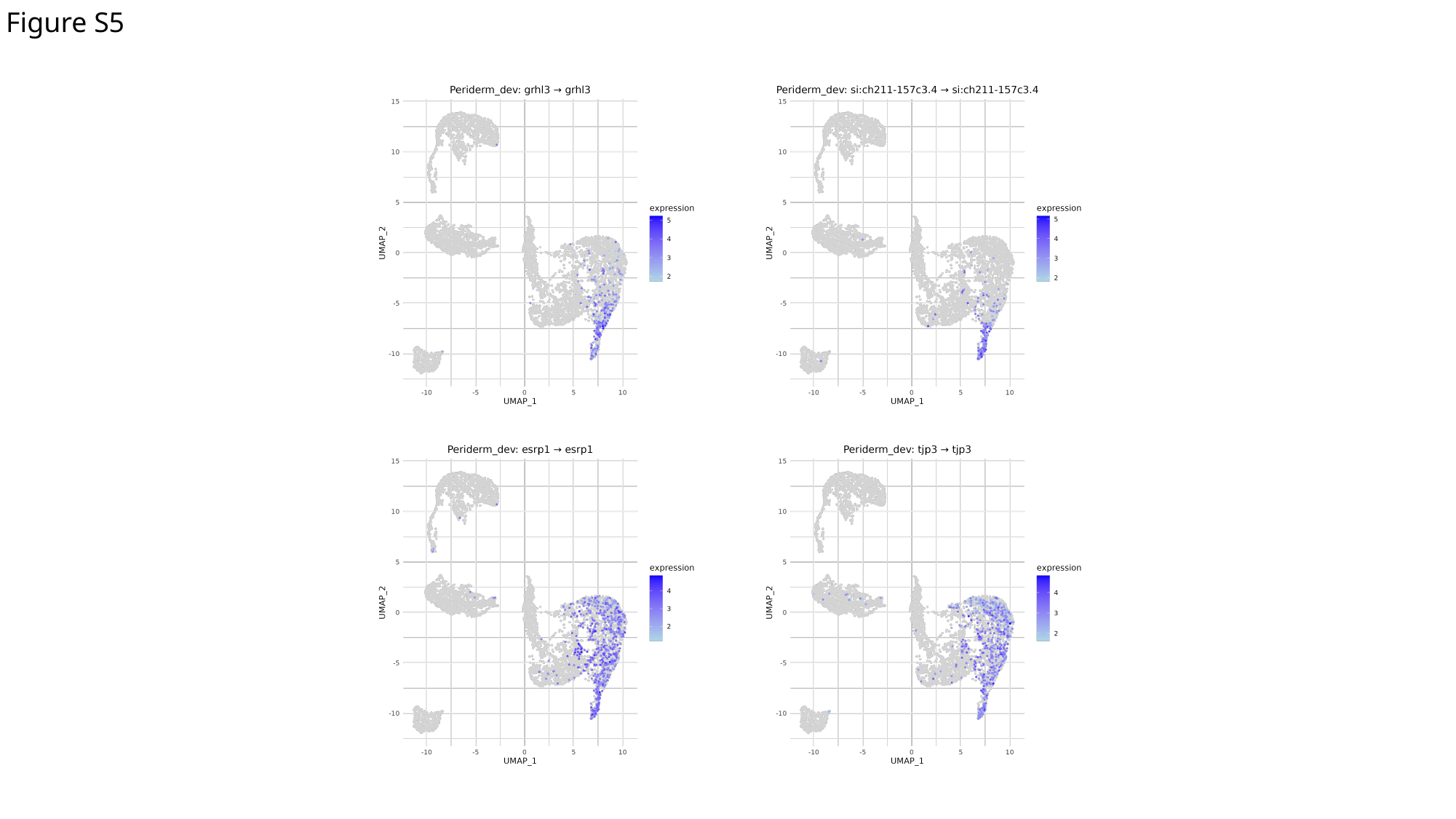

Figure S5

### Slide 6
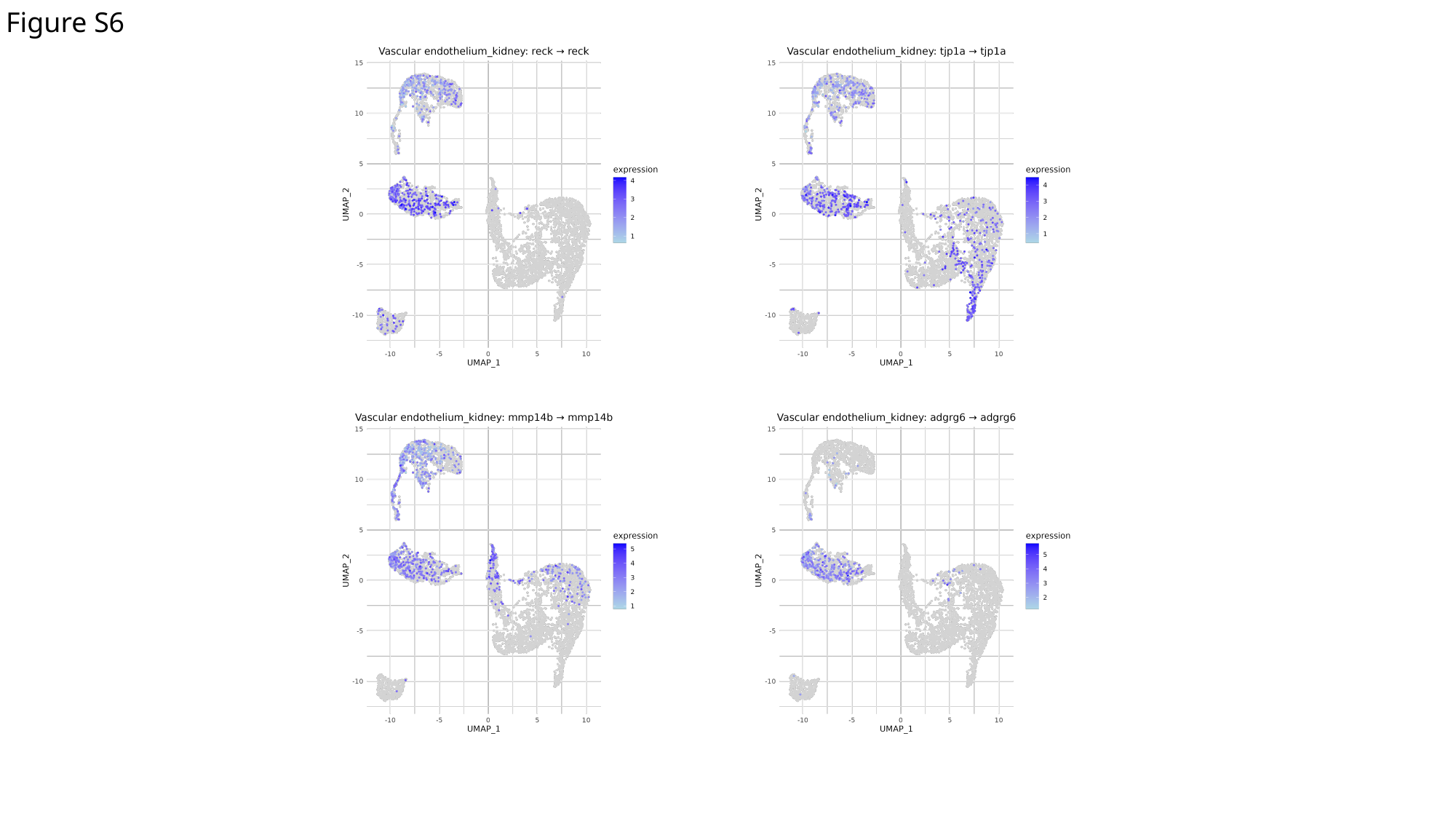

Figure S6

### Slide 7
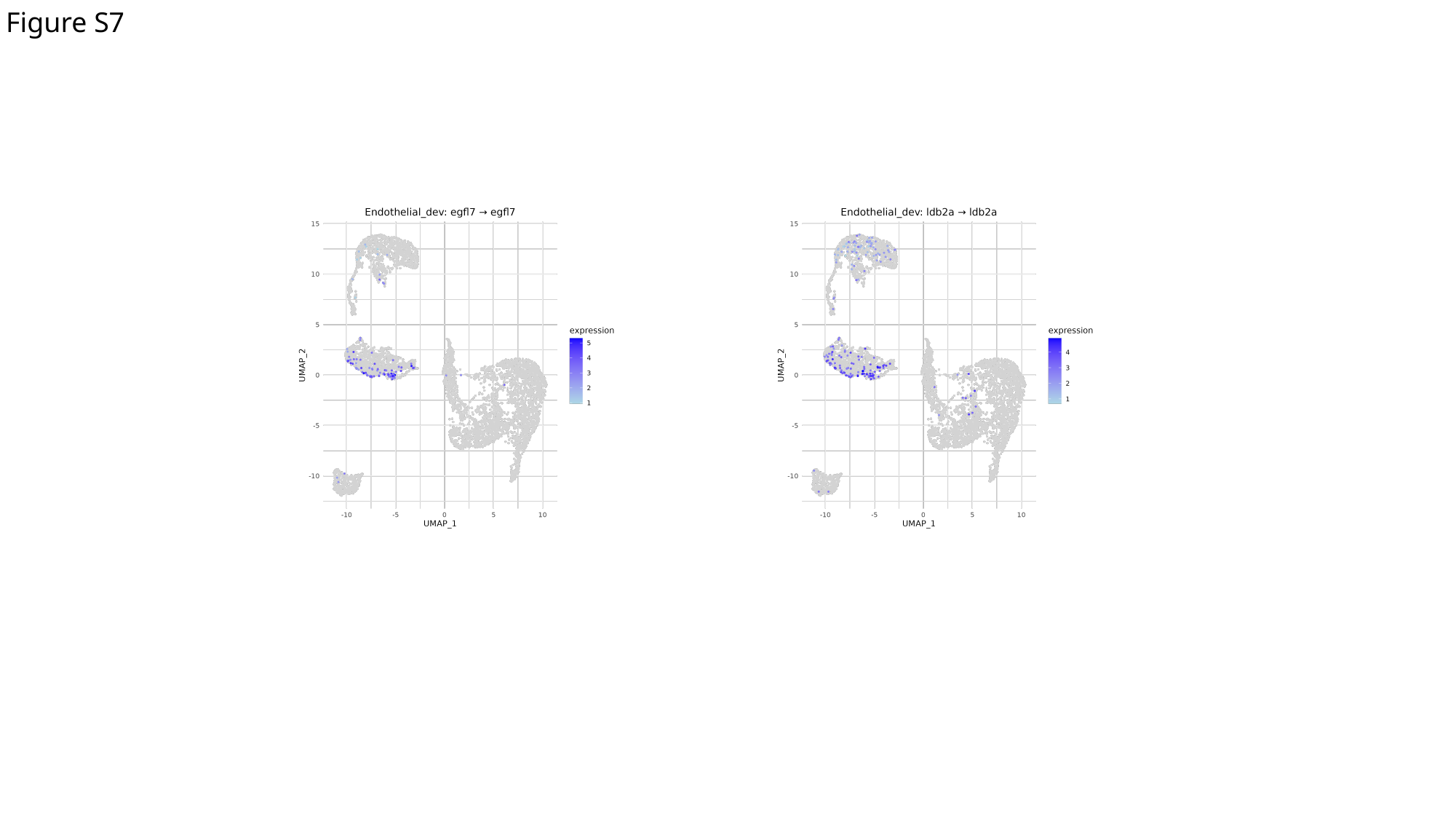

Figure S7

### Slide 8
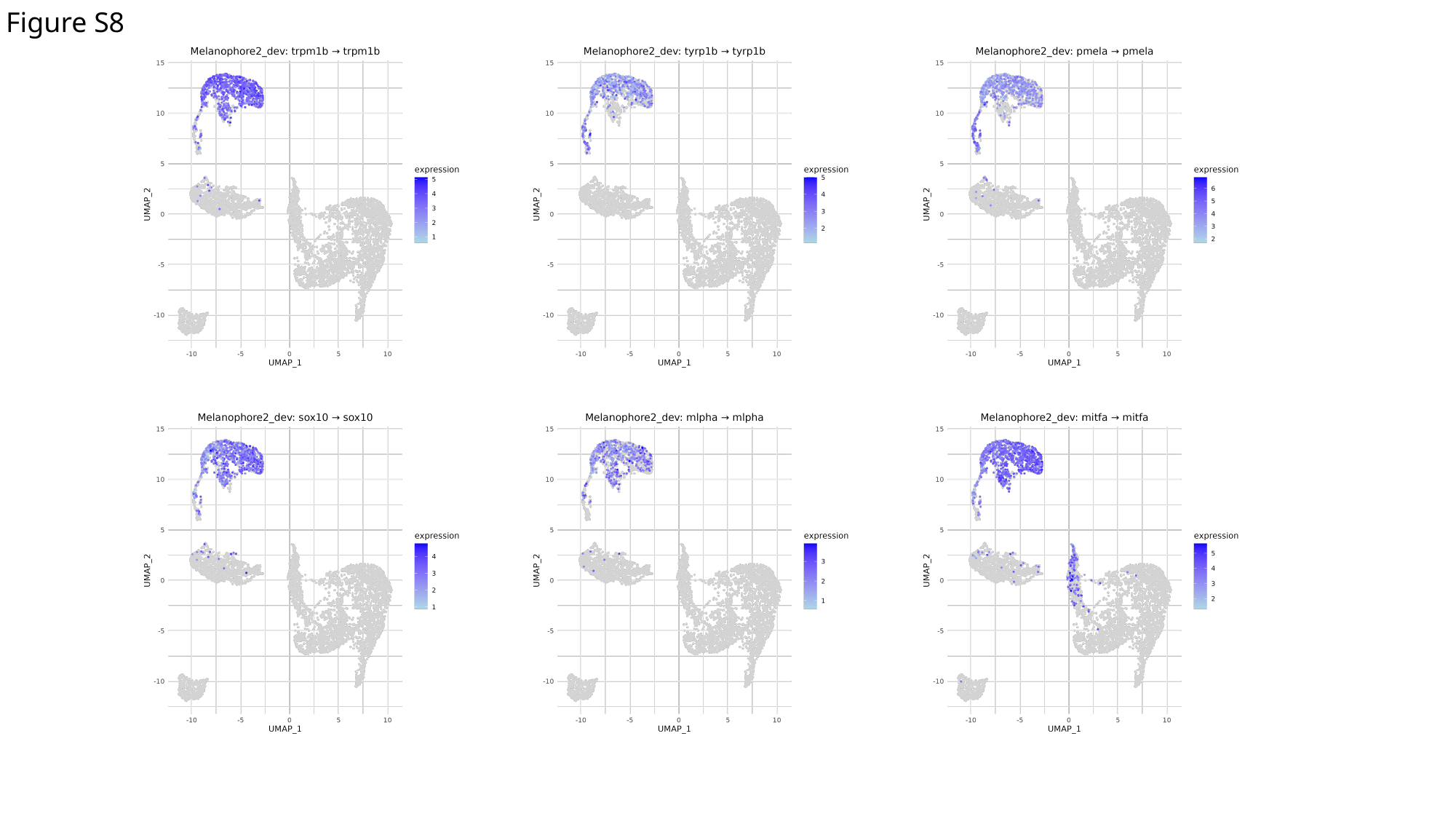

Figure S8

### Slide 9
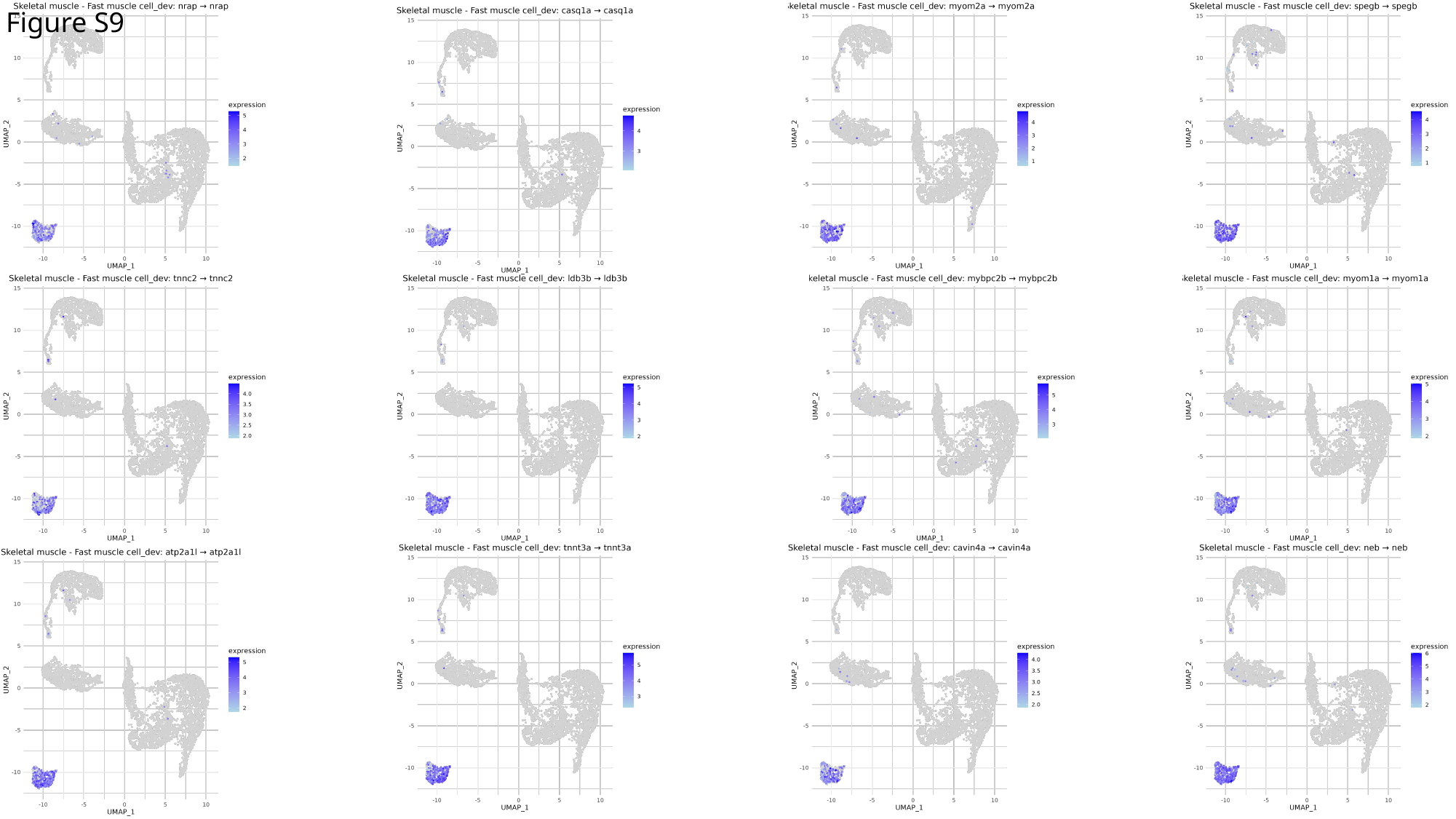

Figure S9

### Slide 10
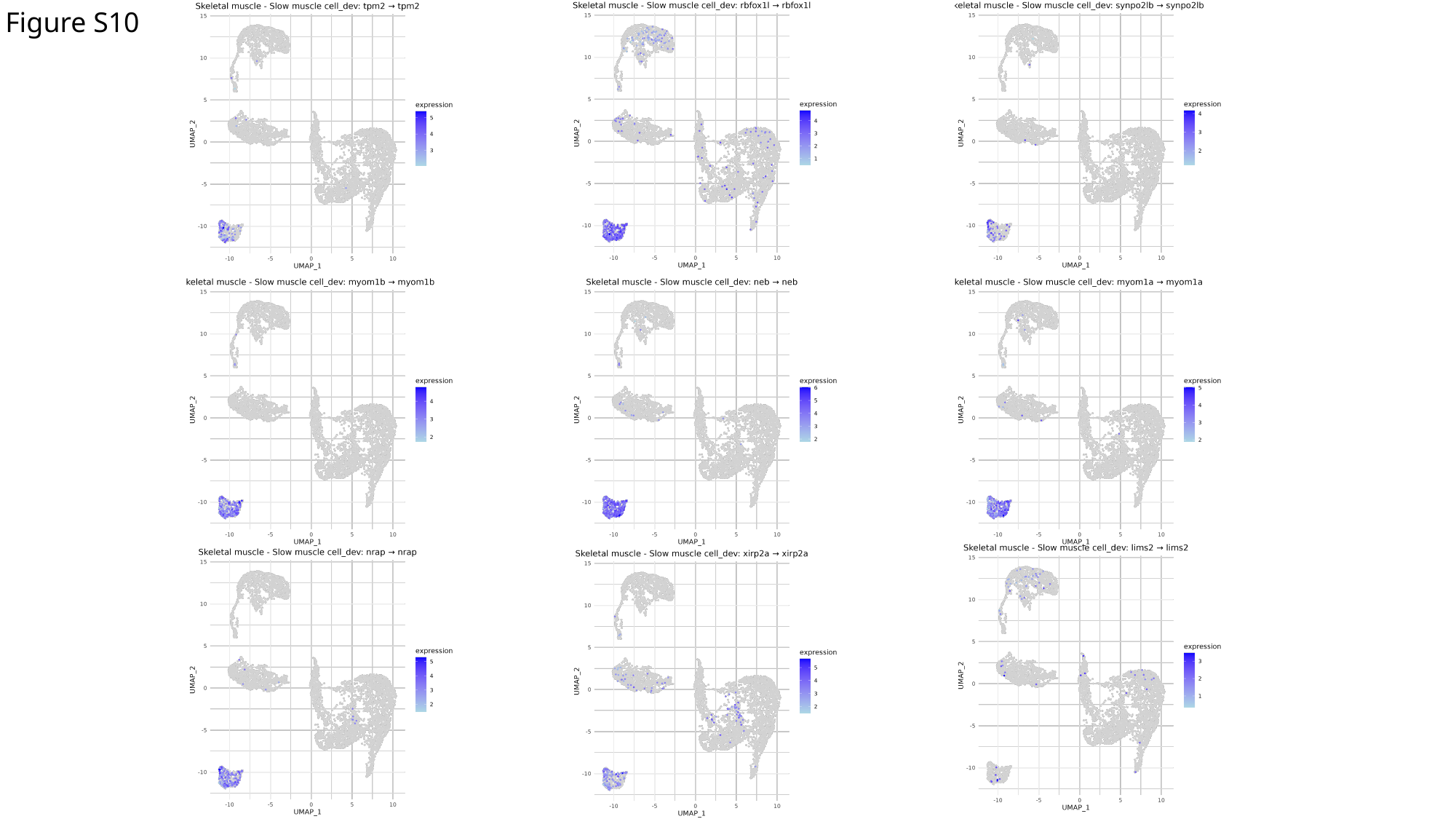

Figure S10

### Slide 11
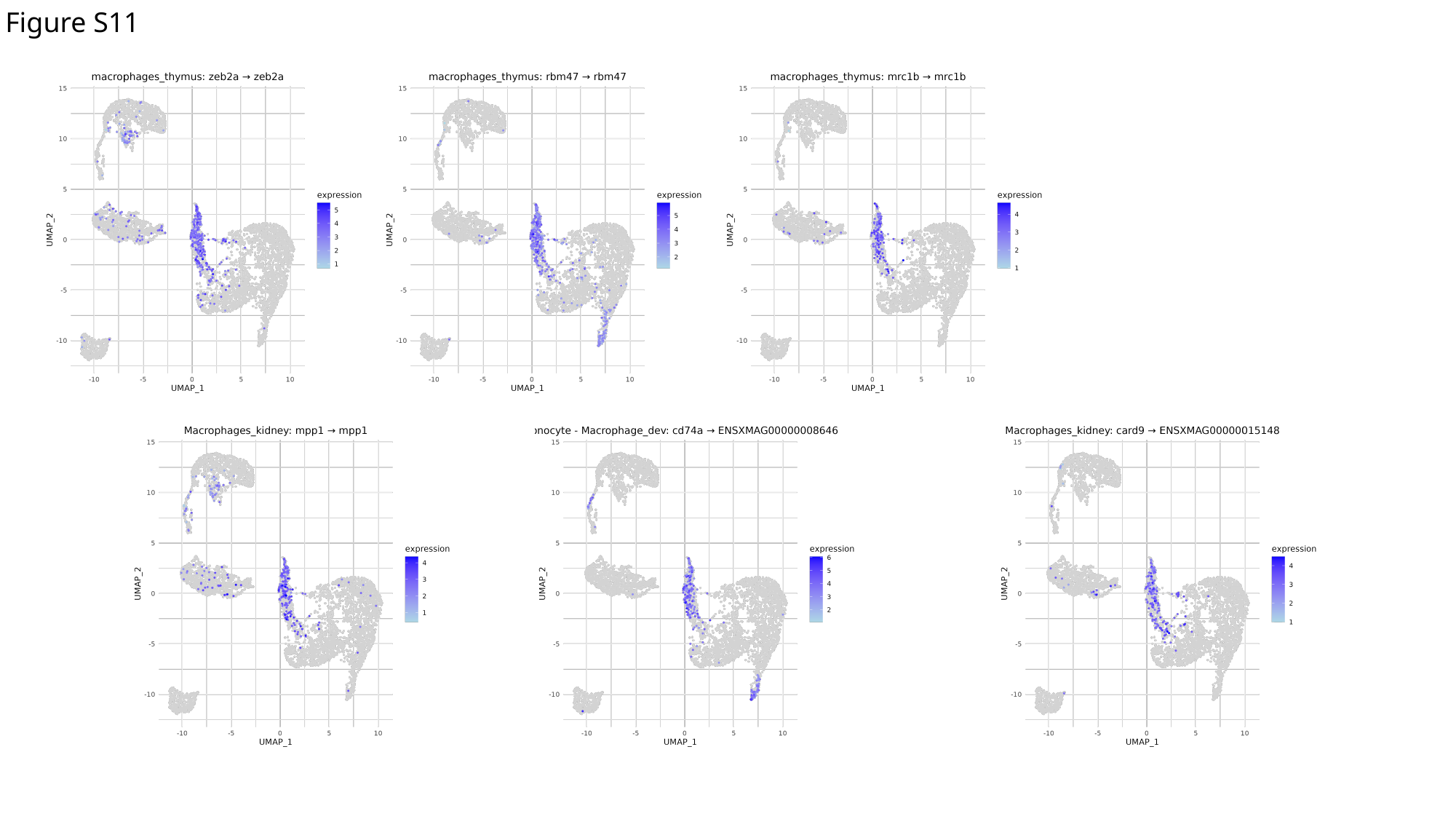

Figure S11

### Slide 12
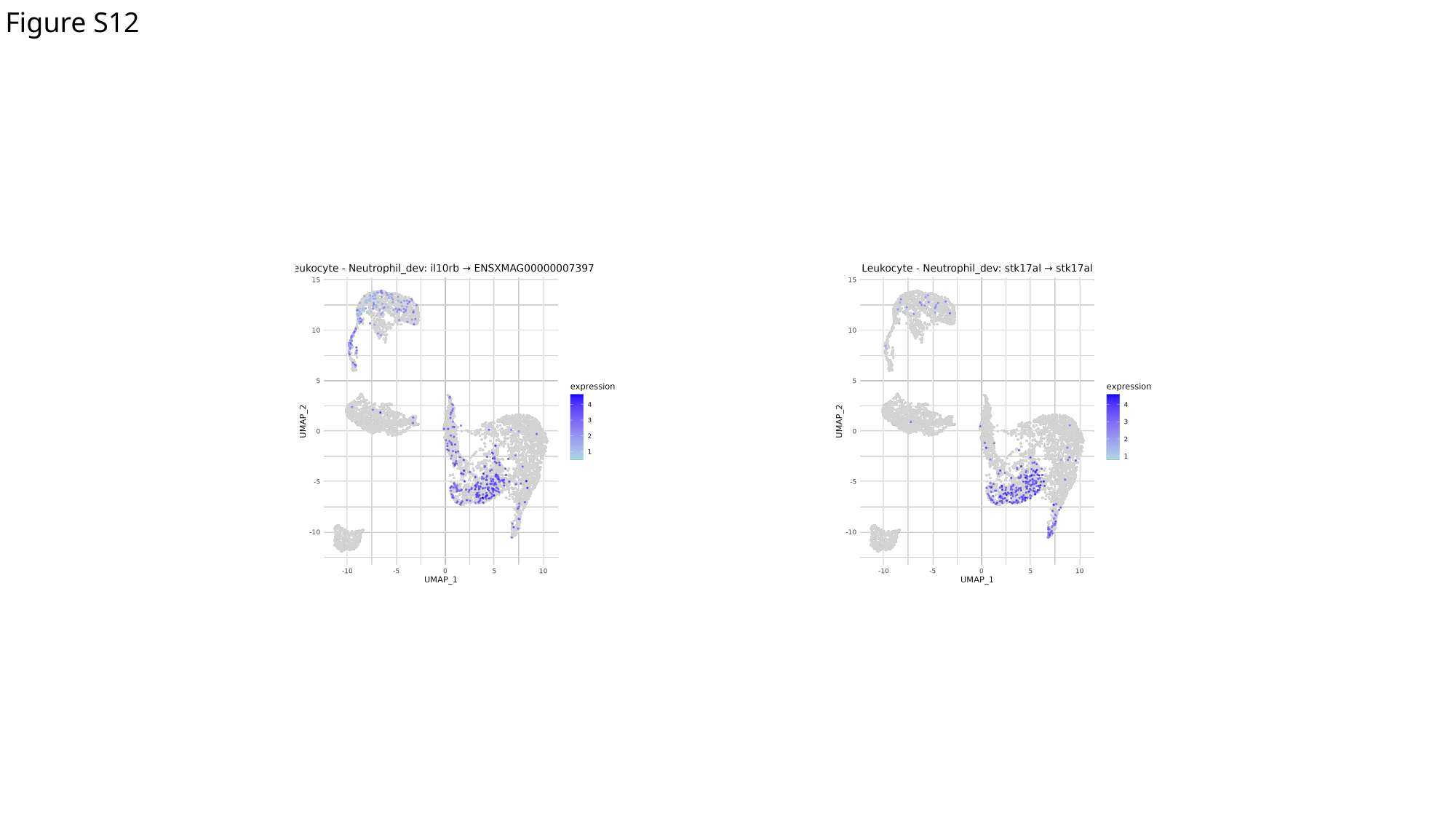

Figure S12

### Slide 13
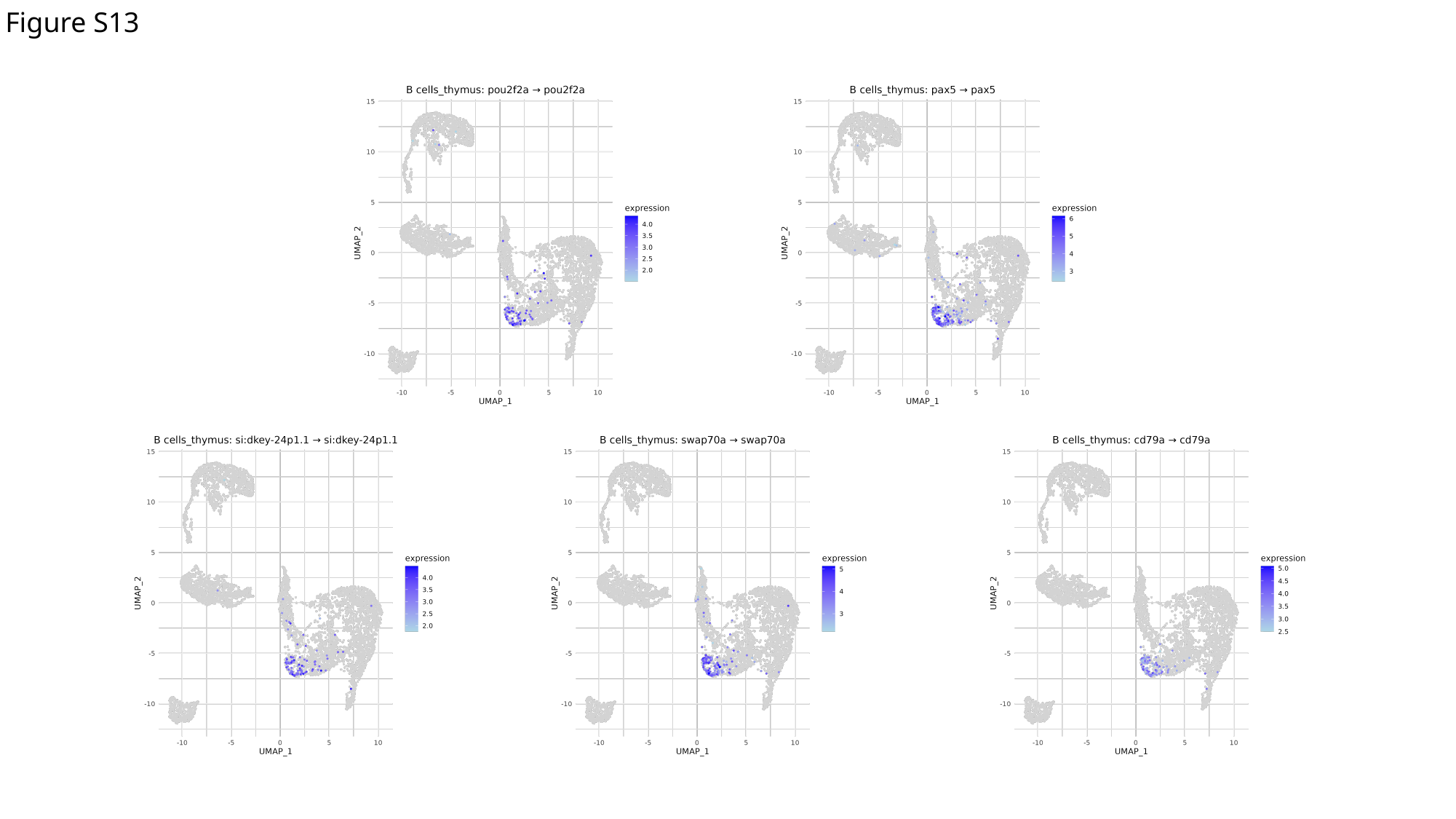

Figure S13

### Slide 14
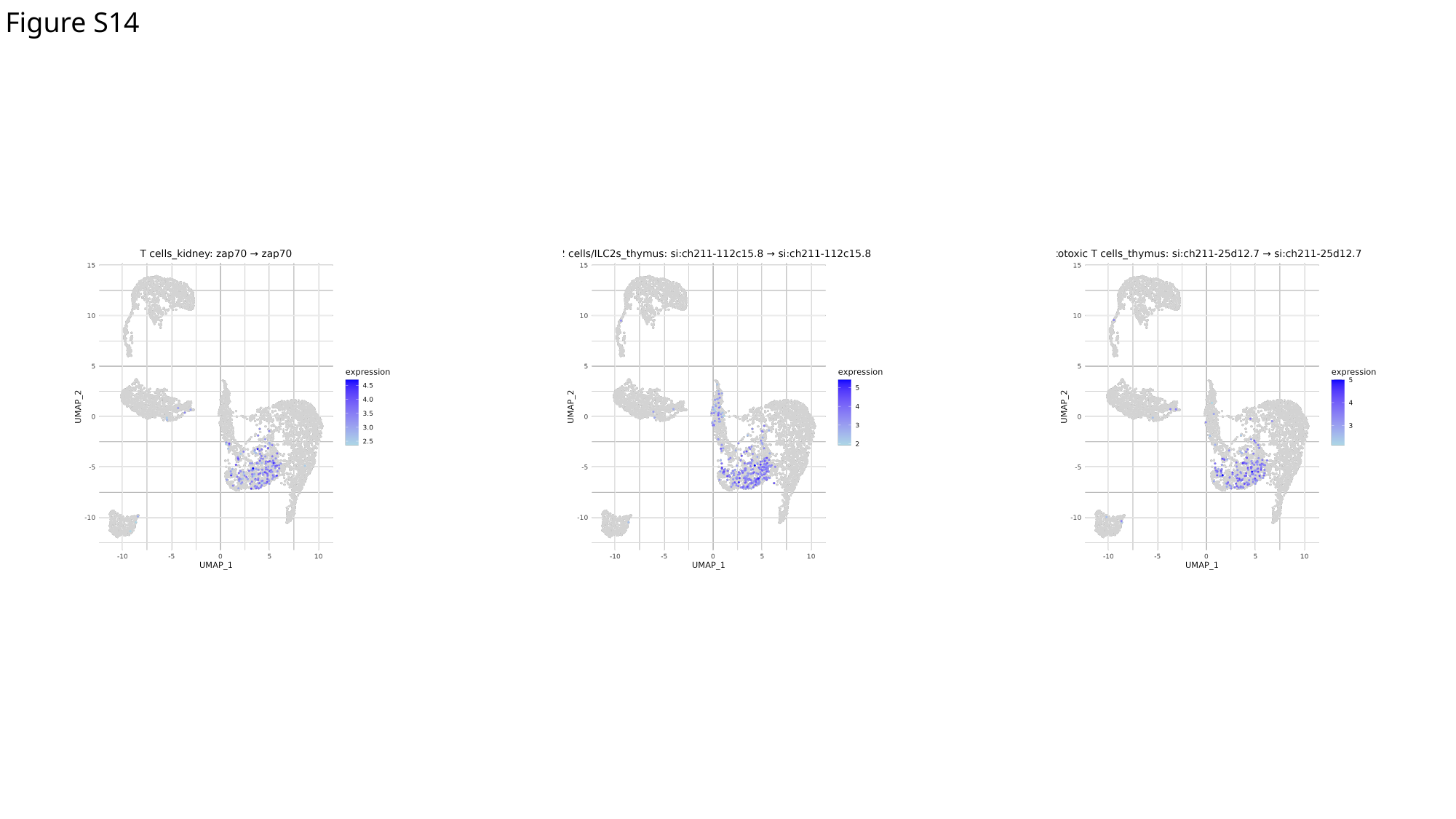

Figure S14

### Slide 15
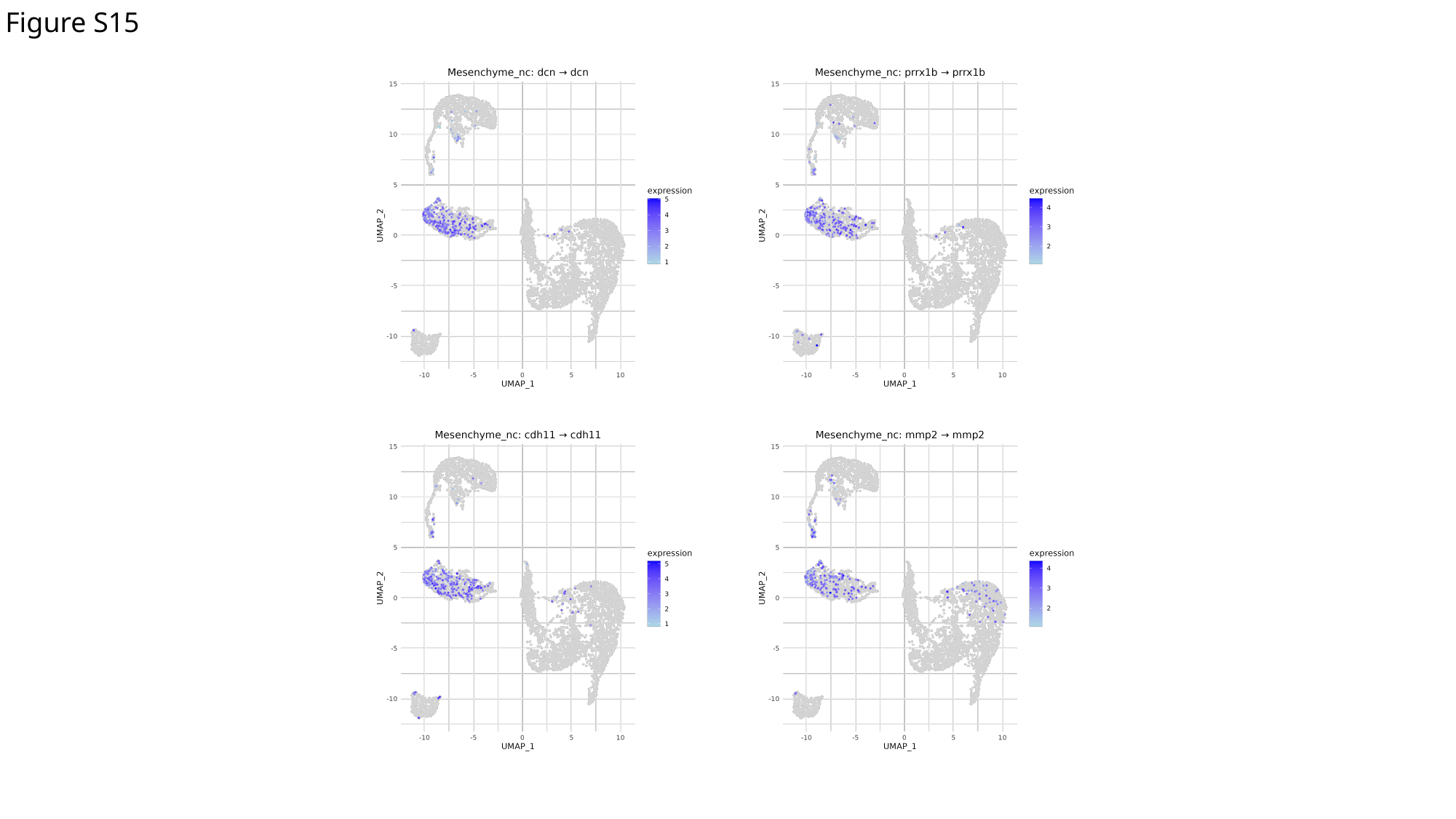

Figure S15

### Slide 16
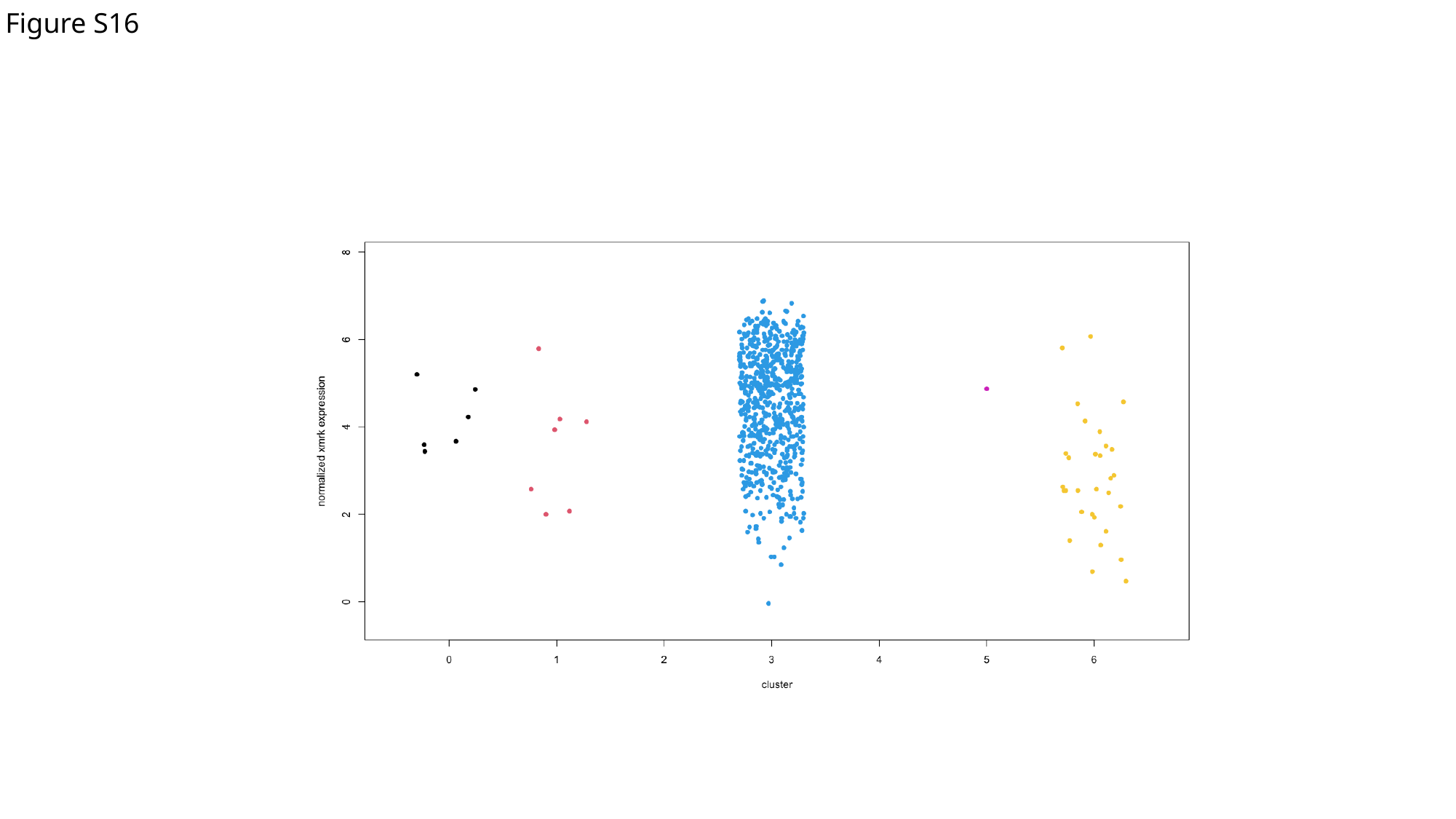

Figure S16

### Slide 17
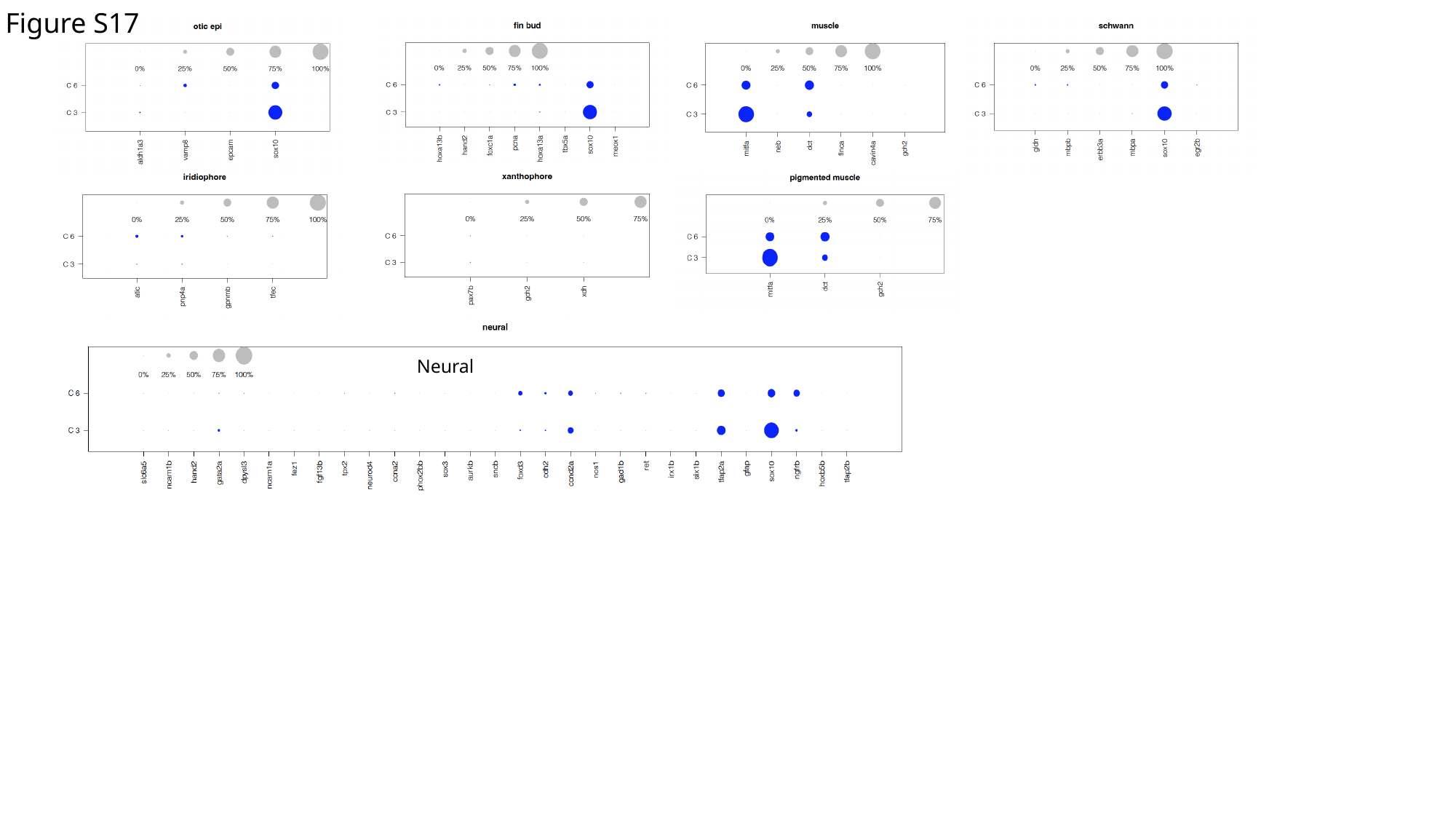

Figure S17
Neural

### Slide 18
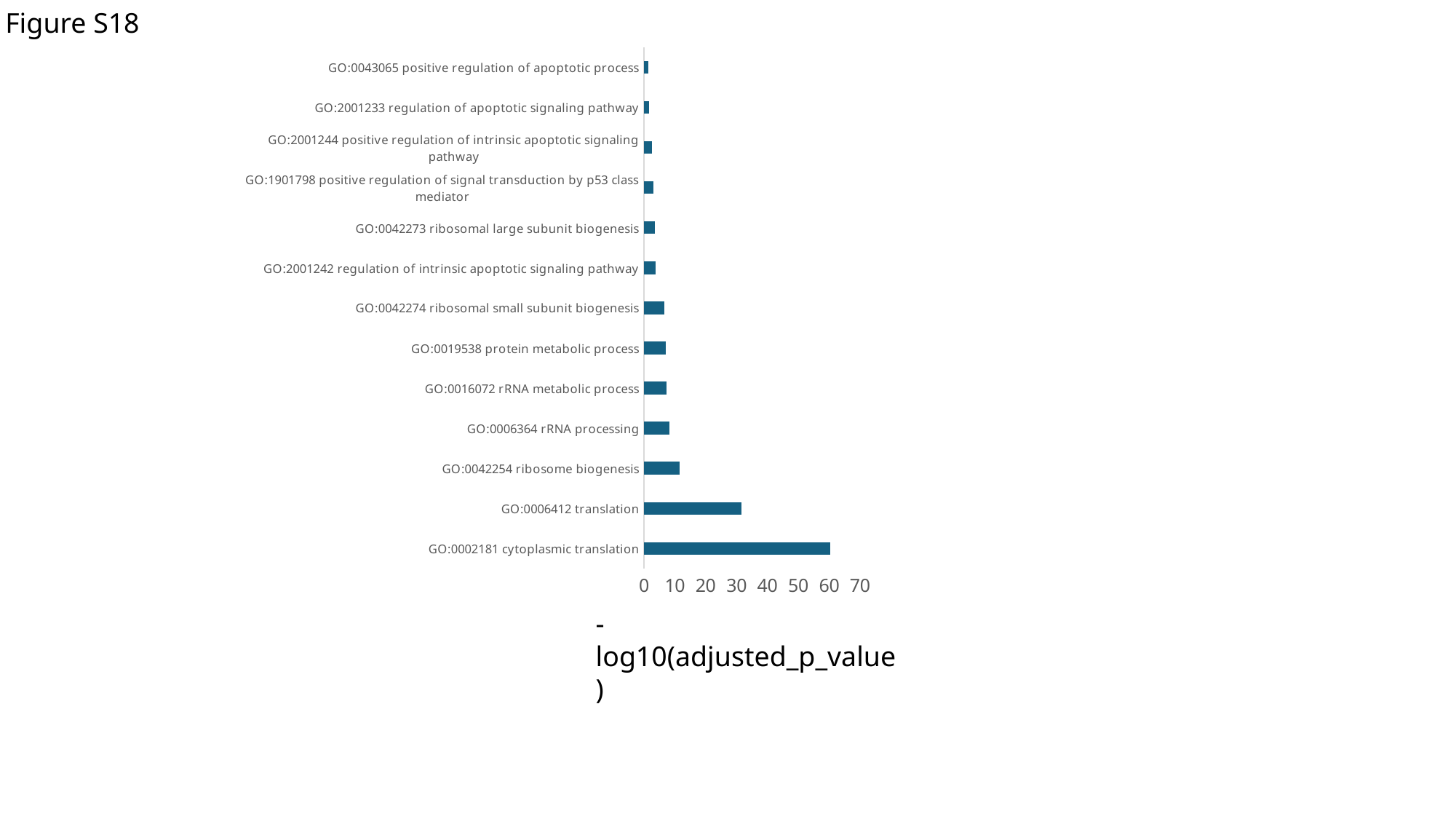

Figure S18
#### Chart
| Category | |
|---|---|
| GO:0002181 cytoplasmic translation | 60.27044763 |
| GO:0006412 translation | 31.55197579 |
| GO:0042254 ribosome biogenesis | 11.56377377 |
| GO:0006364 rRNA processing | 8.317769937 |
| GO:0016072 rRNA metabolic process | 7.182194378 |
| GO:0019538 protein metabolic process | 6.93682348 |
| GO:0042274 ribosomal small subunit biogenesis | 6.604596223 |
| GO:2001242 regulation of intrinsic apoptotic signaling pathway | 3.677310342 |
| GO:0042273 ribosomal large subunit biogenesis | 3.549006658 |
| GO:1901798 positive regulation of signal transduction by p53 class mediator | 3.123916598 |
| GO:2001244 positive regulation of intrinsic apoptotic signaling pathway | 2.593082134 |
| GO:2001233 regulation of apoptotic signaling pathway | 1.701439813 |
| GO:0043065 positive regulation of apoptotic process | 1.395066812 |-log10(adjusted_p_value)
