## Supplementary material for "Comparison of Human Melanoma Single Cell Profiles to Evolutionary Medicine Model *Xiphophorus* Provides Insights in Disease Control": Supp. files: UMAP_AXL_orthologs_expressed.pdf

### AXL program: CTNNA1 ortholog ... ctnna1

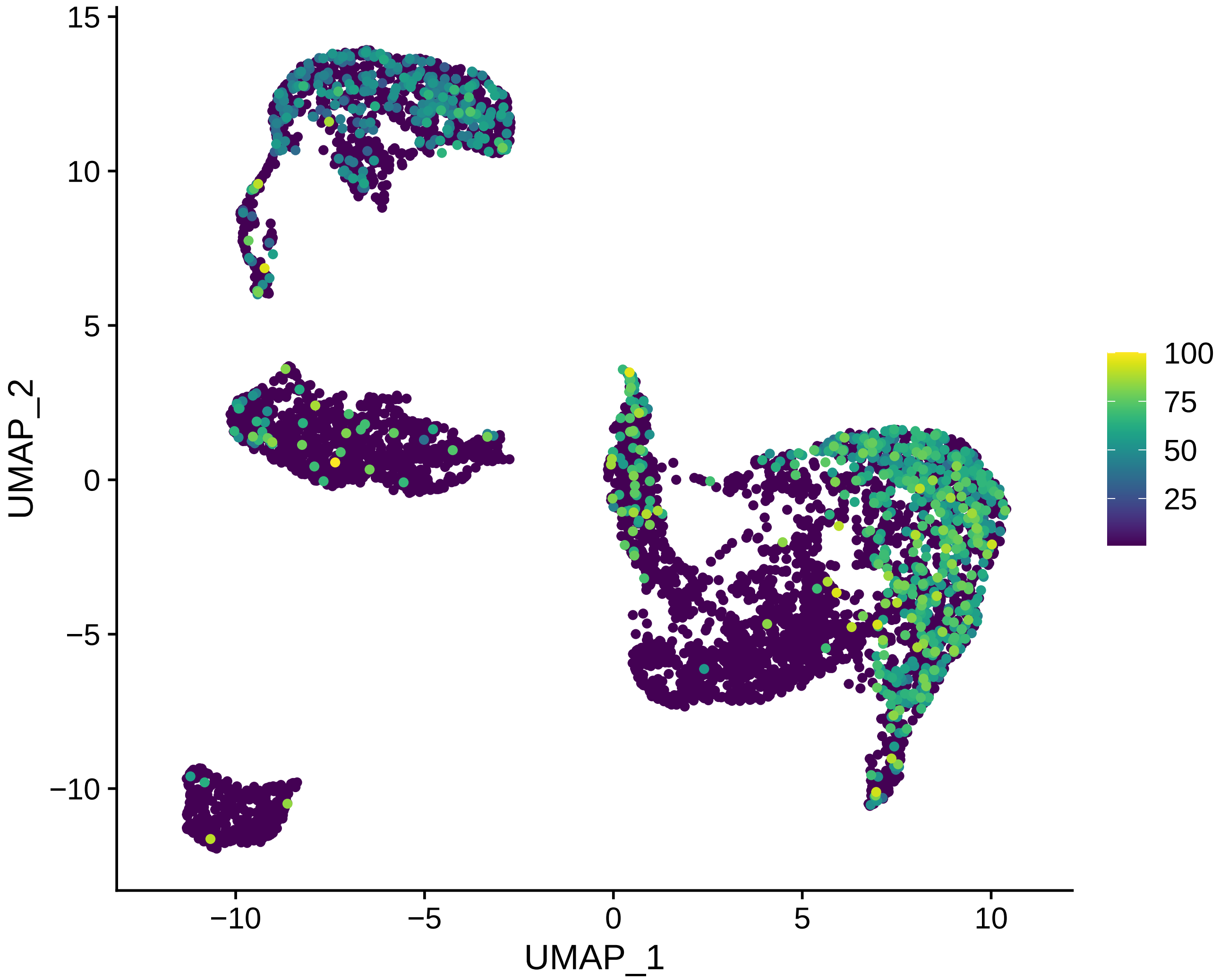

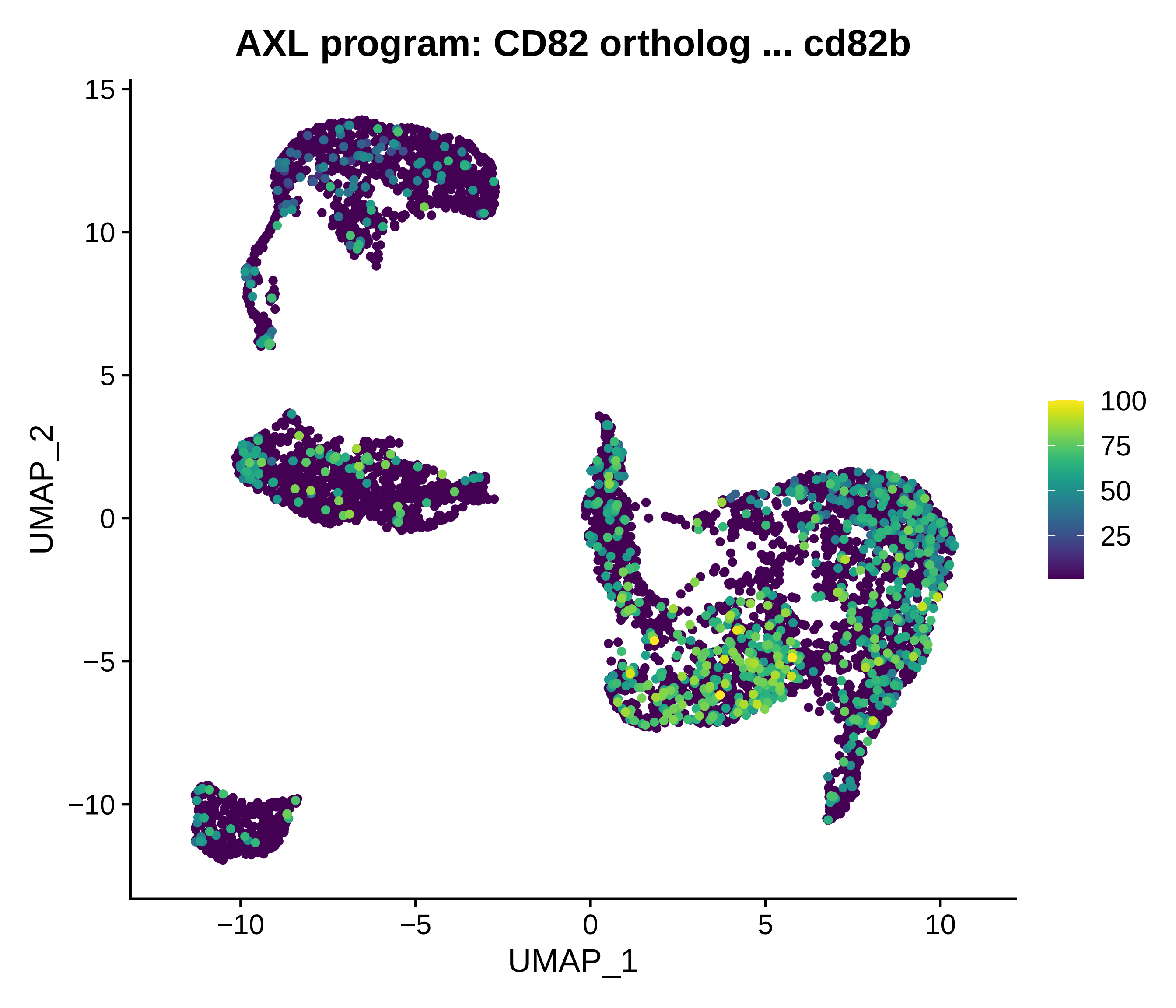
AXL program: CD82 ortholog ... cd82b
15
10
5
0
−5
−10
UMAP_2
−10
−5
0
5
10
UMAP_1
100
75
50
25

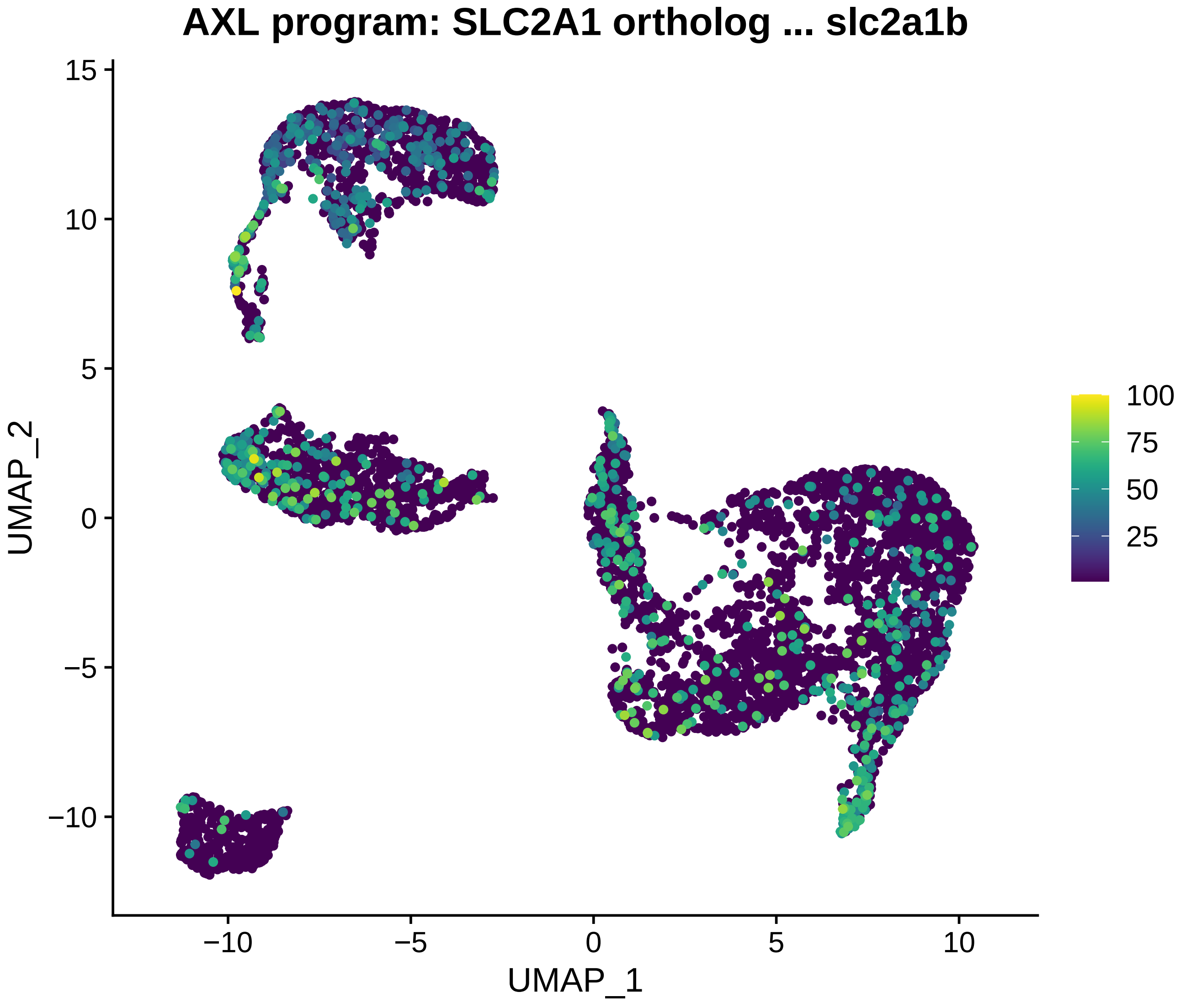
AXL program: SLC2A1 ortholog ... slc2a1b
UMAP_2
UMAP_1
15
10
5
0
−5
−10
−10
−5
0
5
10
100
75
50
25

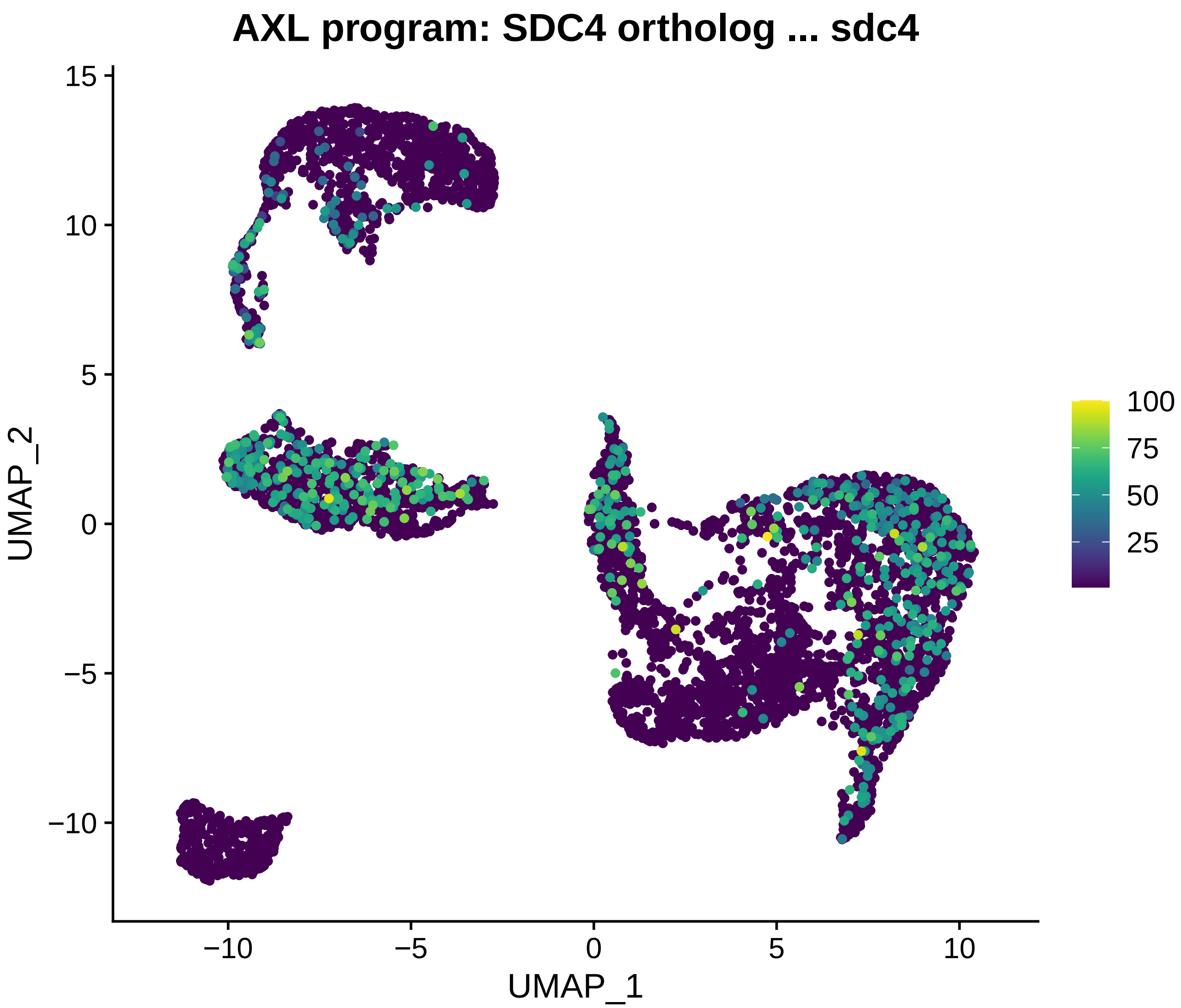

AXL program: SDC4 ortholog ... sdc4
UMAP_2
UMAP_1
15
10
5
0
−5
−10
−10
−5
0
5
10
100
75
50
25

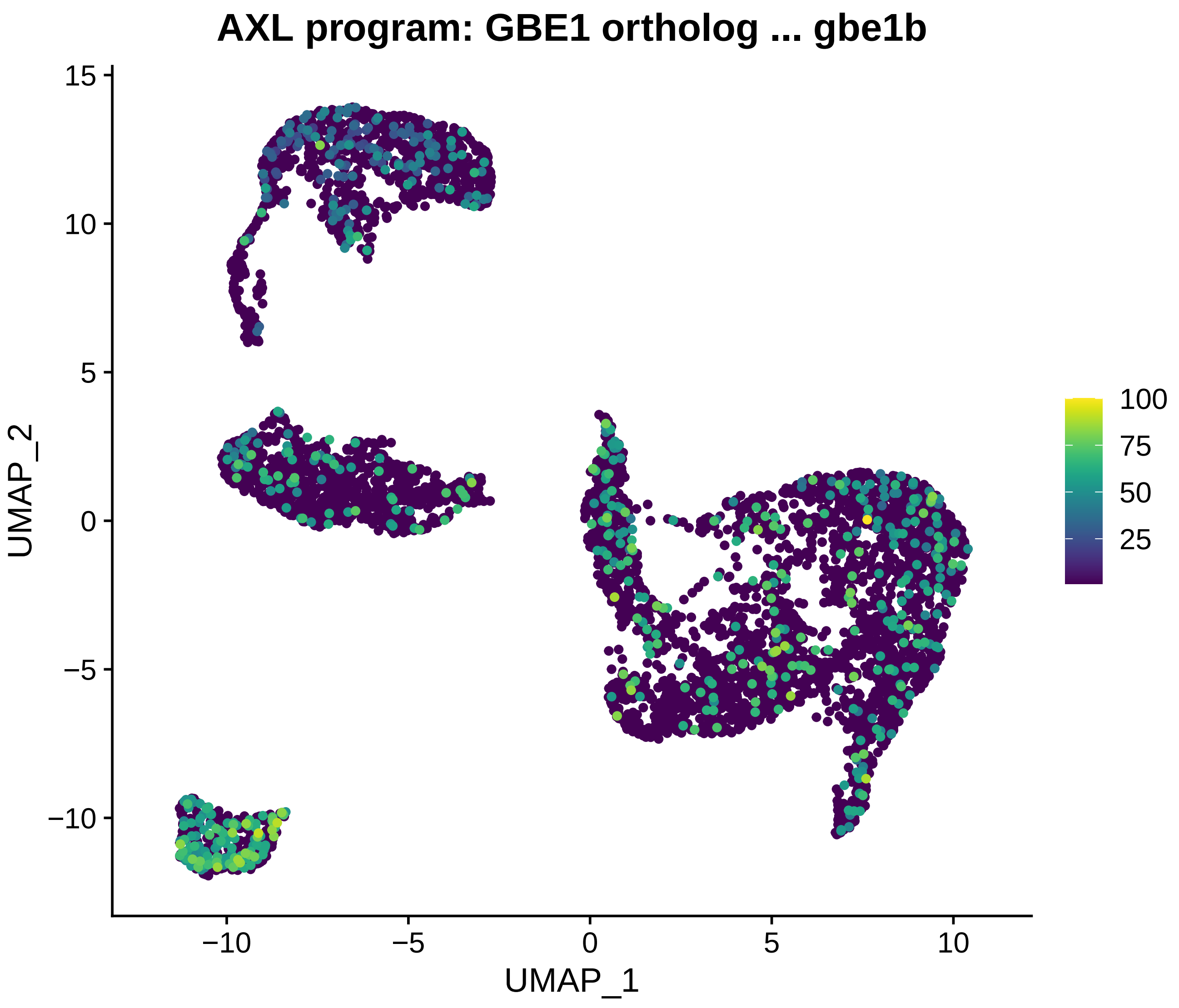
AXL program: GBE1 ortholog ... gbe1b
UMAP_2
UMAP_1
15
10
5
0
−5
−10
−10
−5
0
5
10
100
75
50
25

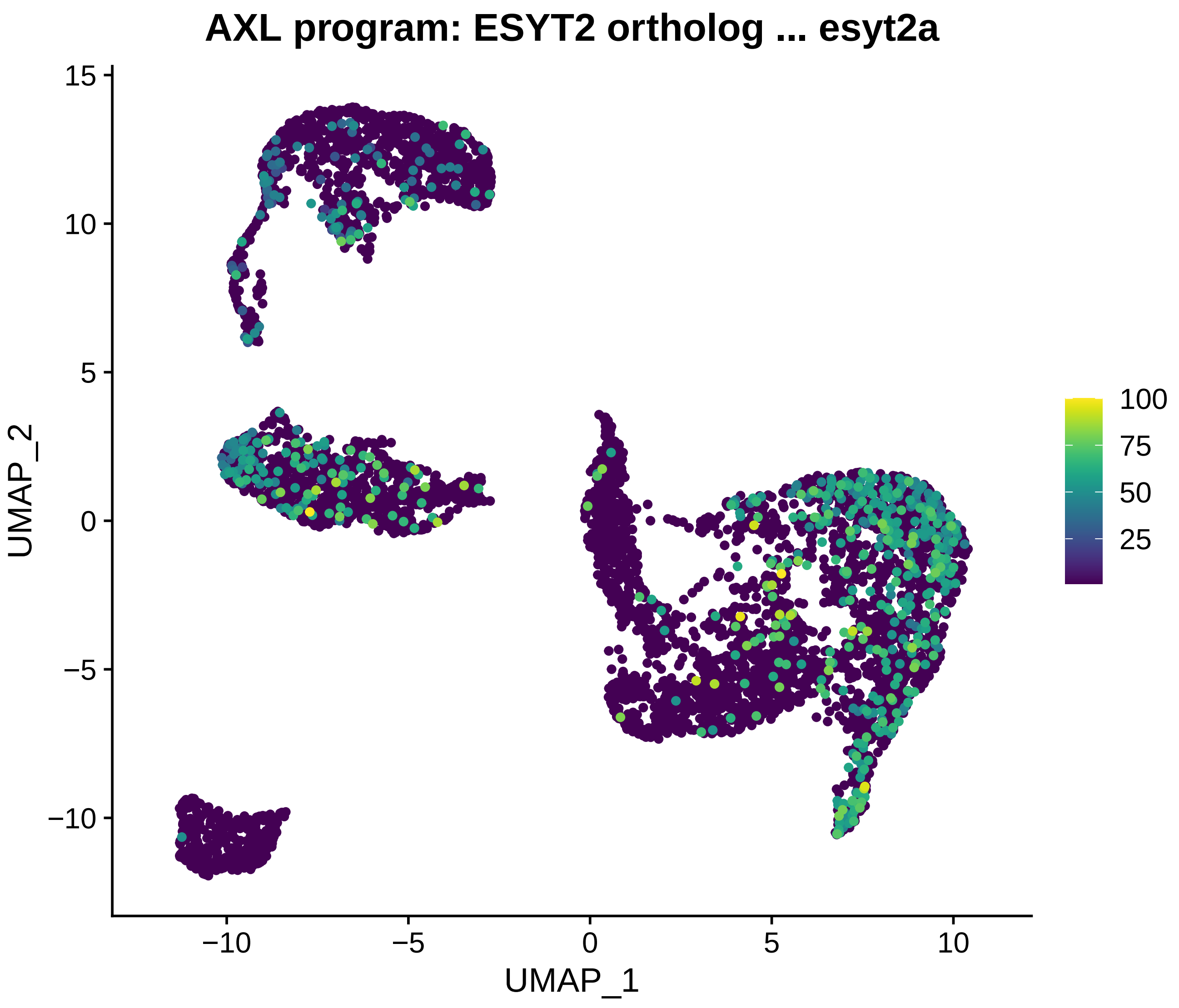
AXL program: ESYT2 ortholog ... esyt2a
UMAP_2
UMAP_1
15
10
5
0
−5
−10
−10
−5
0
5
10
100
75
50
25

#### AXL program: LTBP3 ortholog ... LTBP3

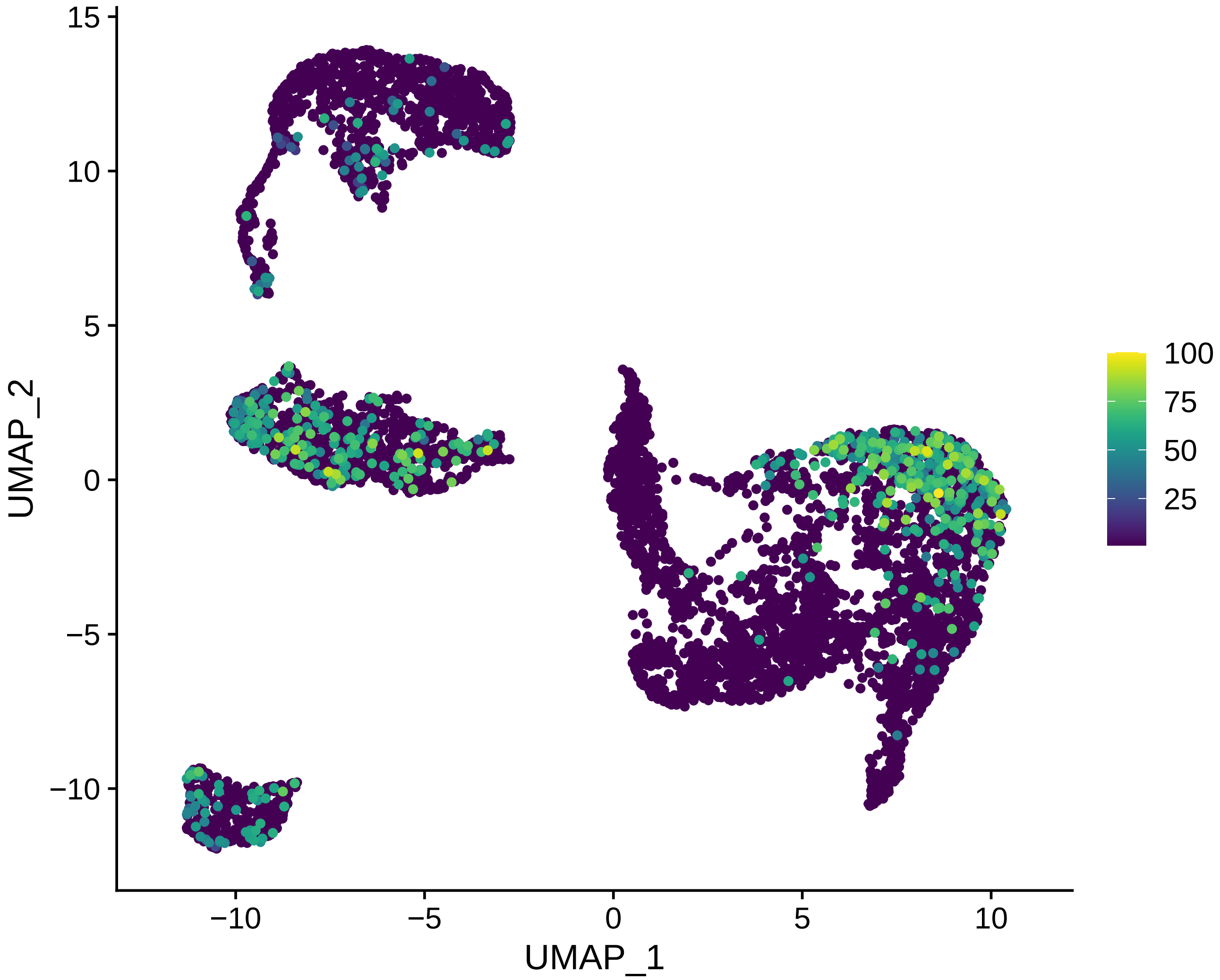

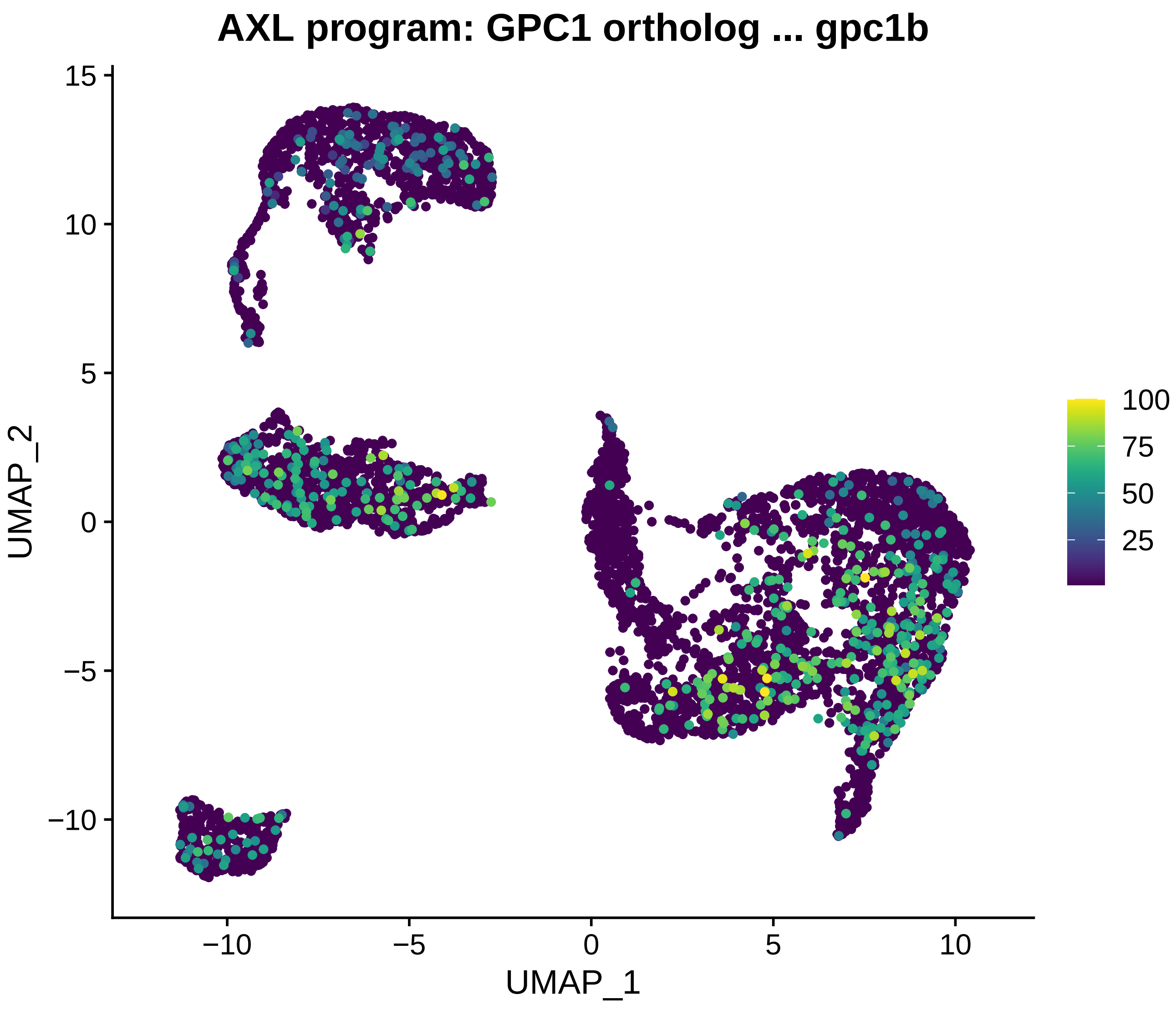
AXL program: GPC1 ortholog ... gpc1b
UMAP_2
UMAP_1
15
10
5
0
−5
−10
−10
−5
0
5
10
100
75
50
25

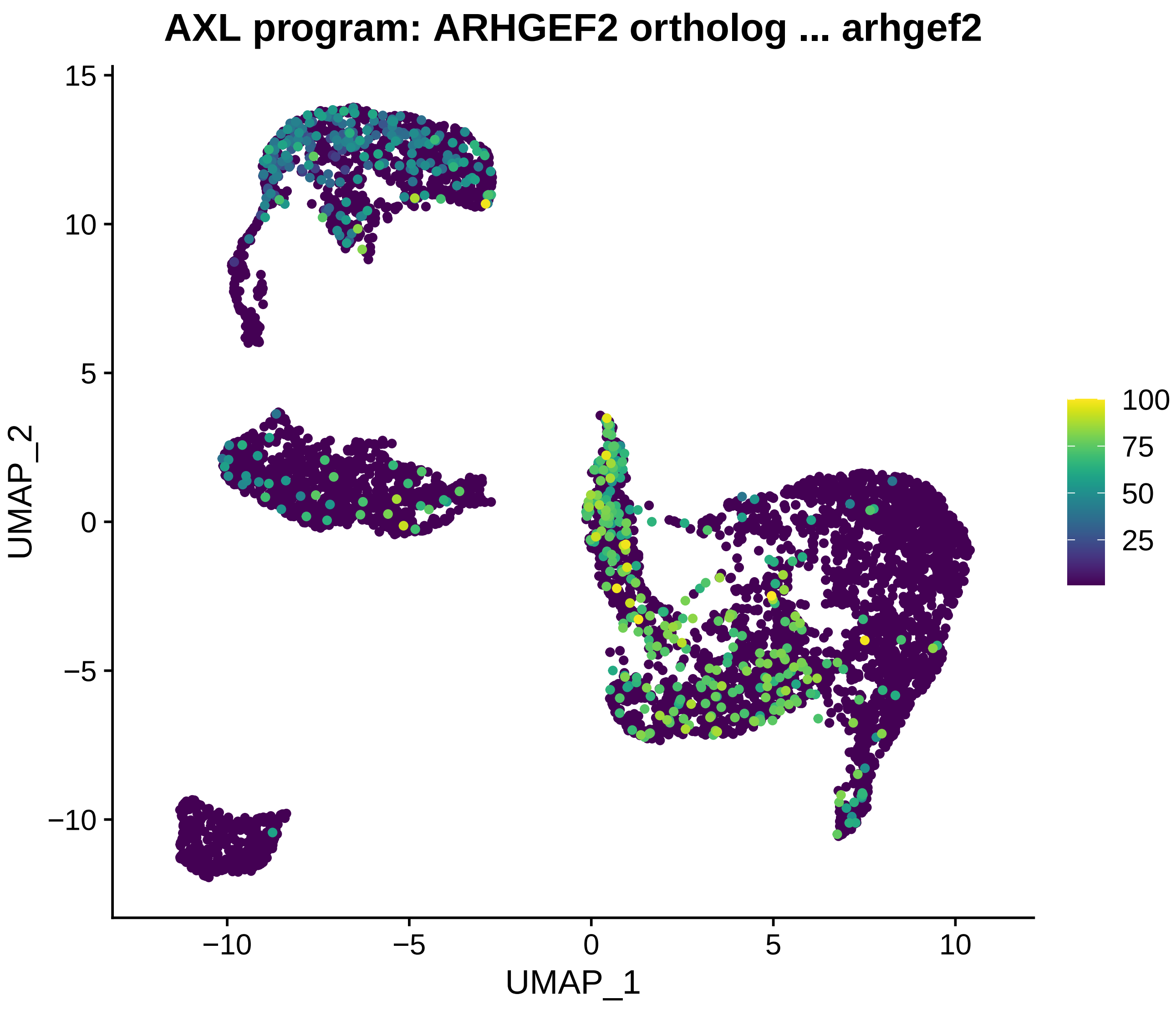
AXL program: ARHGEF2 ortholog ... arhgef2
UMAP_2
UMAP_1
15
10
5
0
−5
−10
−10
−5
0
5
10
100
75
50
25

### AXL program: TBC1D8 ortholog ... TBC1D8

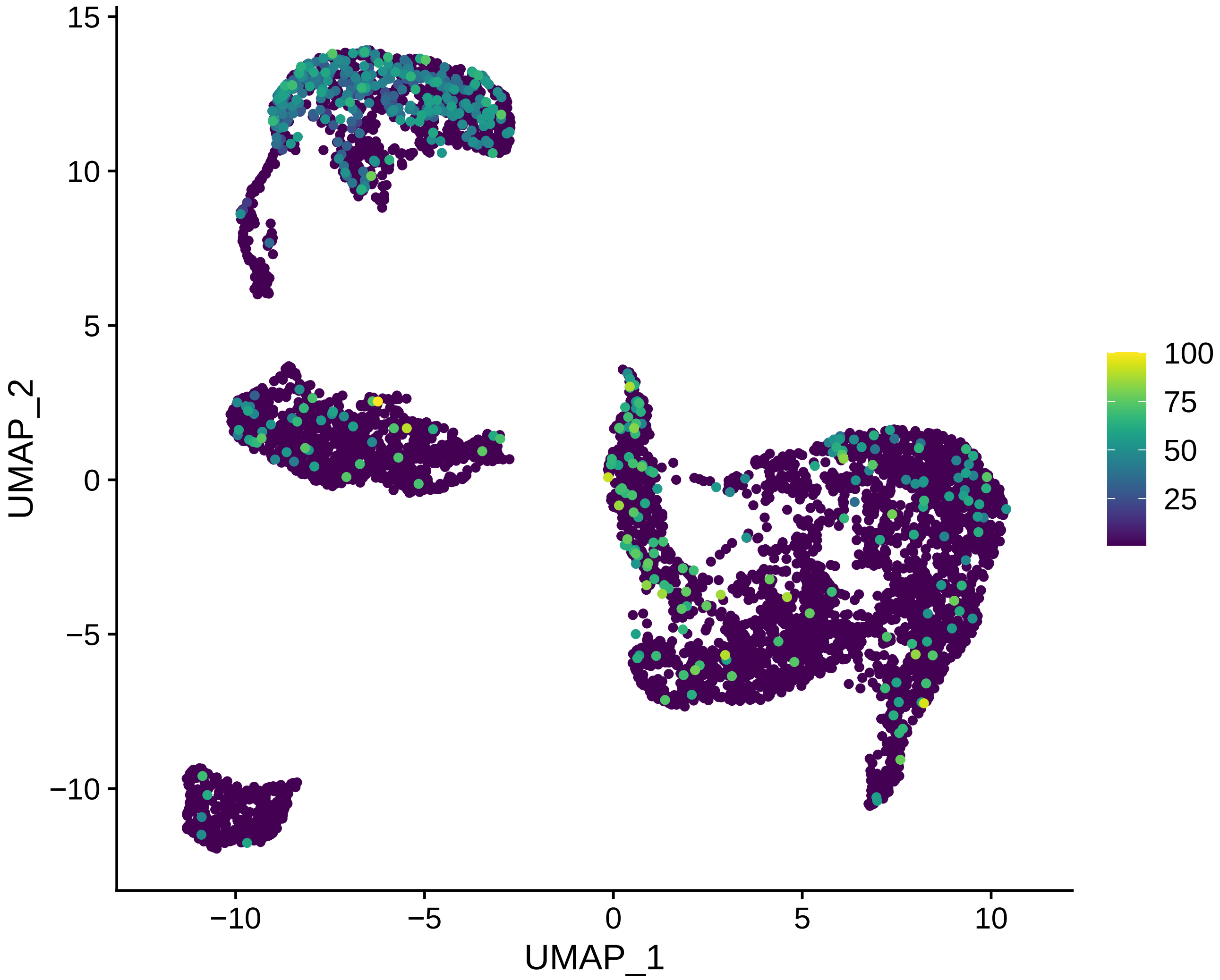

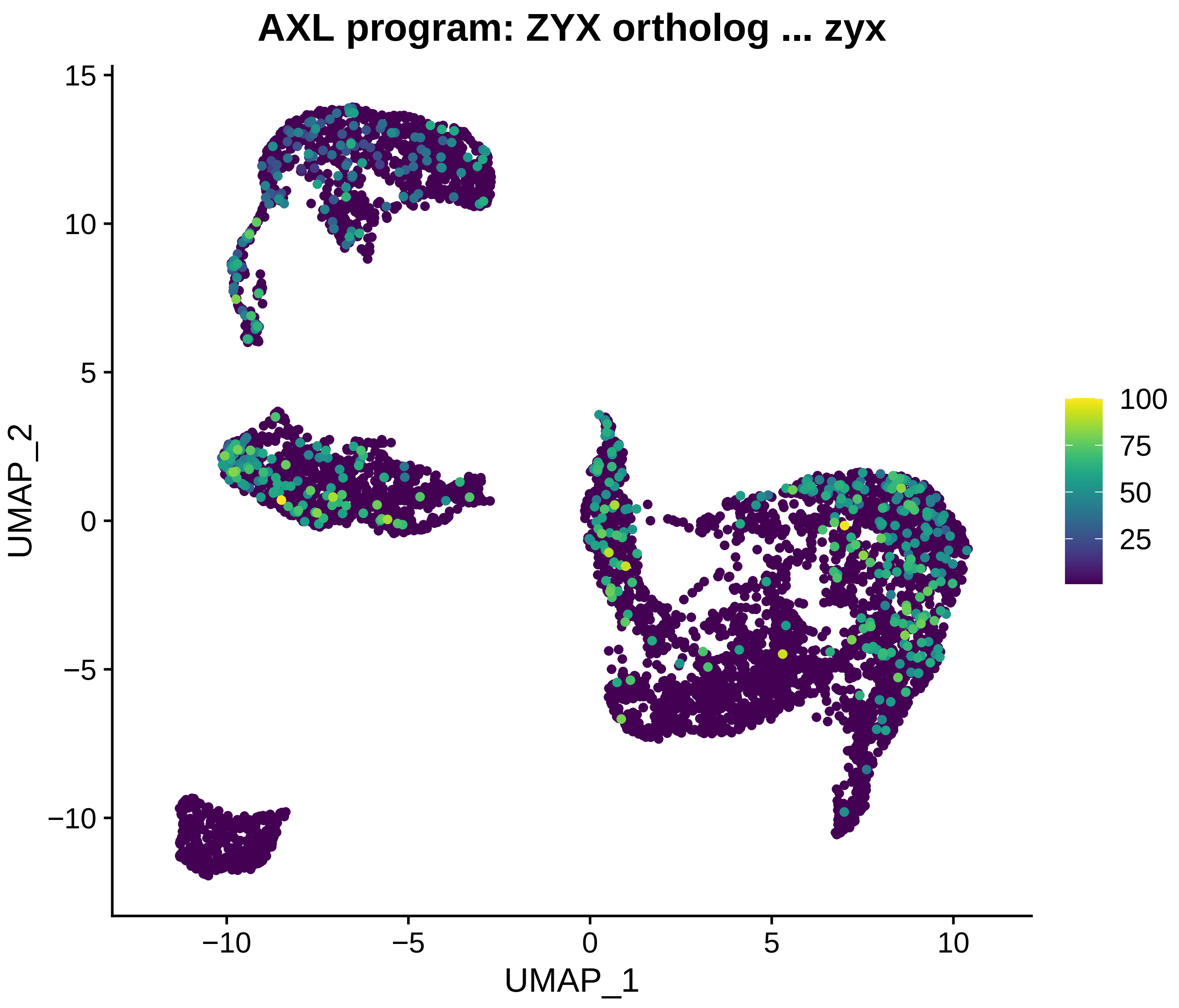

AXL program: ZYX ortholog ... zyx
UMAP_2
UMAP_1
15
10
5
0
−5
−10
−10
−5
0
5
10
100
75
50
25

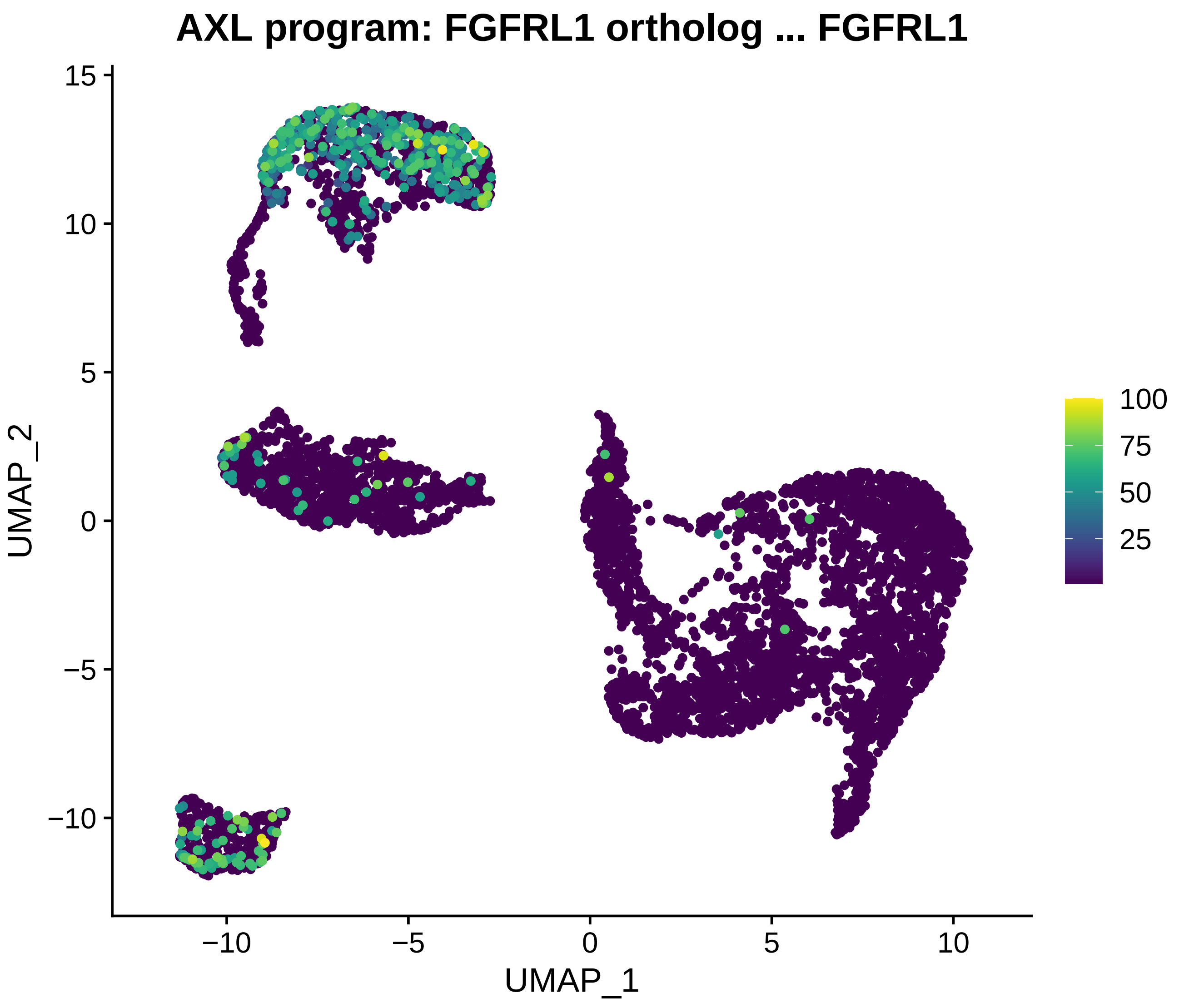
AXL program: FGFRL1 ortholog ... FGFRL1
UMAP_2
UMAP_1
15
10
5
0
−5
−10
−10
−5
0
5
10
100
75
50
25

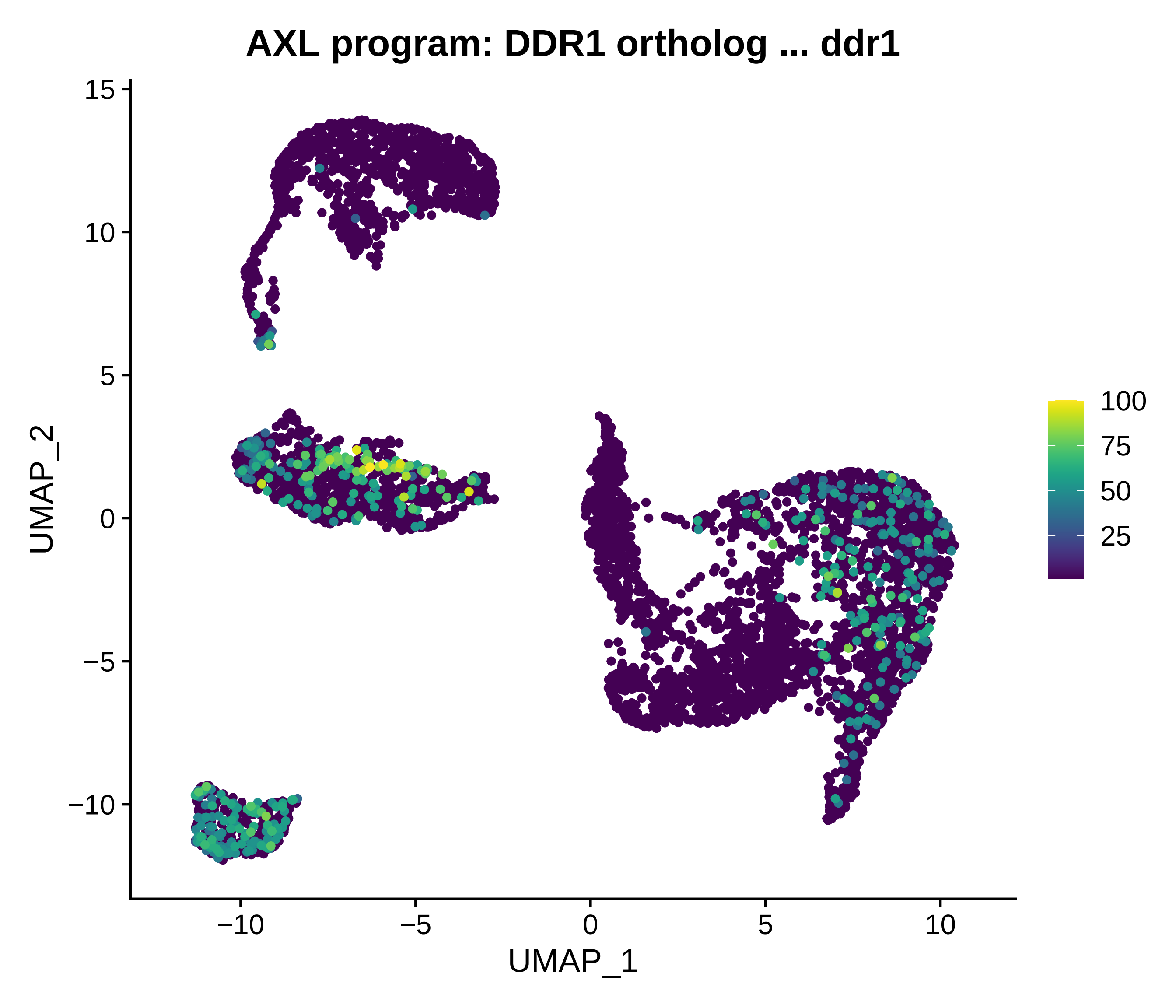
AXL program: DDR1 ortholog ... ddr1
UMAP_2
UMAP_1
15
10
5
0
−5
−10
−10
−5
0
5
10
100
75
50
25

### AXL program: PPFIBP1 ortholog ... ppfibp1b

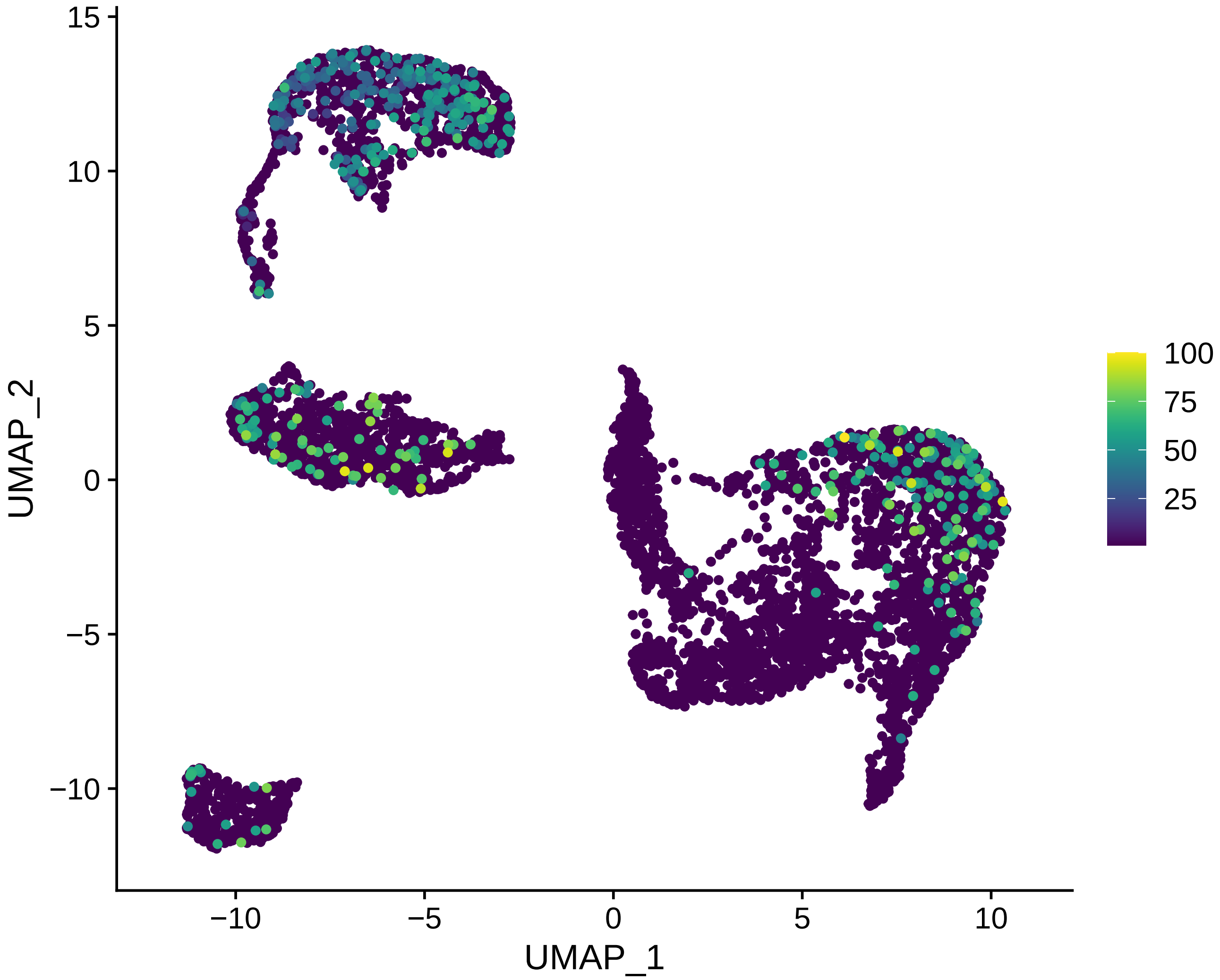

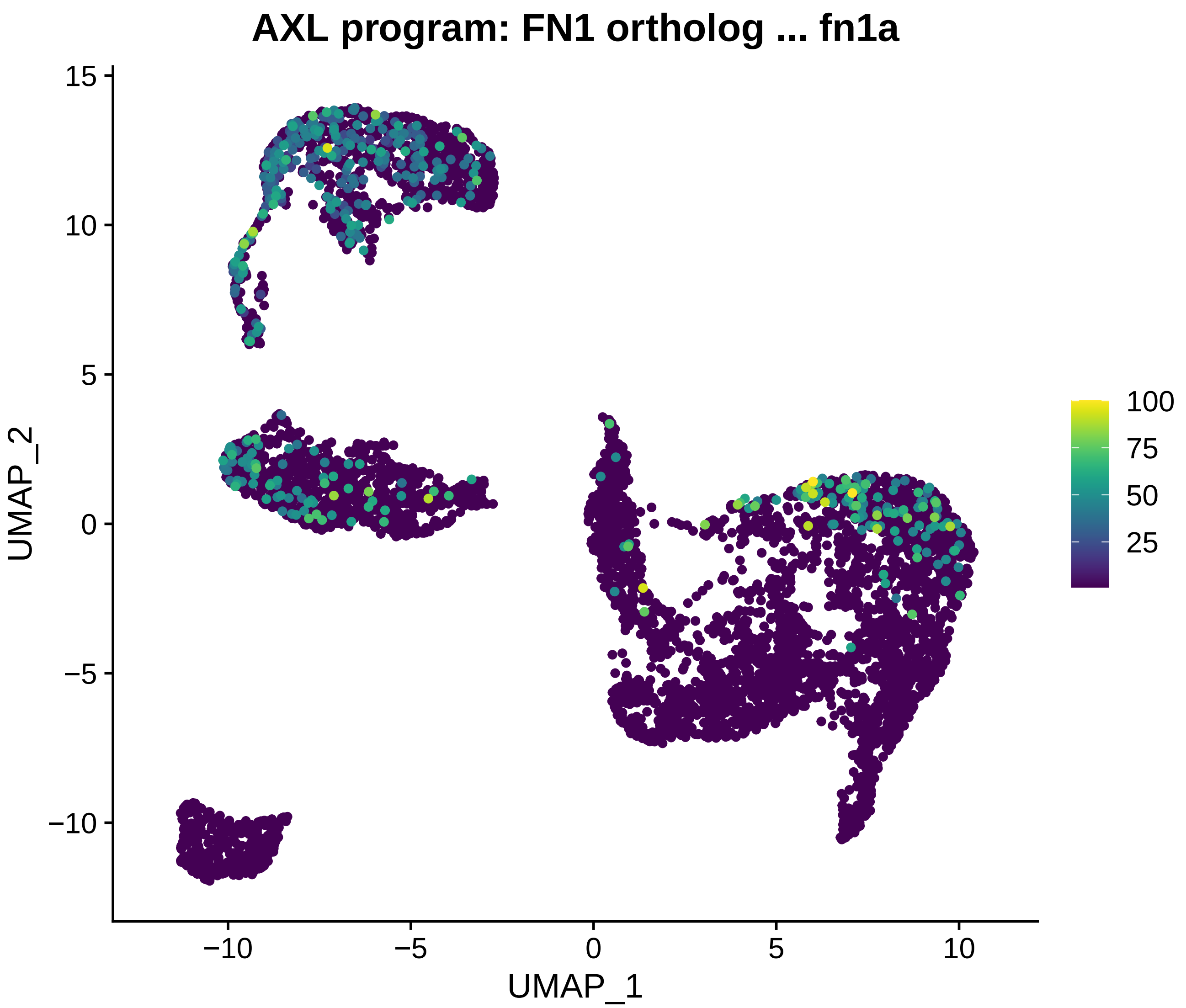
AXL program: FN1 ortholog ... fn1a
UMAP_2
UMAP_1
15
10
5
0
−5
−10
10
100
75
50
25

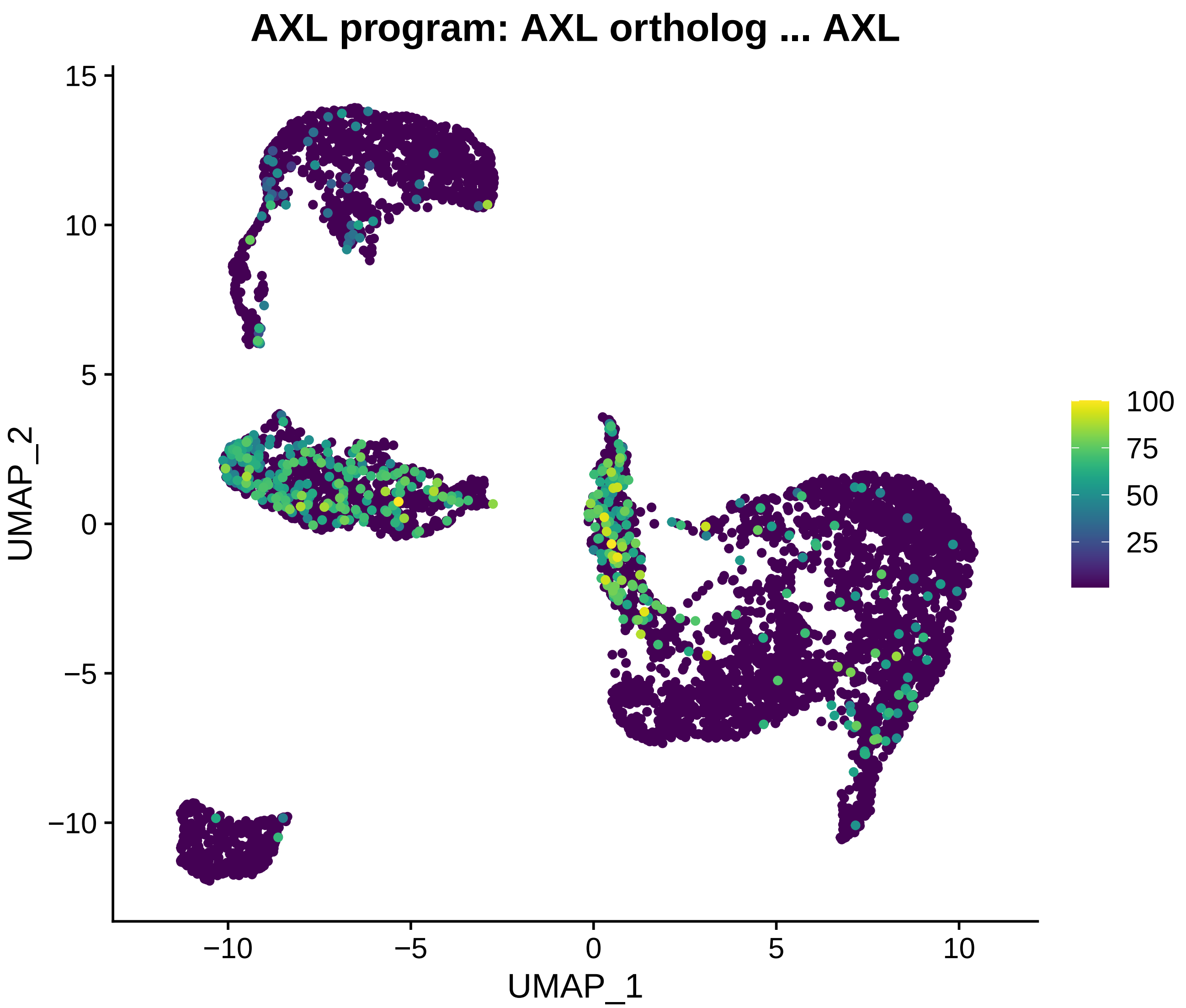
AXL program: AXL ortholog ... AXL
UMAP_2
UMAP_1
15
10
5
0
−5
−10
−10
−5
0
5
10
100
75
50
25

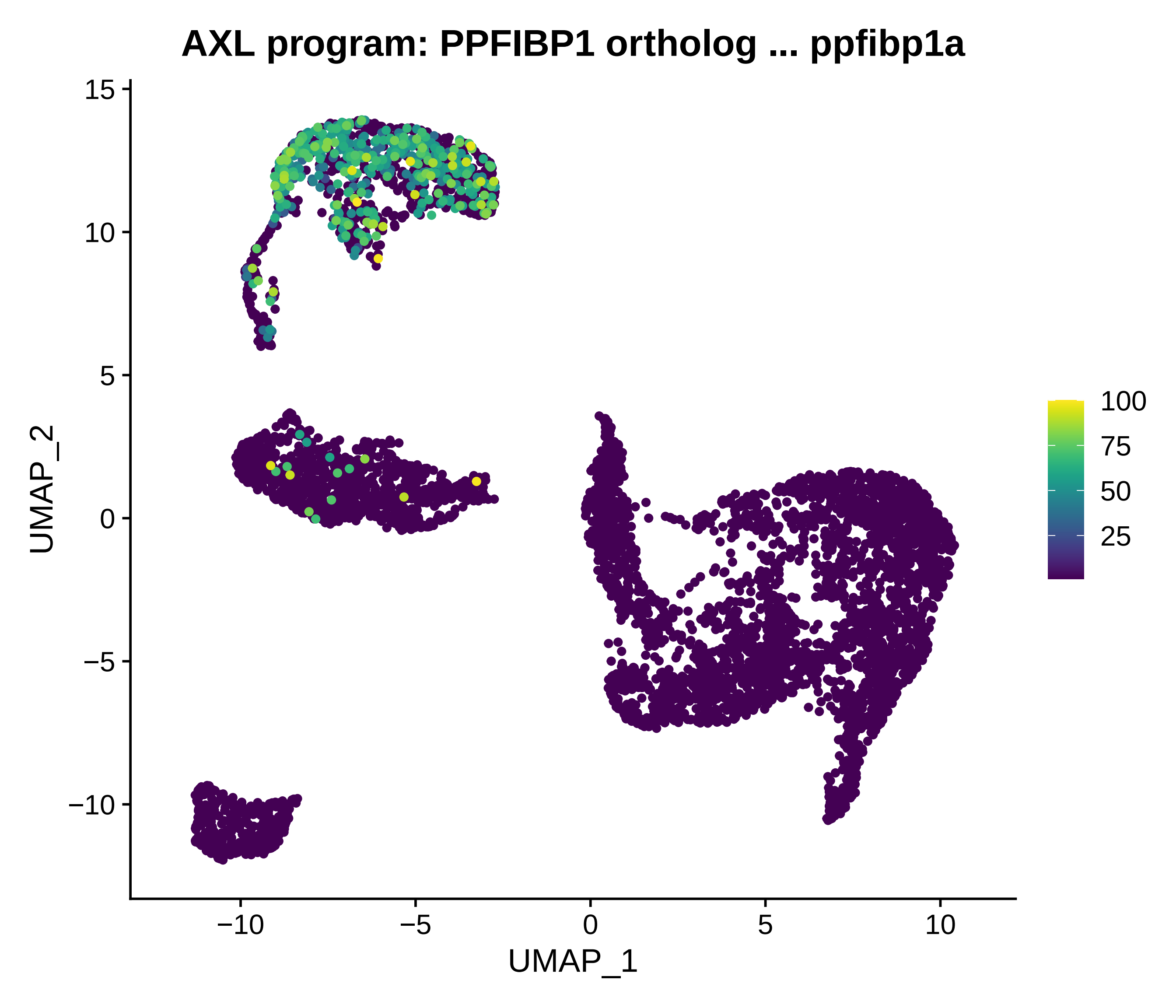
AXL program: PPFIBP1 ortholog ... ppfibp1a
UMAP_2
UMAP_1
15
10
5
0
−5
−10
−10
−5
0
5
10
100
75
50
25

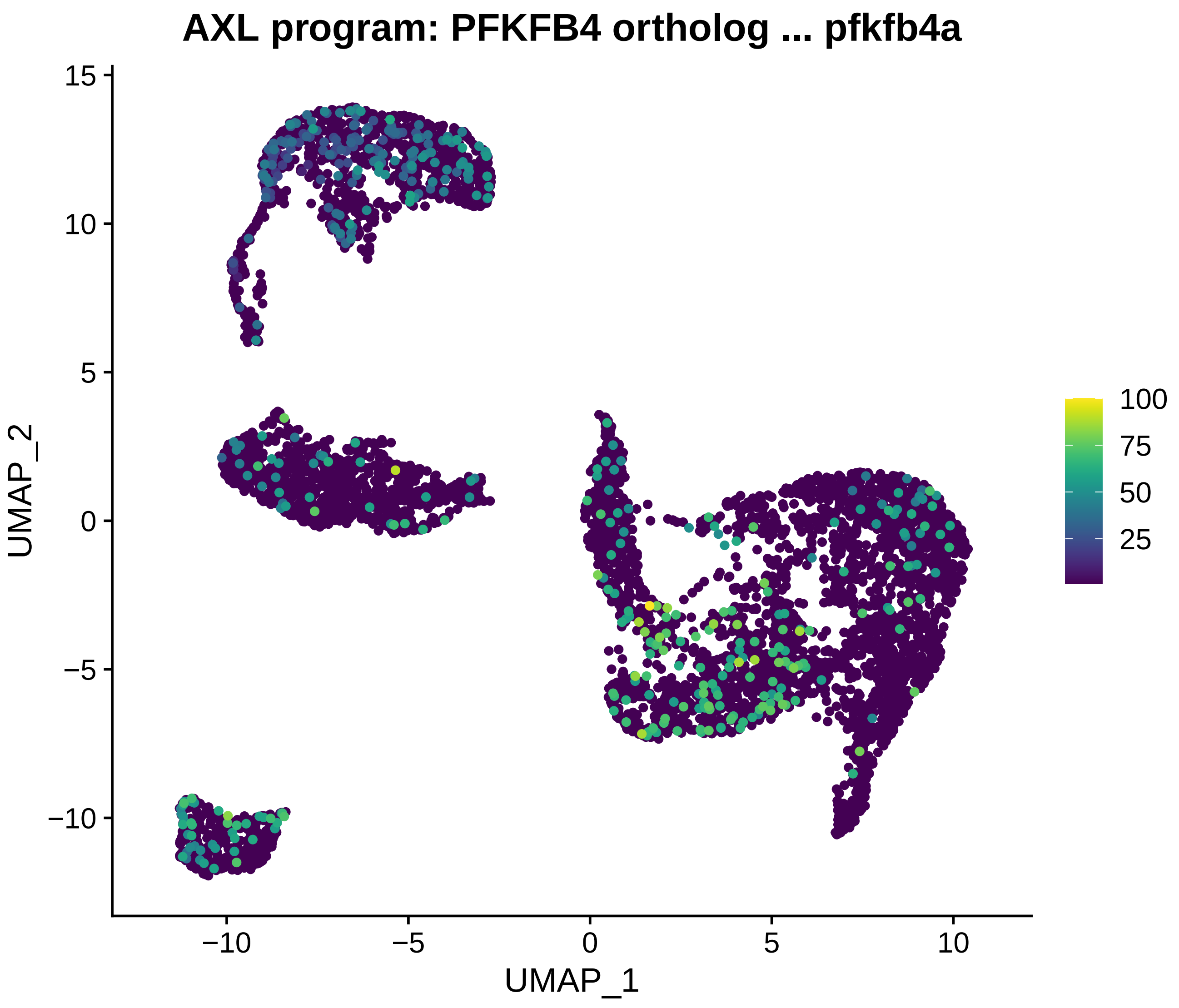

AXL program: PFKFB4 ortholog ... pfkfb4a
UMAP_2
UMAP_1
15
10
5
0
−5
−10
100
75
50
25

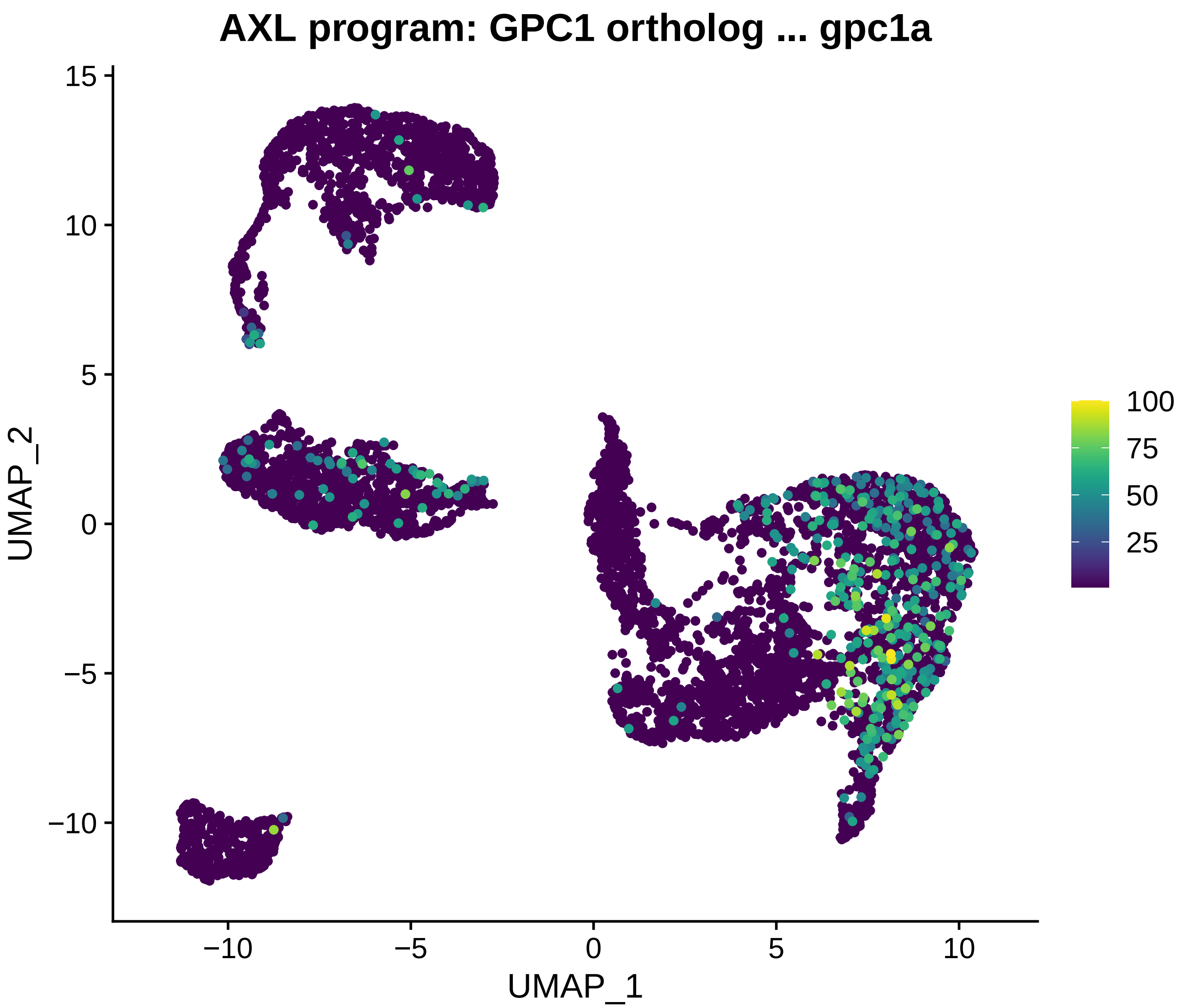
AXL program: GPC1 ortholog ... gpc1a
UMAP_2
UMAP_1
15
10
5
0
−5
−10
−10
−5
0
5
10
100
75
50
25

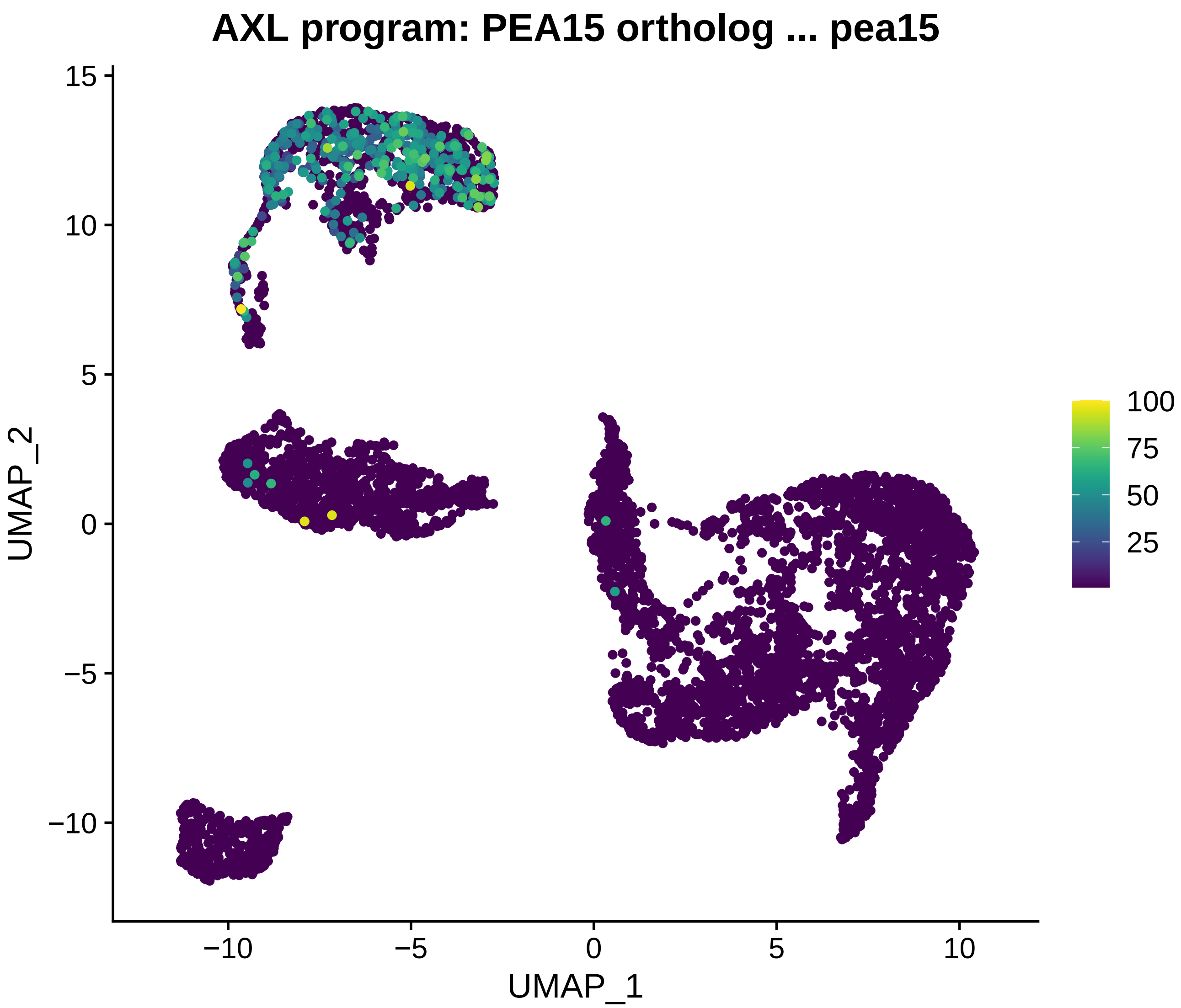
AXL program: PEA15 ortholog ... pea15
UMAP_2
UMAP_1
15
10
5
0
−5
−10
−10
−5
0
5
10
100
75
50
25

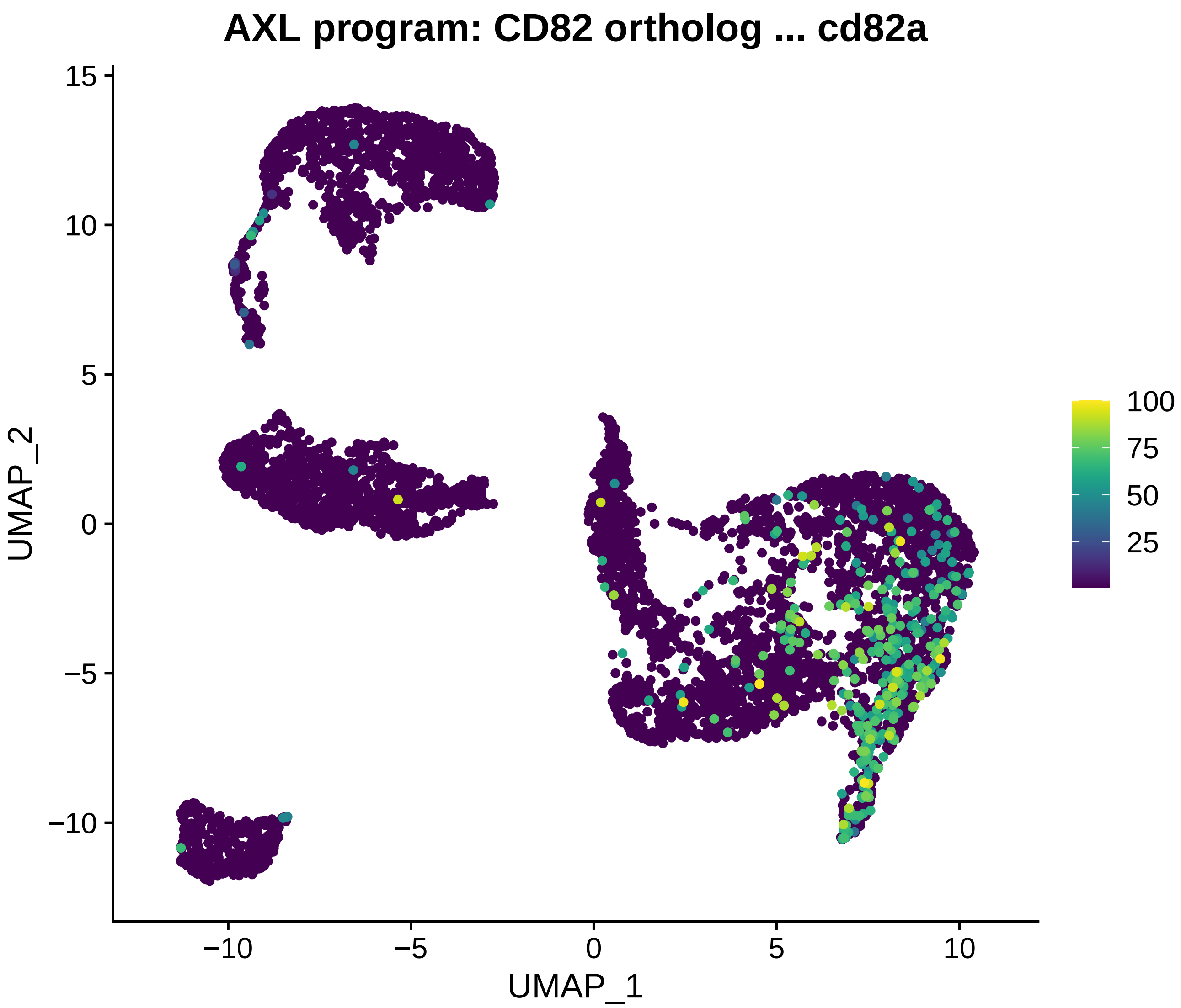
AXL program: CD82 ortholog ... cd82a
UMAP_2
UMAP_1
15
10
5
0
−5
−10
−10
−5
0
5
10
100
75
50
25

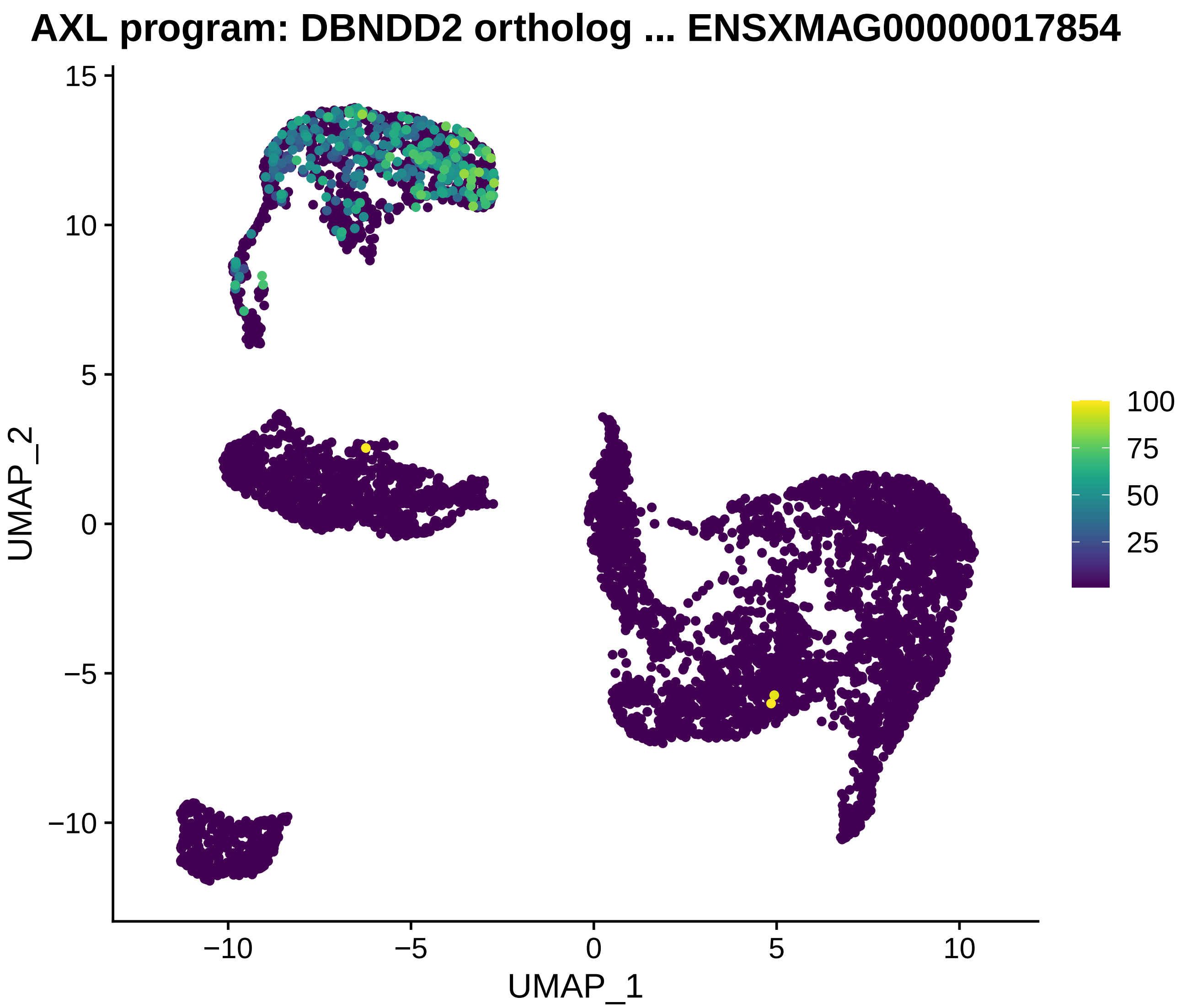

AXL program: DBNDD2 ortholog ... ENSXMAG00000017854
UMAP_2
UMAP_1
15
10
5
0
−5
−10
−10
−5
0
5
10
100
75
50
25

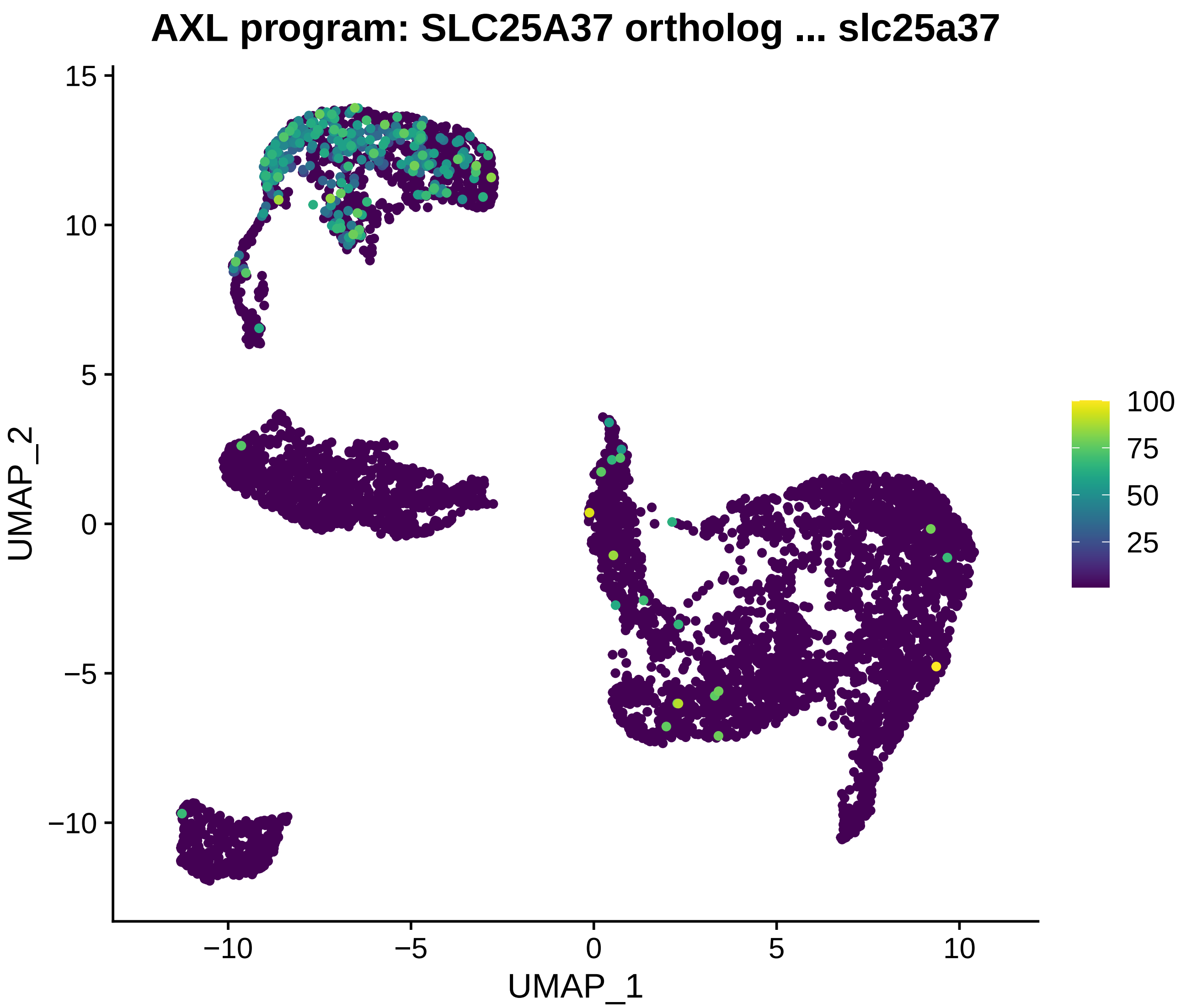
AXL program: SLC25A37 ortholog ... slc25a37
UMAP_1
UMAP_2
15
10
5
0
−5
−10
−10
−5
0
5
10
100
75
50
25

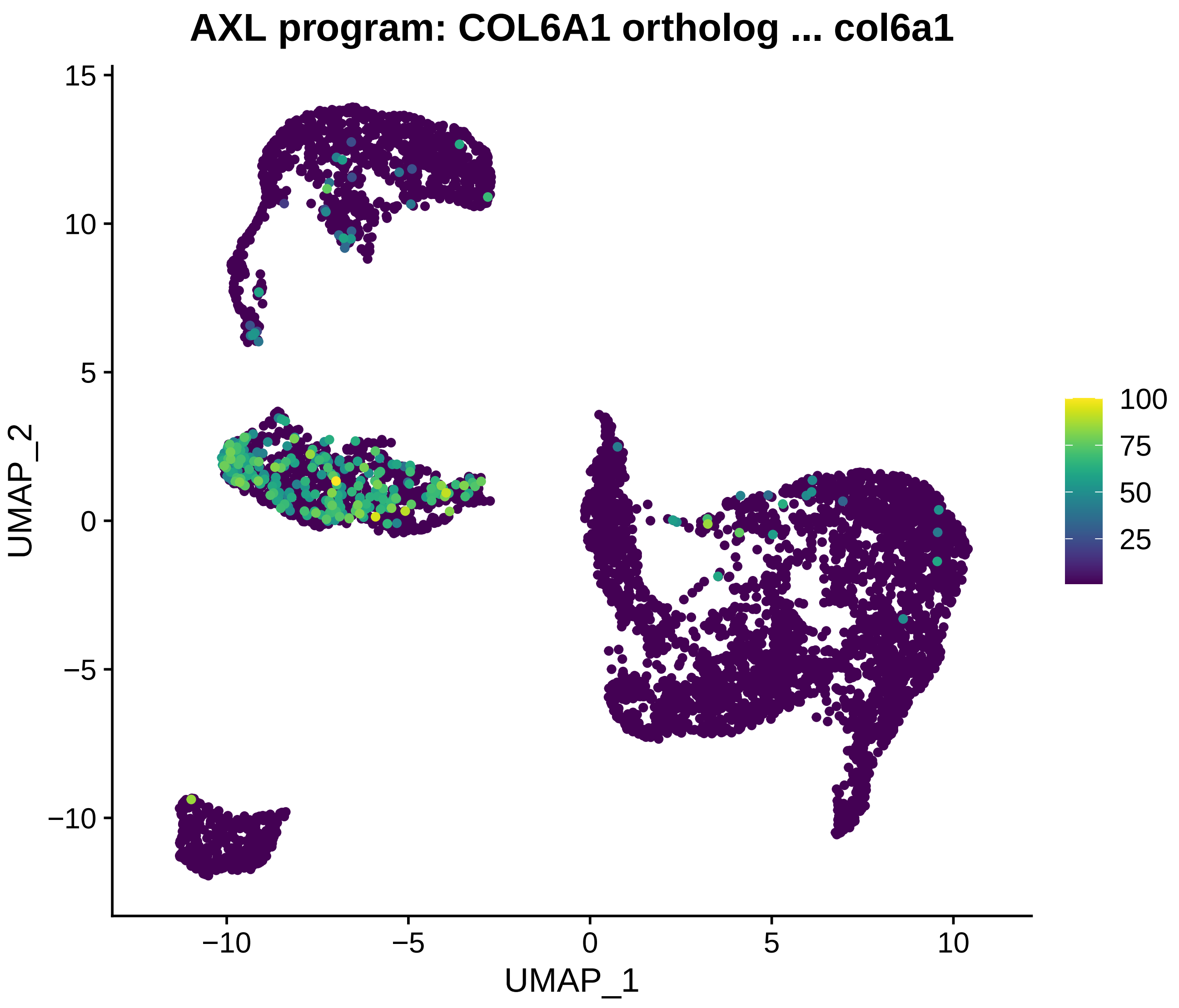
AXL program: COL6A1 ortholog ... col6a1
UMAP_2
UMAP_1
15
10
5
0
−5
−10
−10
−5
0
5
10
100
75
50
25

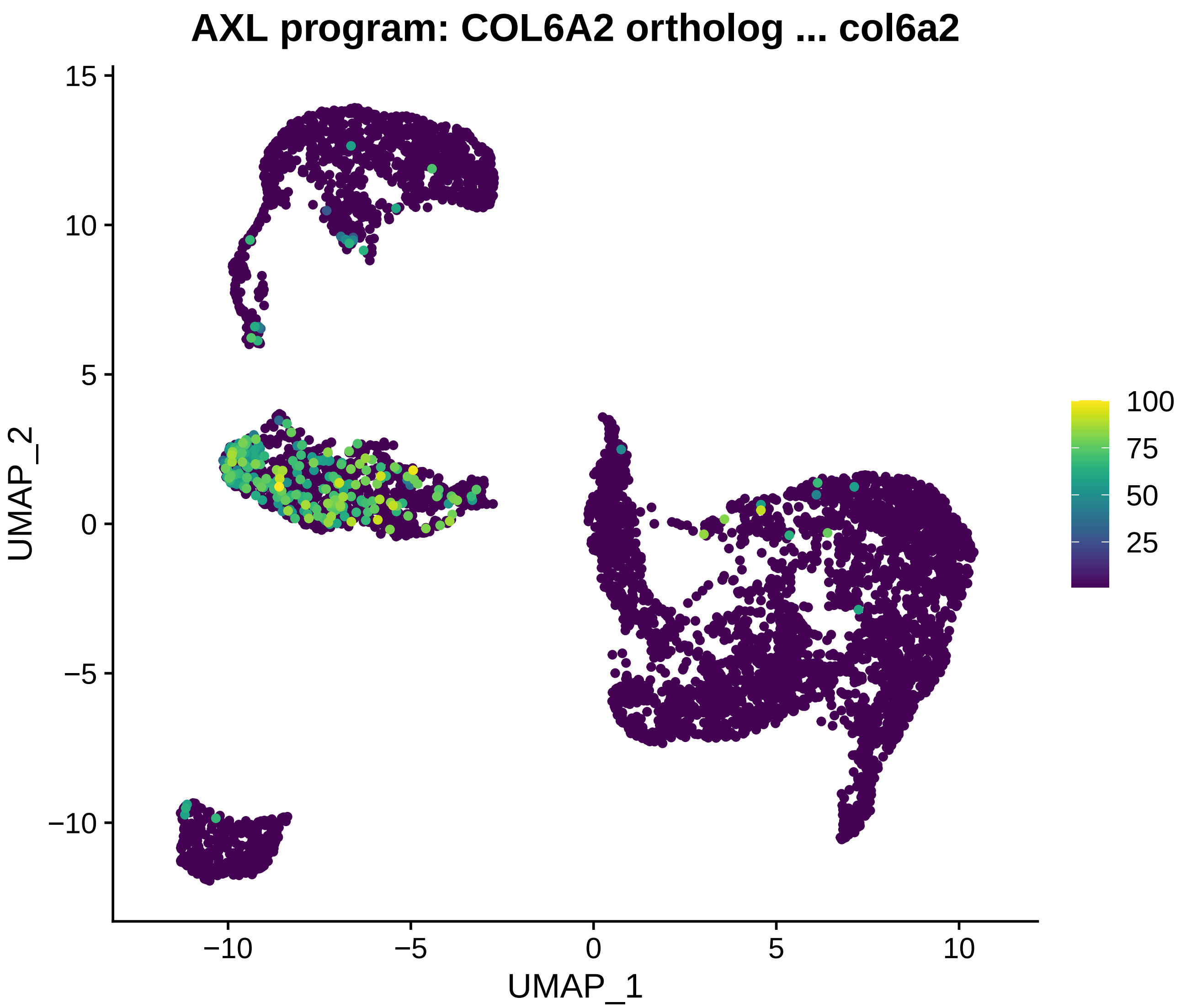
AXL program: COL6A2 ortholog ... col6a2
UMAP_2
UMAP_1
15
10
5
0
−5
−10
−10
−5
0
5
10
100
75
50
25

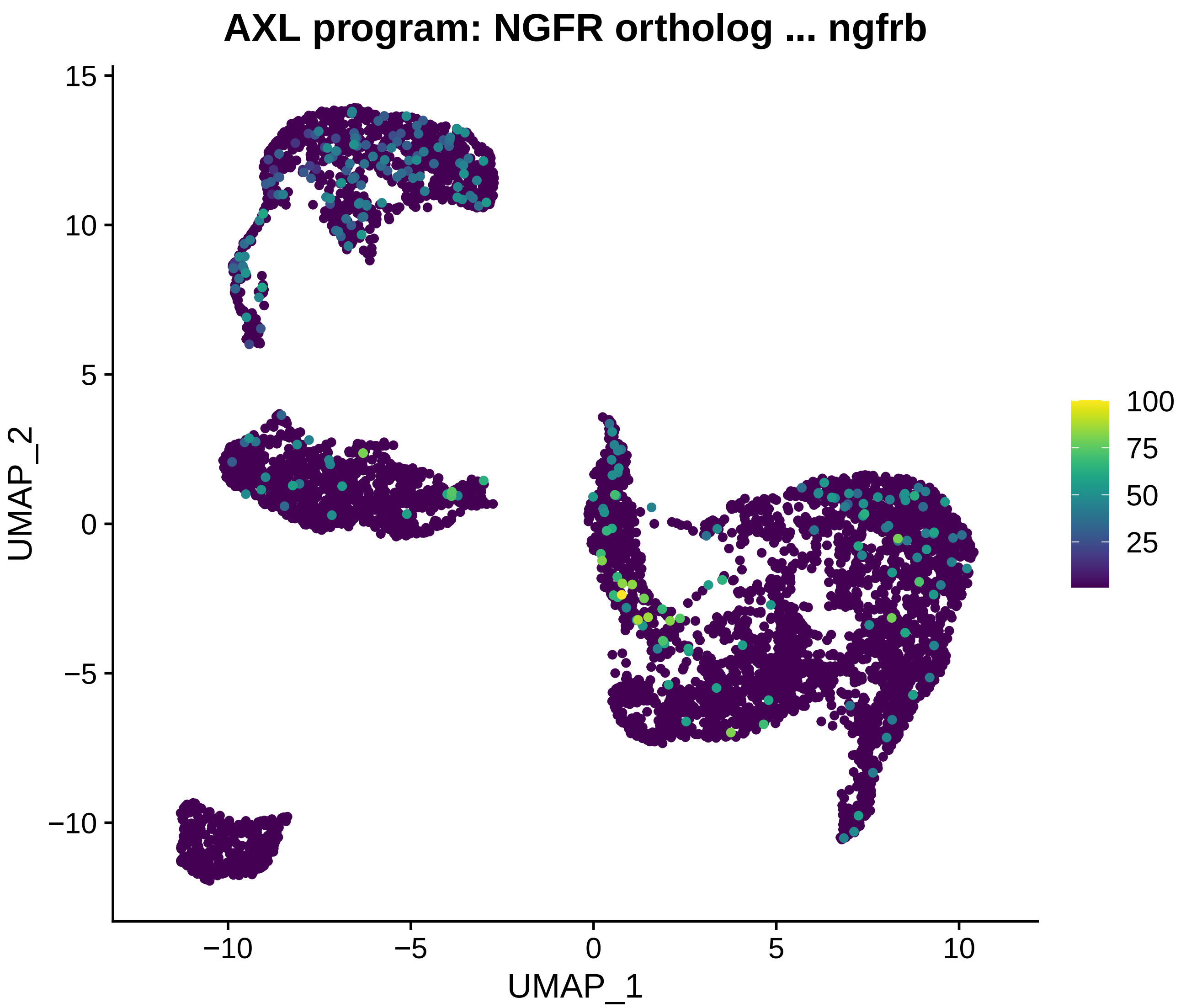

AXL program: NGFR ortholog ... ngfrb
UMAP_2
UMAP_1
15
10
5
0
−5
−10
−10
−5
0
5
10
100
75
50
25

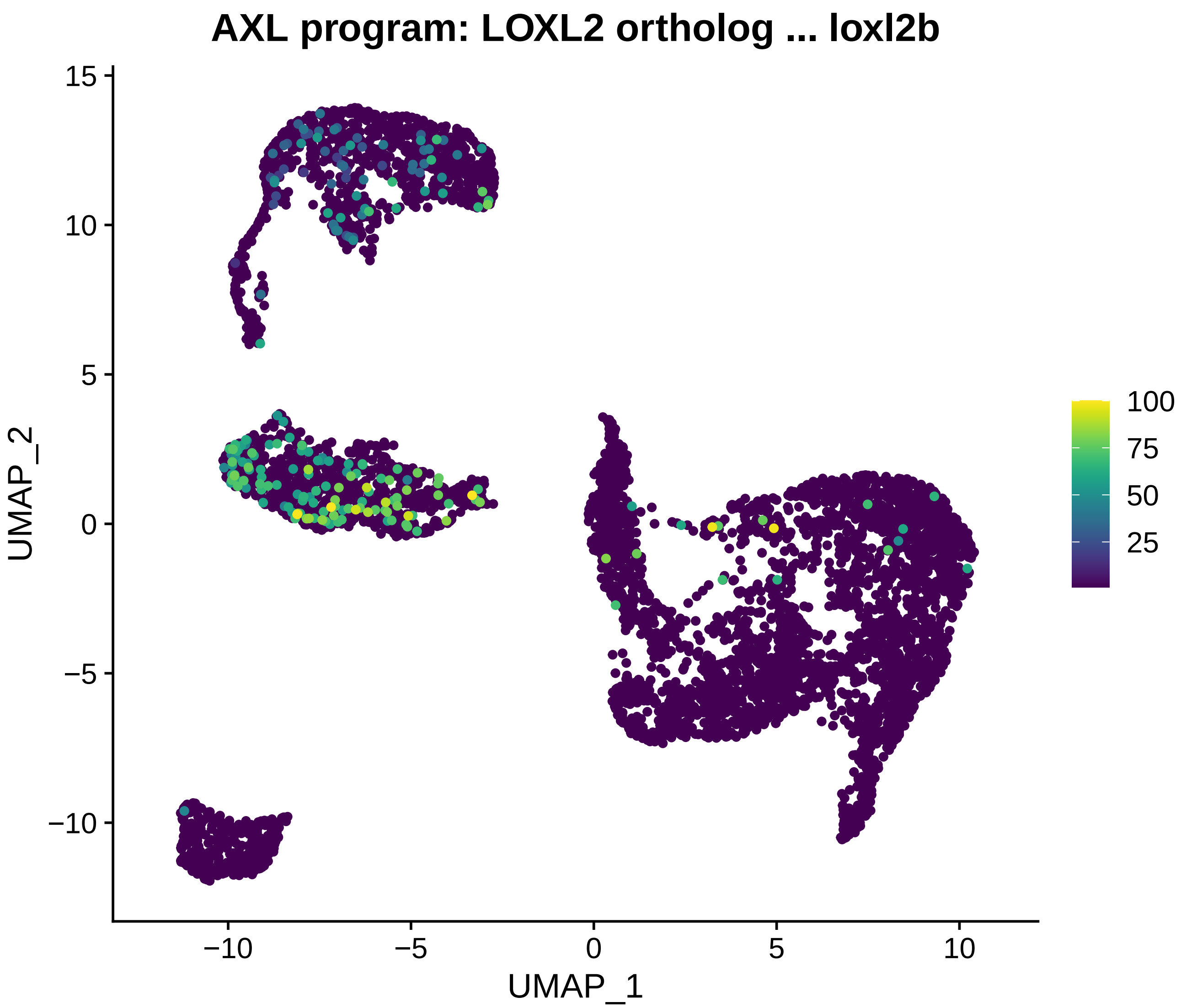
AXL program: LOXL2 ortholog ... loxl2b
UMAP_1
UMAP_2
−10
−5
0
5
10
15
100
75
50
25

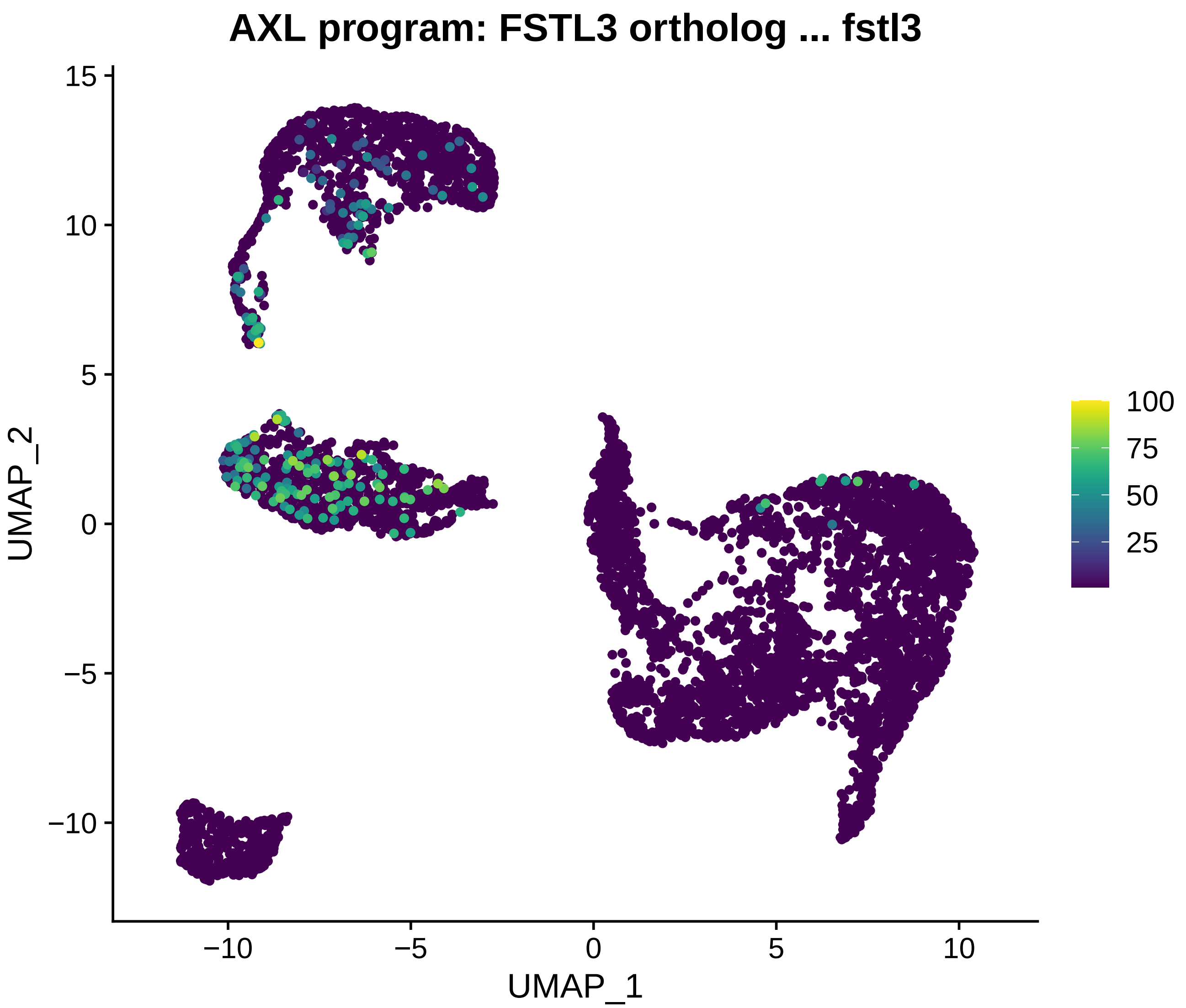
AXL program: FSTL3 ortholog ... fstl3
UMAP_1
UMAP_2
15
10
5
0
−5
−10
−10
−5
0
5
10
100
75
50
25

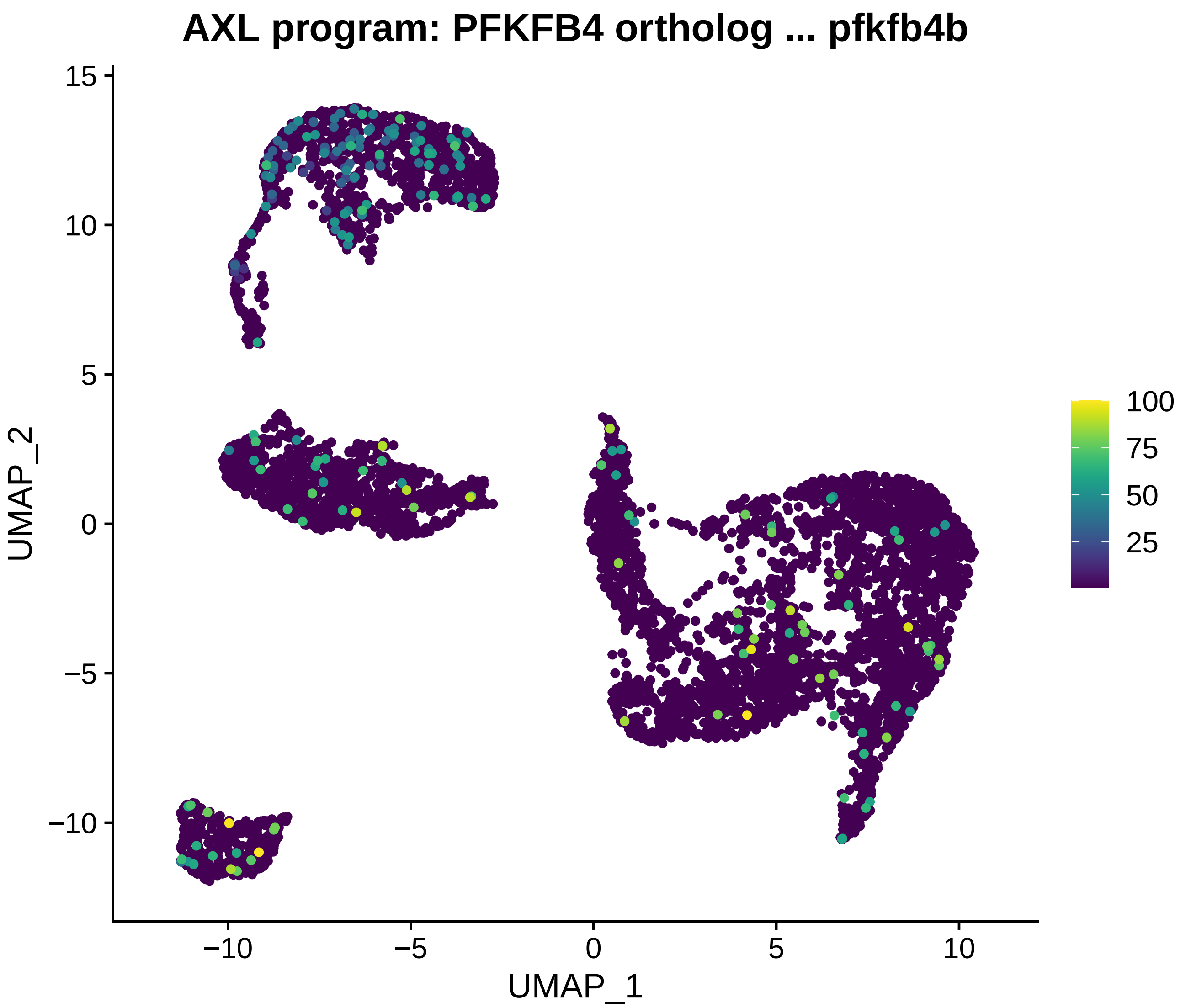
AXL program: PFKFB4 ortholog ... pfkfb4b
UMAP_1
UMAP_2
15
10
5
0
−5
−10
−10
−5
0
5
10
100
75
50
25

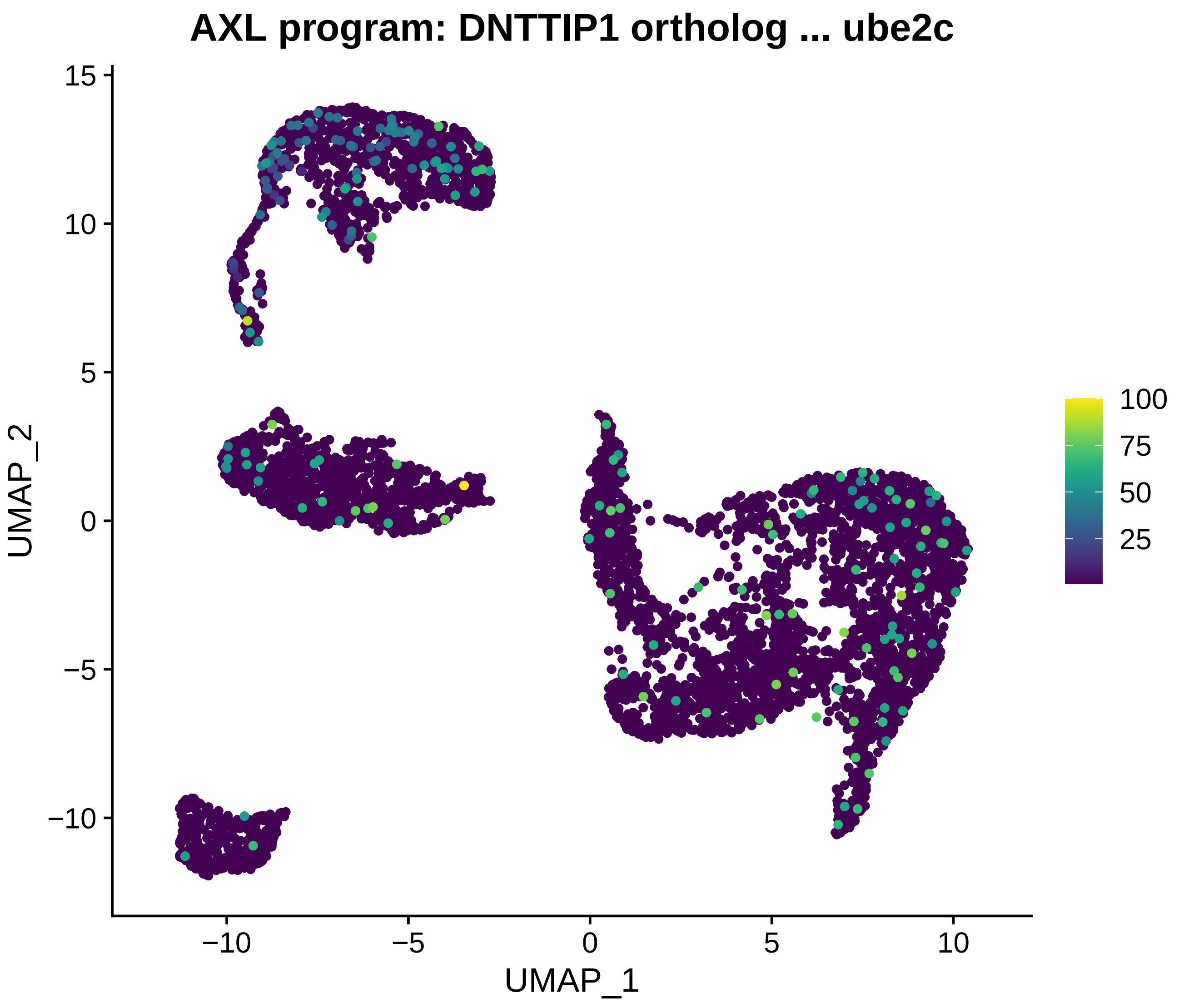

AXL program: DNTTIP1 ortholog ... ube2c
UMAP_2
UMAP_1
15
10
5
0
−5
−10
−10
−5
0
5
10
100
75
50
25

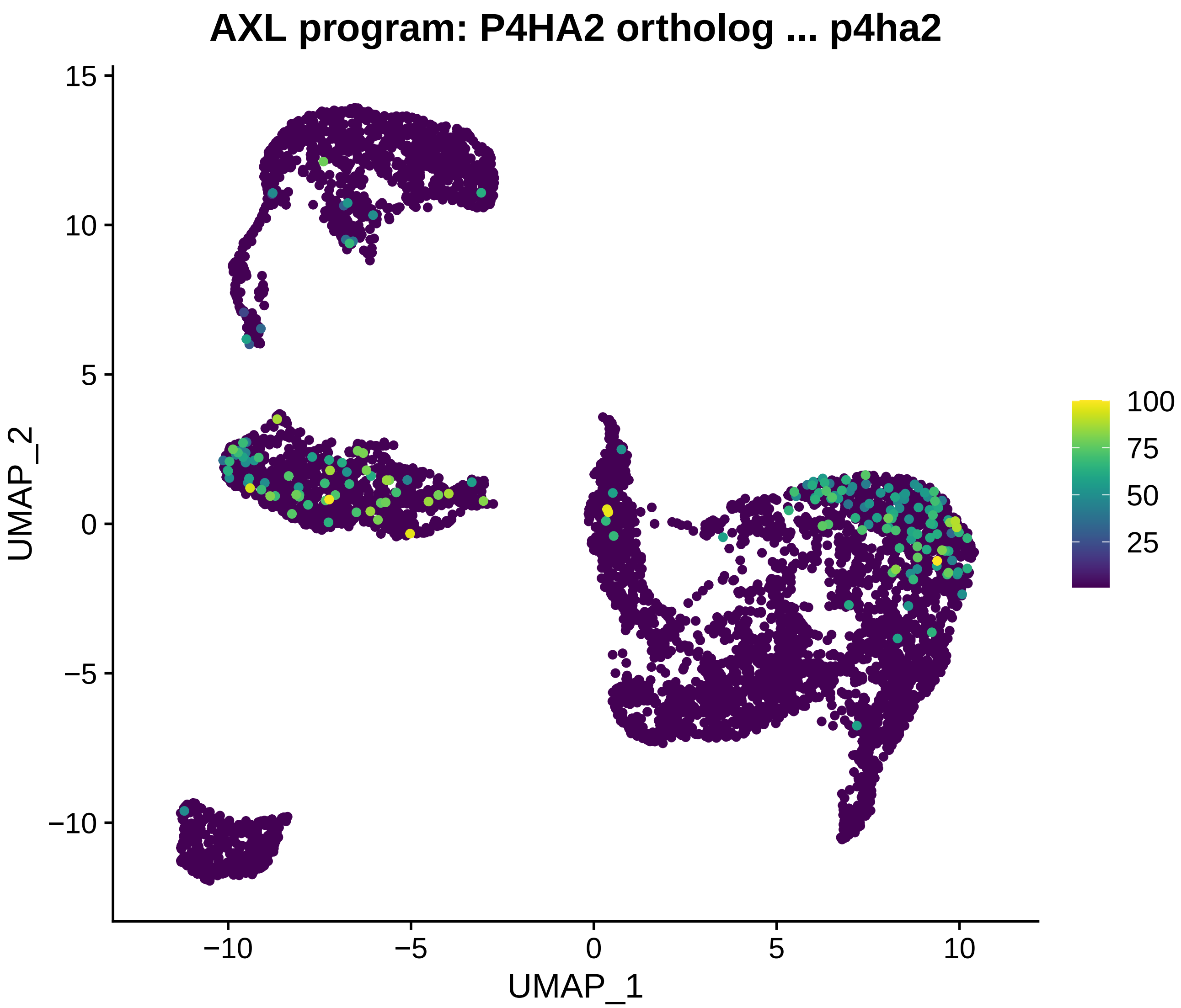
AXL program: P4HA2 ortholog ... p4ha2
UMAP_2
UMAP_1
15
10
5
0
−5
−10
−10
−5
0
5
10
100
75
50
25

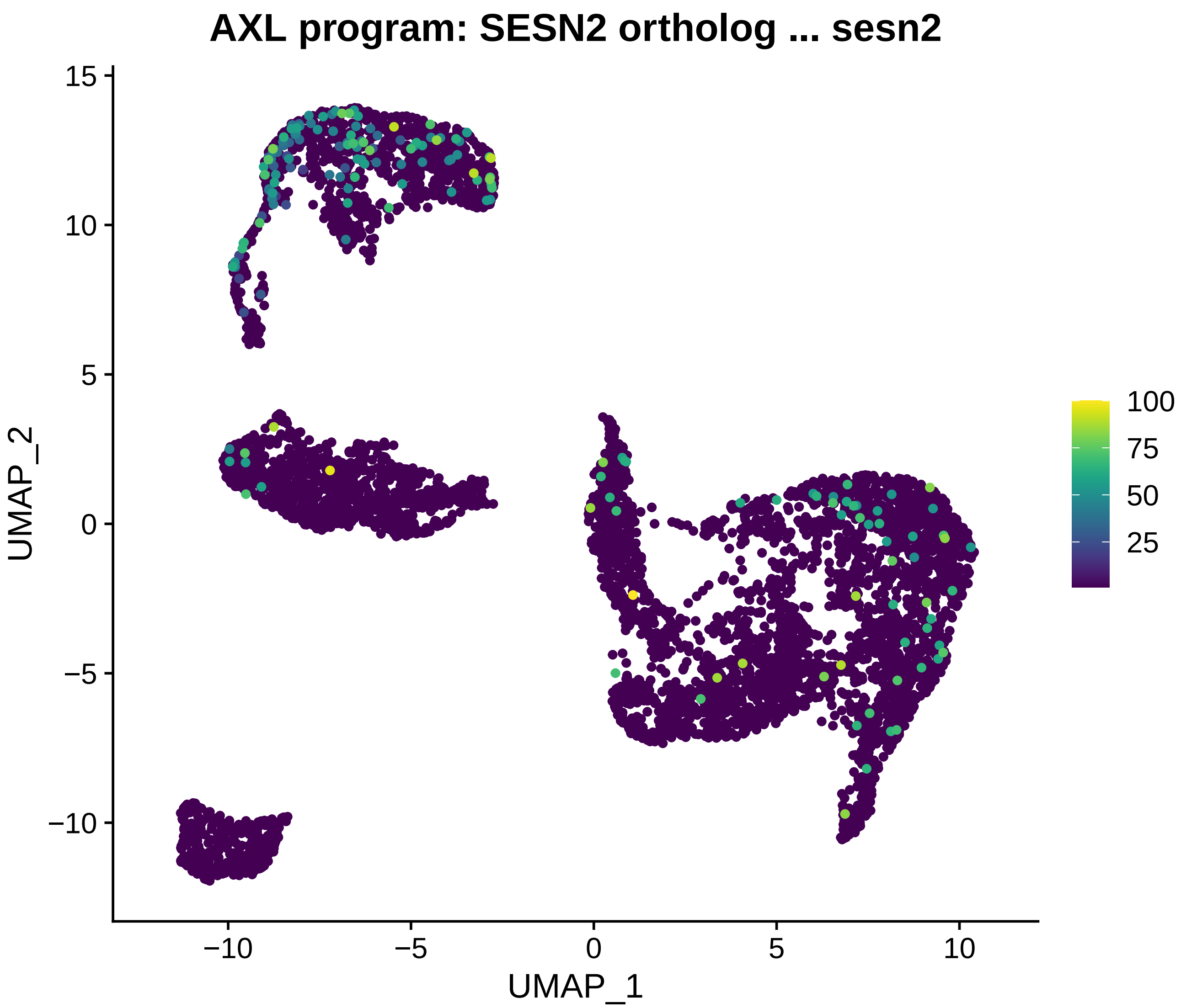
AXL program: SESN2 ortholog ... sesn2
UMAP_1
UMAP_2
−10
−5
0
5
10
15
100
75
50
25

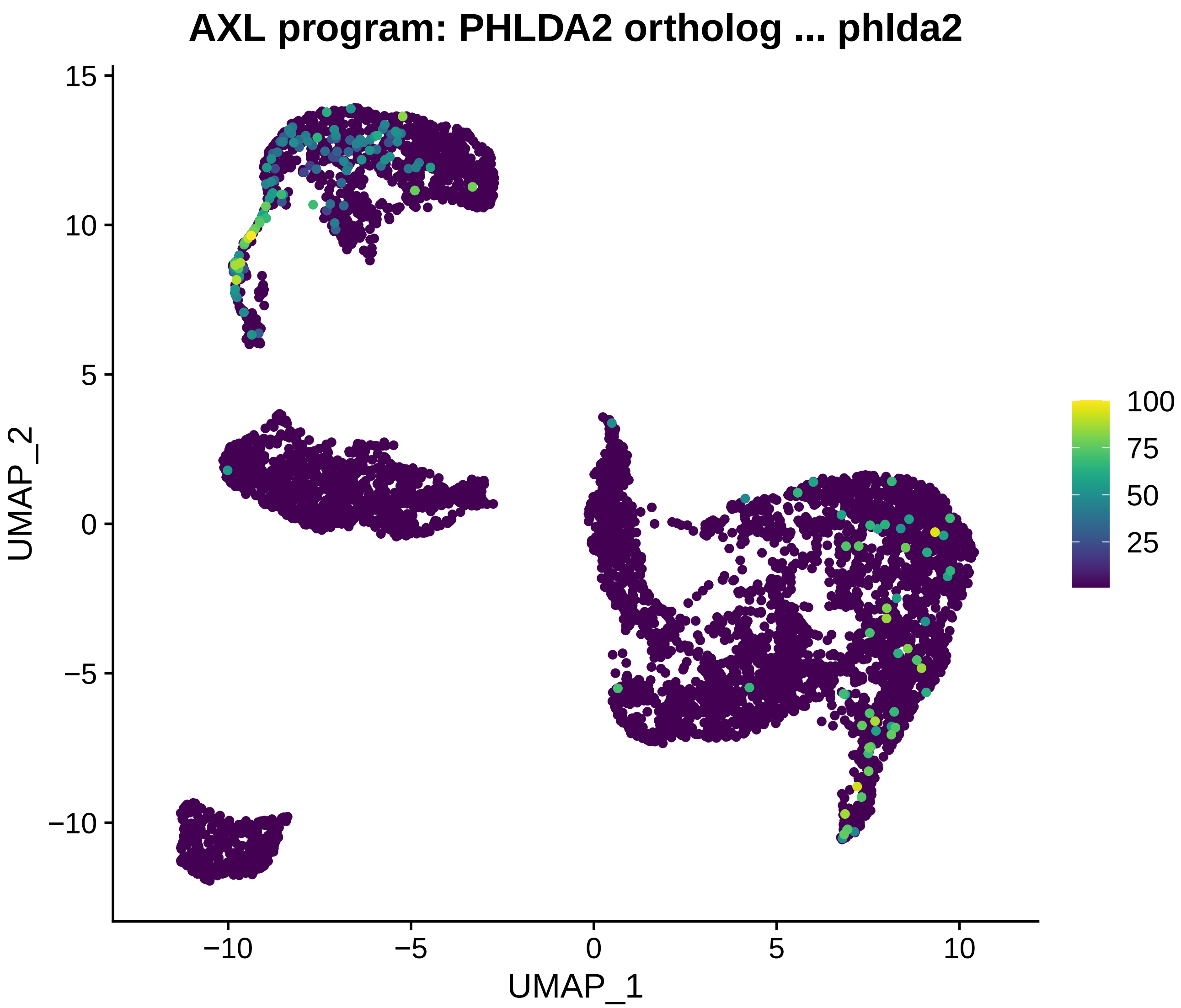
AXL program: PHLDA2 ortholog ... phlda2
UMAP_2
UMAP_1
15
10
5
0
−5
−10
−10
−5
0
5
10
100
75
50
25

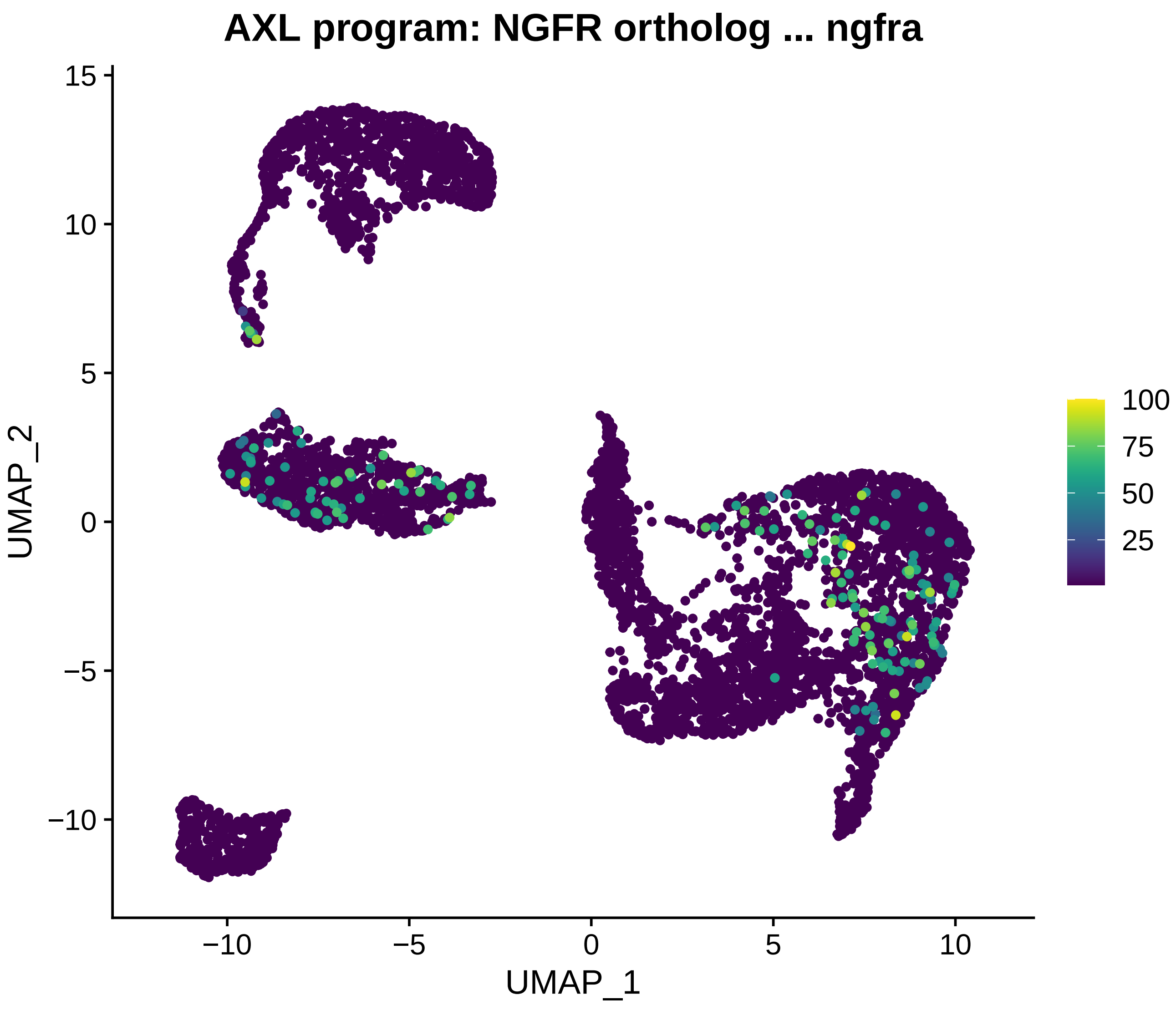
AXL program: NGFR ortholog ... ngfra
UMAP_2
UMAP_1
15
10
5
0
−5
−10
−10
−5
0
5
10
100
75
50
25

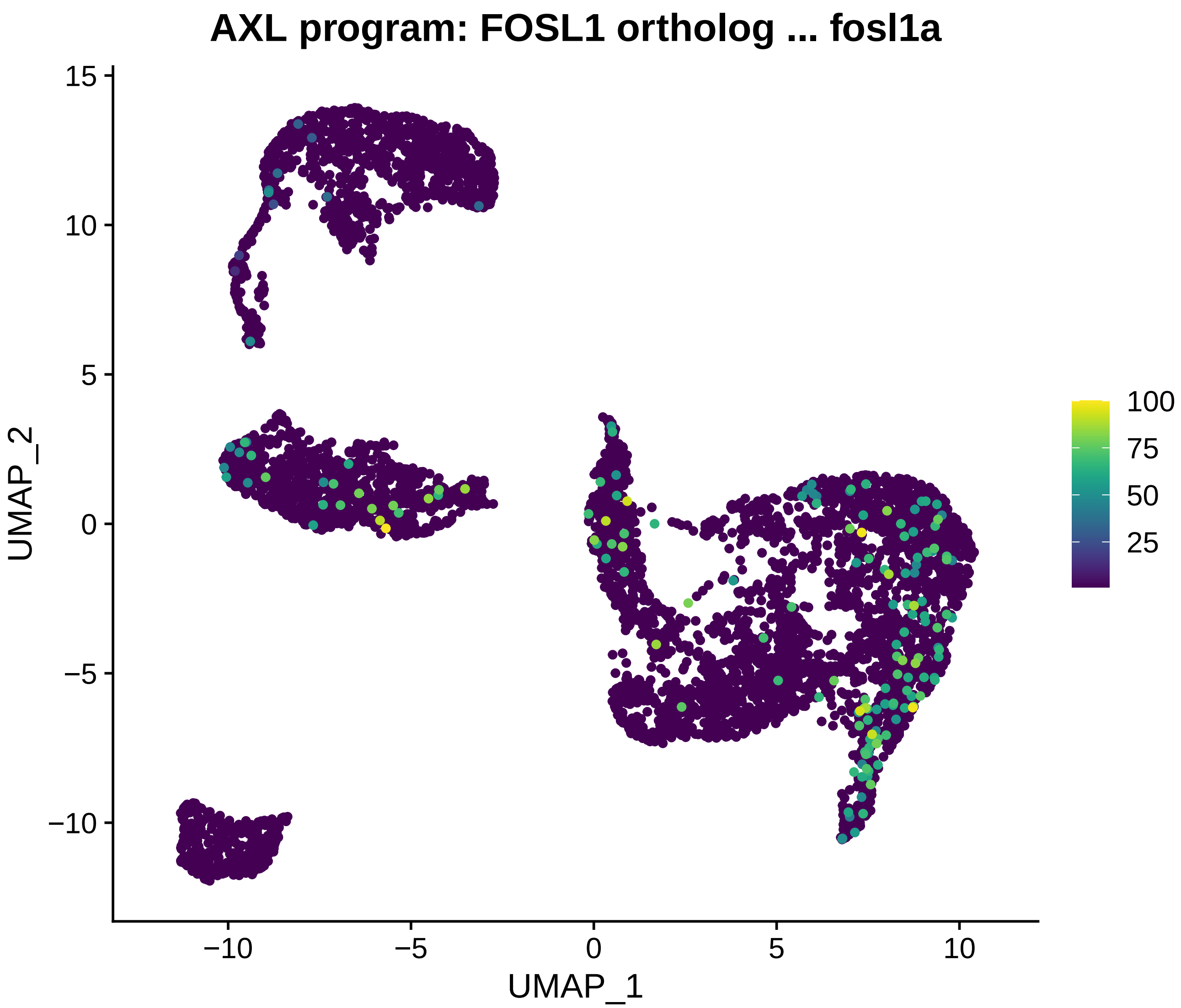

AXL program: FOSL1 ortholog ... fosl1a
UMAP_2
UMAP_1
15
10
5
0
−5
−10
−10
−5
0
5
10
100
75
50
25

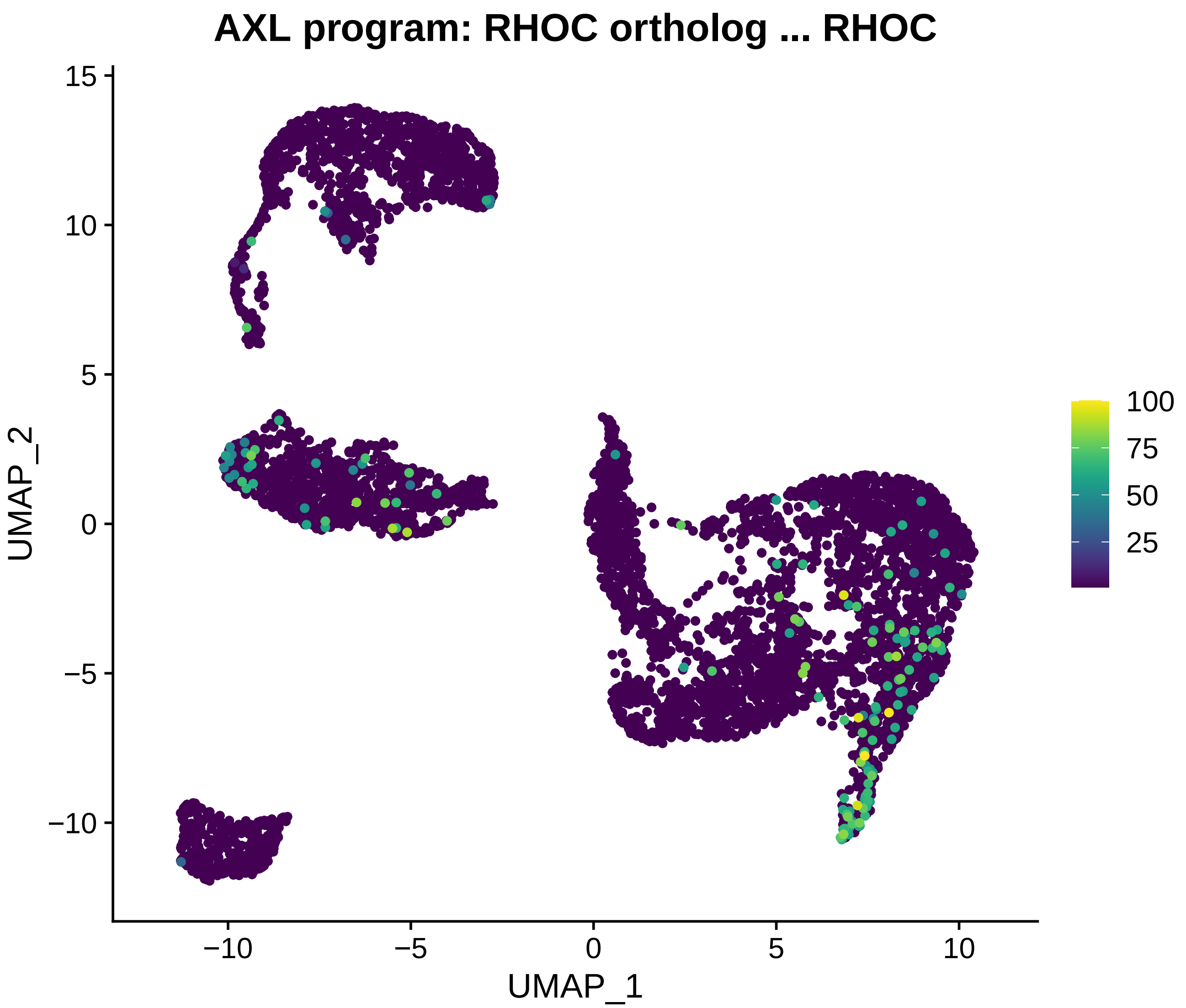
AXL program: RHOC ortholog ... RHOC
UMAP_2
UMAP_1
15
10
5
0
−5
−10
−10
−5
0
5
10
100
75
50
25

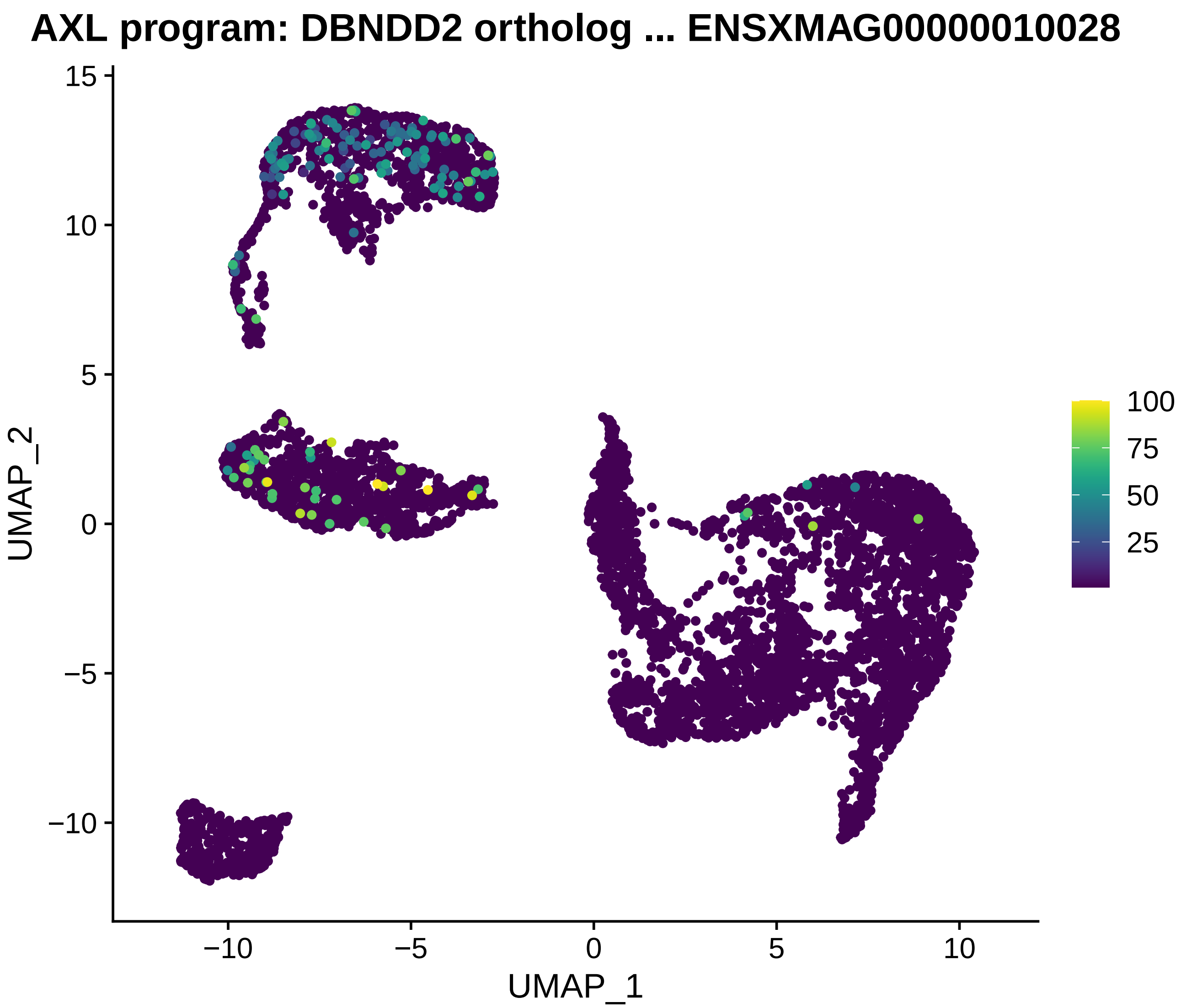
AXL program: DBNDD2 ortholog ... ENSXMAG00000010028
UMAP_2
UMAP_1
15
10
5
0
−5
−10
−10
−5
0
5
10
100
75
50
25

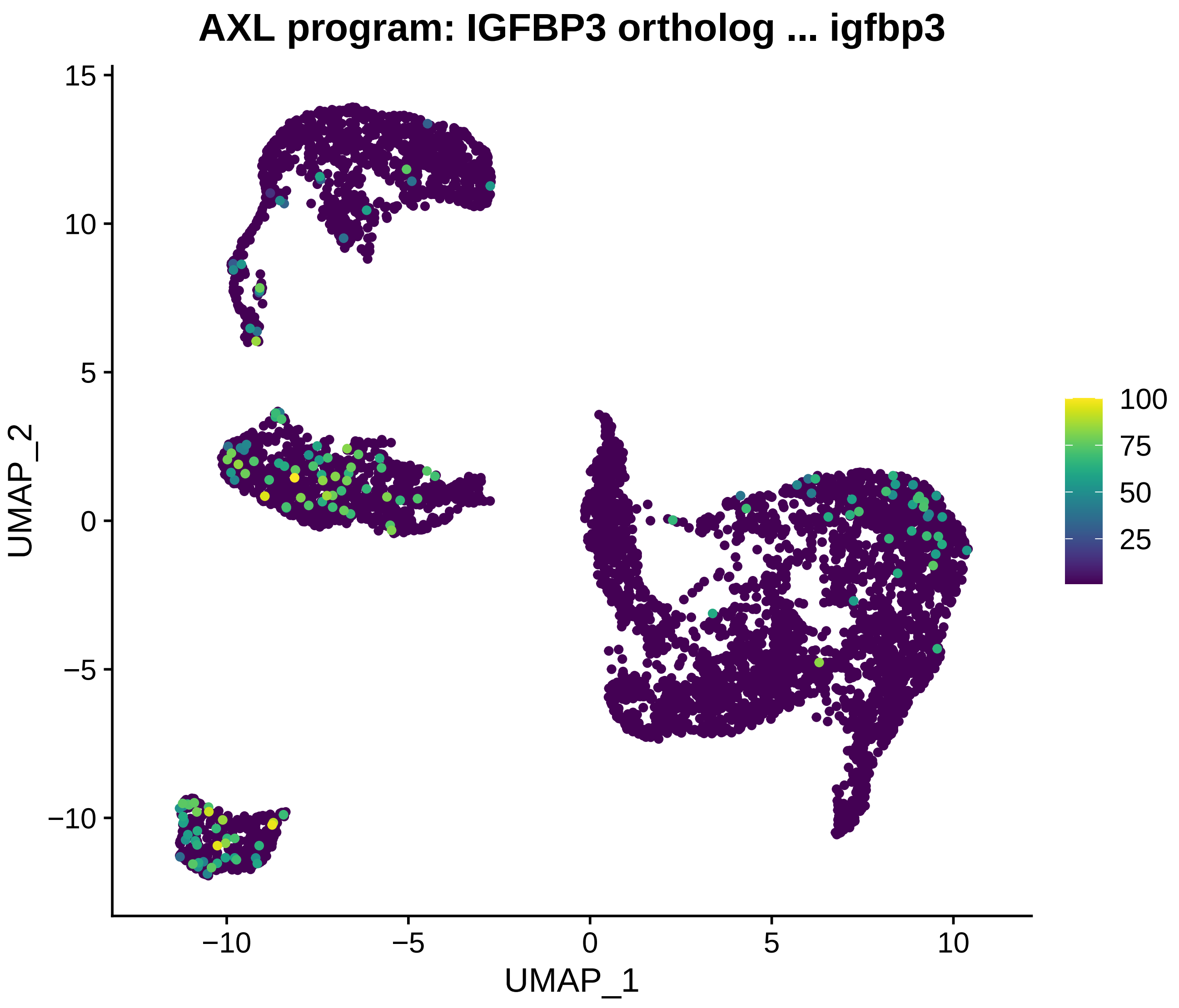
AXL program: IGFBP3 ortholog ... igfbp3
UMAP_2
UMAP_1
15
10
5
0
−5
−10
−10
−5
0
5
10
100
75
50
25

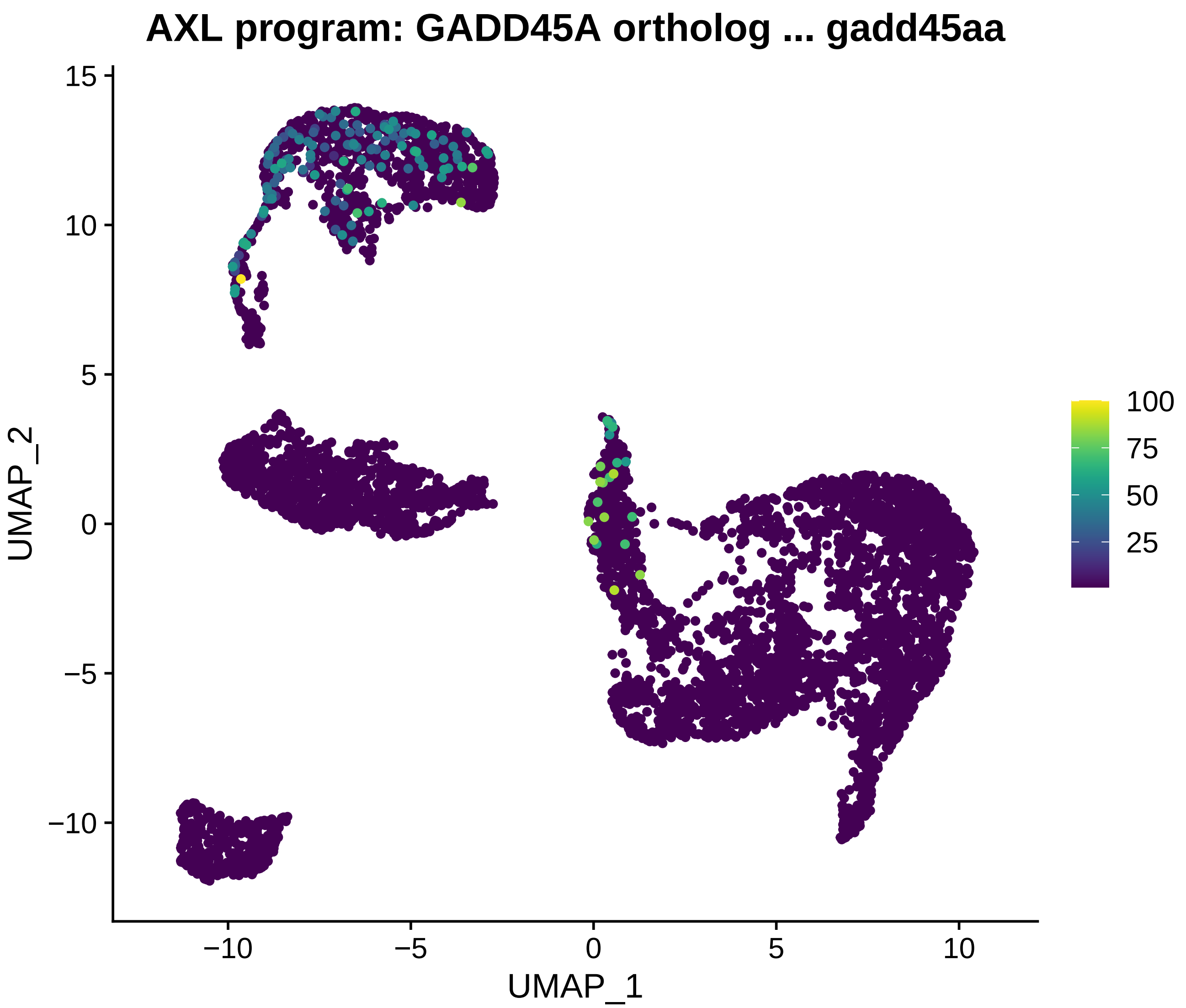
AXL program: GADD45A ortholog ... gadd45aa
15
10
5
0
−5
−10
UMAP_2
−10
−5
0
5
10
UMAP_1
100
75
50
25

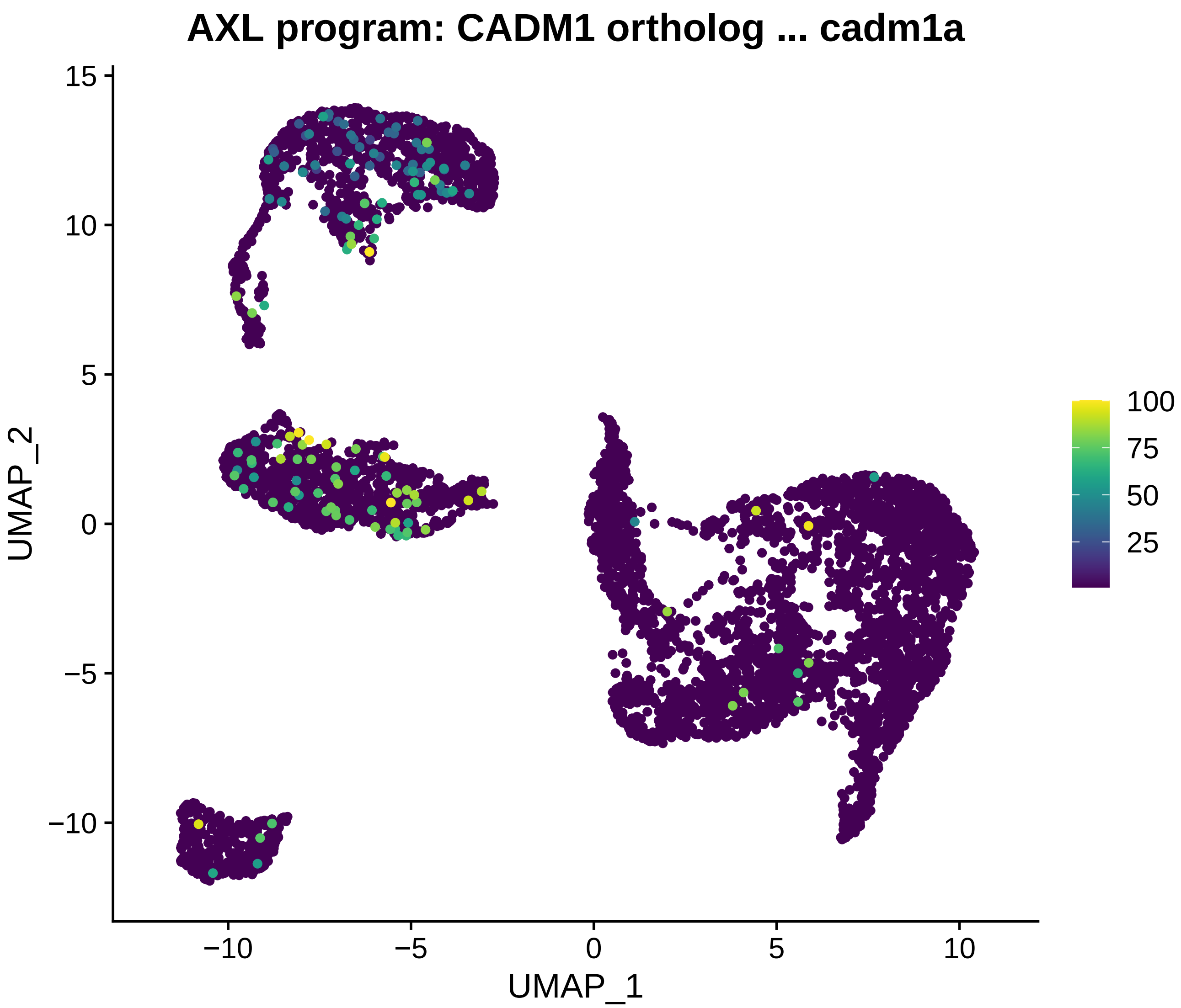

AXL program: CADM1 ortholog ... cadm1a
UMAP_2
UMAP_1
15
10
5
0
−5
−10
−10
−5
0
5
10
100
75
50
25

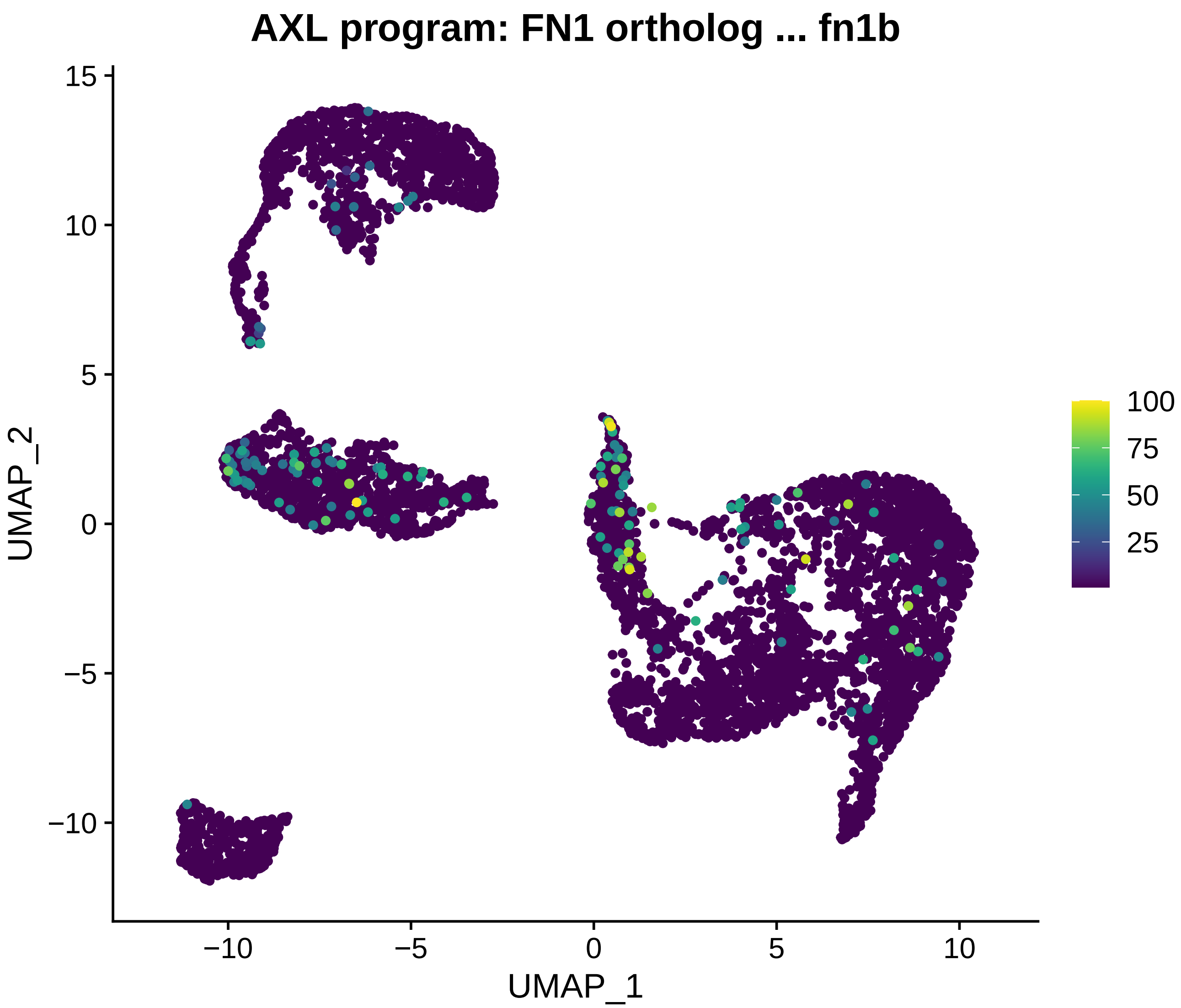

AXL program: FN1 ortholog ... fn1b
15
10
5
0
−5
−10
UMAP_2
−10
−5
0
5
10
UMAP_1
100
75
50
25

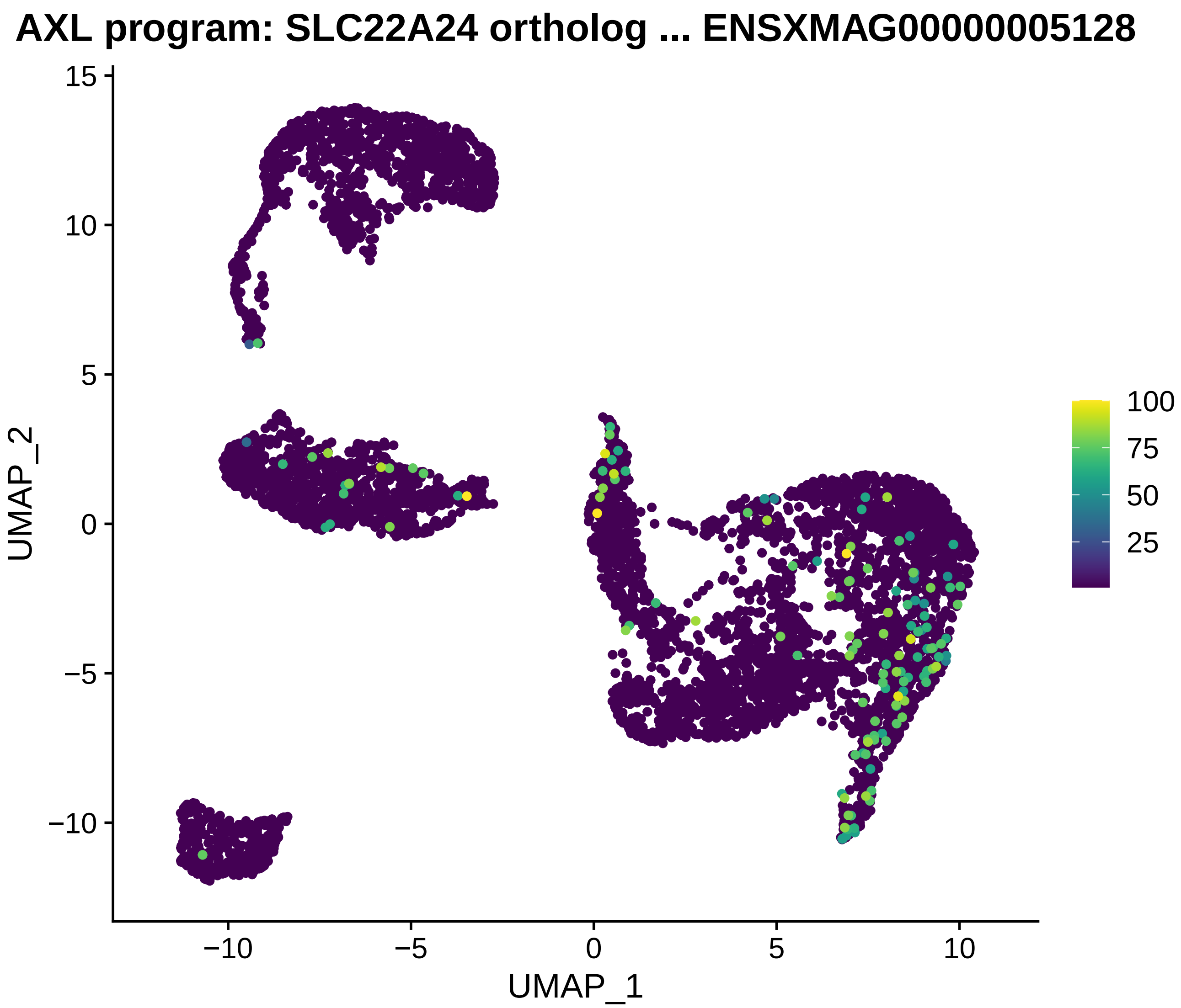
AXL program: SLC22A24 ortholog ... ENSXMAG00000005128
UMAP_2
UMAP_1
15
10
5
0
−5
−10
−10
−5
0
5
10
100
75
50
25

#### AXL program: UBE2J1 ortholog ... ube2j1

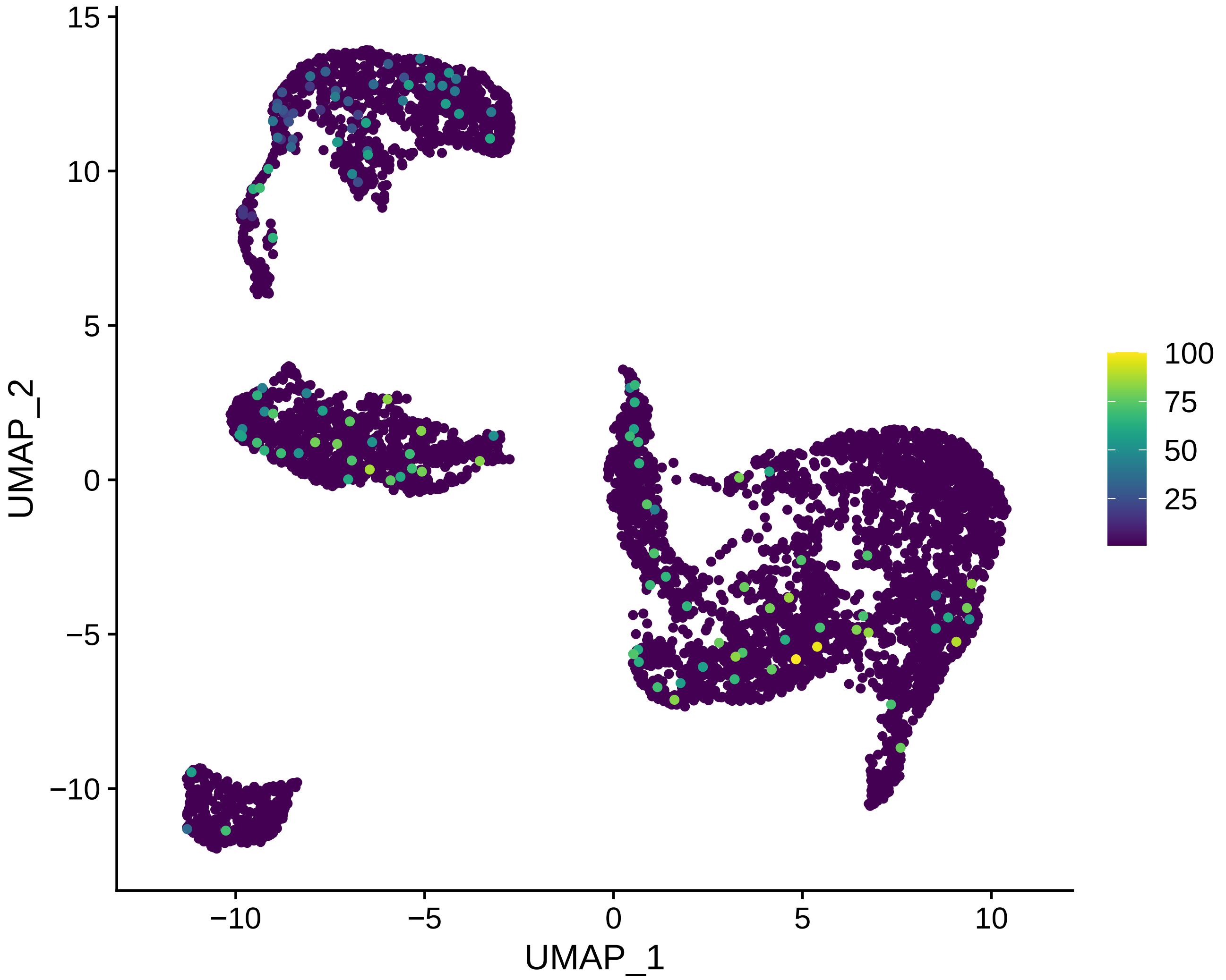

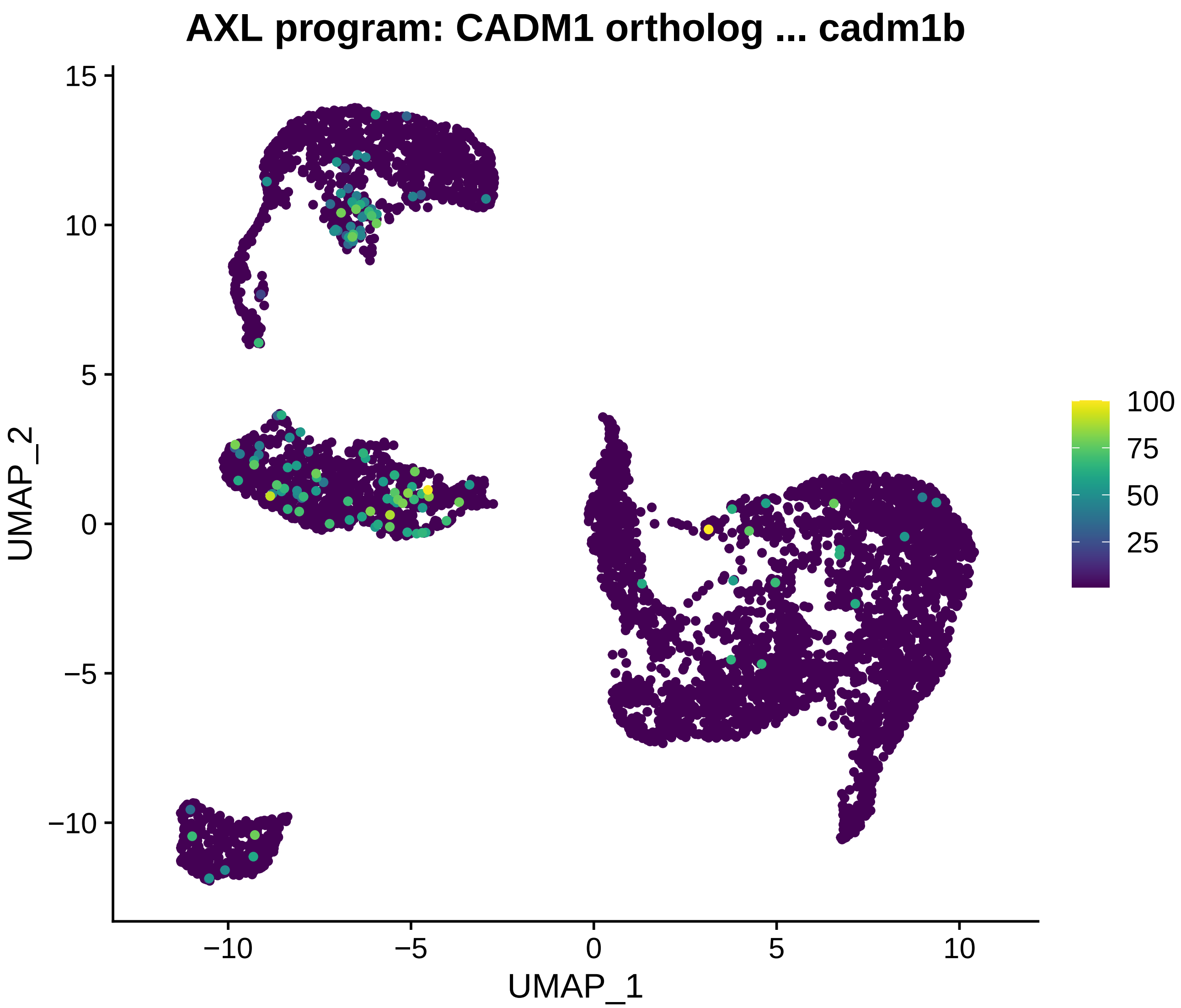
AXL program: CADM1 ortholog ... cadm1b
UMAP_2
UMAP_1
15
10
5
0
−5
−10
−10
−5
0
5
10
100
75
50
25

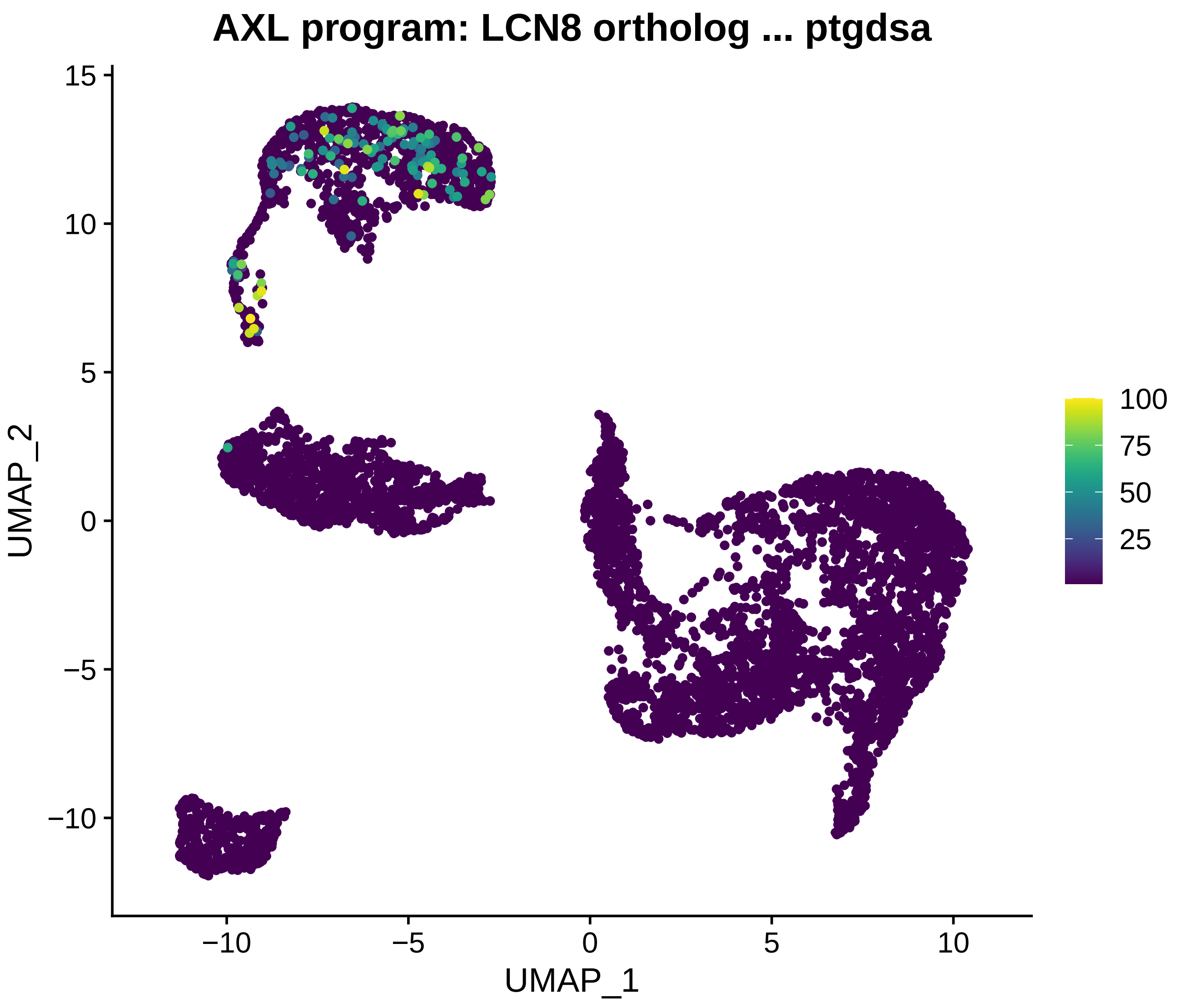
AXL program: LCN8 ortholog ... ptgdsa
UMAP_2
UMAP_1
15
10
5
0
−5
−10
−10
−5
0
5
10
100
75
50
25

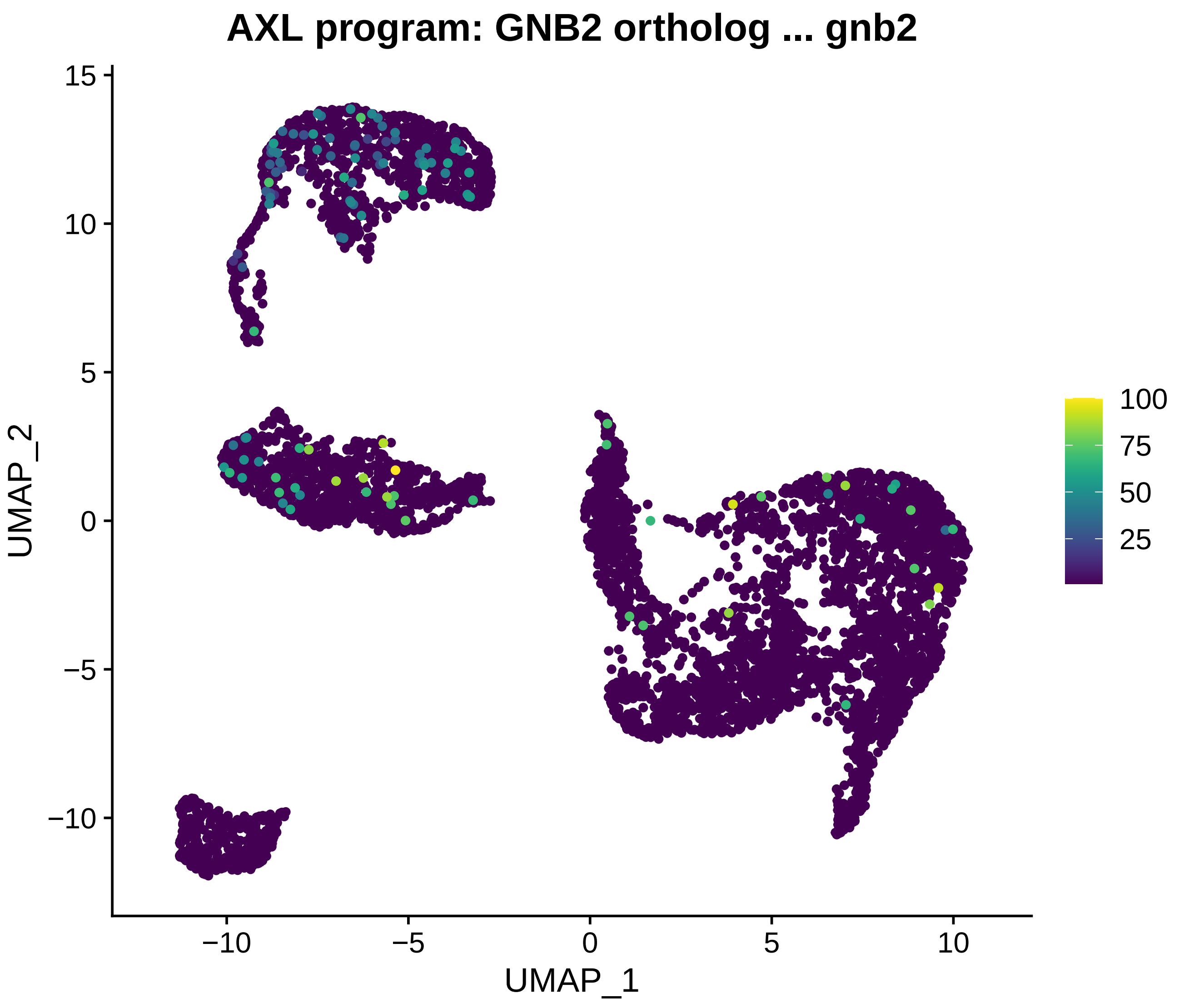

AXL program: GNB2 ortholog ... gnb2
UMAP_2
UMAP_1
15
10
5
0
−5
−10
−10
−5
0
5
10
100
75
50
25

**AXL program: BACH1 ortholog ... ENSXMAG00000026281**

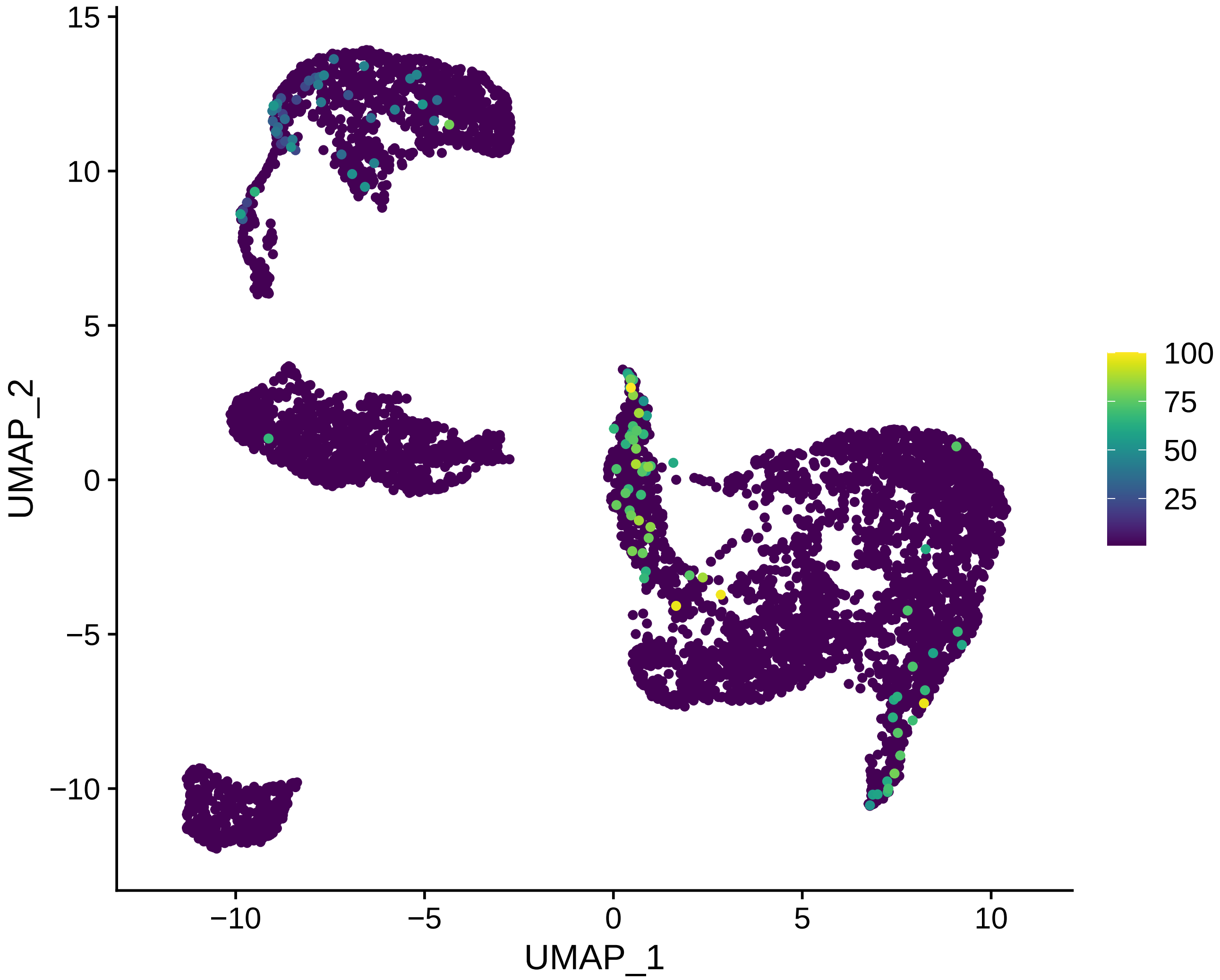

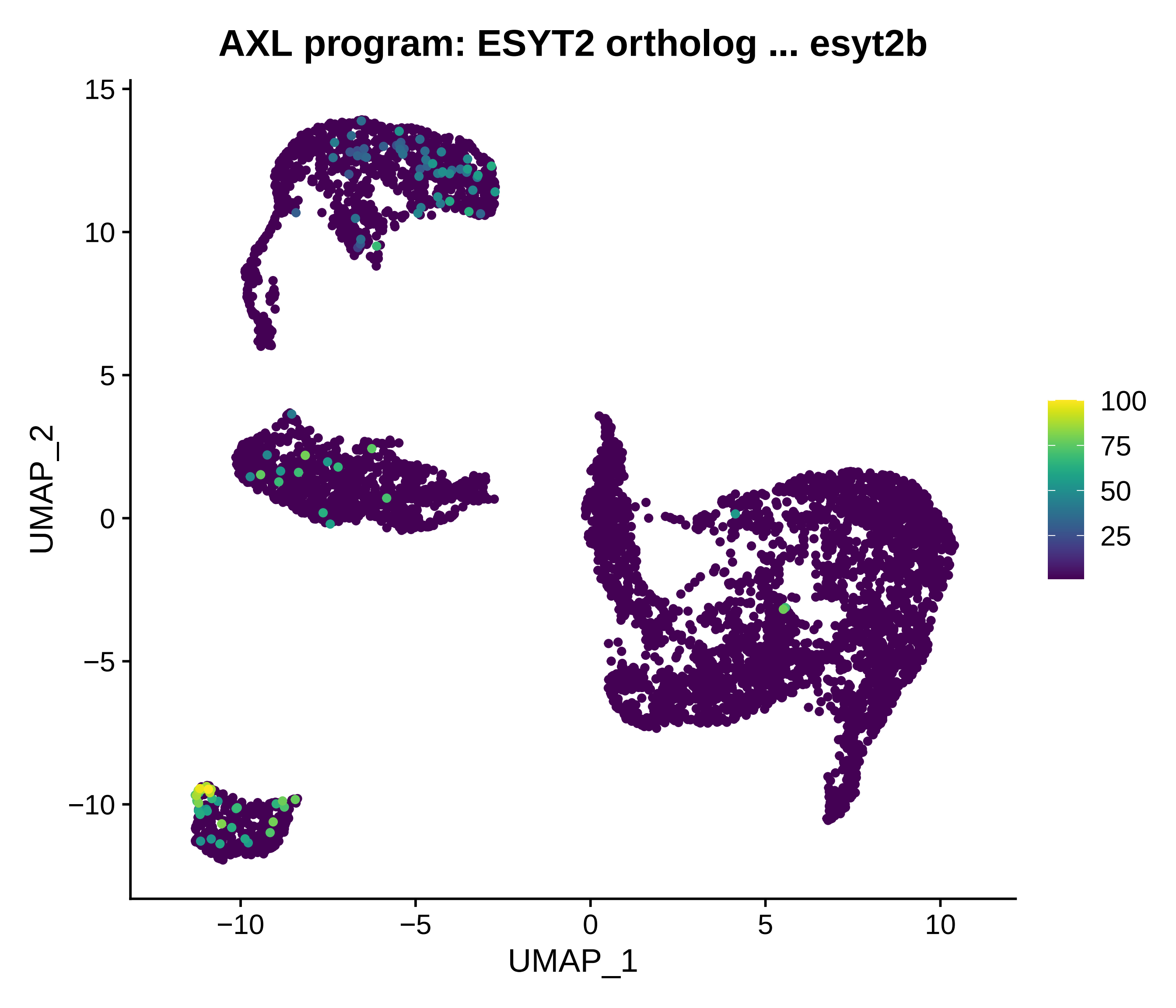
AXL program: ESYT2 ortholog ... esyt2b
UMAP_2
15
10
5
0
−5
−10
−10
−5
0
5
10
UMAP_1
100
75
50
25

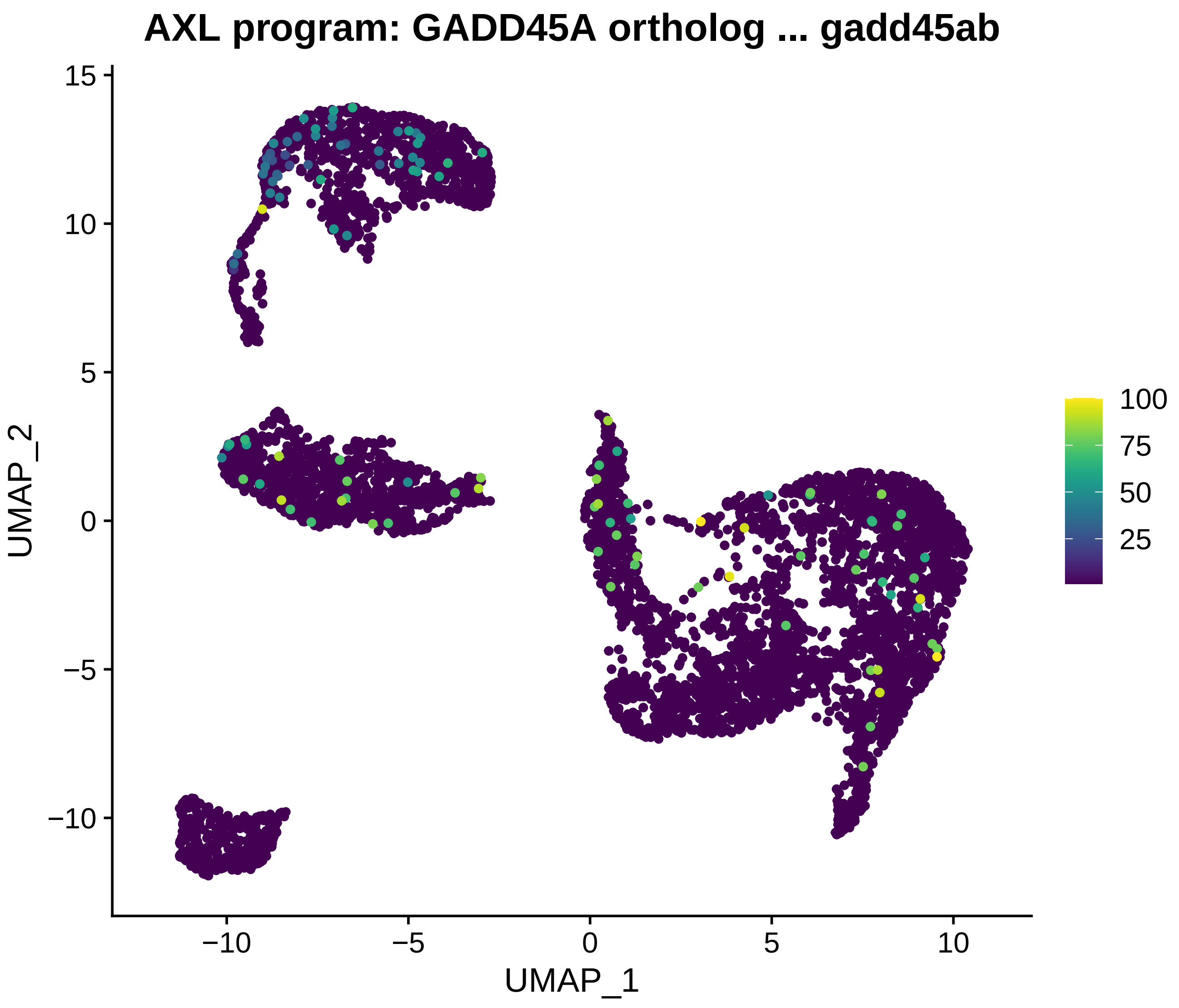

AXL program: GADD45A ortholog ... gadd45ab
UMAP_2
UMAP_1
15
10
5
0
−5
−10
−10
−5
0
5
10
100
75
50
25

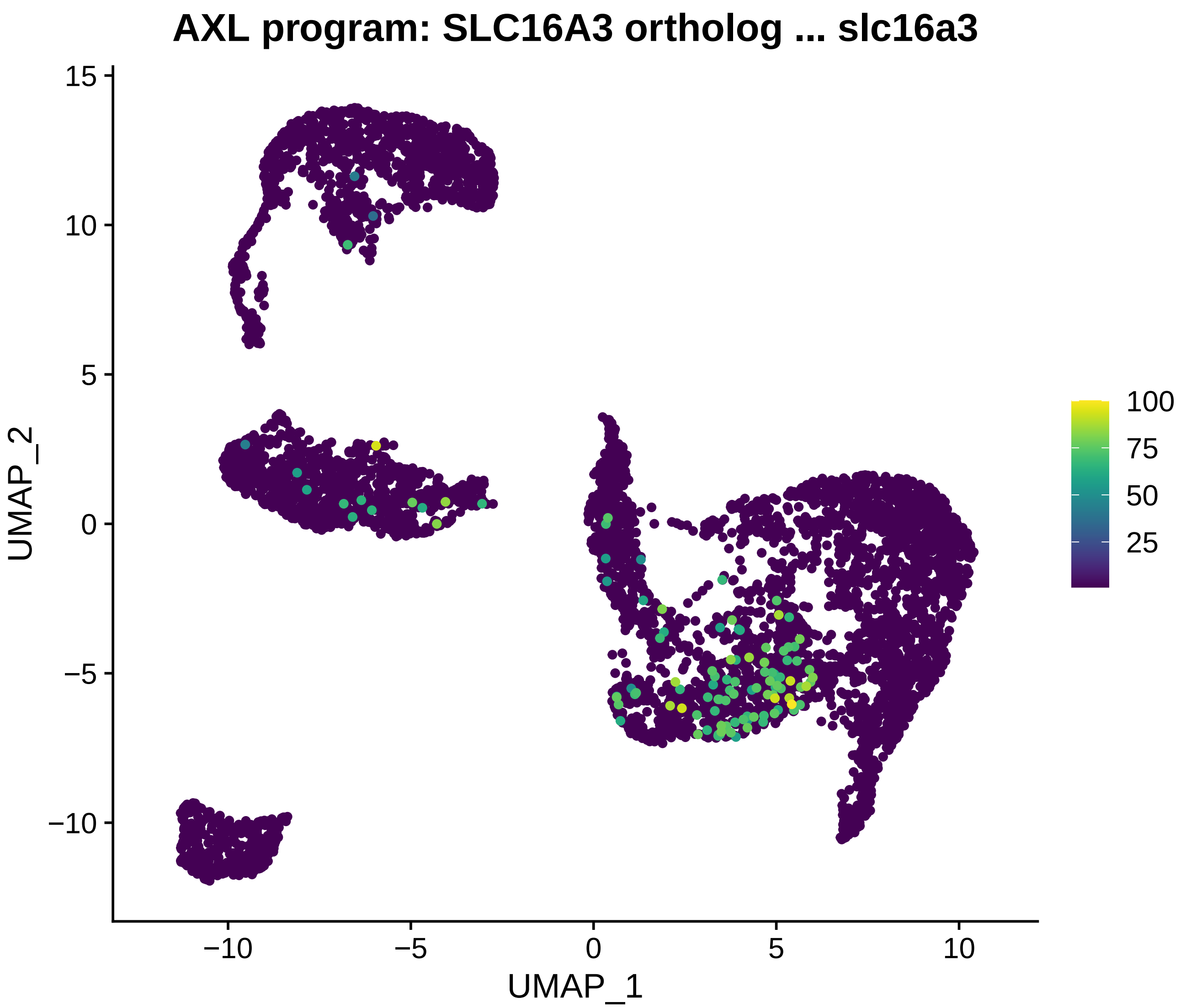
AXL program: SLC16A3 ortholog ... slc16a3
UMAP_2
UMAP_1
15
10
5
0
−5
−10
−10
−5
0
5
10
100
75
50
25

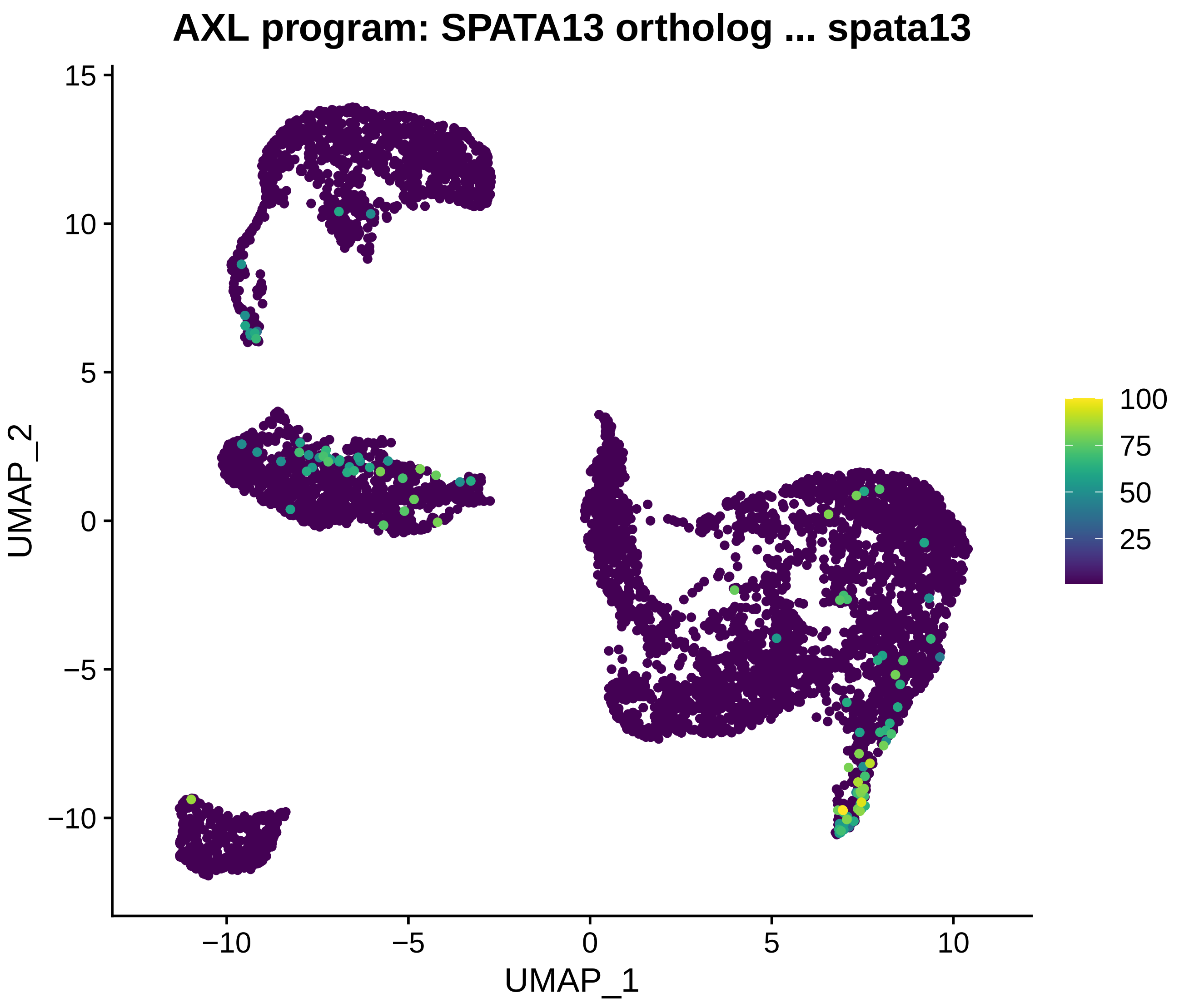
AXL program: SPATA13 ortholog ... spata13
UMAP_2
UMAP_1
15
10
5
0
−5
−10
−10
−5
0
5
10
100
75
50
25

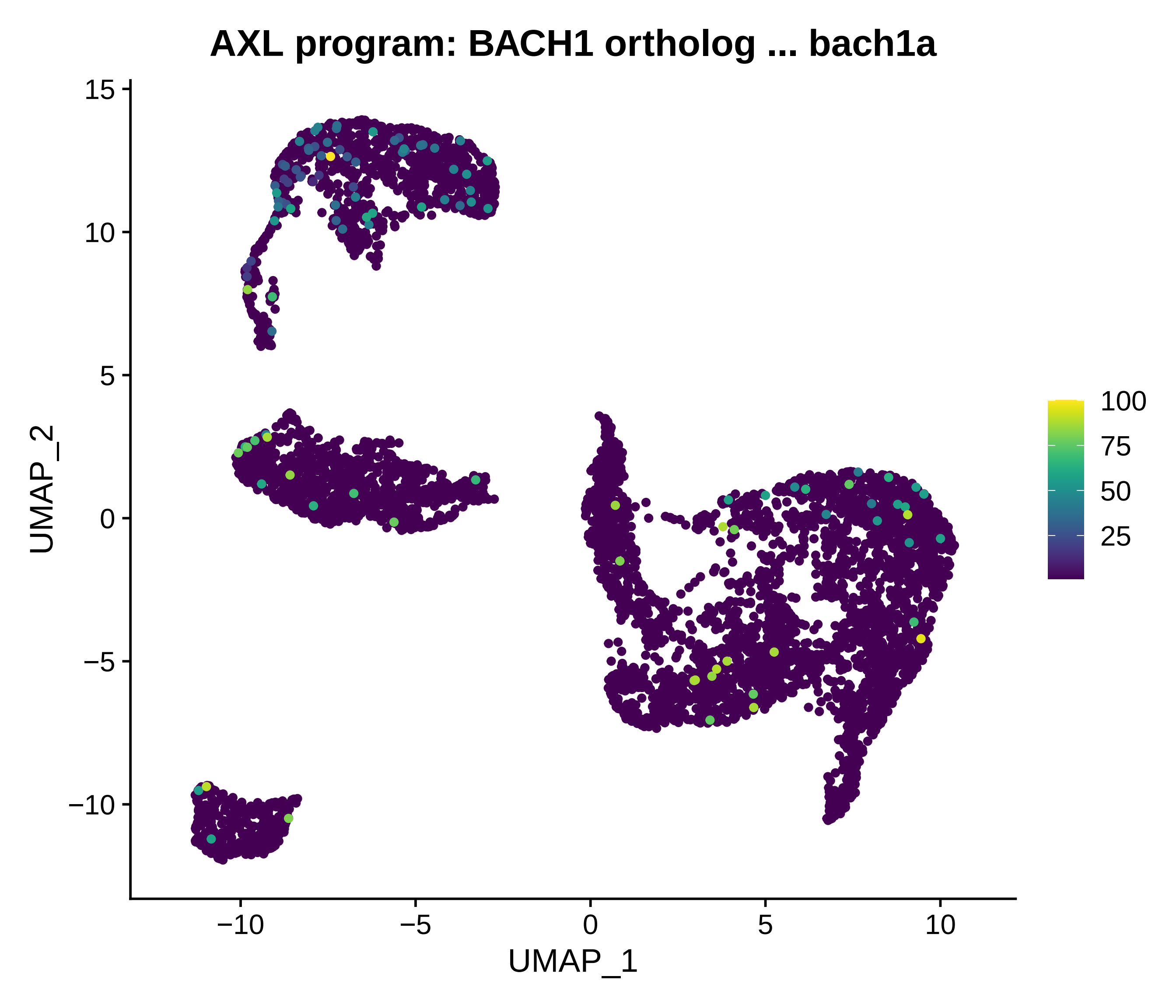
AXL program: BACH1 ortholog ... bach1a
UMAP_2
UMAP_1
15
10
5
0
−5
−10
−10
−5
0
5
10
100
75
50
25

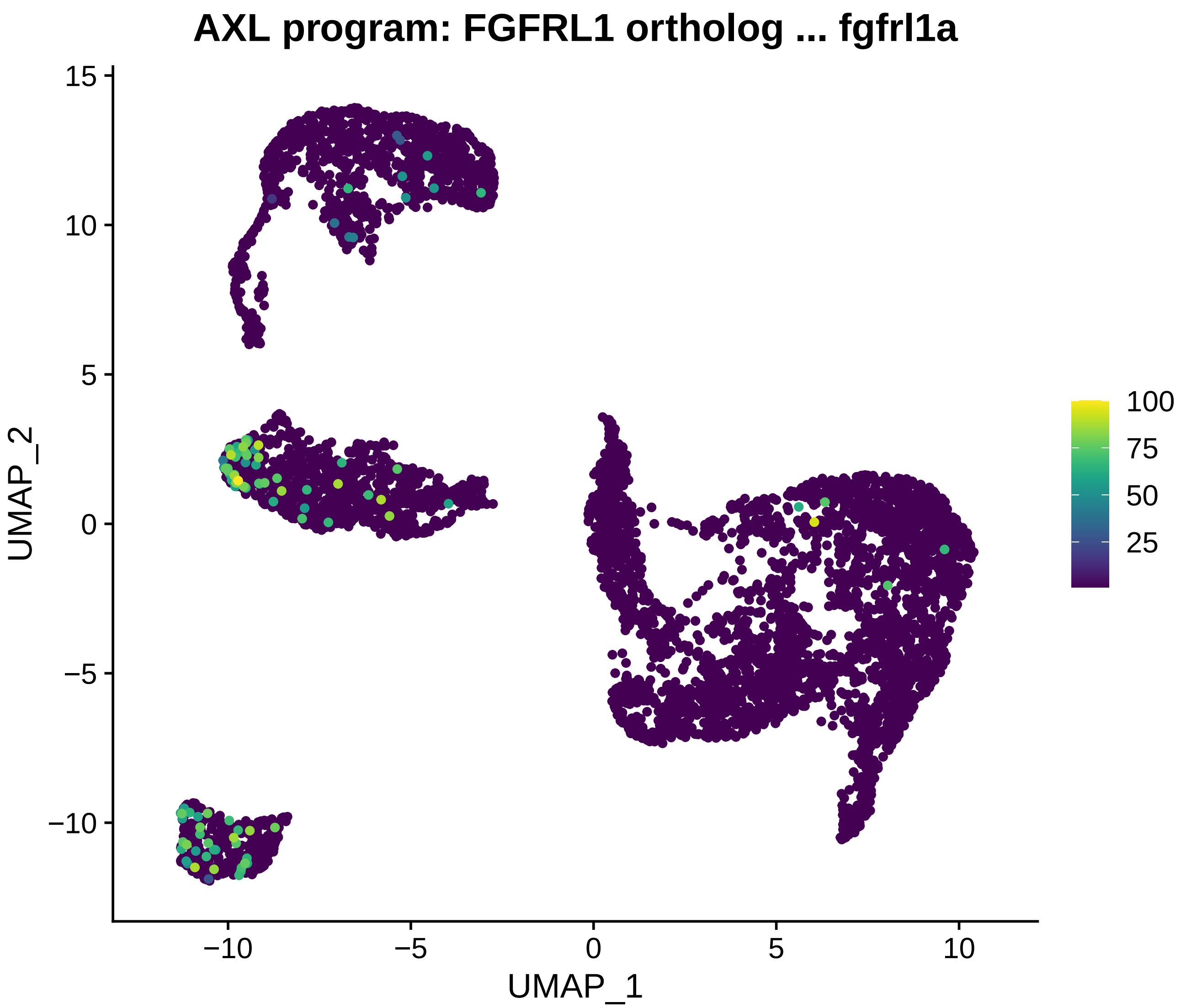
AXL program: FGFRL1 ortholog ... fgfrl1a
UMAP_1
UMAP_2
15
10
5
0
−5
−10
100
75
50
25

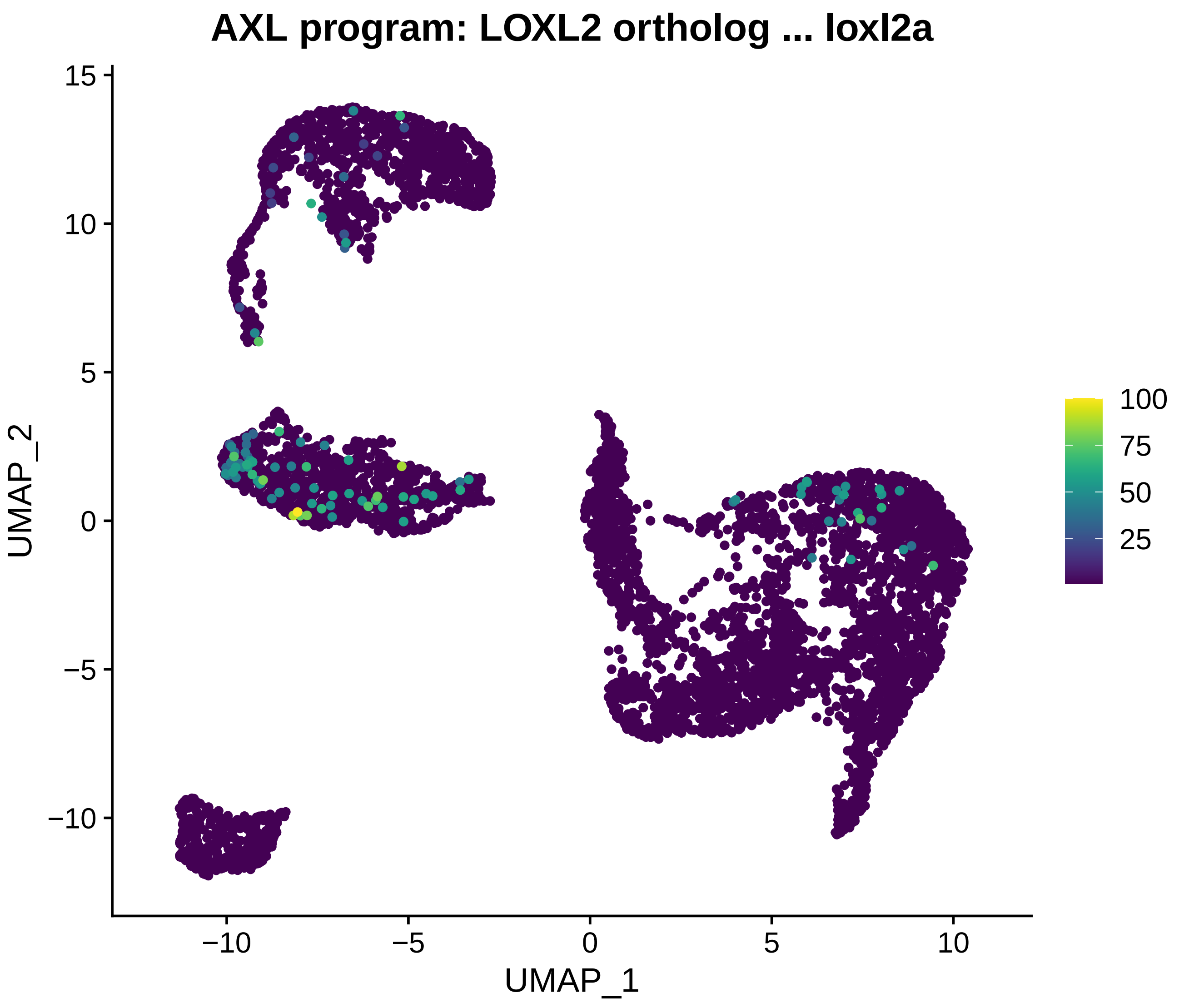
AXL program: LOXL2 ortholog ... loxl2a
UMAP_1
UMAP_2
−10
−5
0
5
10
15
100
75
50
25

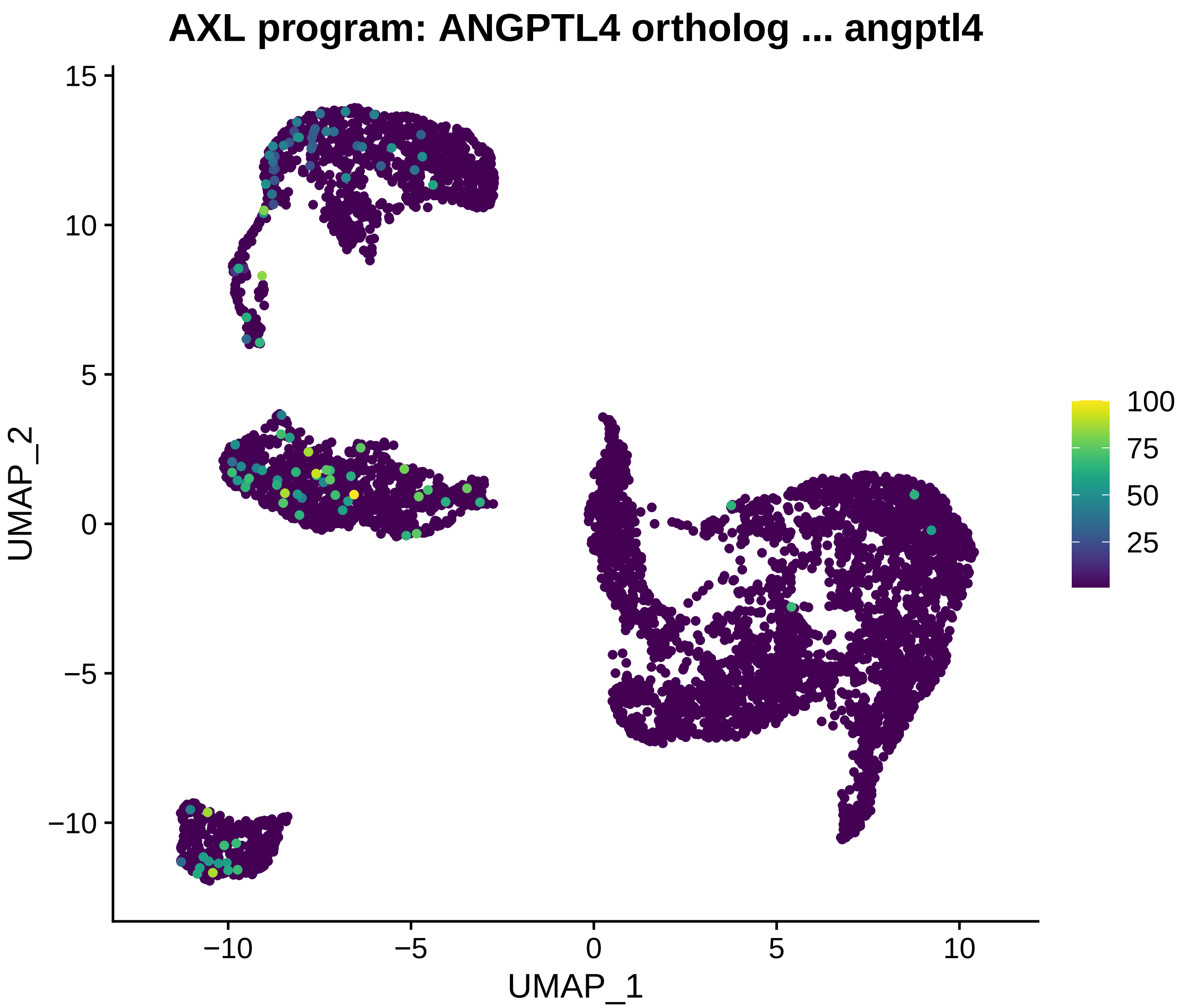
AXL program: ANGPTL4 ortholog ... angptl4
UMAP_2
UMAP_1
15
10
5
0
−5
−10
−10
−5
0
5
10
100
75
50
25

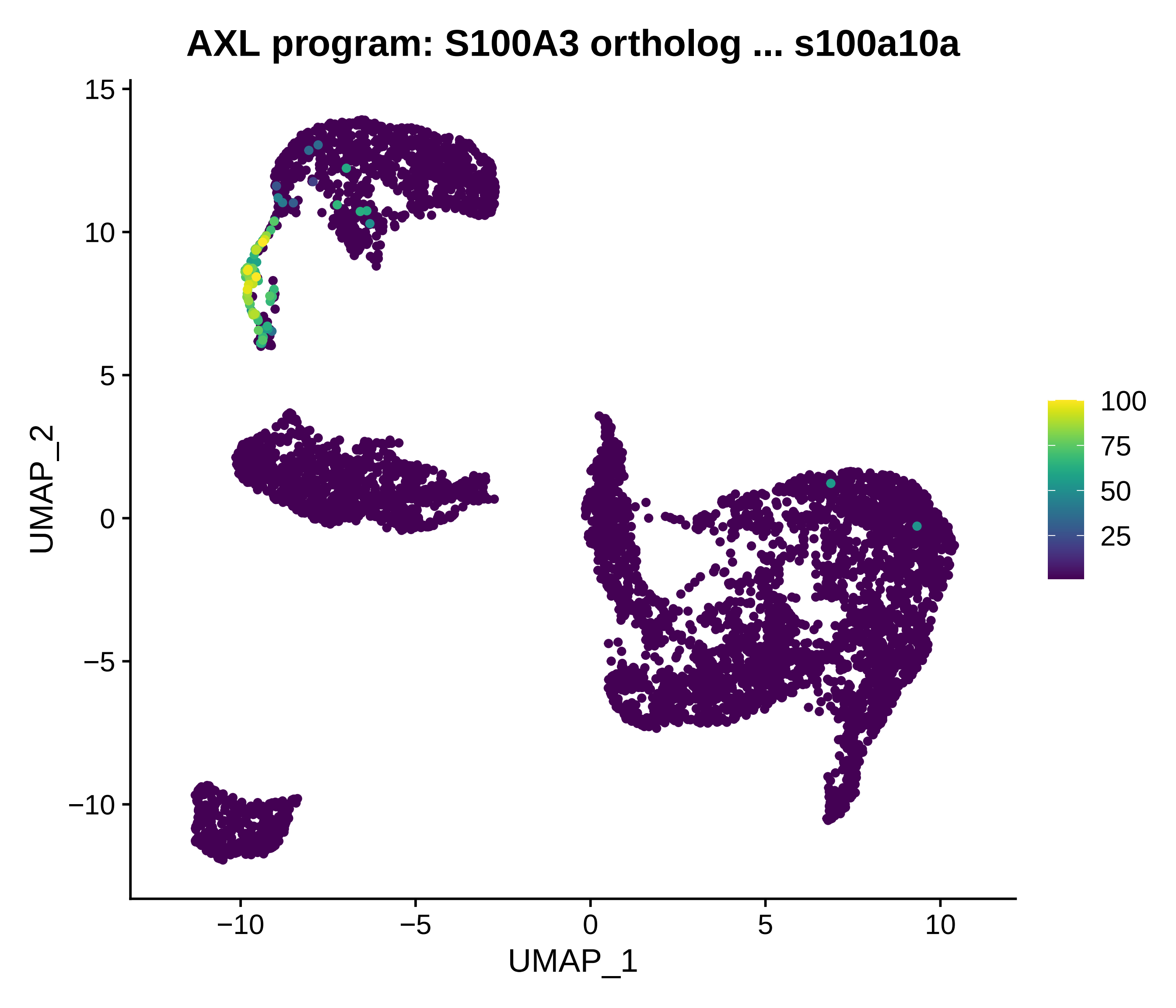
AXL program: S100A3 ortholog ... s100a10a
UMAP_1
UMAP_2
−10
−5
0
5
10
15
100
75
50
25

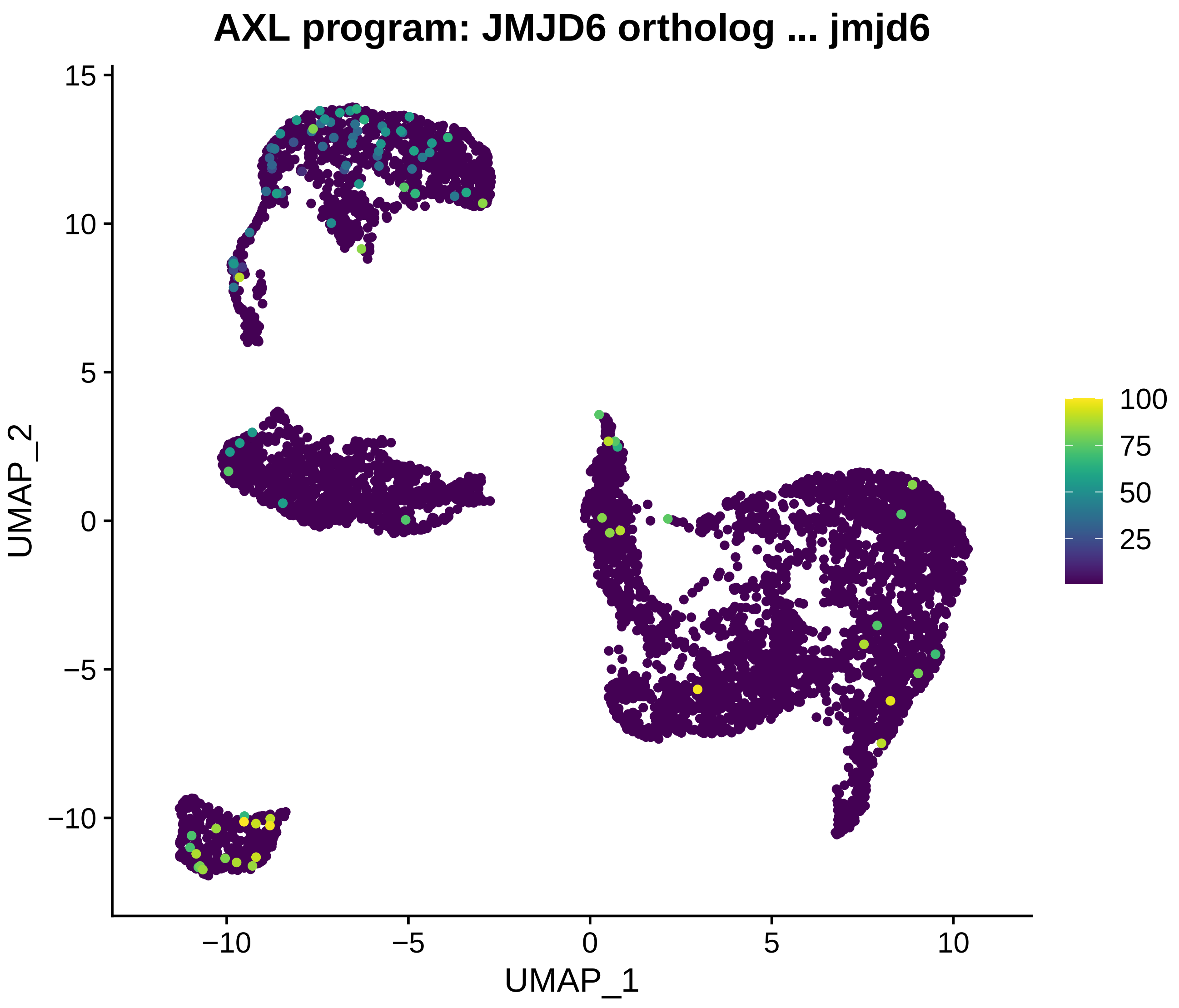
AXL program: JMJD6 ortholog ... jmjd6
UMAP_2
UMAP_1
15
10
5
0
−5
−10
−10
−5
0
5
10
100
75
50
25

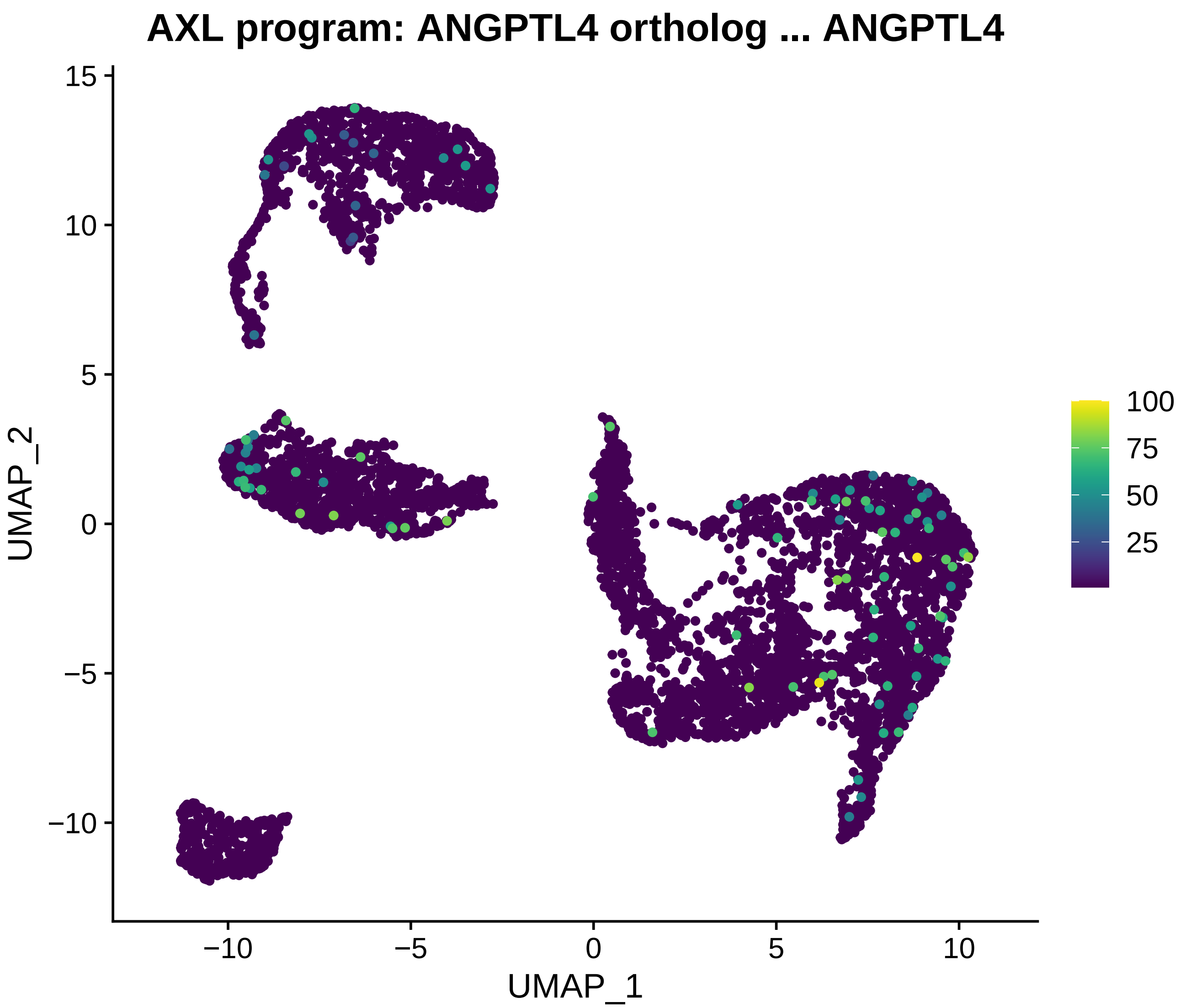

AXL program: ANGPTL4 ortholog ... ANGPTL4
15
10
5
0
−5
−10
UMAP_2
−10
−5
0
5
10
UMAP_1
100
75
50
25

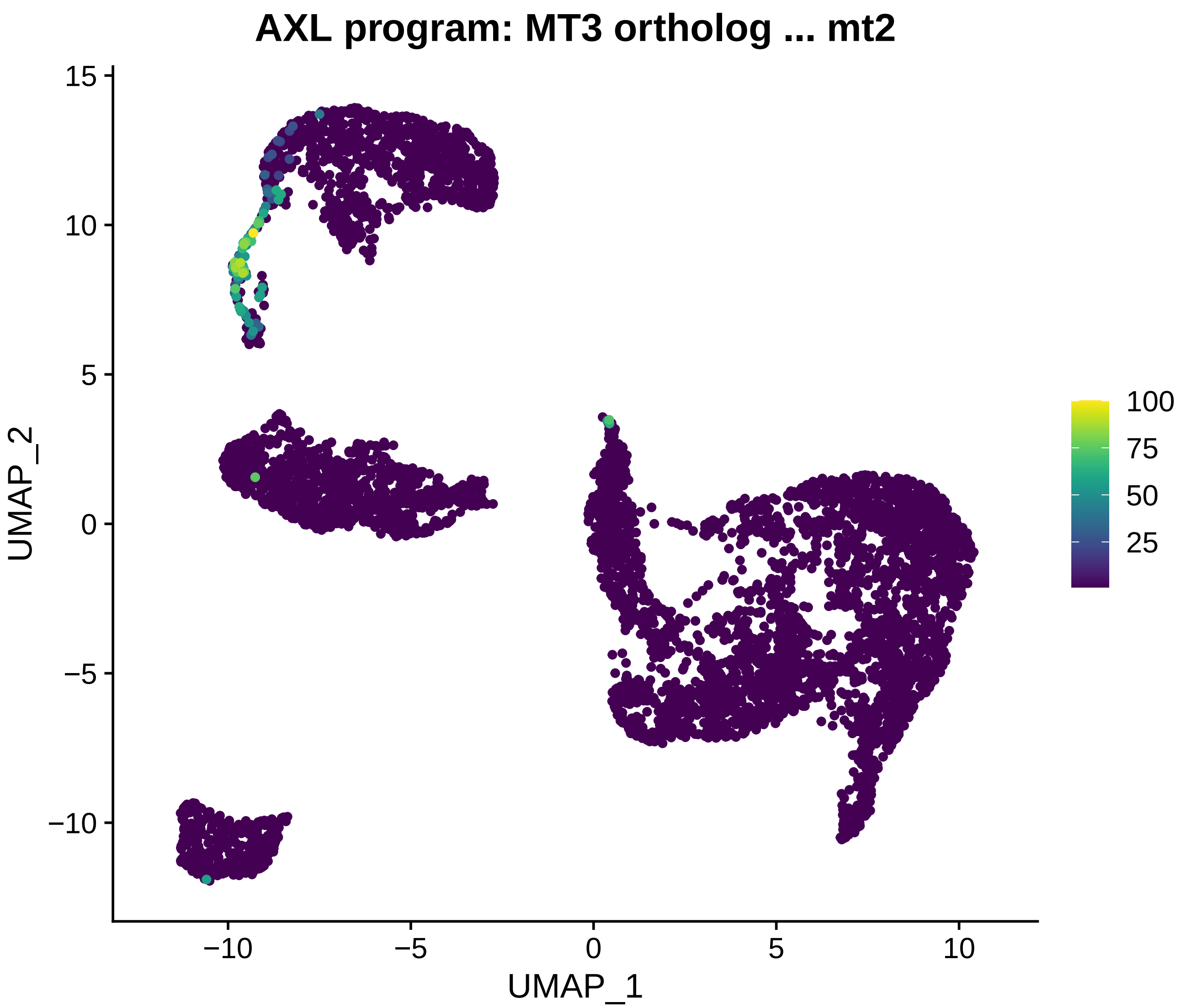
AXL program: MT3 ortholog ... mt2
UMAP_2
UMAP_1
15
10
5
0
−5
−10
−10
−5
0
5
10
100
75
50
25

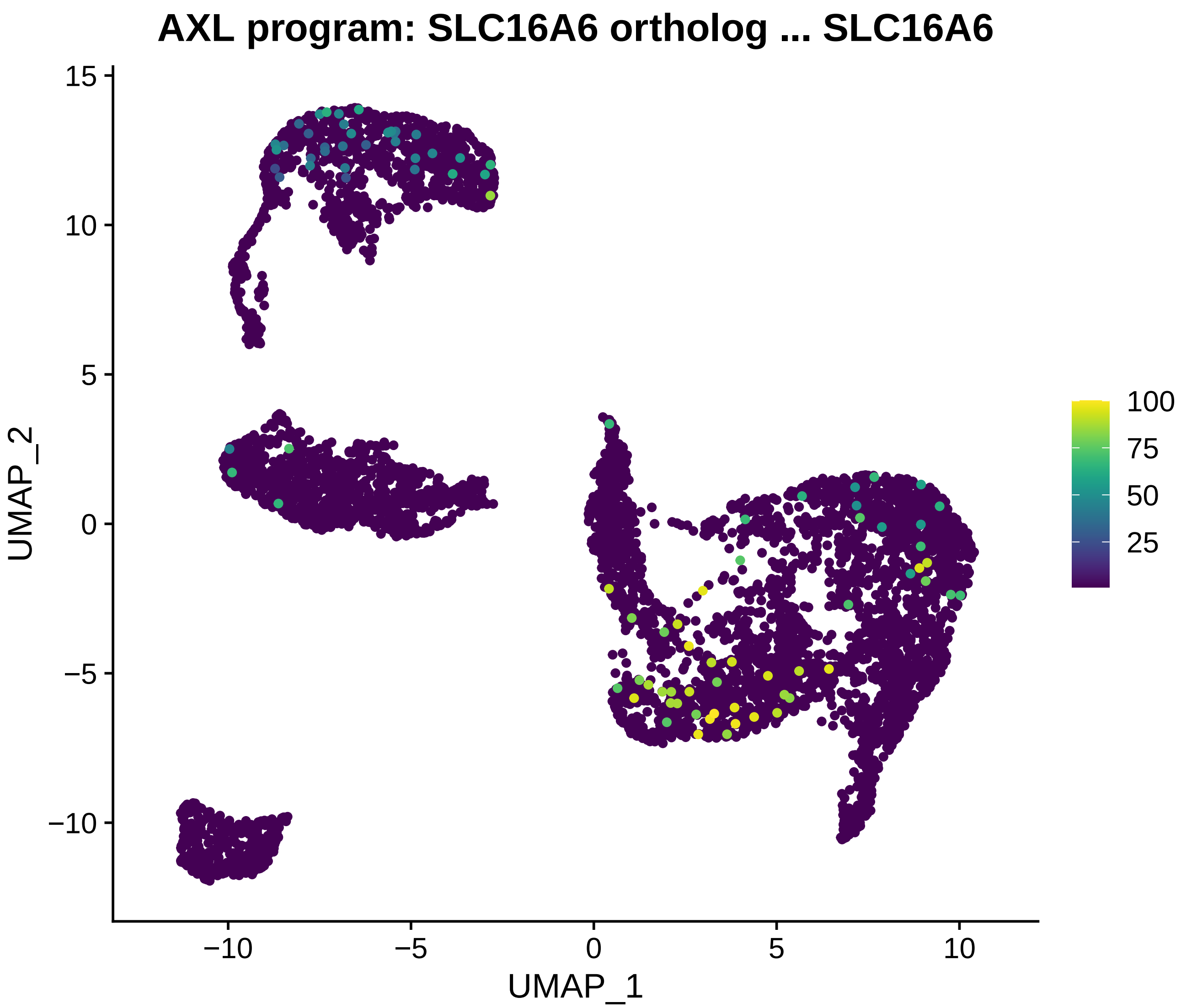

AXL program: SLC16A6 ortholog ... SLC16A6
UMAP_2
UMAP_1
15
10
5
0
−5
−10
−10
−5
0
5
10
100
75
50
25

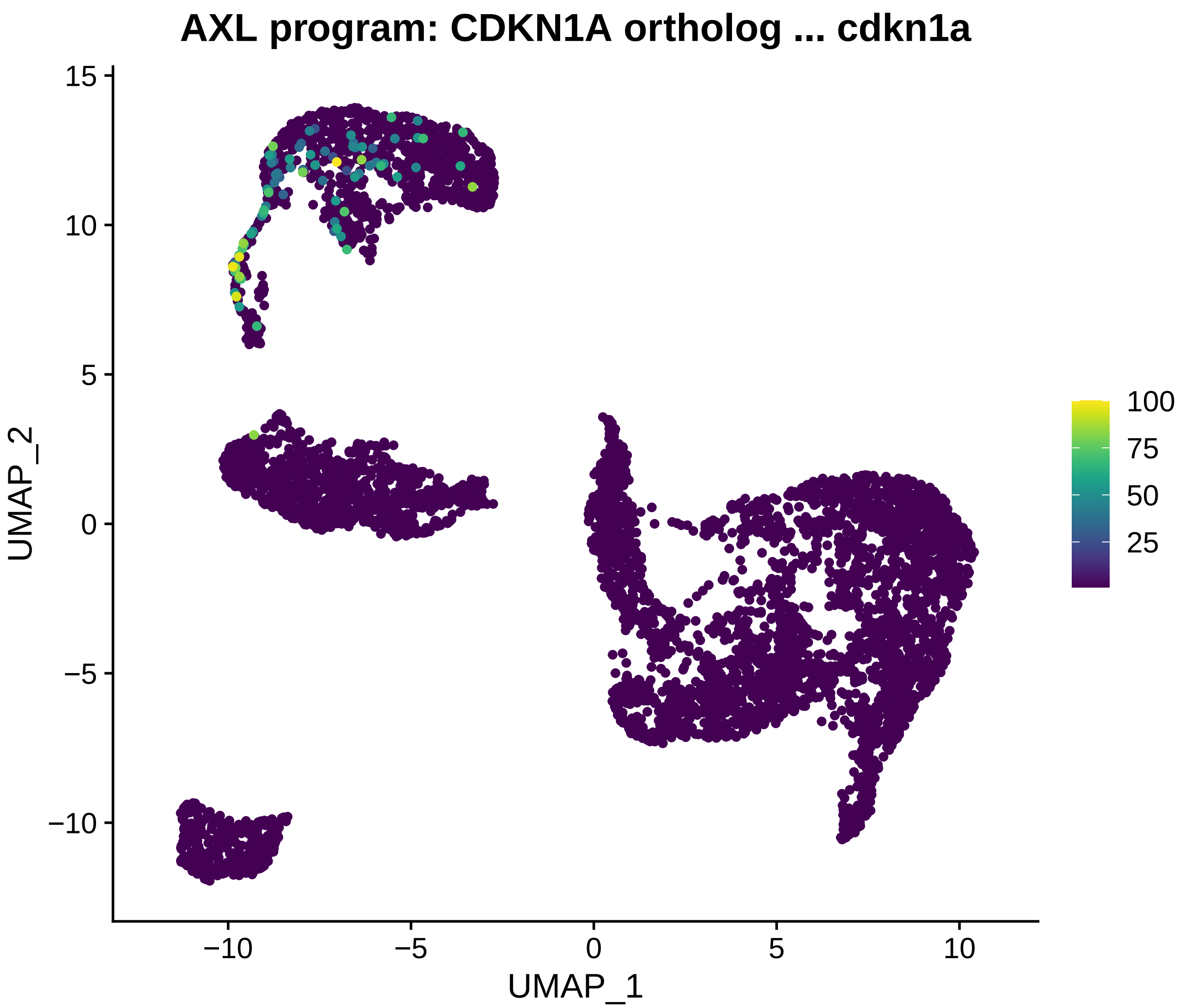
AXL program: CDKN1A ortholog ... cdkn1a
UMAP_1
UMAP_2
15
10
5
0
−5
−10
100
75
50
25

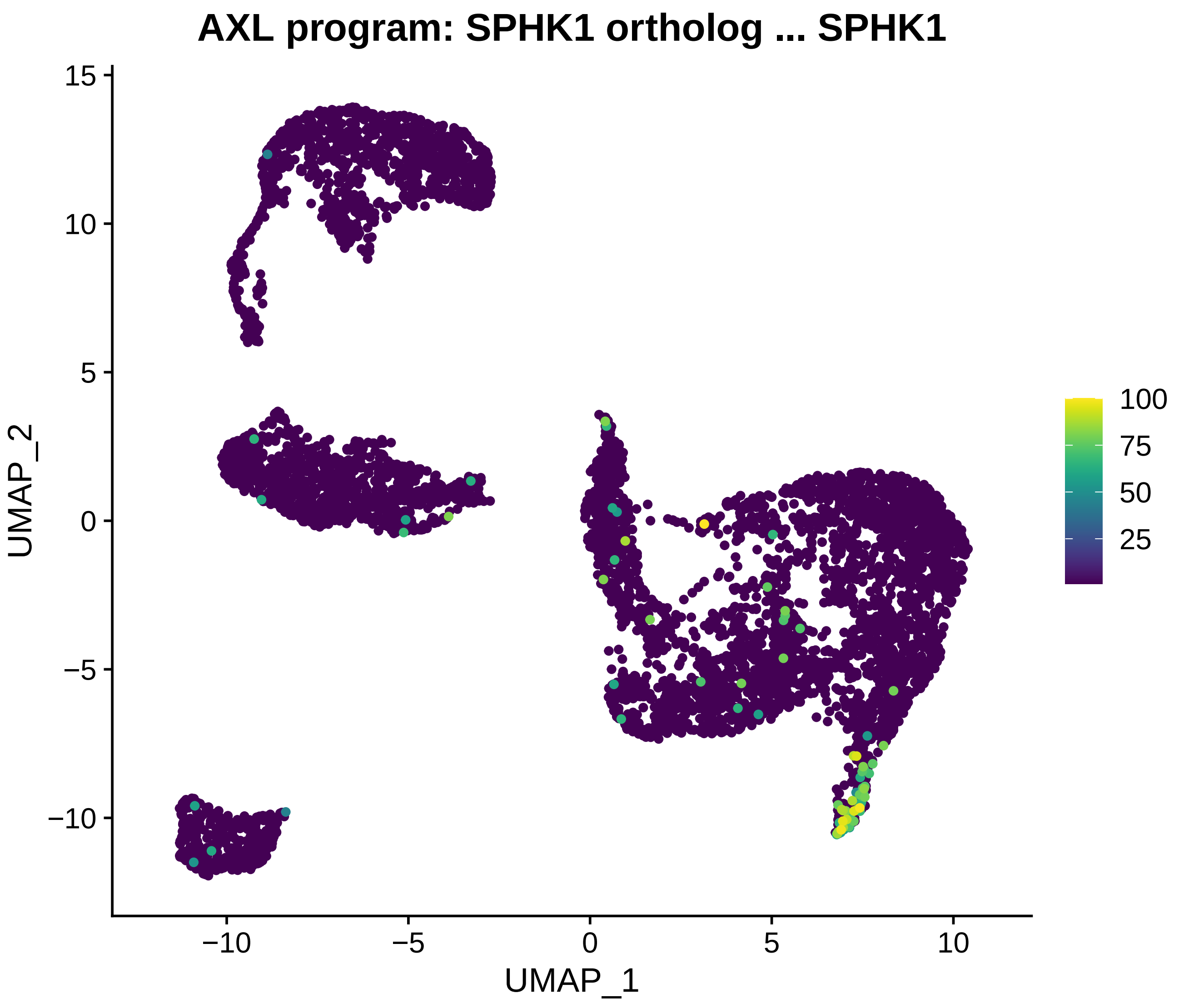
AXL program: SPHK1 ortholog ... SPHK1
UMAP_2
UMAP_1
15
10
5
0
−5
−10
−10
−5
0
5
10
100
75
50
25

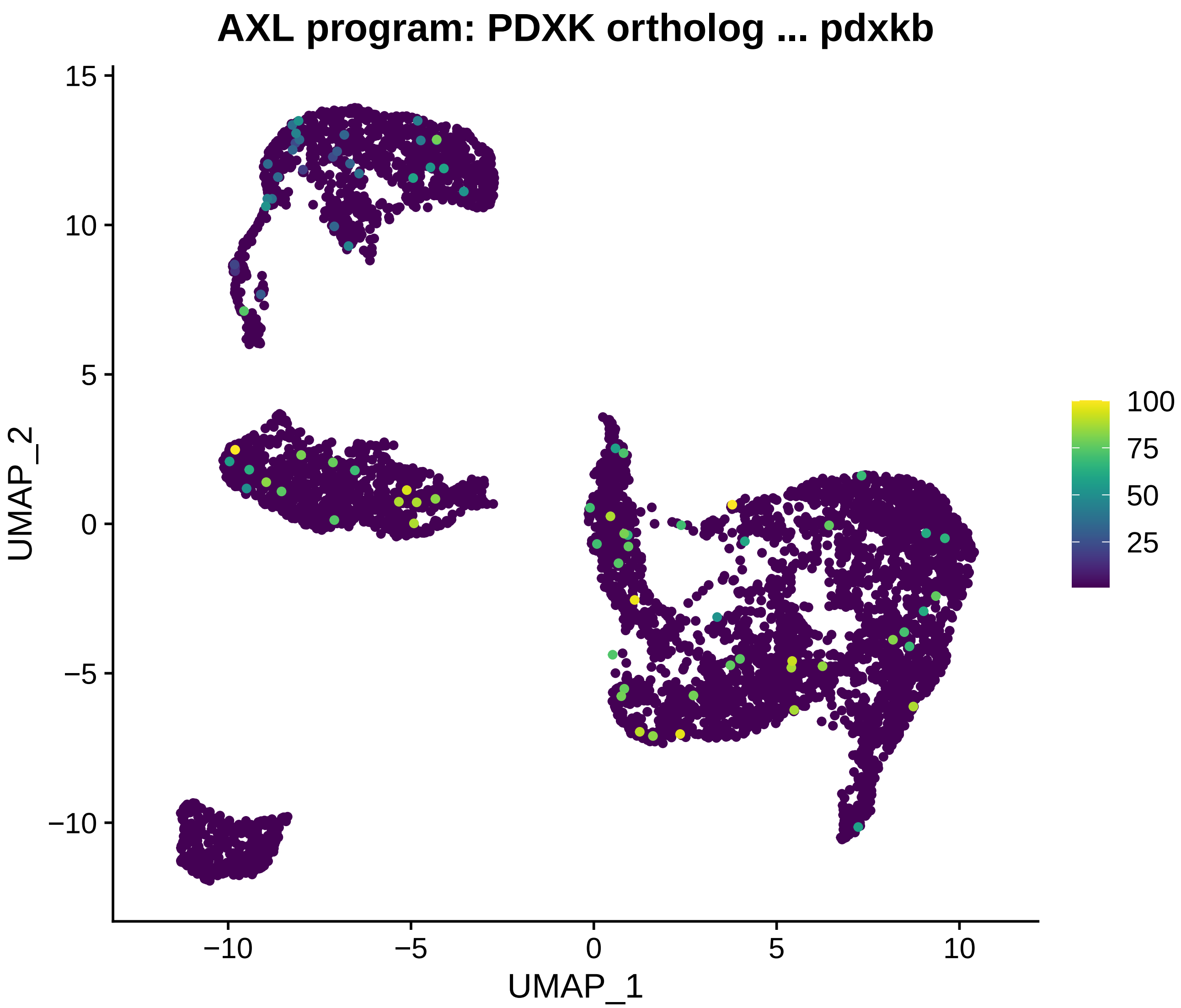
AXL program: PDXK ortholog ... pdxkb
UMAP_2
15
10
5
0
−5
−10
UMAP_1
−10
−5
0
5
10
100
75
50
25

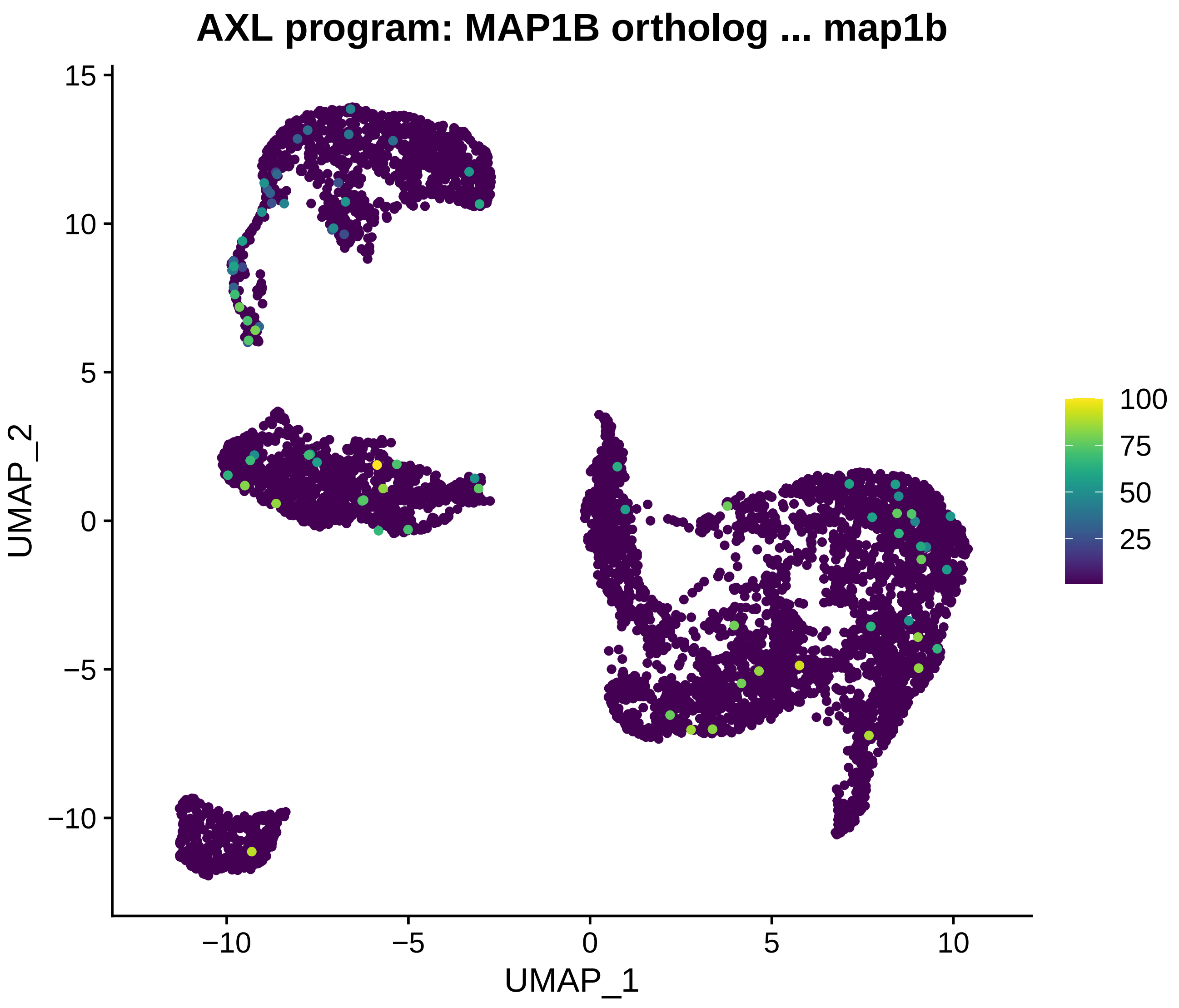
AXL program: MAP1B ortholog ... map1b
UMAP_2
UMAP_1
15
10
5
0
−5
−10
−10
−5
0
5
10
100
75
50
25

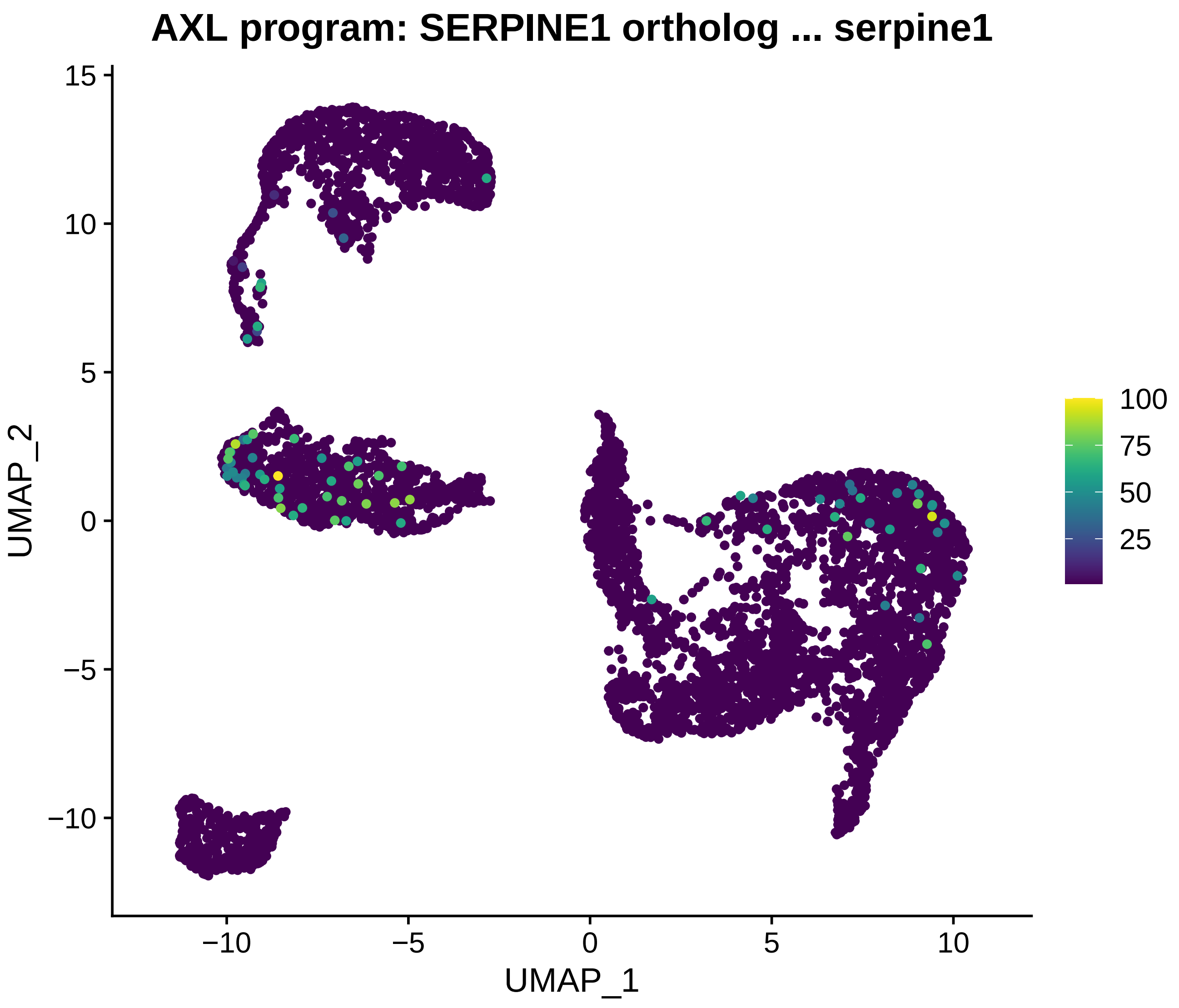

AXL program: SERPINE1 ortholog ... serpine1
UMAP_2
UMAP_1
15
10
5
0
−5
−10
−10
−5
0
5
10
100
75
50
25

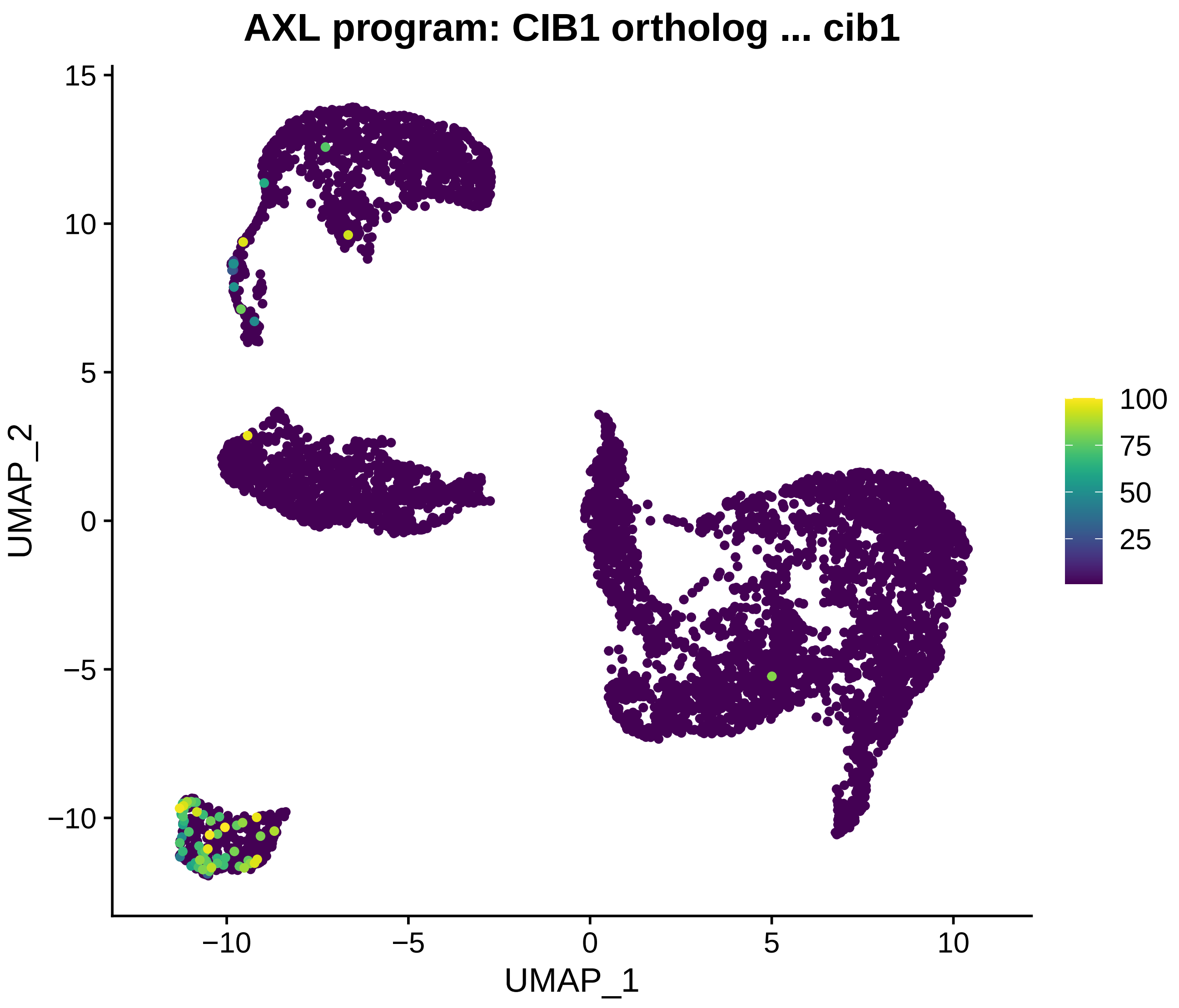

AXL program: CIB1 ortholog ... cib1
UMAP_1
UMAP_2
15
10
5
0
−5
−10
10
100
75
50
25

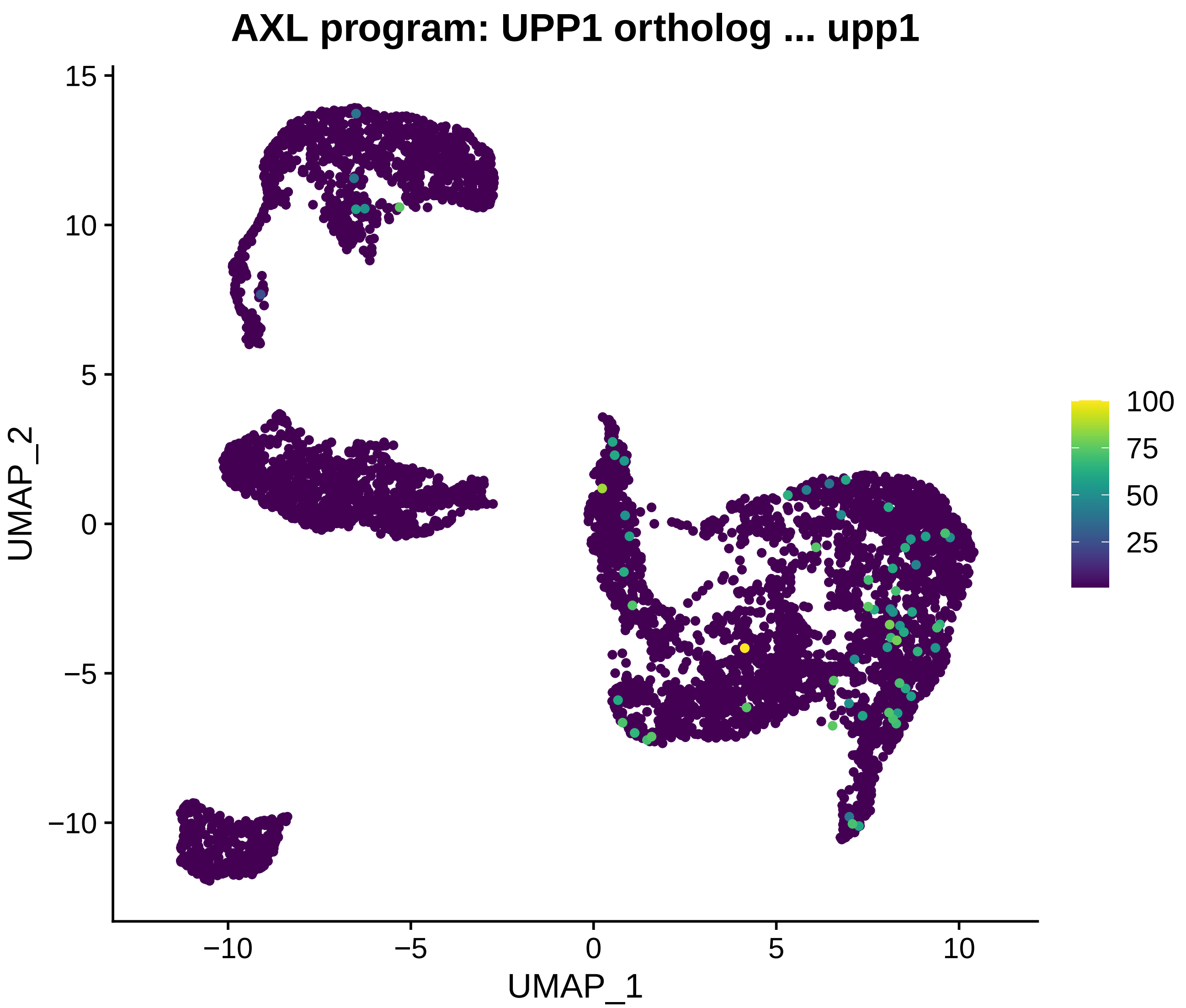
AXL program: UPP1 ortholog ... upp1
UMAP_1
UMAP_2
−10
−5
0
5
10
15
100
75
50
25

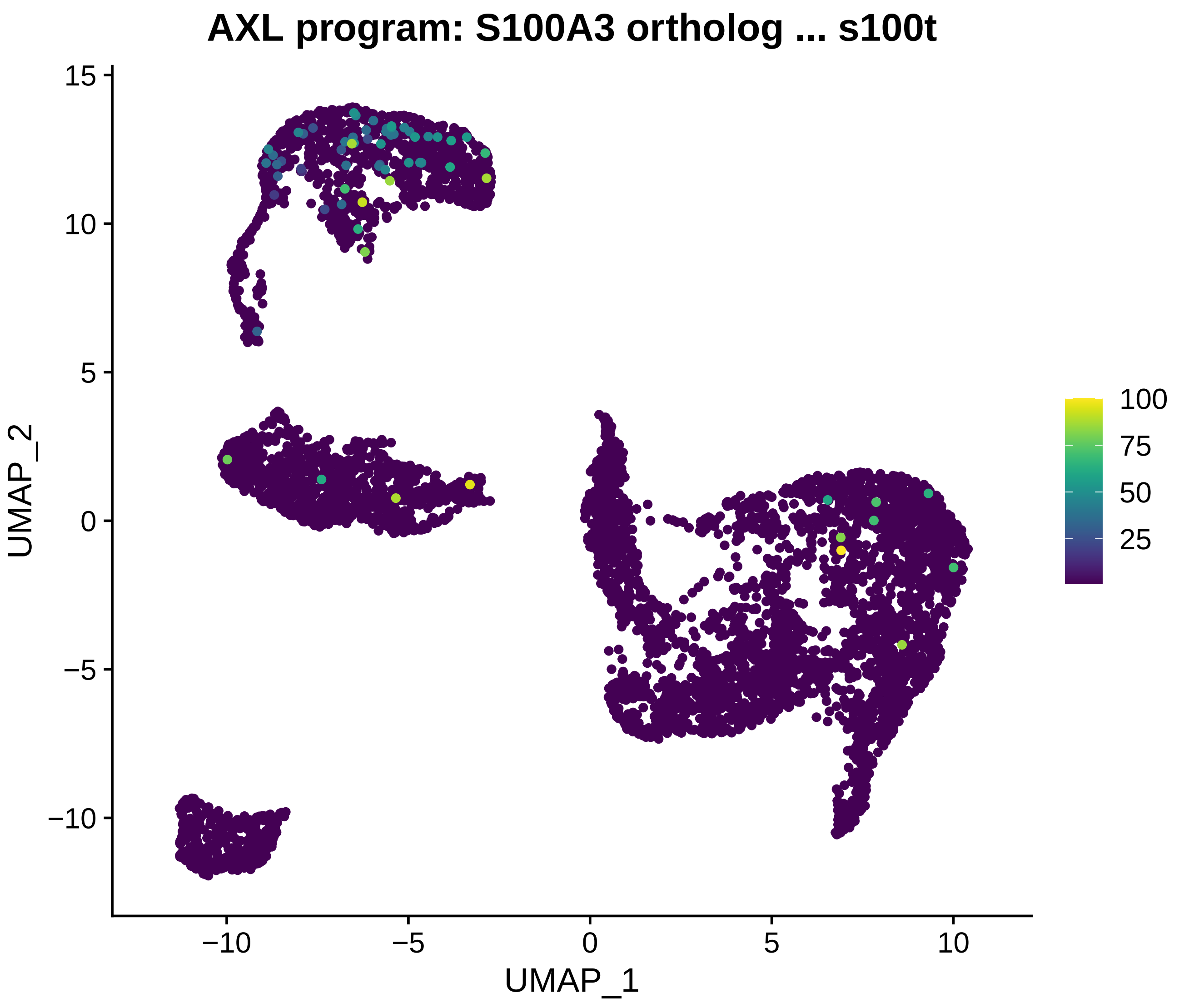
AXL program: S100A3 ortholog ... s100t
UMAP_2
UMAP_1
15
10
5
0
−5
−10
−10
−5
0
5
10
100
75
50
25

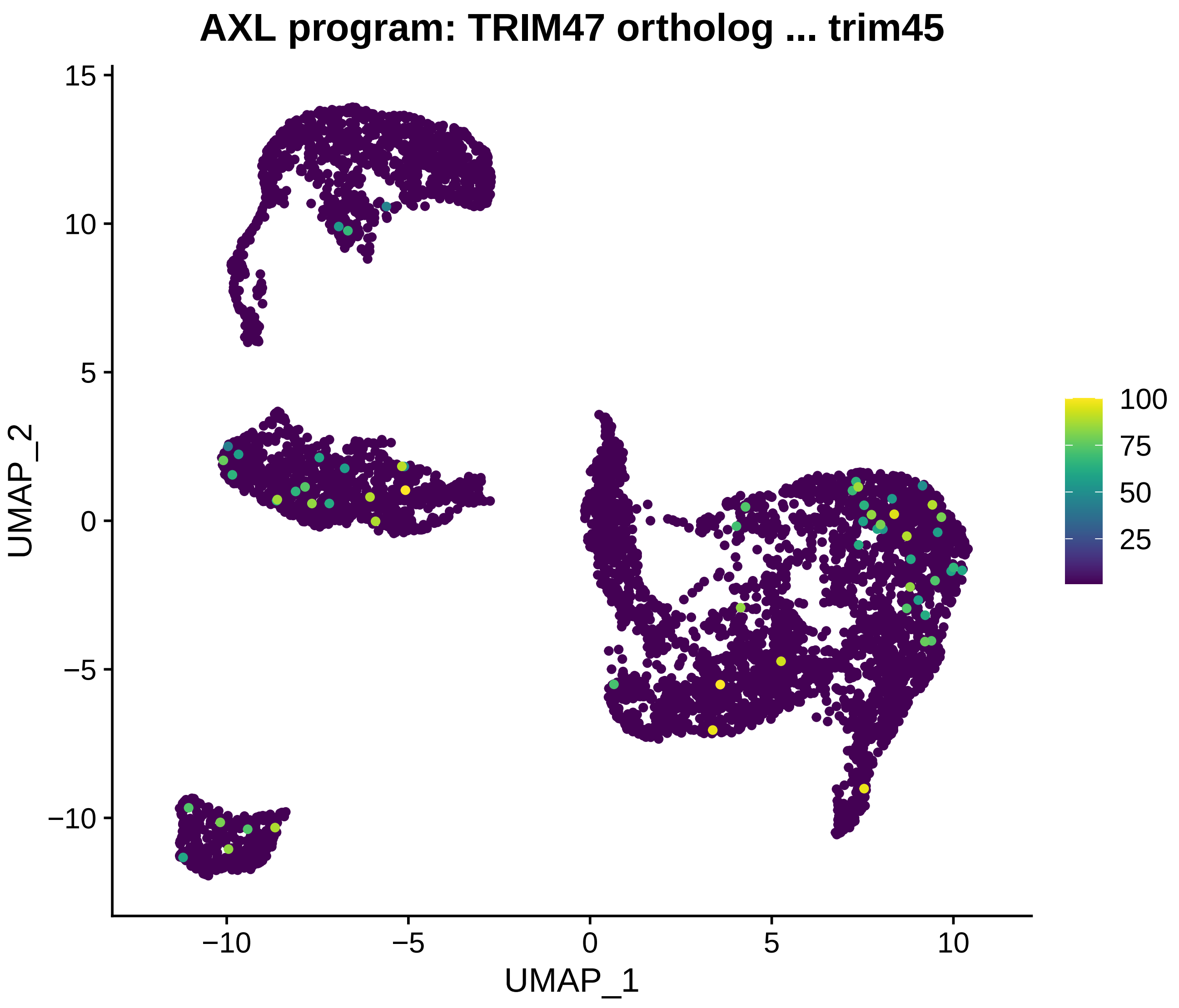

AXL program: TRIM47 ortholog ... trim45
UMAP_2
UMAP_1
15
10
5
0
−5
−10
−10
−5
0
5
10
100
75
50
25

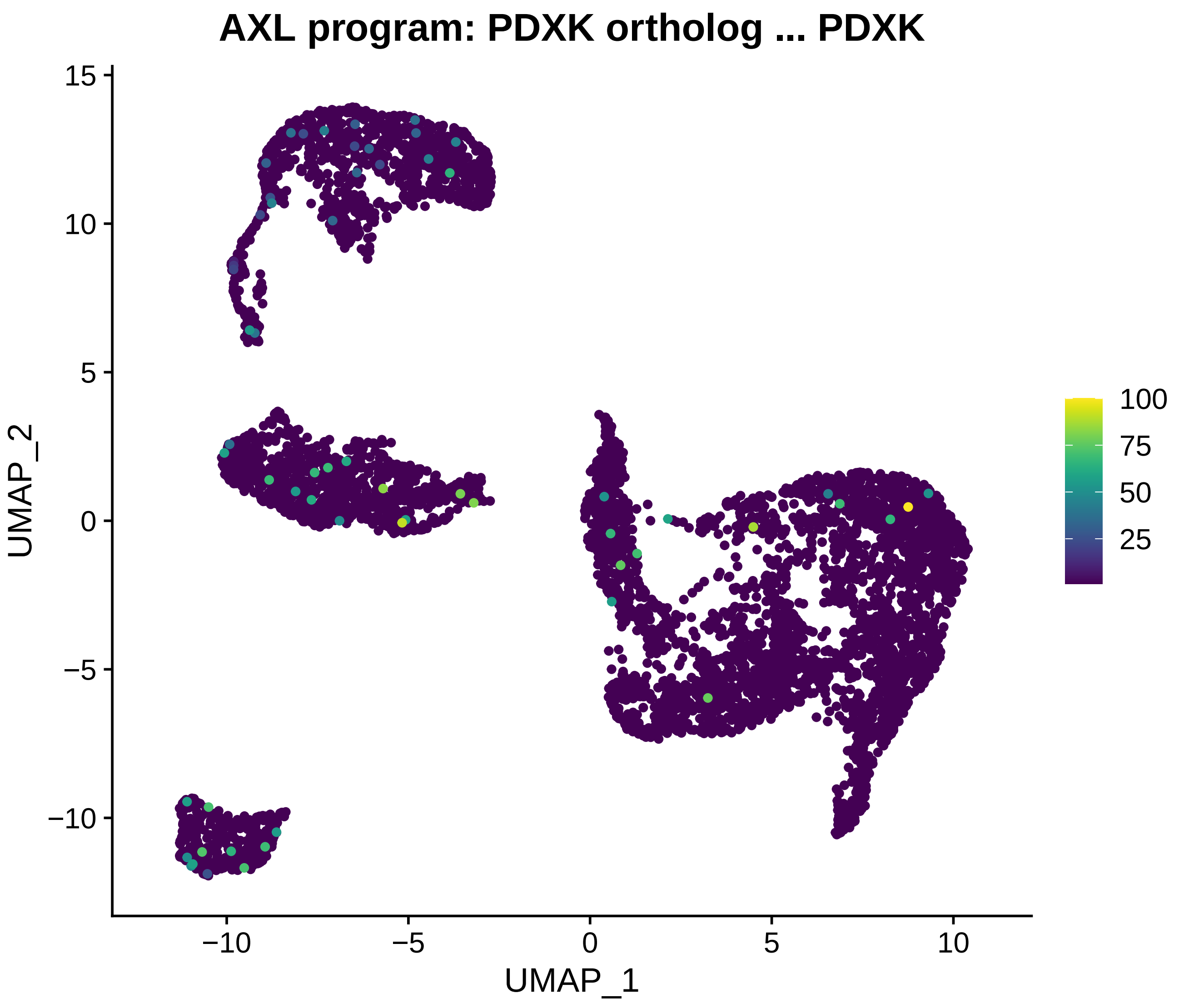
AXL program: PDXK ortholog ... PDXK
UMAP_2
UMAP_1
15
10
5
0
−5
−10
−10
−5
0
5
10
100
75
50
25

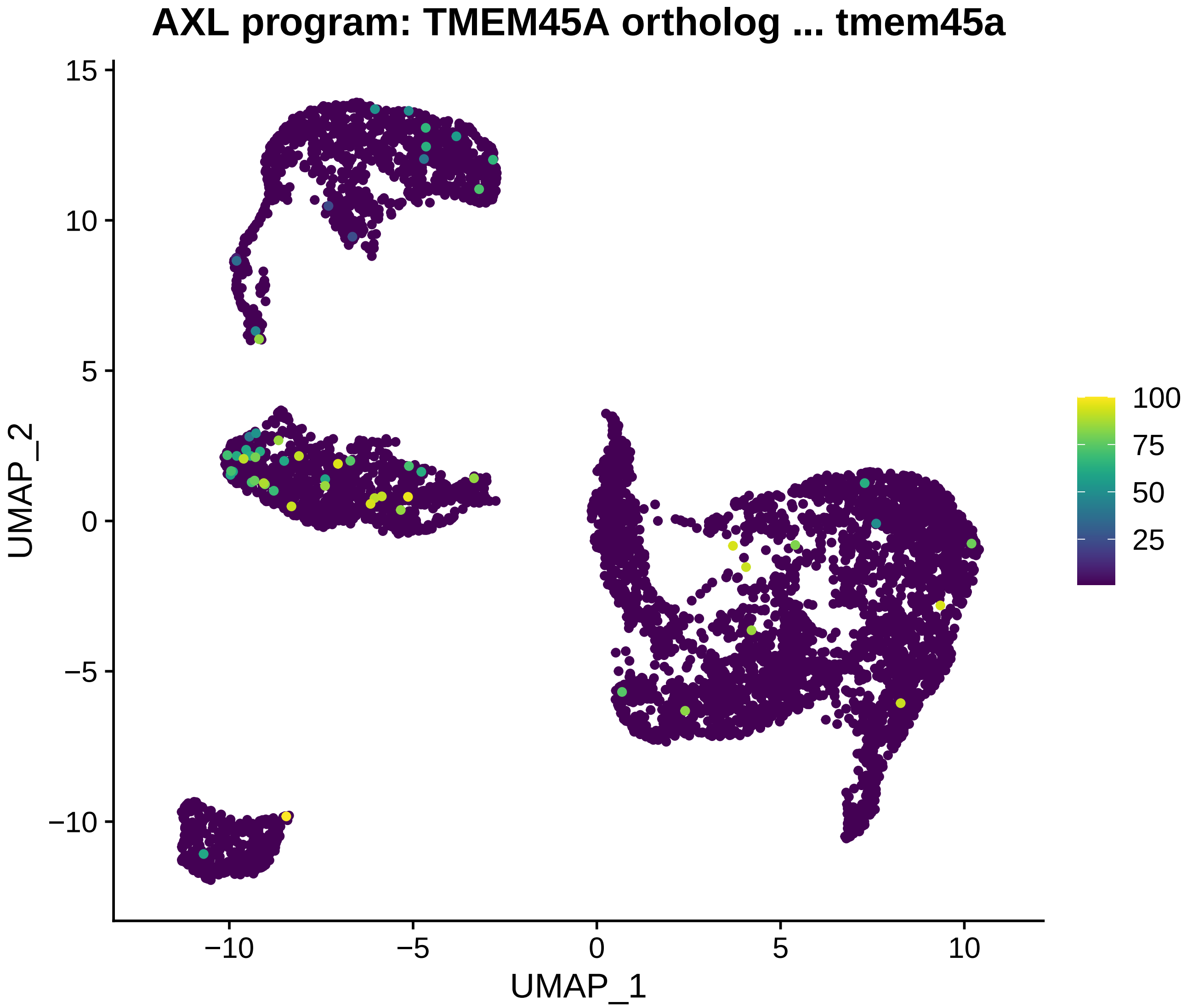
AXL program: TMEM45A ortholog ... tmem45a
UMAP_2
UMAP_1
15
10
5
0
−5
−10
−10
−5
0
5
10
100
75
50
25

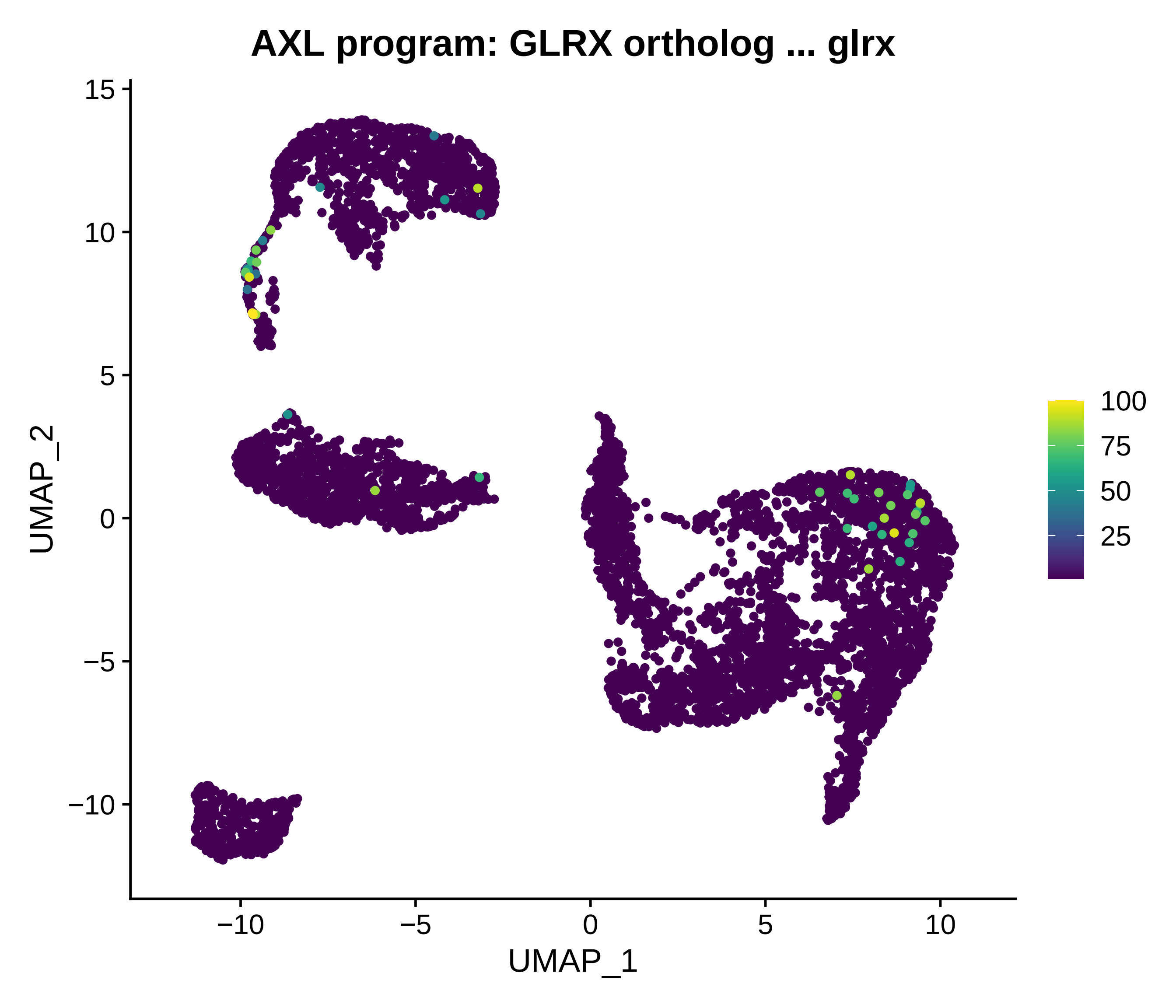
AXL program: GLRX ortholog ... glrx
UMAP_2
UMAP_1
15
10
5
0
−5
−10
−10
−5
0
5
10
100
75
50
25

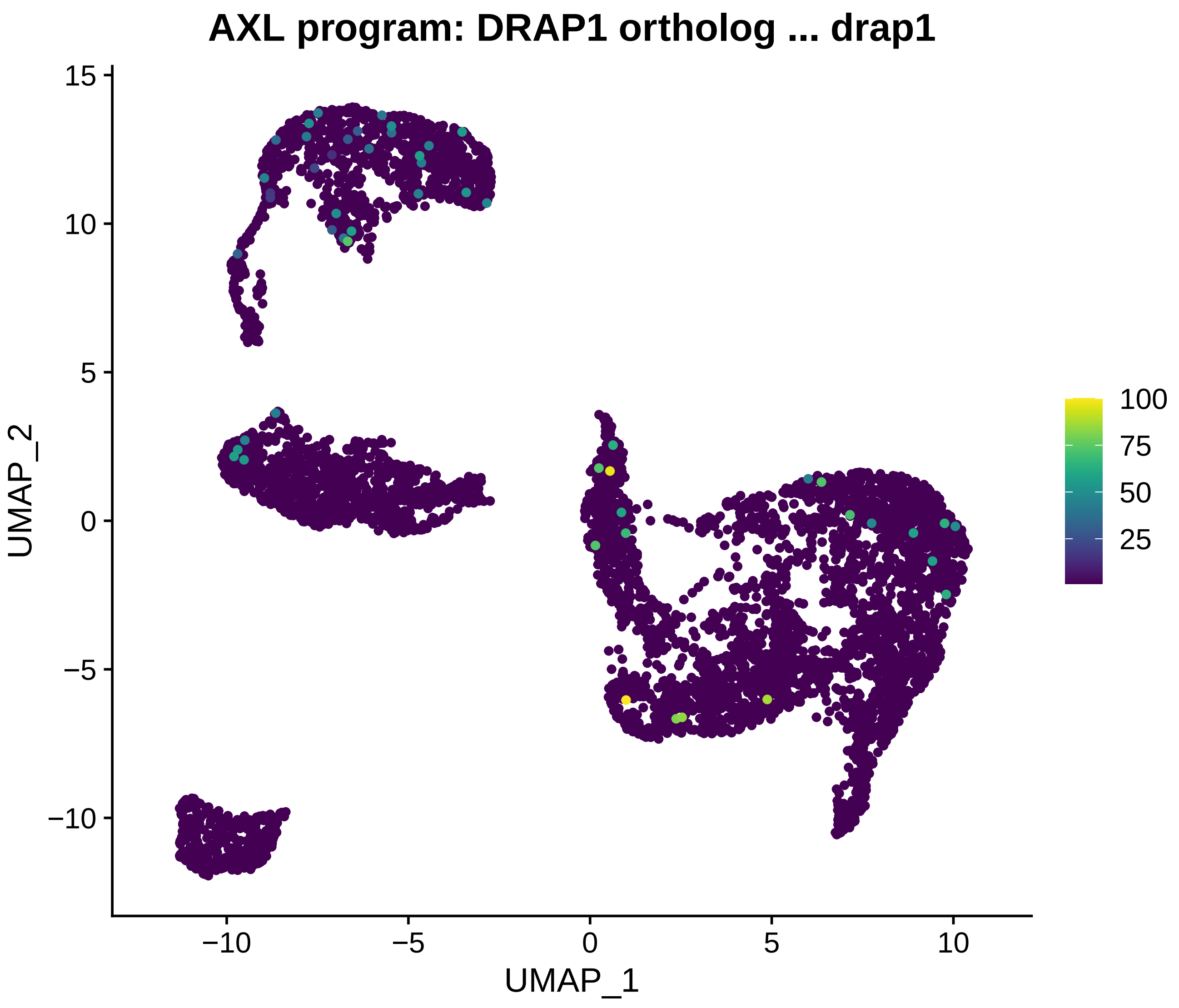
AXL program: DRAP1 ortholog ... drap1
UMAP_2
UMAP_1
15
10
5
0
−5
−10
−10
−5
0
5
10
100
75
50
25

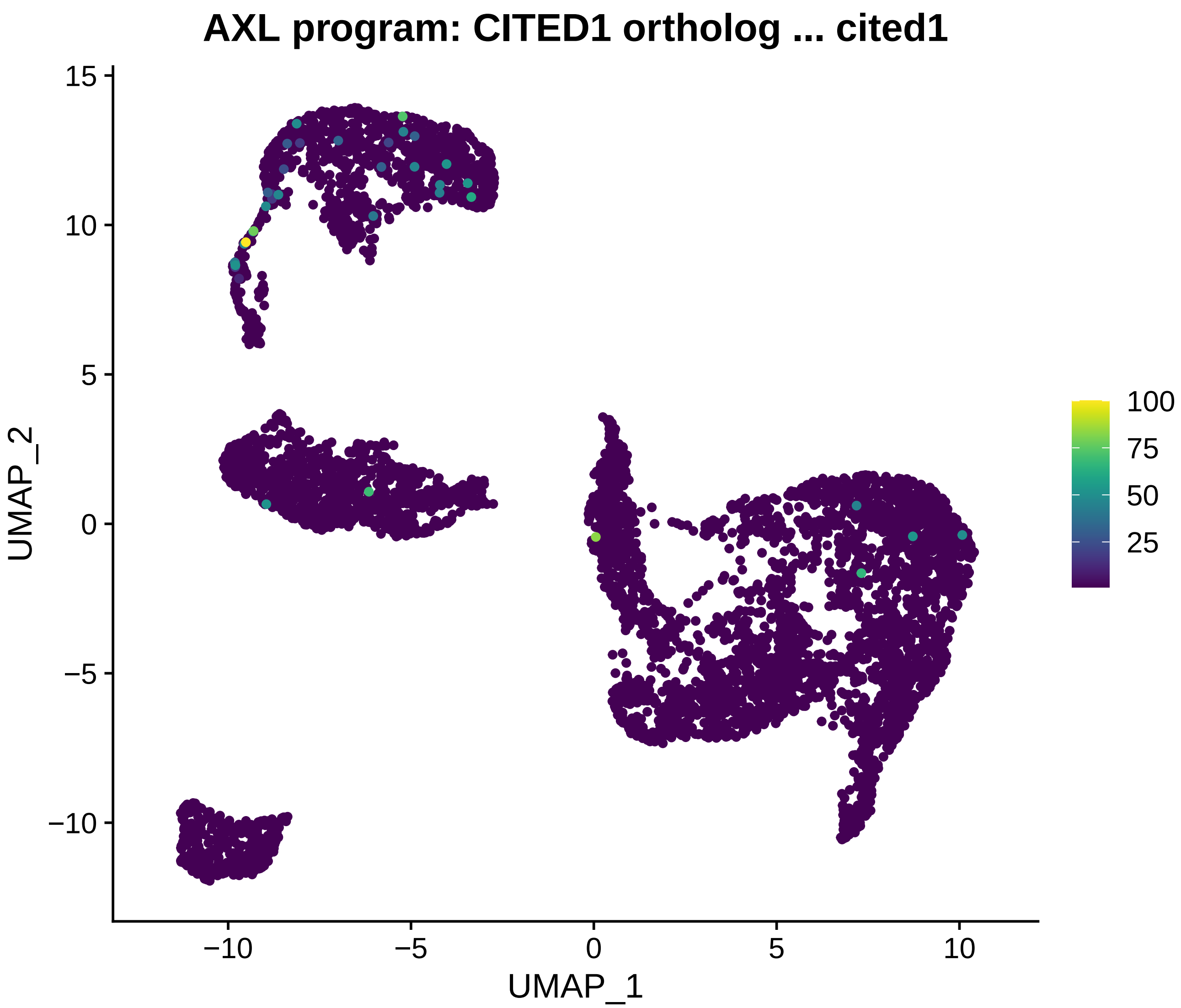
AXL program: CITED1 ortholog ... cited1
UMAP_1
UMAP_2
15
10
5
0
−5
−10
0
5
10
100
75
50
25

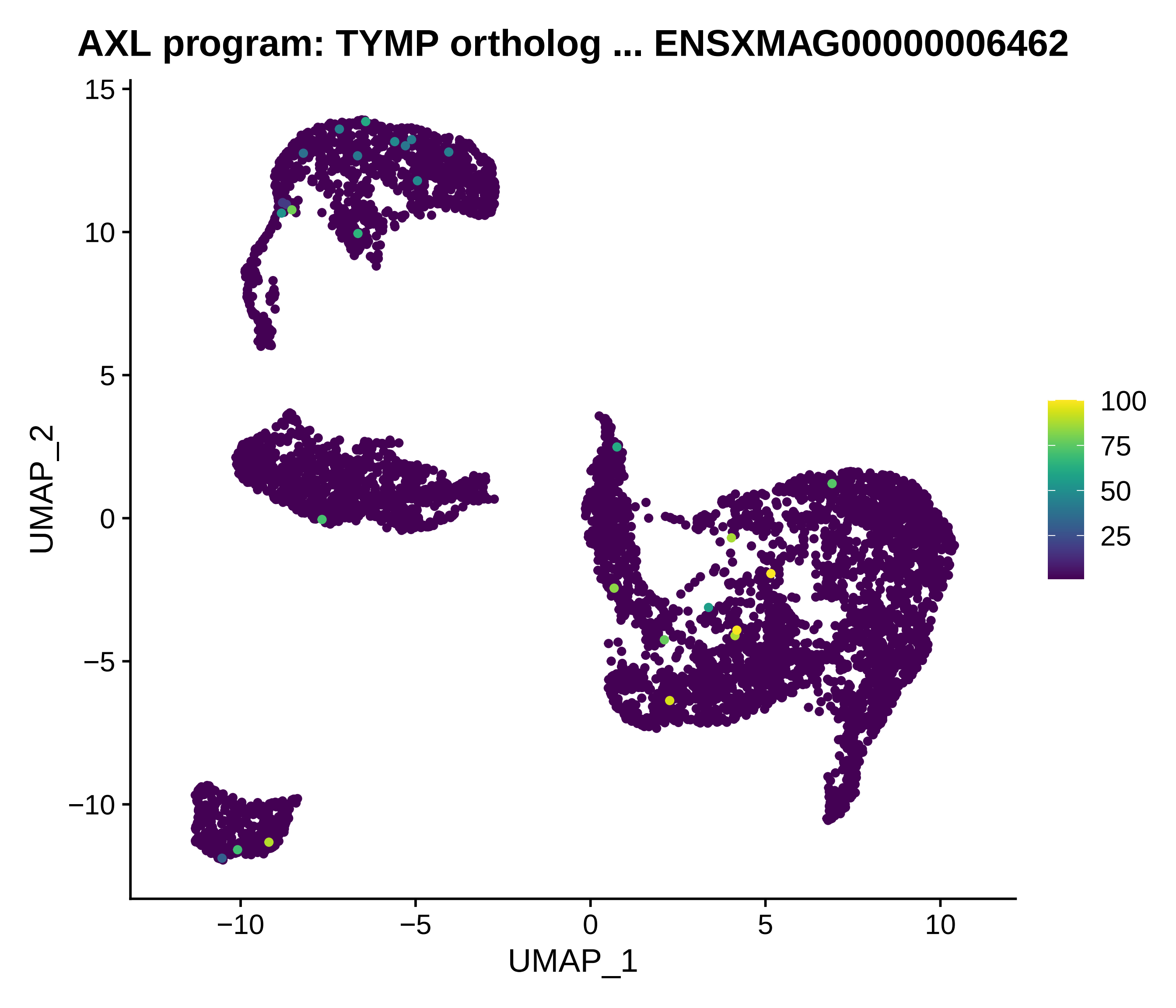
AXL program: TYMP ortholog ... ENSXMAG00000006462
UMAP_2
UMAP_1
15
10
5
0
−5
−10
−10
−5
0
5
10
100
75
50
25

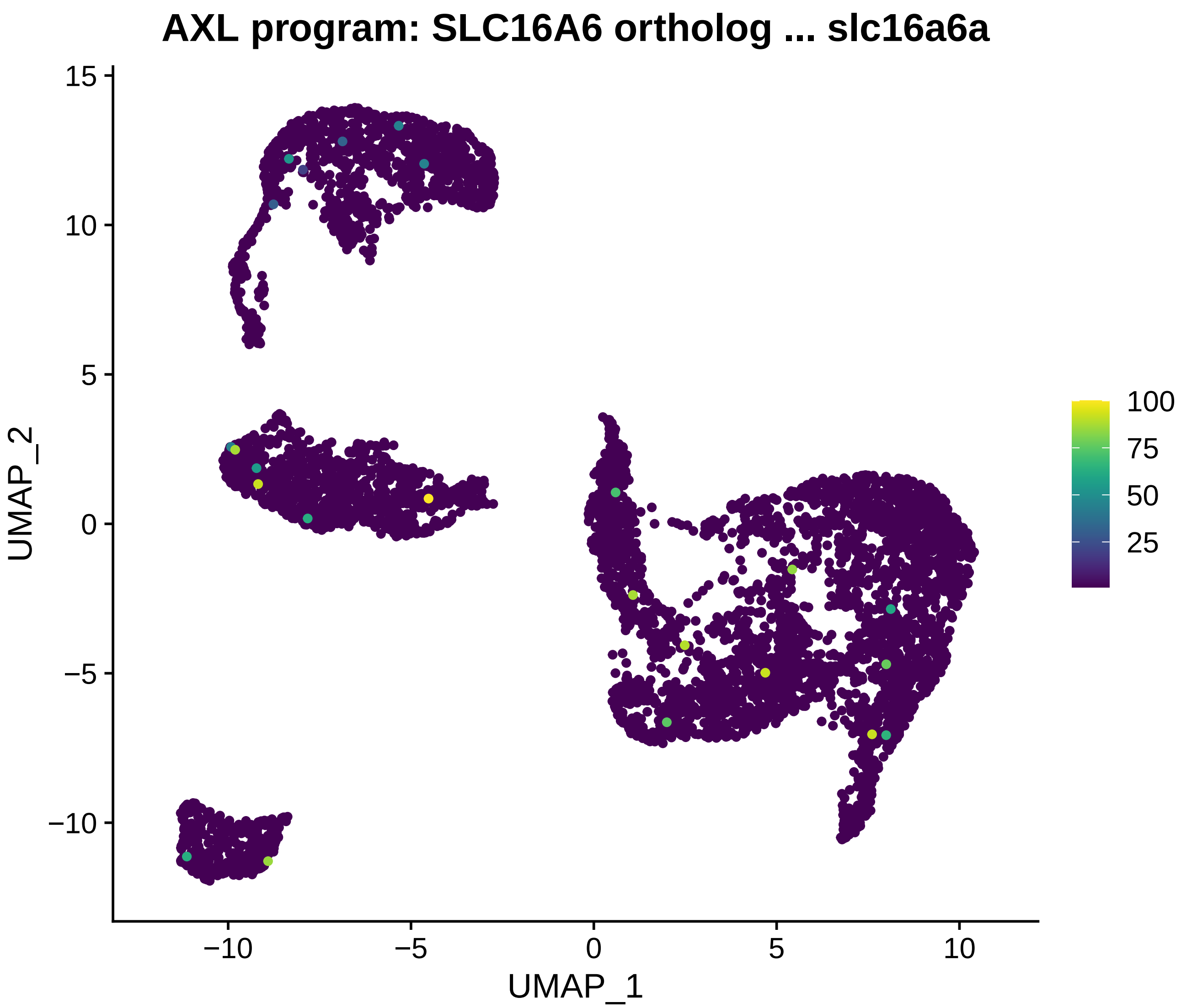
AXL program: SLC16A6 ortholog ... slc16a6a
UMAP_2
UMAP_1
15
10
5
0
−5
−10
−10
−5
0
5
10
100
75
50
25

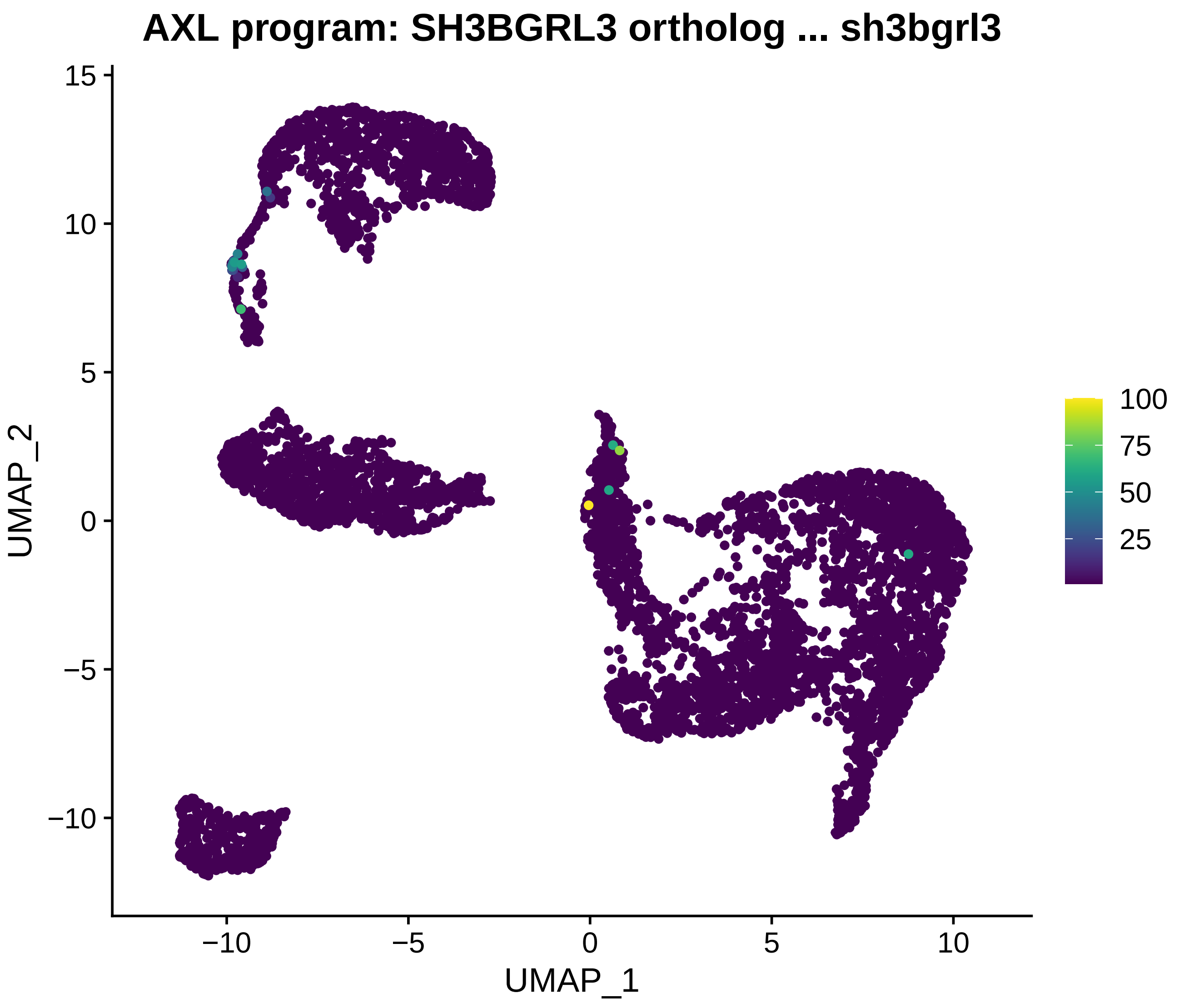
AXL program: SH3BGRL3 ortholog ... sh3bgrl3
UMAP_2
UMAP_1
15
10
5
0
−5
−10
−10
−5
0
5
10
100
75
50
25

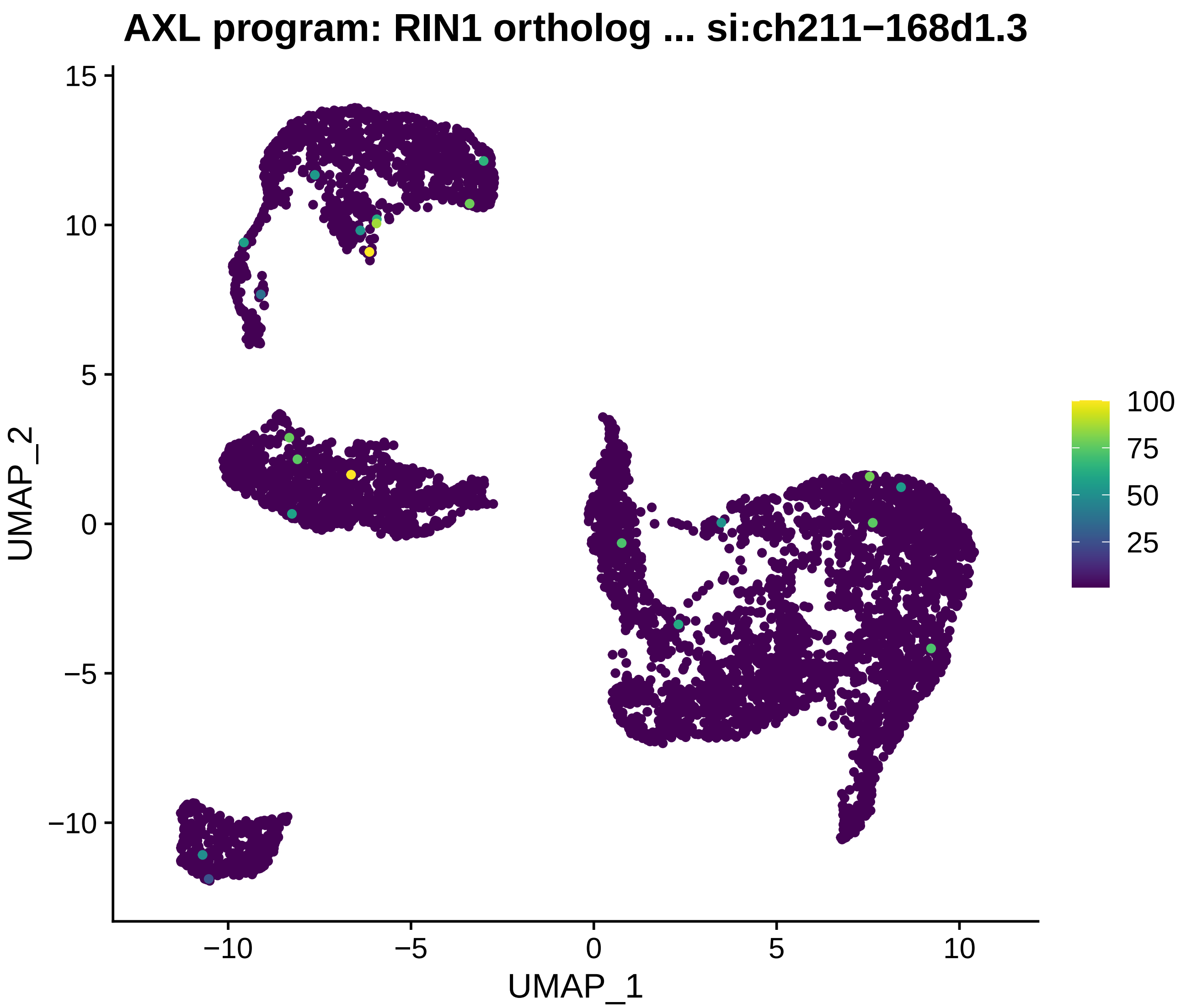
AXL program: RIN1 ortholog ... si:ch211−168d1.3
UMAP_2
UMAP_1
15
10
5
0
−5
−10
−10
−5
0
5
10
100
75
50
25

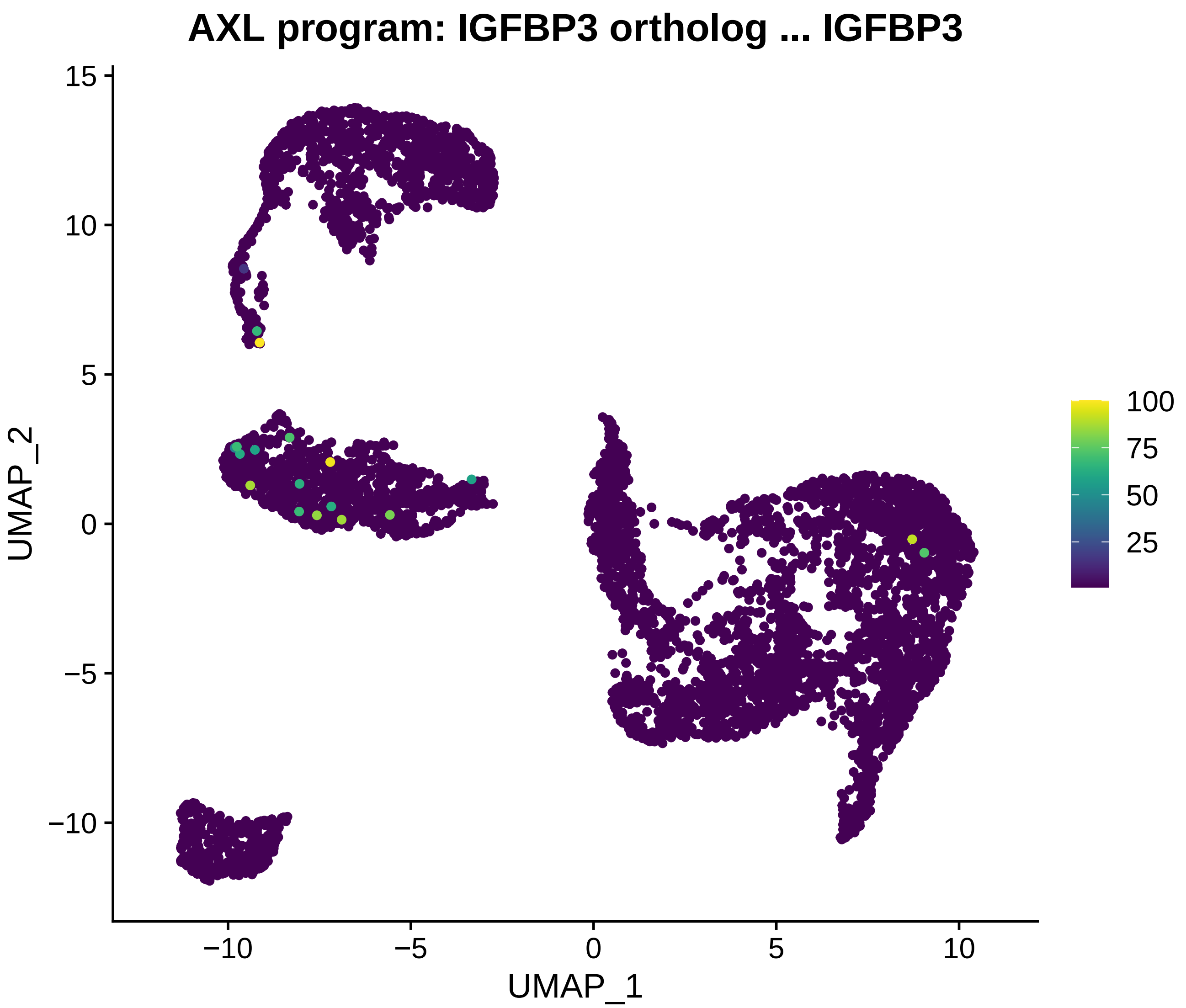
AXL program: IGFBP3 ortholog ... IGFBP3
UMAP_2
UMAP_1
15
10
5
0
−5
−10
−10
−5
0
5
10
100
75
50
25

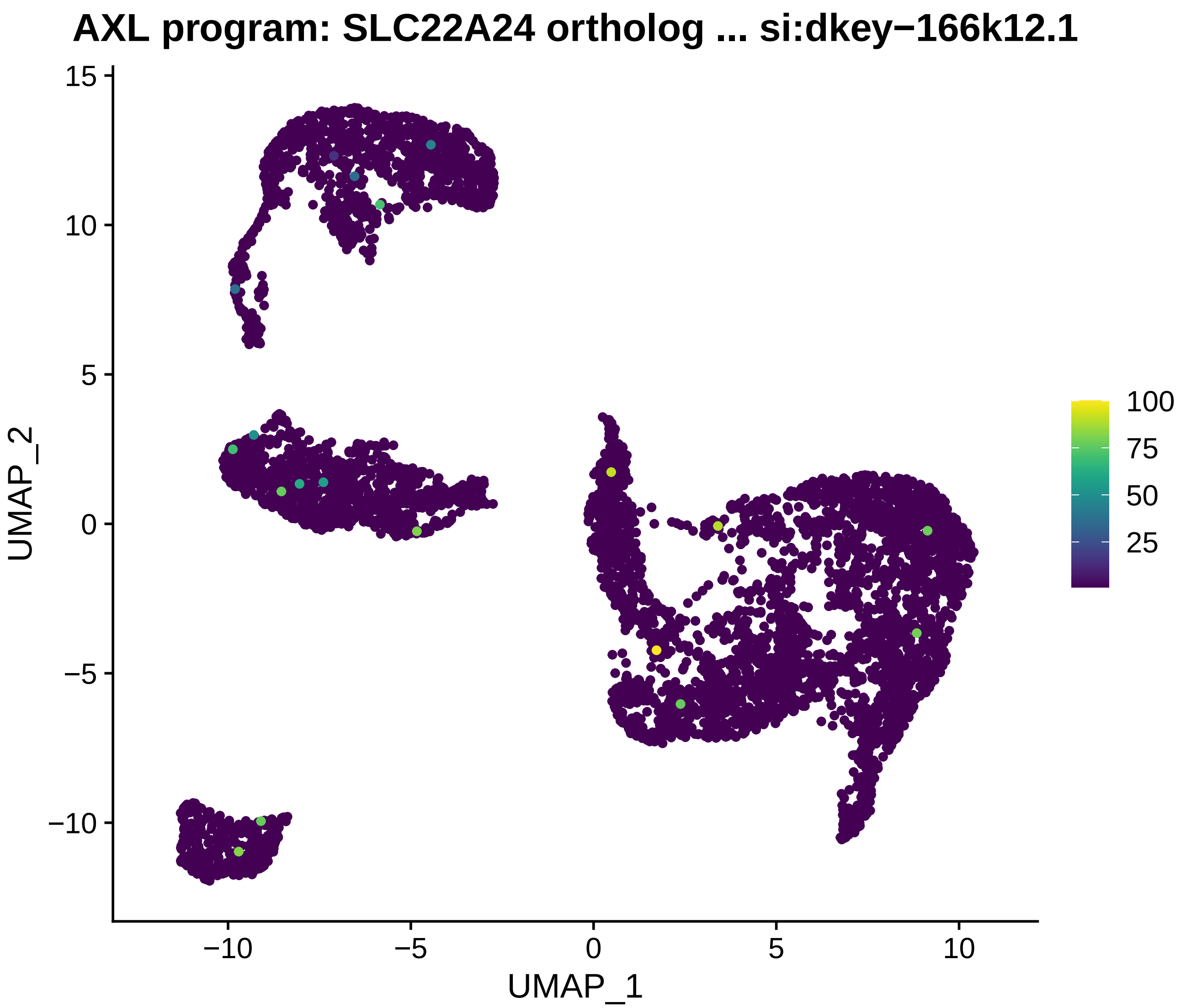
AXL program: SLC22A24 ortholog ... si:dkey−166k12.1
UMAP_1
UMAP_2
−10
−5
0
5
10
15
100
75
50
25

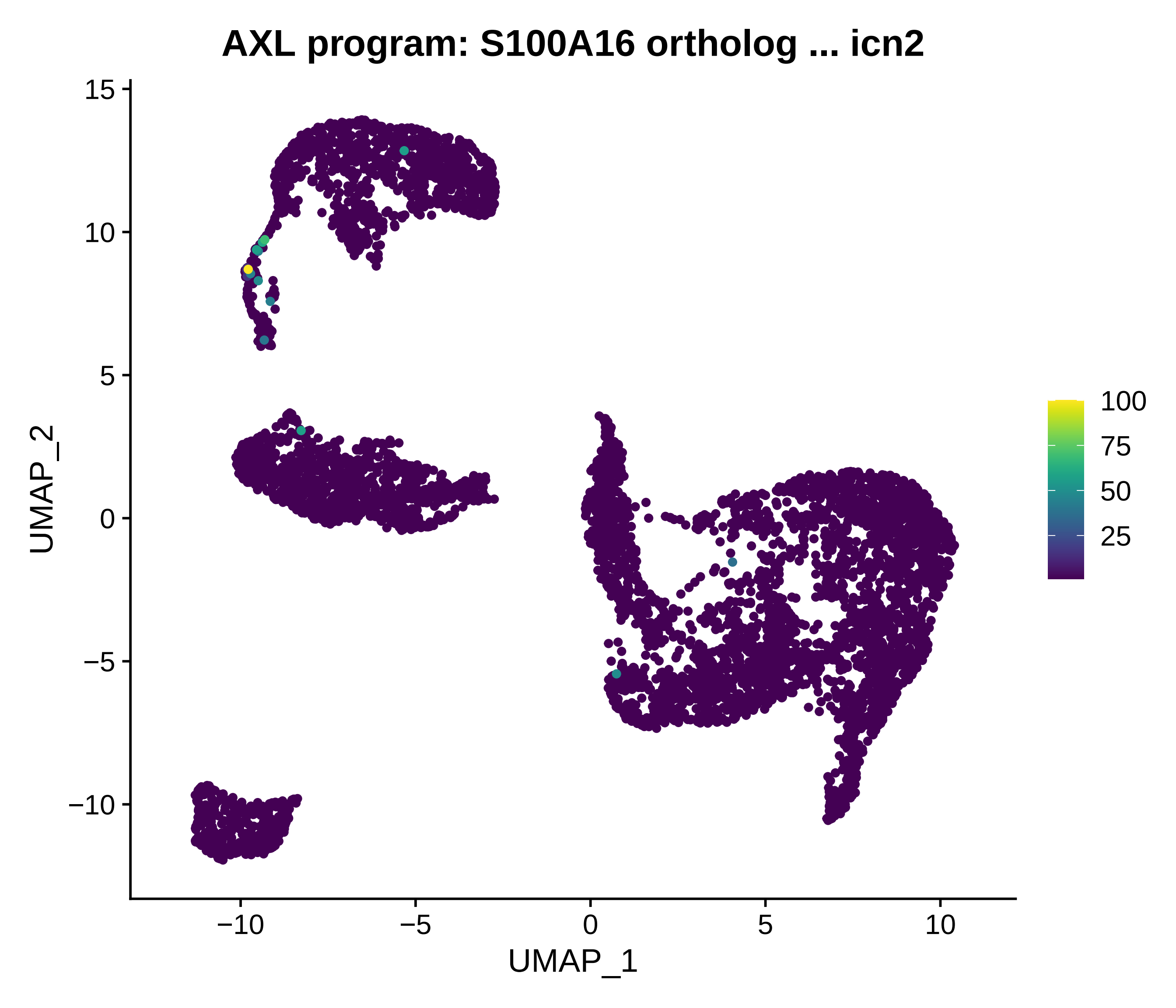
AXL program: S100A16 ortholog ... icn2
UMAP_1
UMAP_2
−10
−5
0
5
10
15
100
75
50
25

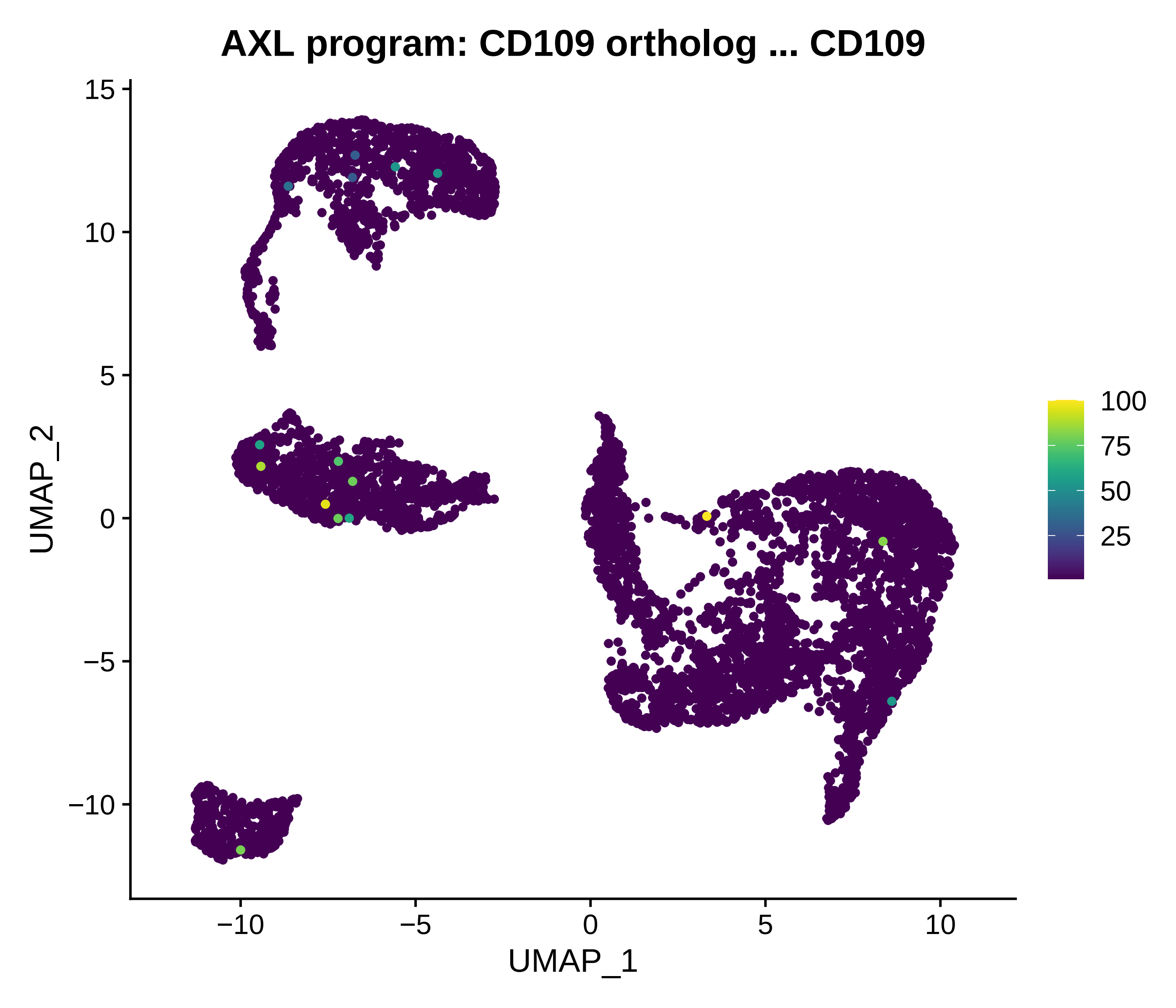
AXL program: CD109 ortholog ... CD109
UMAP_2
UMAP_1
15
10
5
0
−5
−10
−10
−5
0
5
10
100
75
50
25

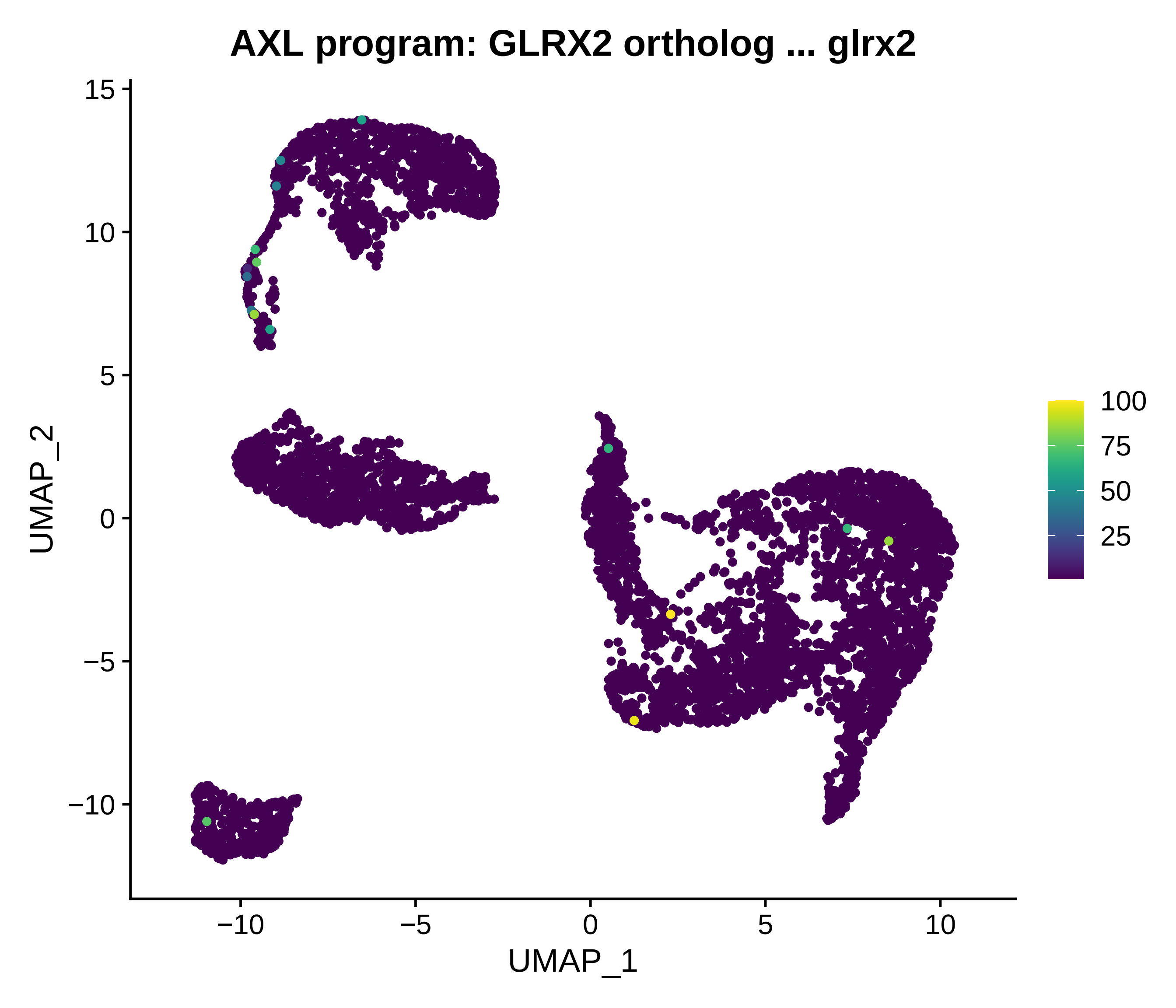
AXL program: GLRX2 ortholog ... glrx2
UMAP_1
UMAP_2
−10
−5
0
5
10
15
100
75
50
25

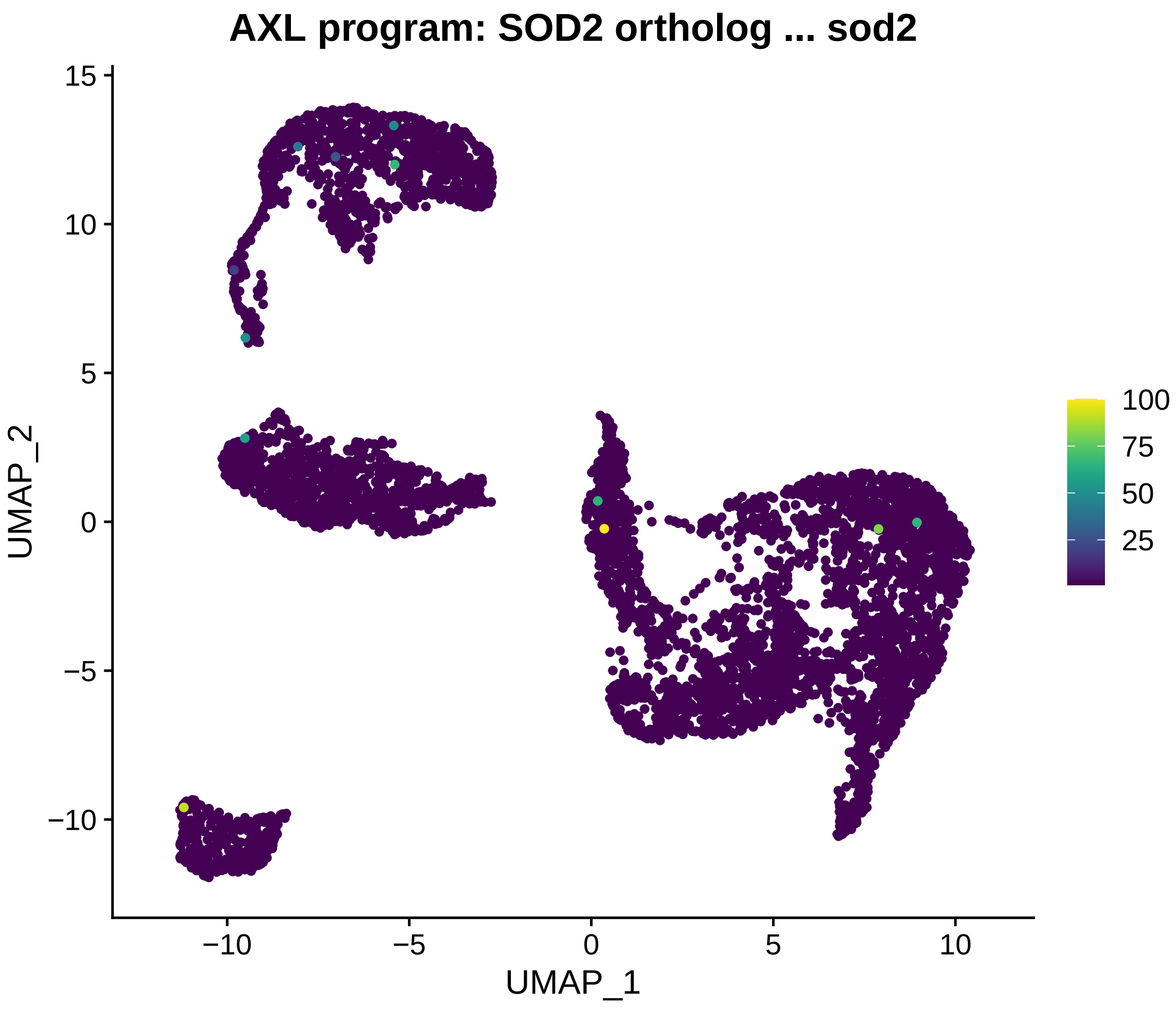
AXL program: SOD2 ortholog ... sod2
UMAP_2
UMAP_1
15
10
5
0
−5
−10
−10
−5
0
5
10
100
75
50
25

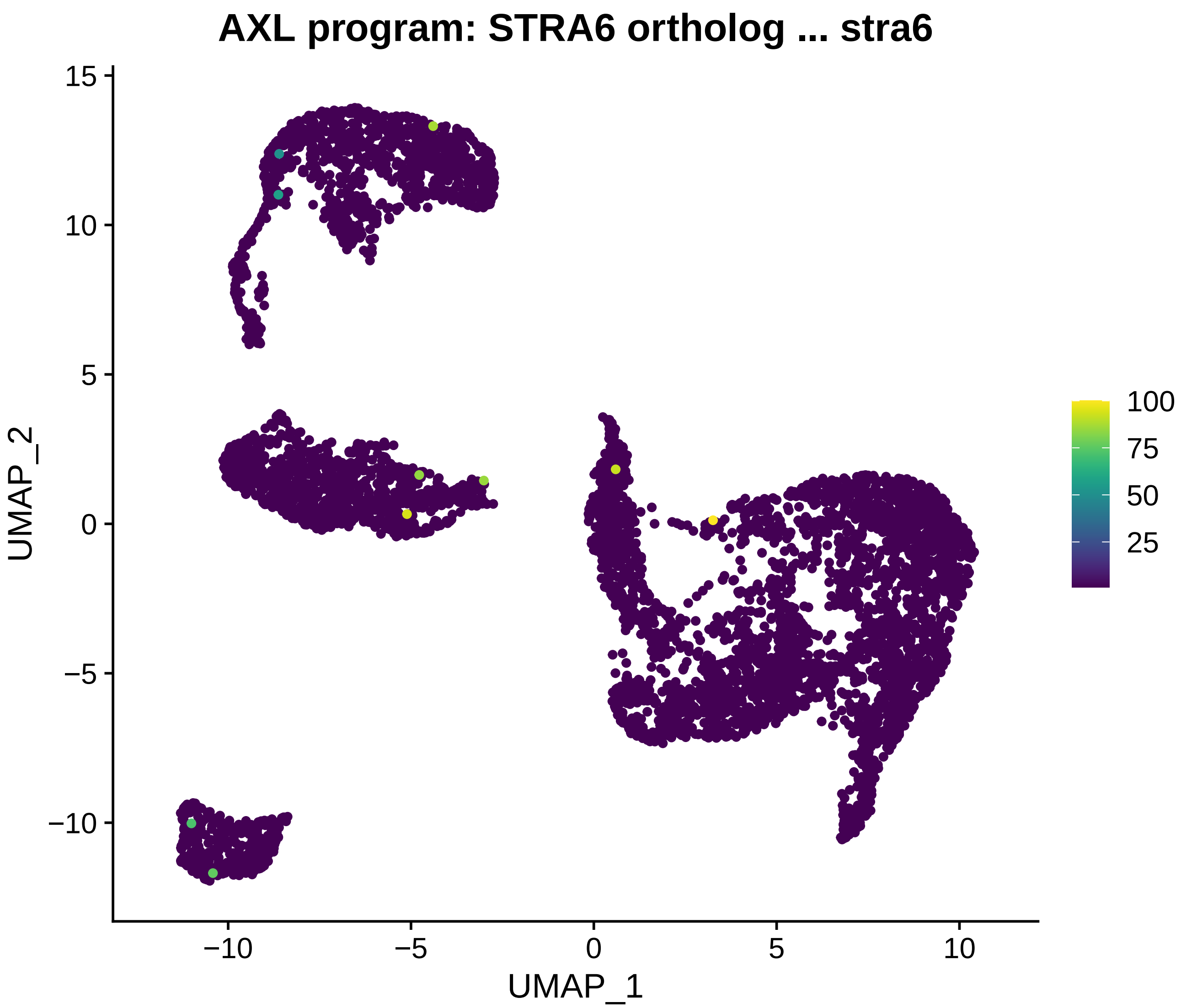
AXL program: STRA6 ortholog ... stra6
UMAP_2
15
10
5
0
−5
−10
UMAP_1
−10
−5
0
5
10
100
75
50
25
