## Supplementary material for "Comparison of Human Melanoma Single Cell Profiles to Evolutionary Medicine Model *Xiphophorus* Provides Insights in Disease Control": Supp. files: UMAP_MITF_orthologs_expressed.pdf

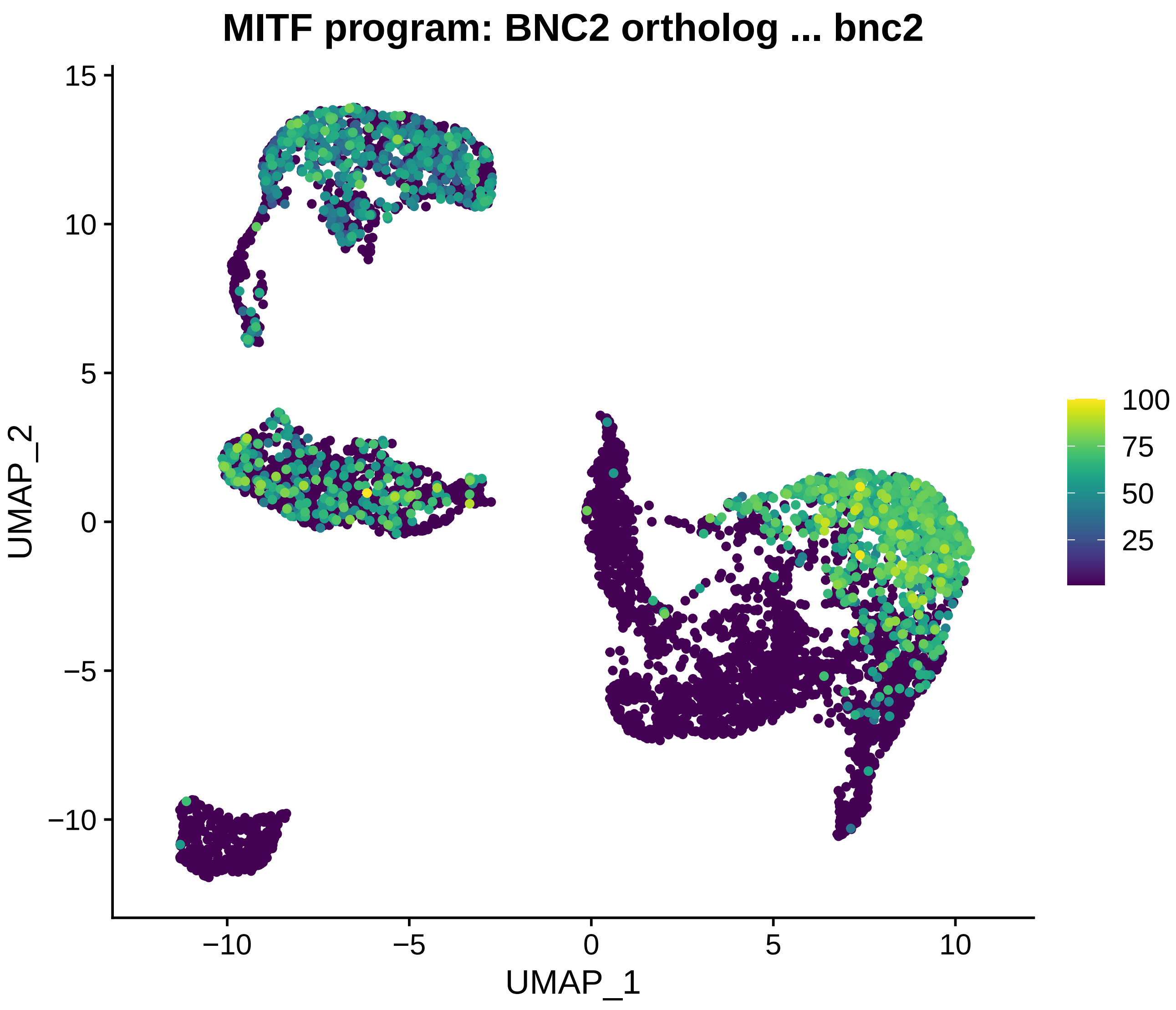

MITF program: BNC2 ortholog ... bnc2
15
10
5
0
−5
−10
UMAP_2
−10
−5
0
5
10
UMAP_1
100
75
50
25

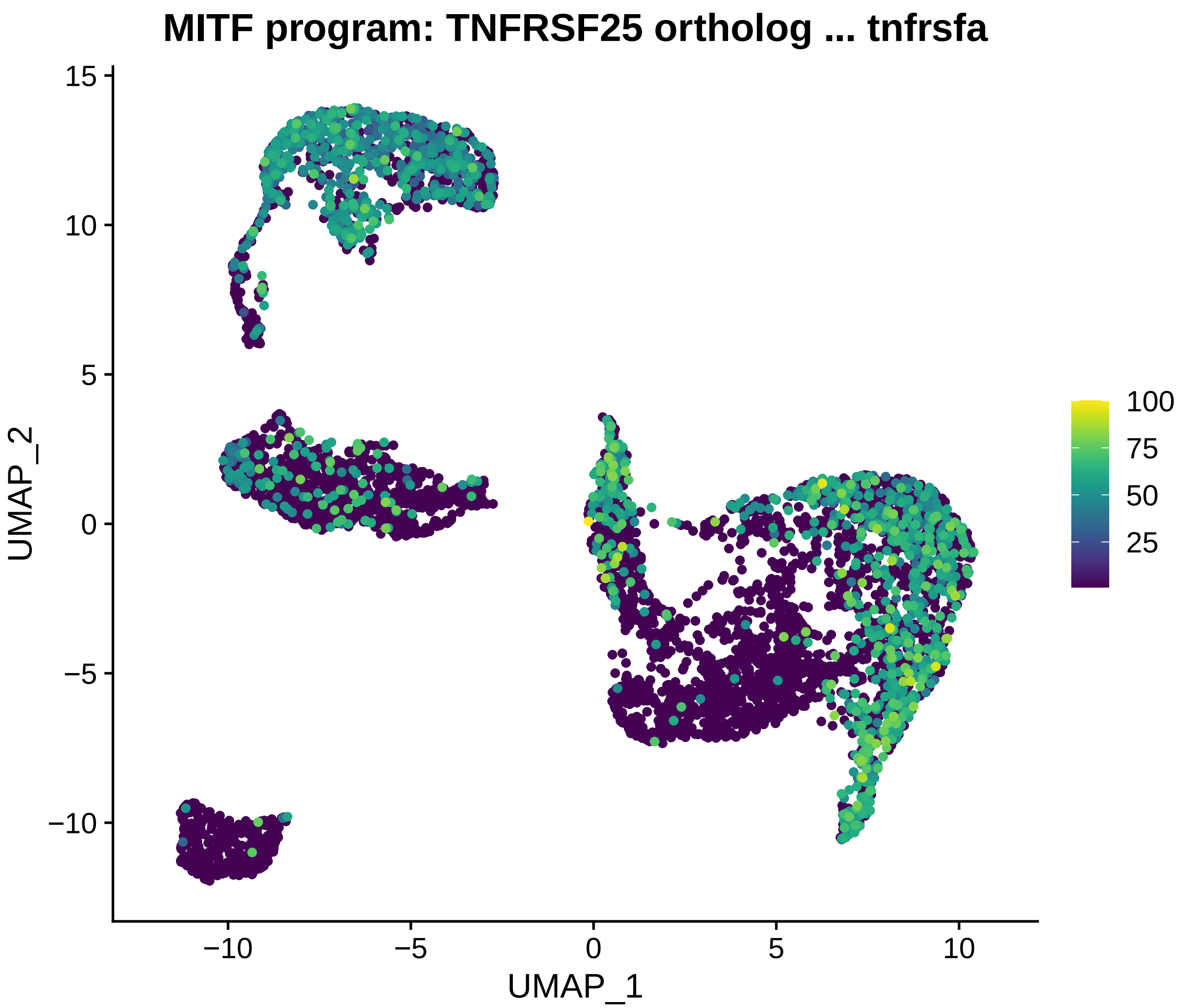
MITF program: TNFRSF25 ortholog ... tnfrsfa
UMAP_1
UMAP_2
−10
−5
0
5
10
15
100
75
50
25

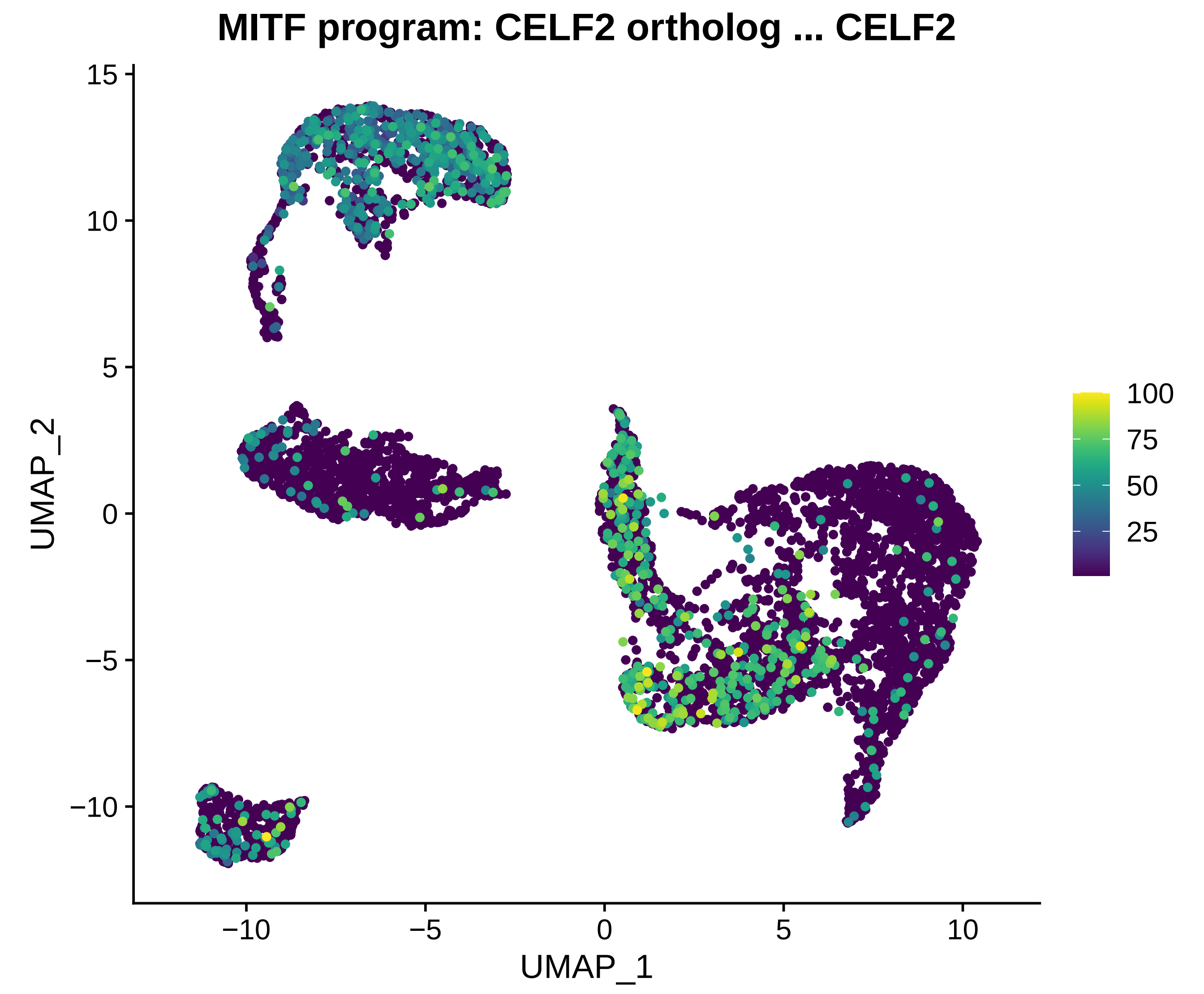
MITF program: CELF2 ortholog ... CELF2
UMAP_2
UMAP_1
15
10
5
0
−5
−10
−10
−5
0
5
10
100
75
50
25

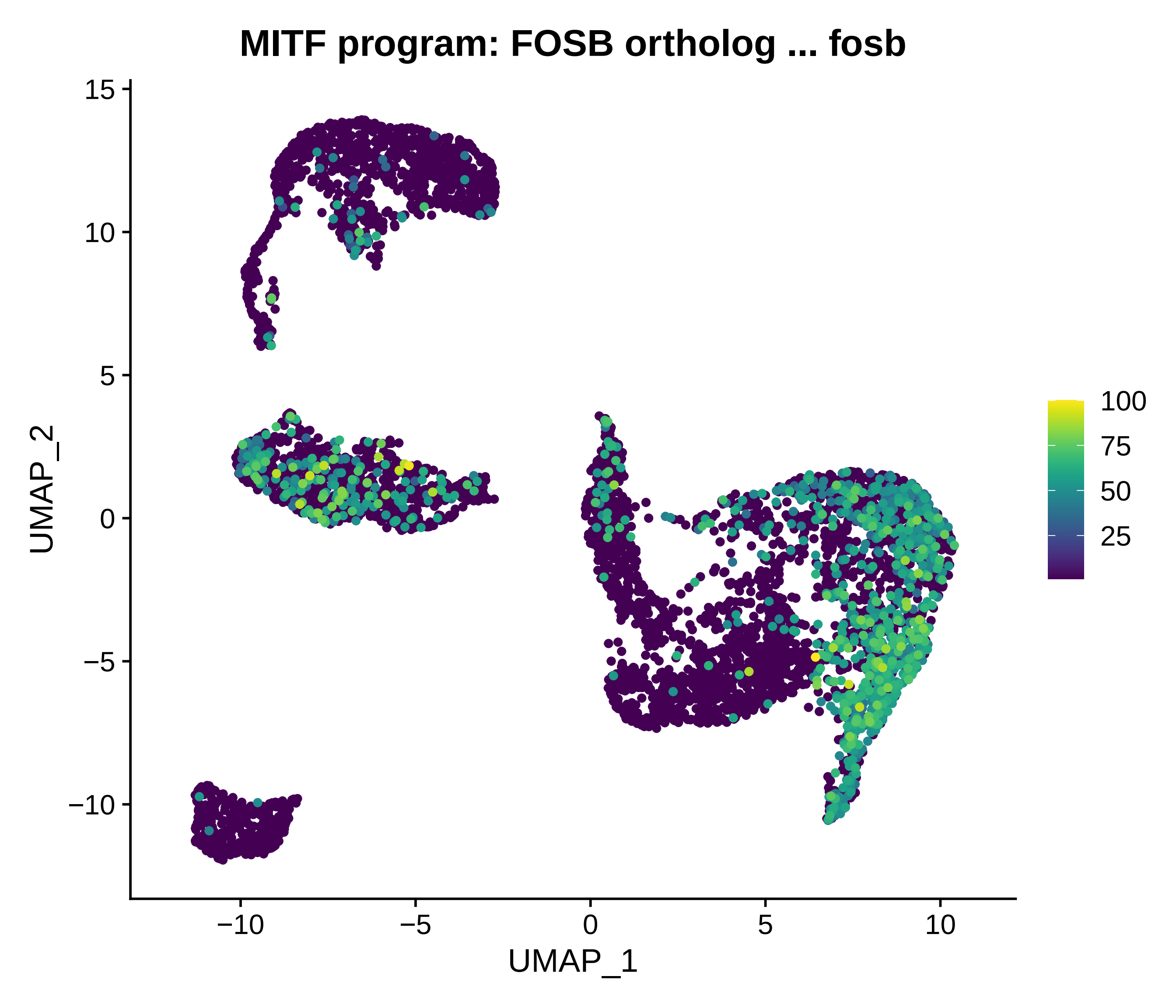

MITF program: FOSB ortholog ... fosb
UMAP_2
UMAP_1
15
10
5
0
−5
−10
−10
−5
0
5
10
100
75
50
25

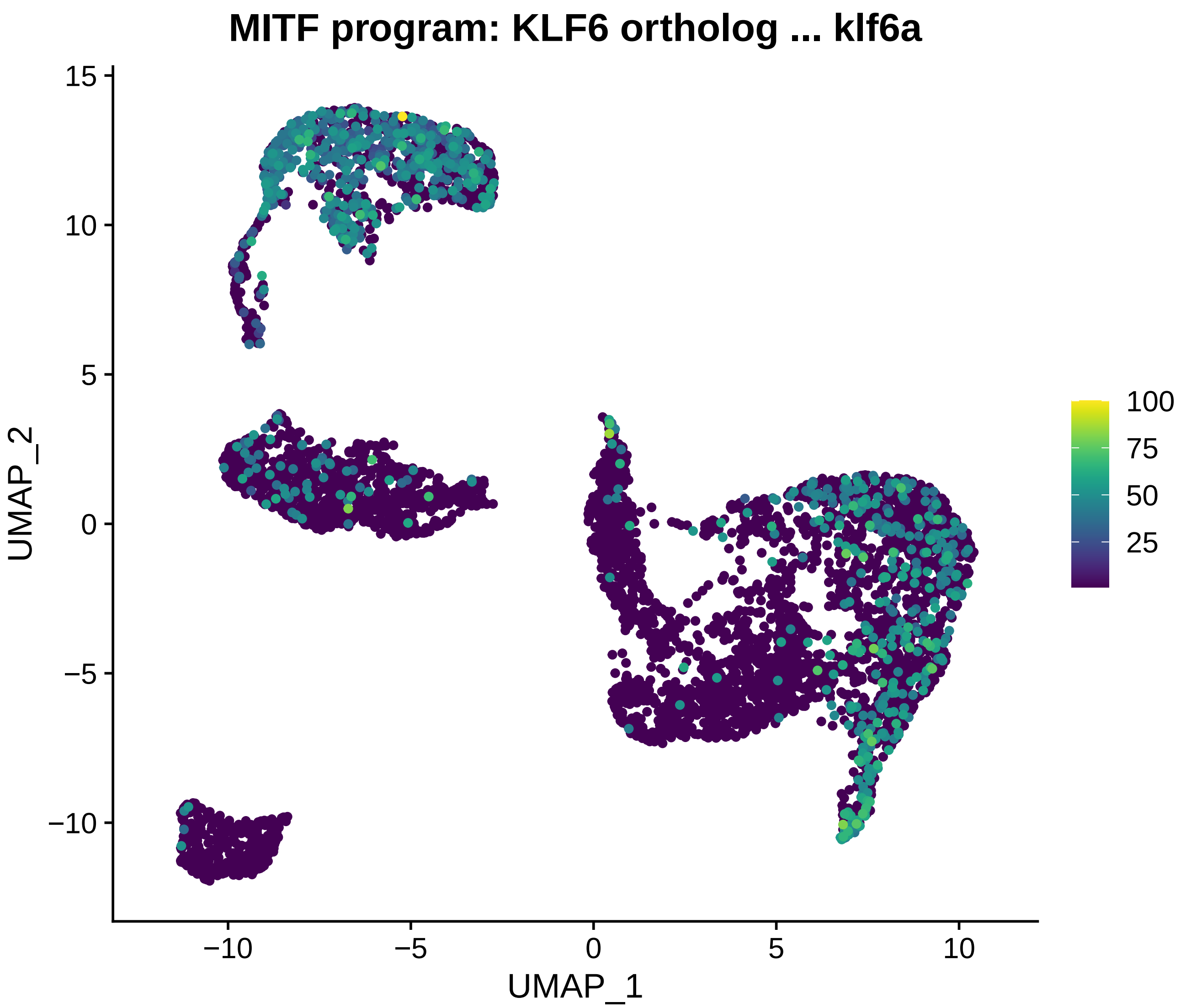
MITF program: KLF6 ortholog ... klf6a
UMAP_2
UMAP_1
15
10
5
0
−5
−10
−10
−5
0
5
10
100
75
50
25

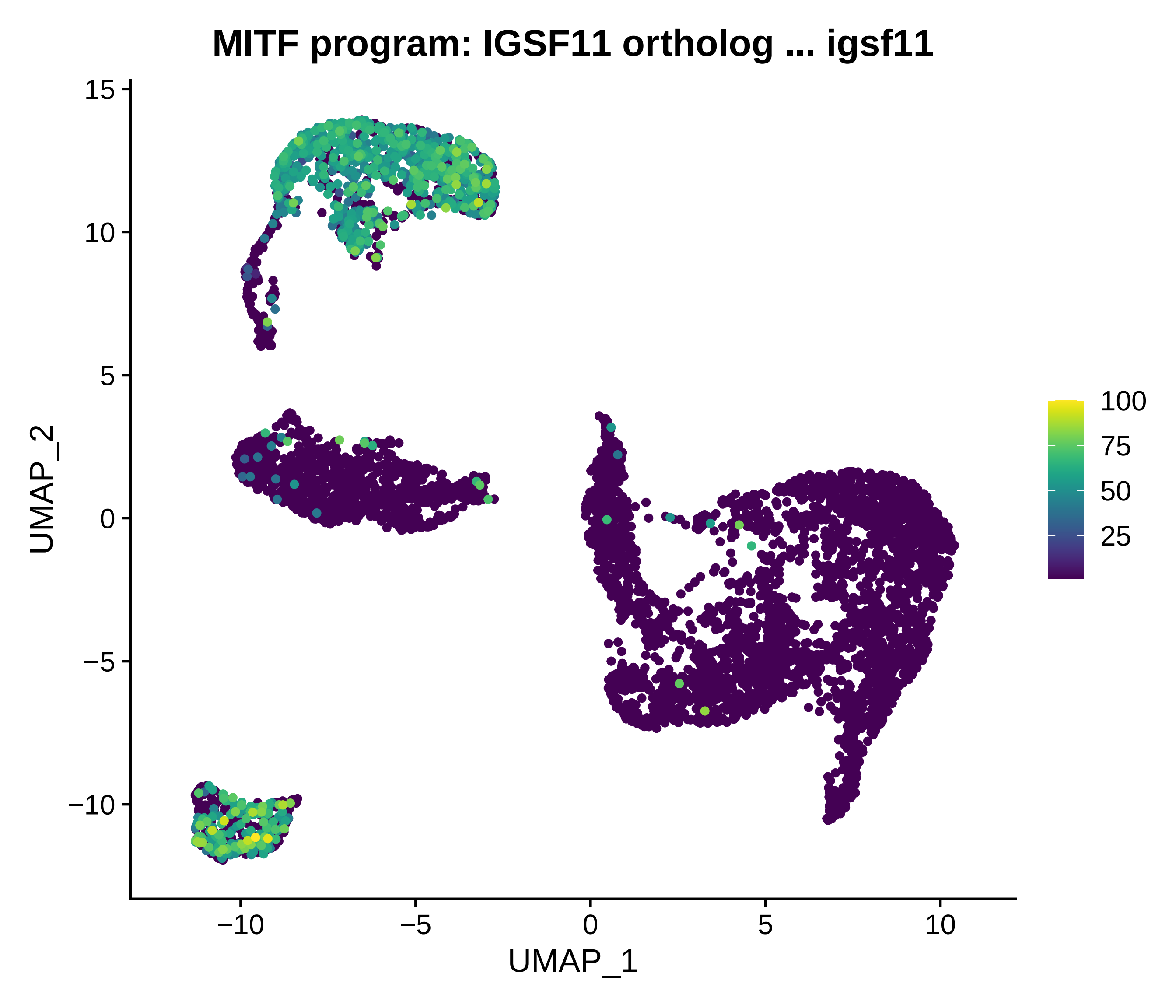
MITF program: IGSF11 ortholog ... igsf11
UMAP_2
15
10
5
0
−5
−10
UMAP_1
−10
−5
0
5
10
100
75
50
25

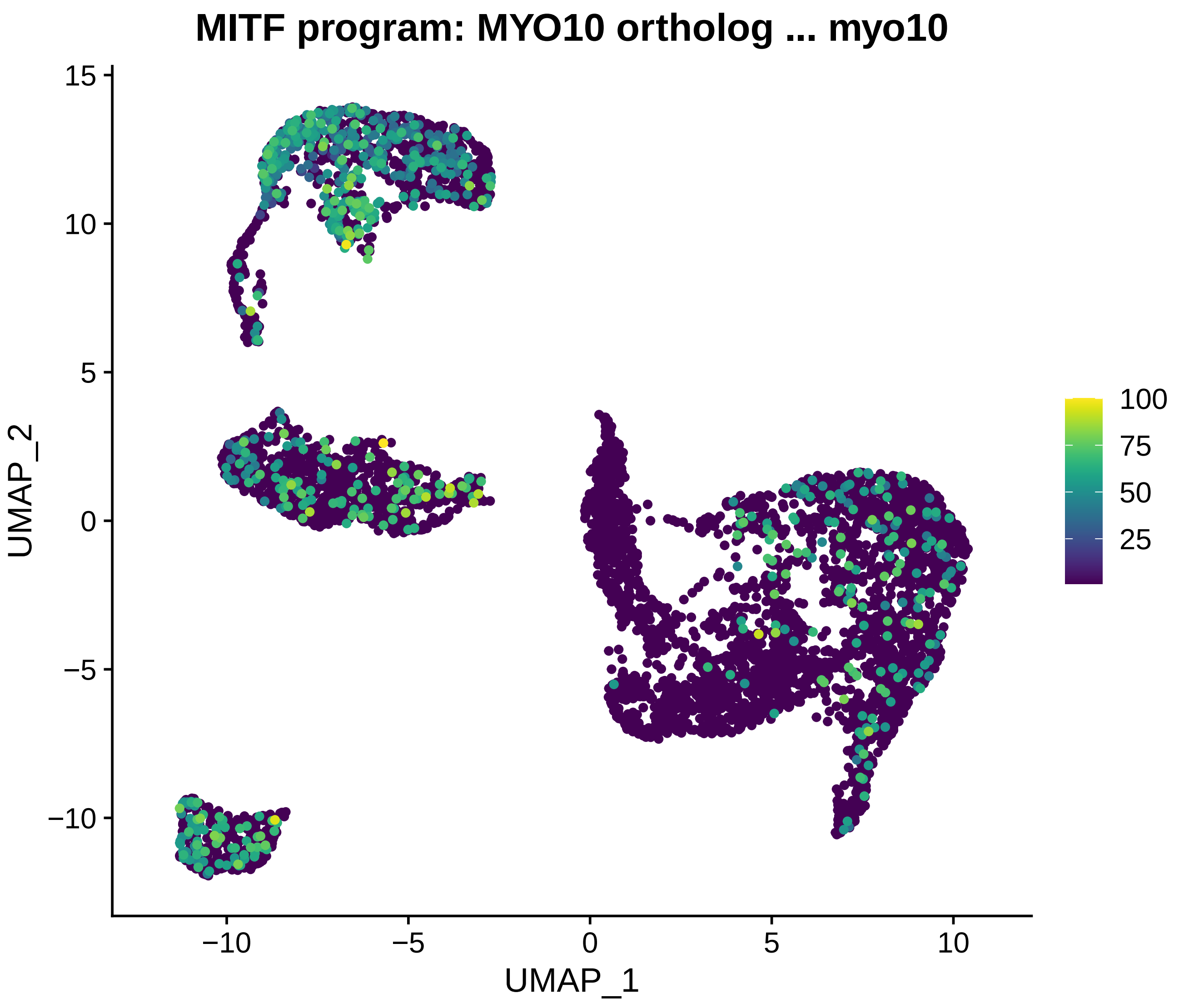
MITF program: MYO10 ortholog ... myo10
UMAP_2
UMAP_1
15
10
5
0
−5
−10
−10
−5
0
5
10
100
75
50
25

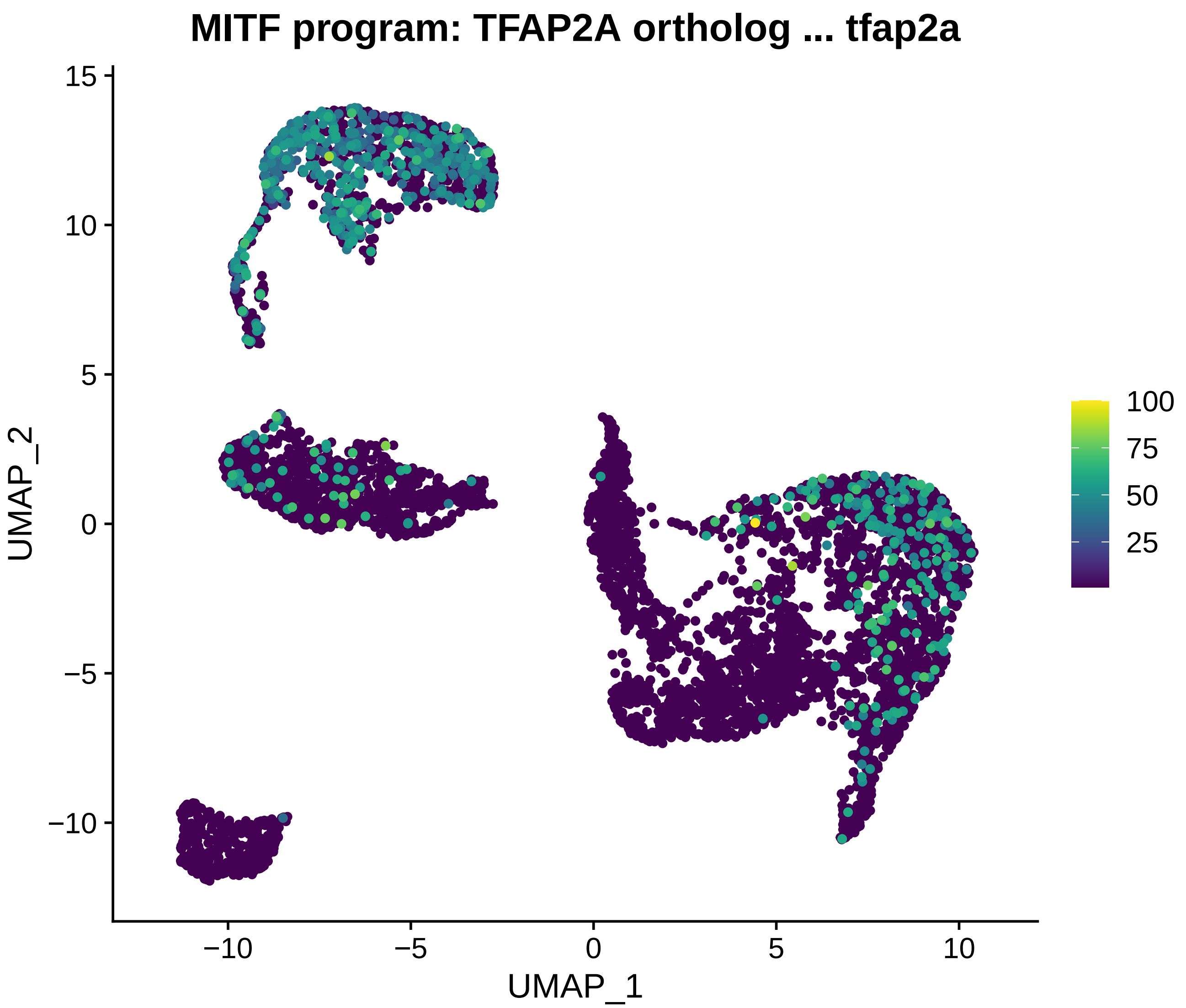
MITF program: TFAP2A ortholog ... tfap2a
UMAP_2
UMAP_1
15
10
5
0
−5
−10
−10
−5
0
5
10
100
75
50
25

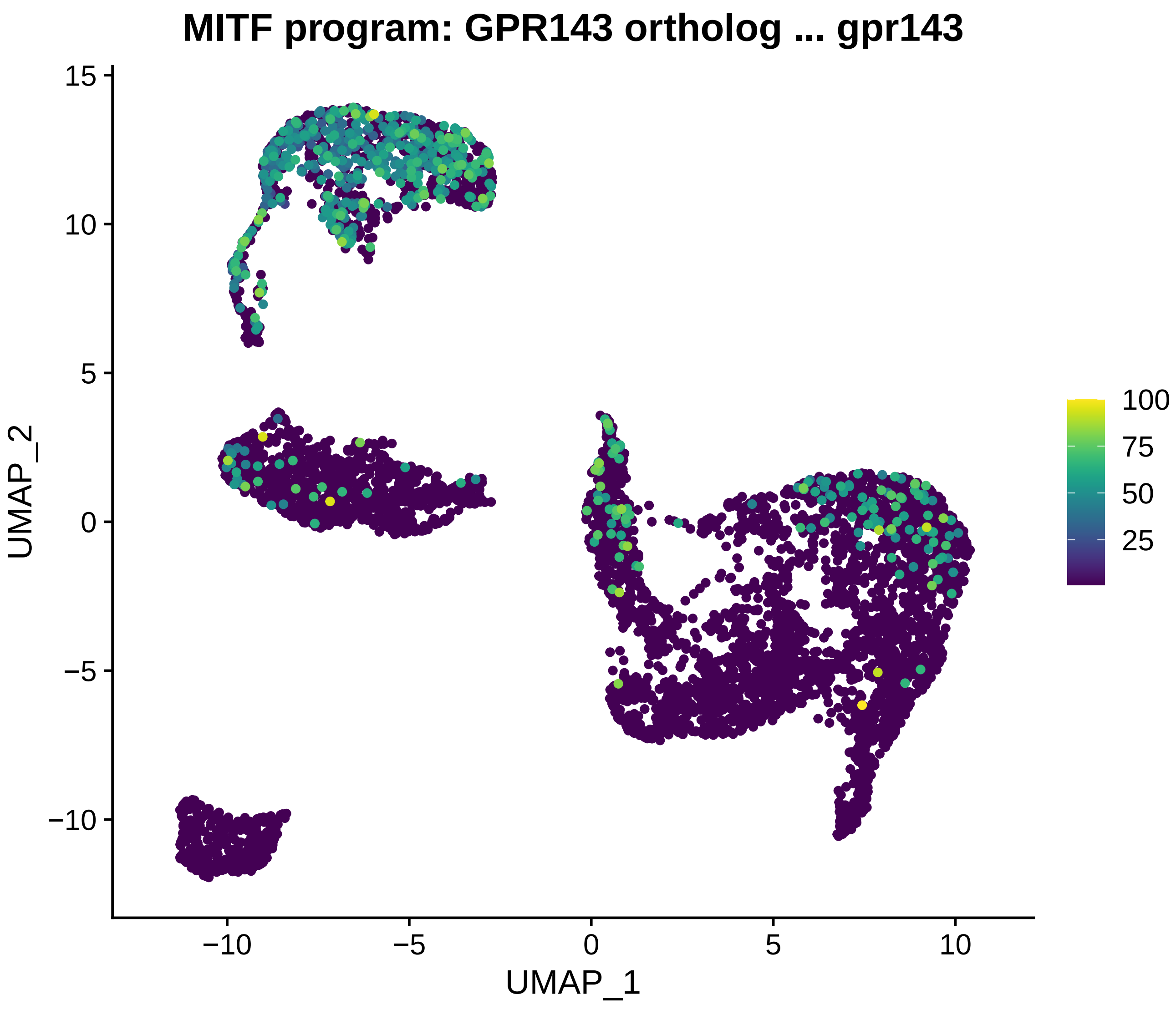
MITF program: GPR143 ortholog ... gpr143
UMAP_2
UMAP_1
15
10
5
0
−5
−10
−10
−5
0
5
10
100
75
50
25

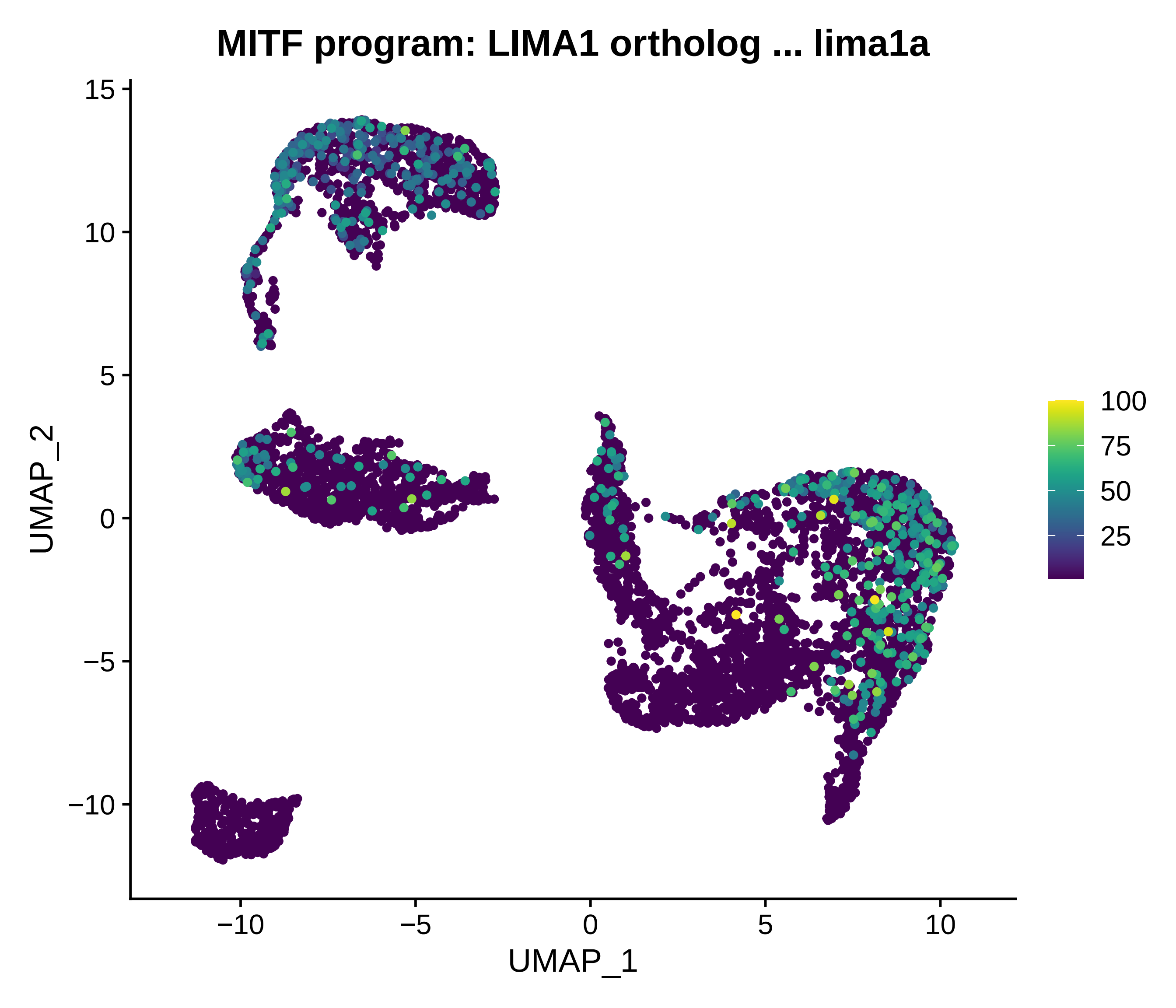

MITF program: LIMA1 ortholog ... lima1a
UMAP_2
UMAP_1
15
10
5
0
−5
−10
−10
−5
0
5
10
100
75
50
25

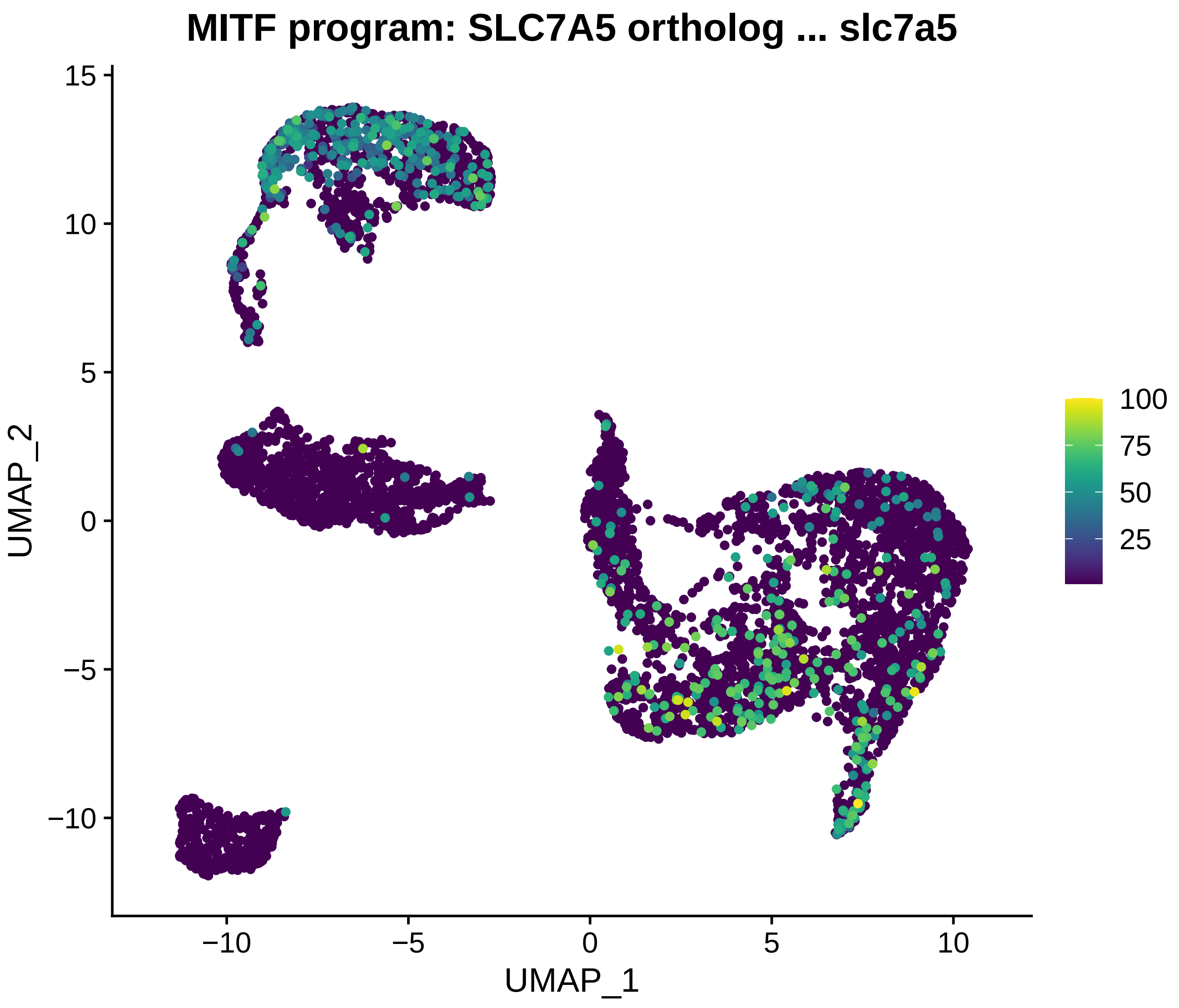
MITF program: SLC7A5 ortholog ... slc7a5
UMAP_2
UMAP_1
15
10
5
0
−5
−10
−10
−5
0
5
10
100
75
50
25

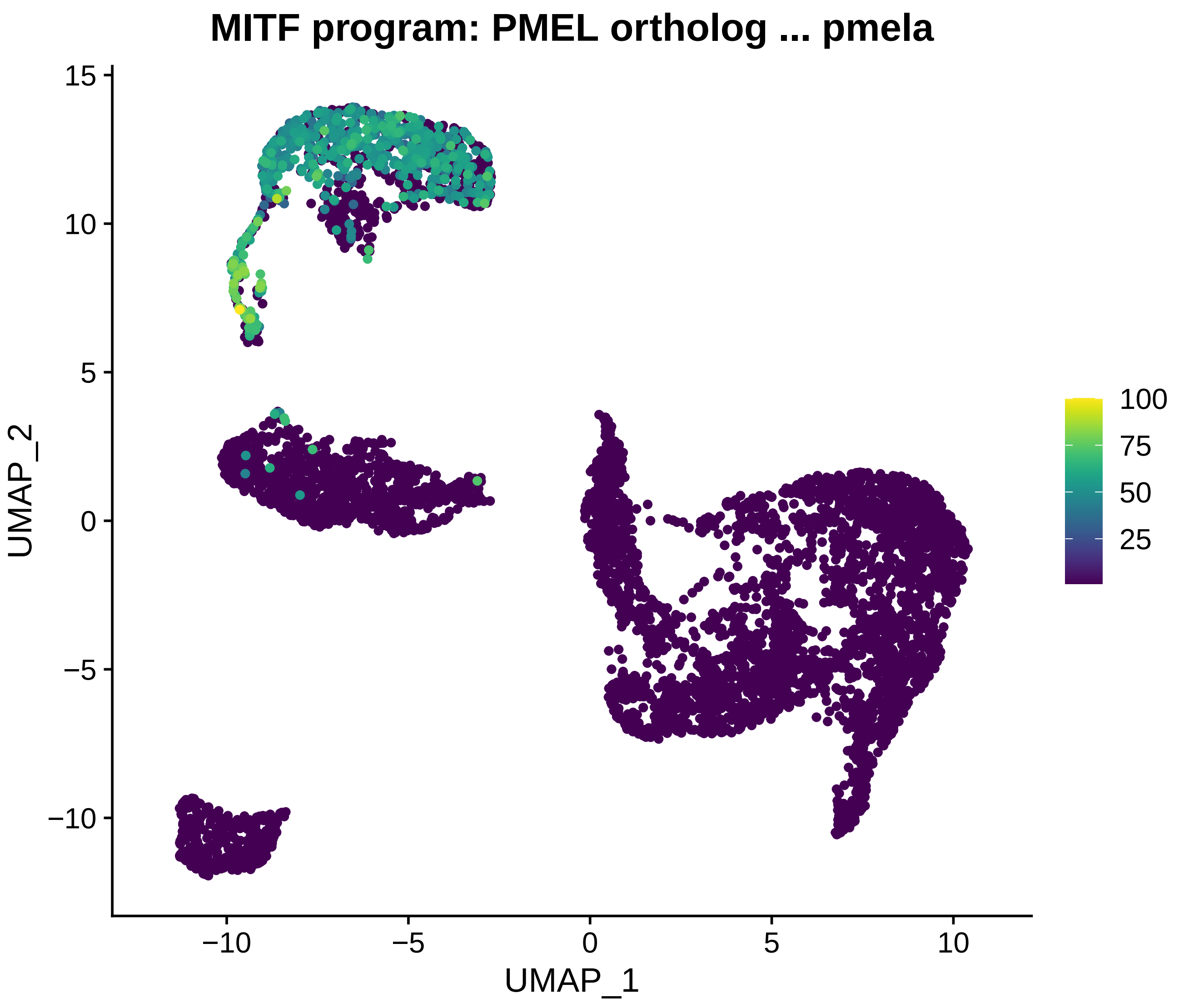
MITF program: PMEL ortholog ... pmela
UMAP_2
UMAP_1
15
10
5
0
−5
−10
−10
−5
0
5
10
100
75
50
25

### MITF program: PLK2 ortholog ... plk2b

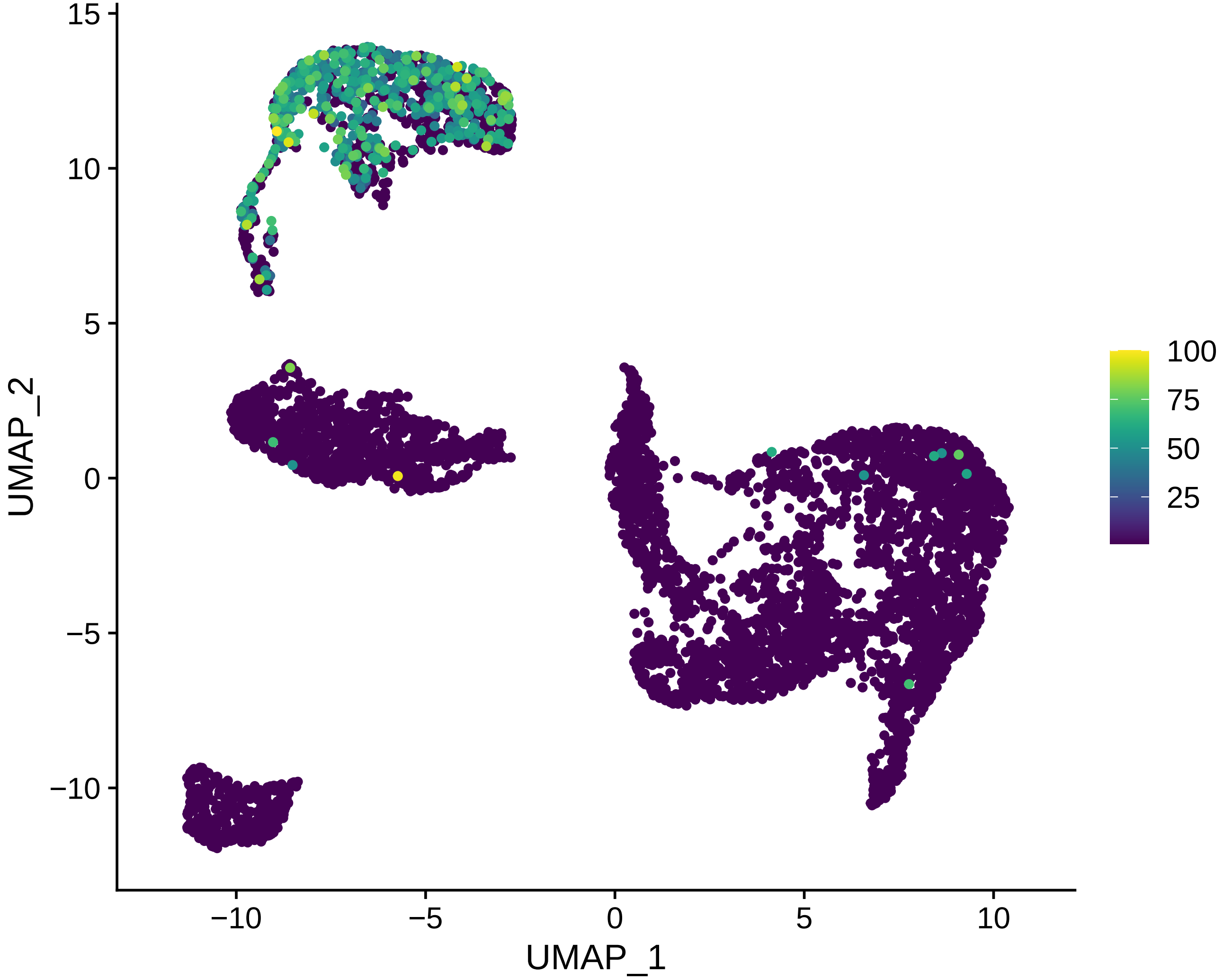

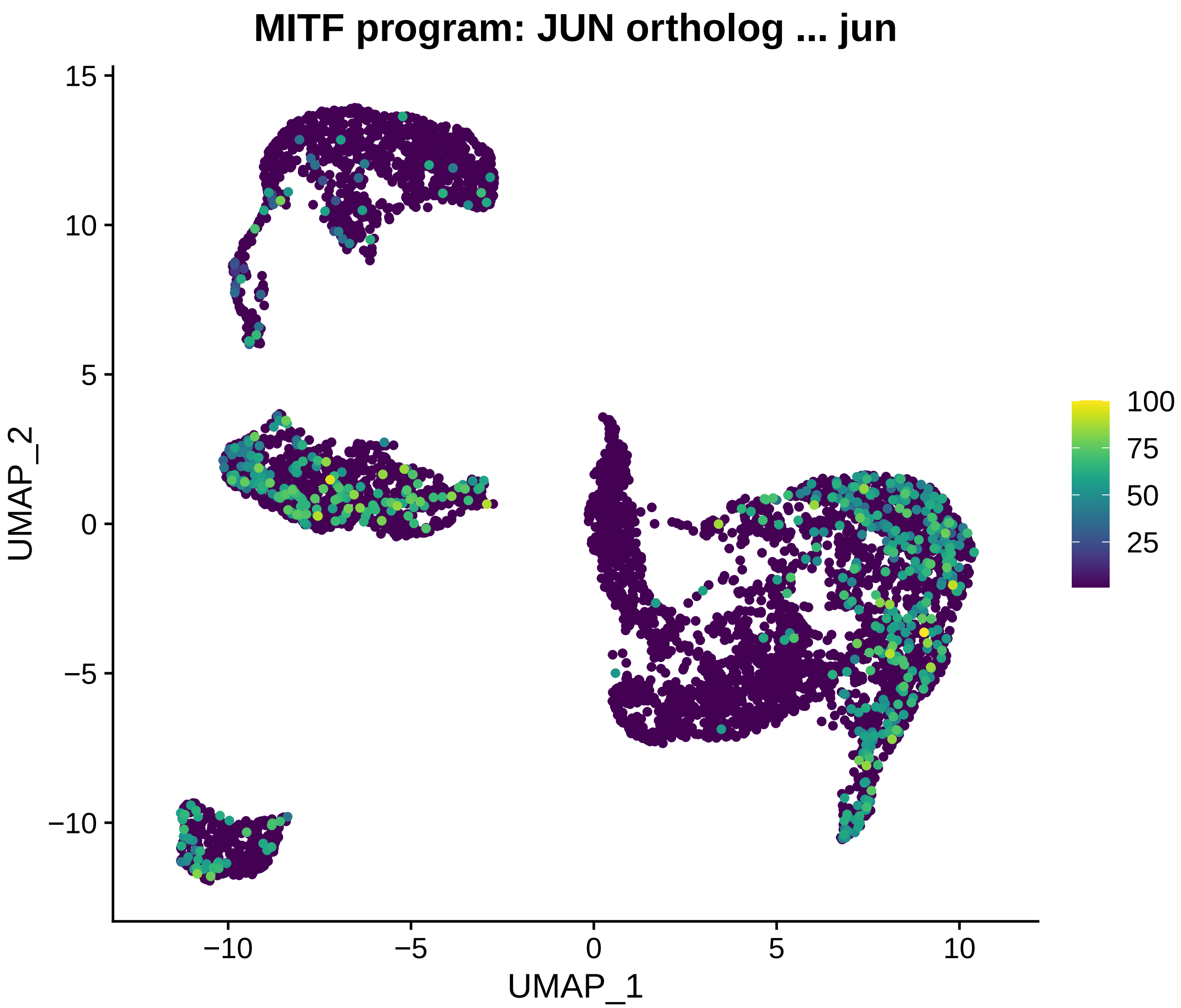
MITF program: JUN ortholog ... jun
UMAP_1
UMAP_2
−10
−5
0
5
10
15
100
75
50
25

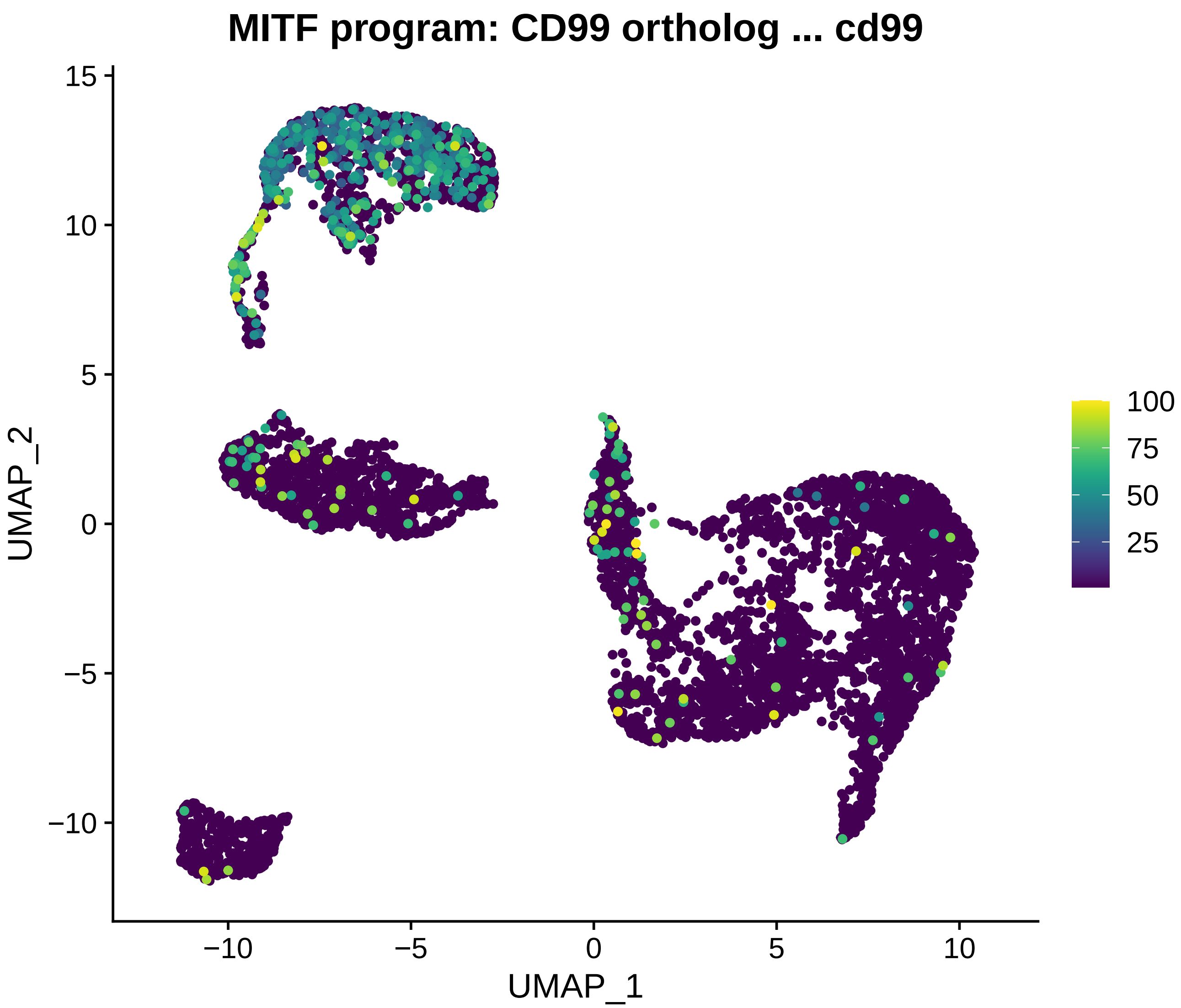
MITF program: CD99 ortholog ... cd99
UMAP_2
UMAP_1
15
10
5
0
−5
−10
−10
−5
0
5
10
100
75
50
25

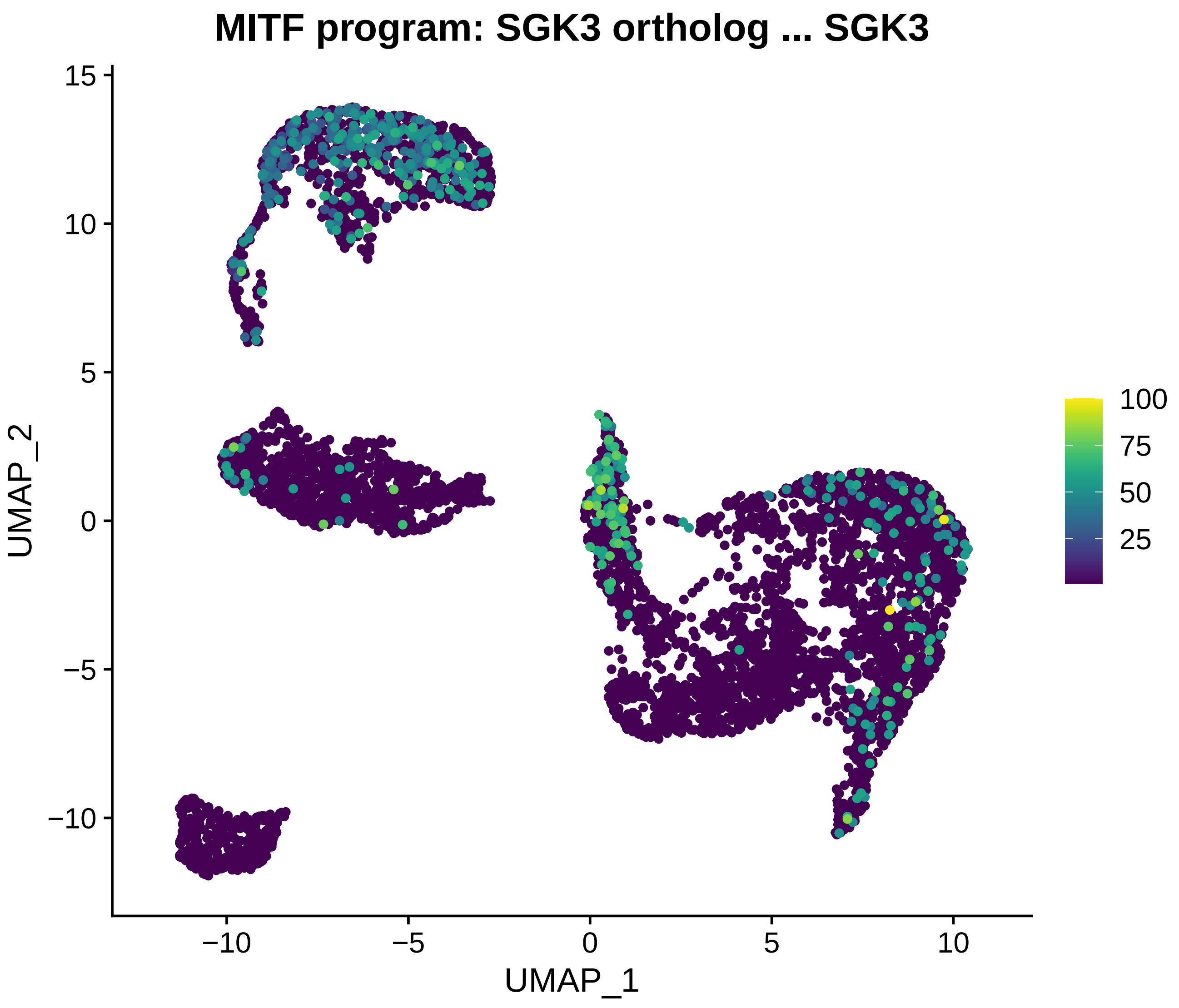
MITF program: SGK3 ortholog ... SGK3
UMAP_2
UMAP_1
15
10
5
0
−5
−10
−10
−5
0
5
10
100
75
50
25

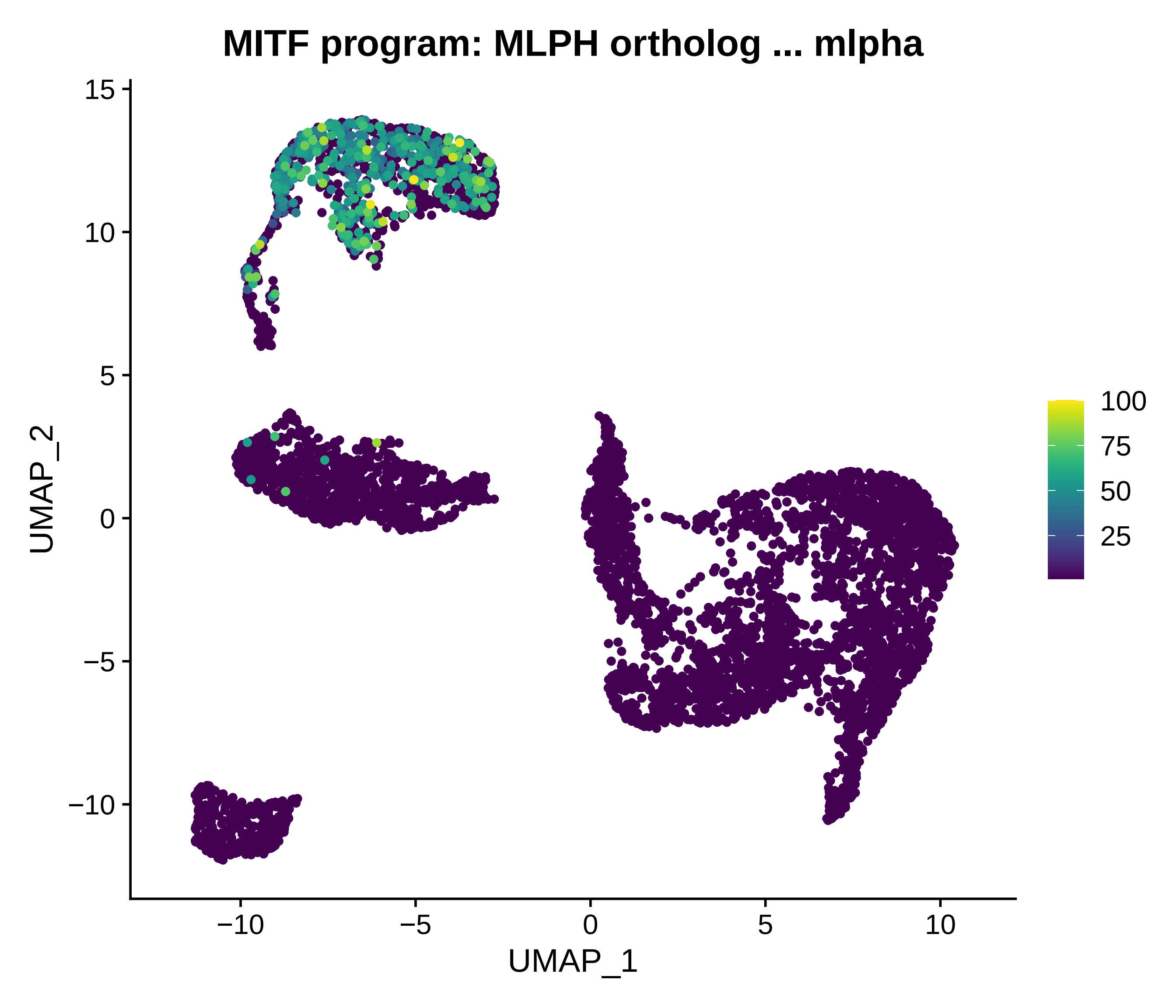

MITF program: MLPH ortholog ... mlpha
UMAP_2
UMAP_1
15
10
5
0
−5
−10
−10
−5
0
5
10
100
75
50
25

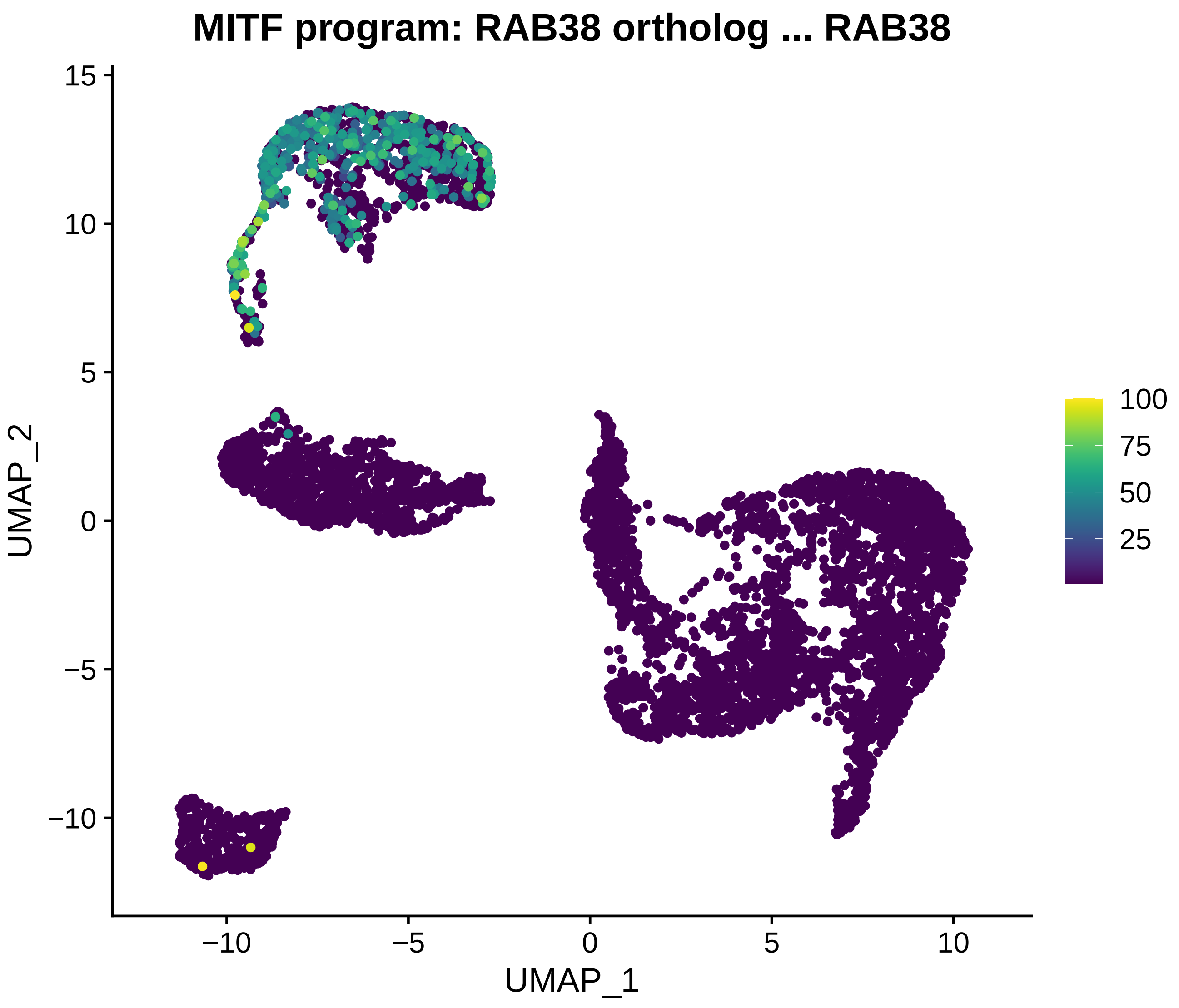
MITF program: RAB38 ortholog ... RAB38
UMAP_1
UMAP_2
−10
−5
0
5
10
15
100
75
50
25

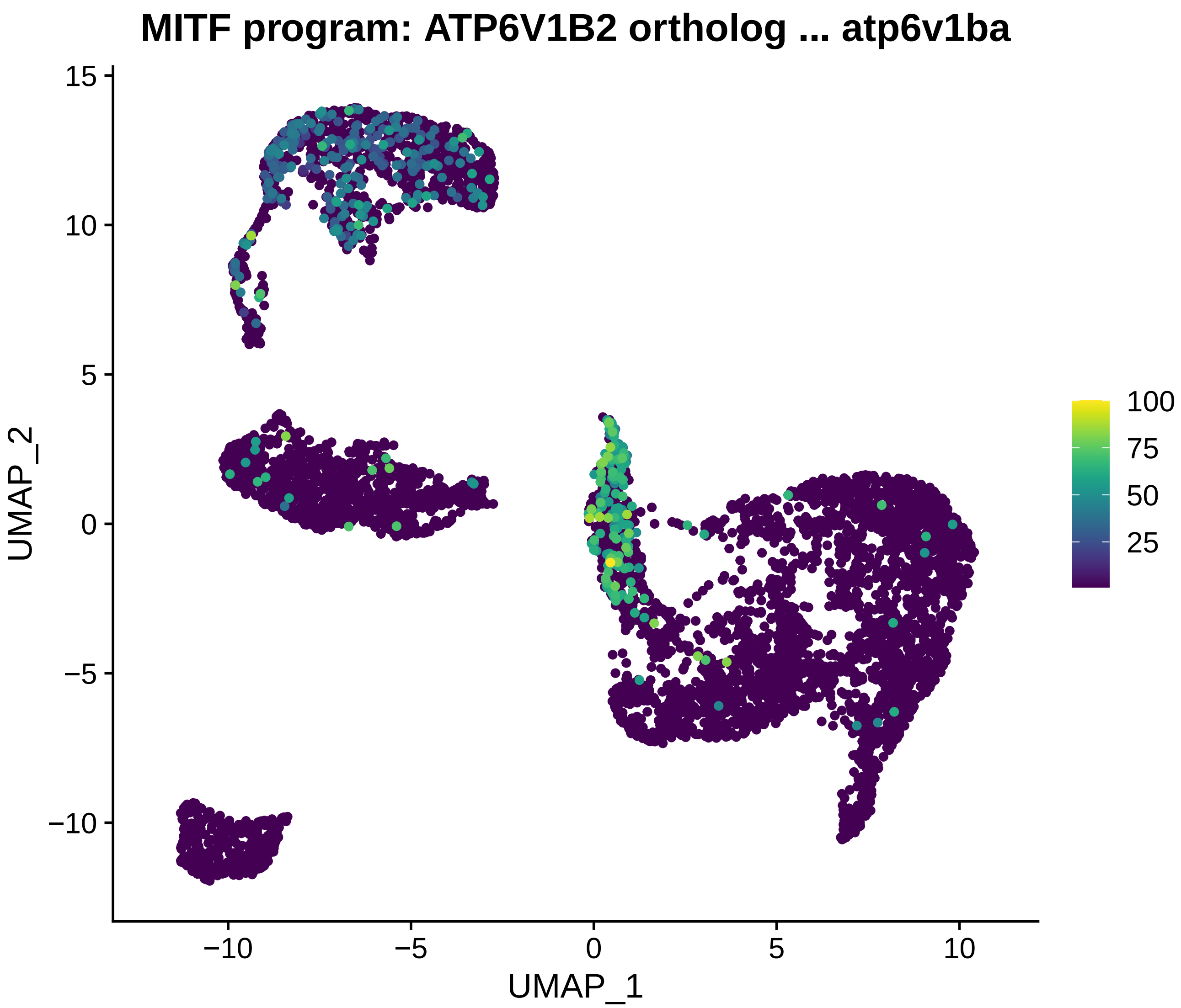

MITF program: ATP6V1B2 ortholog ... atp6v1ba
UMAP_2
UMAP_1
15
10
5
0
−5
−10
−10
−5
0
5
10
100
75
50
25

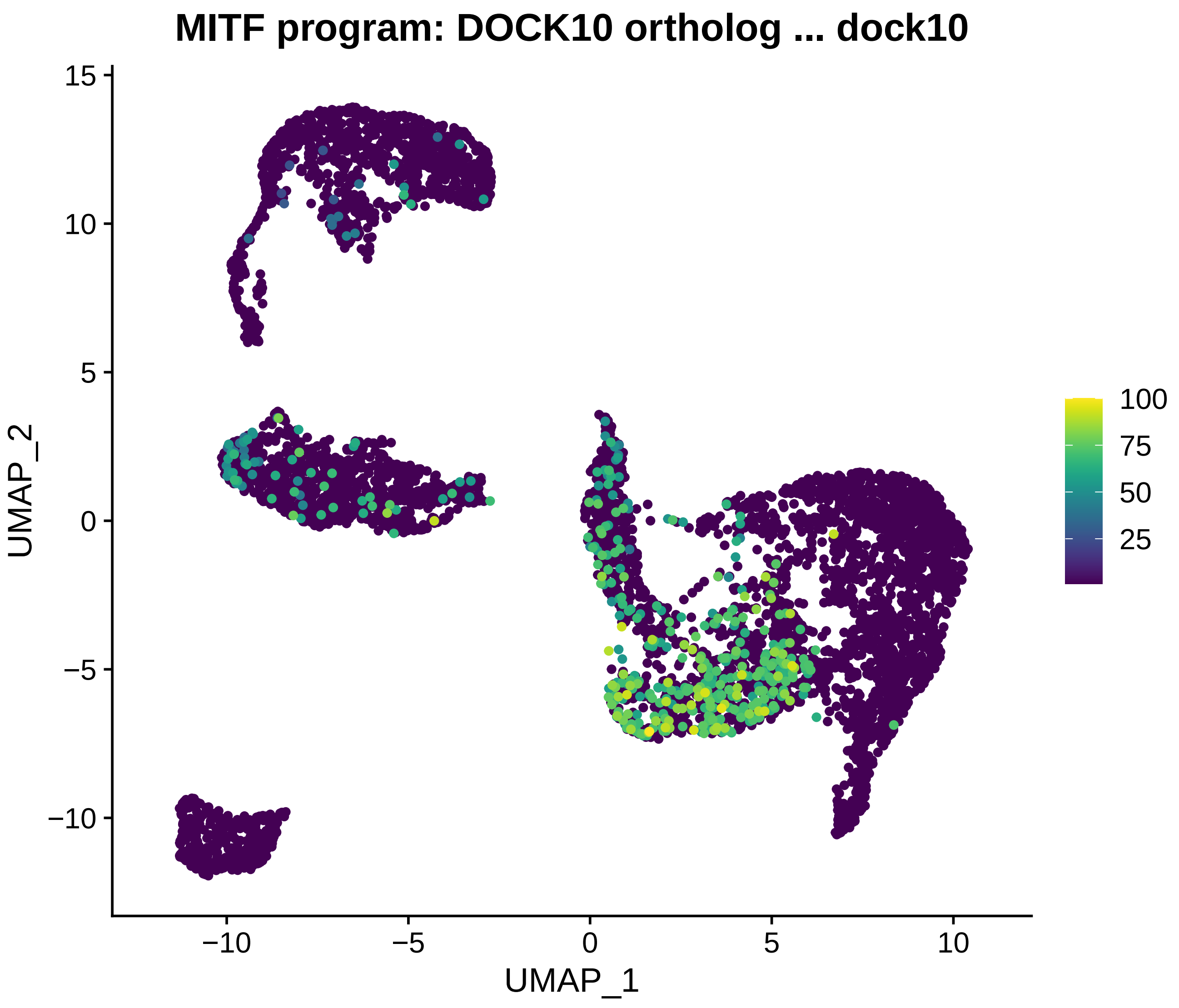
MITF program: DOCK10 ortholog ... dock10
UMAP_2
UMAP_1
15
10
5
0
−5
−10
−10
−5
0
5
10
100
75
50
25

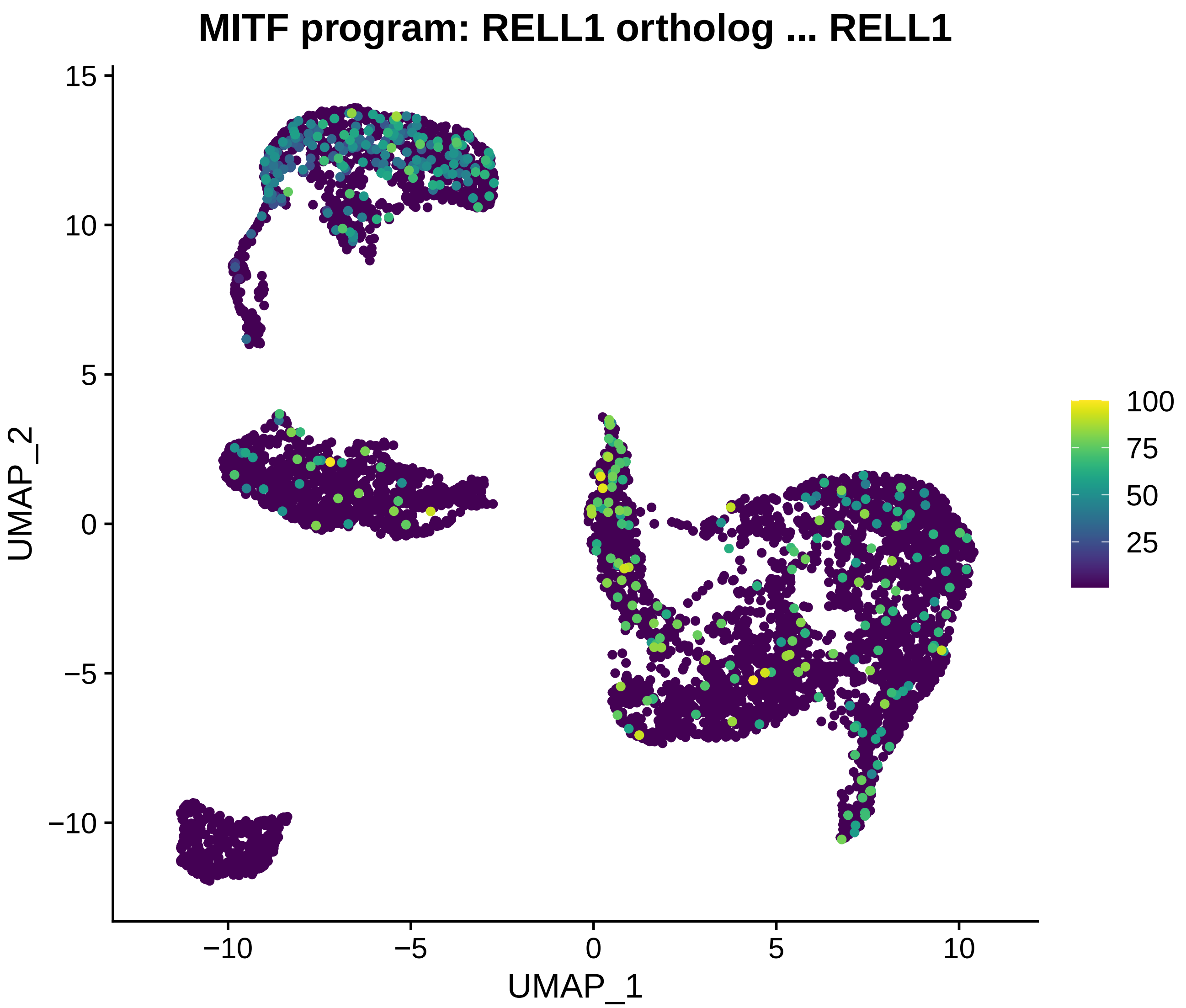
MITF program: RELL1 ortholog ... RELL1
UMAP_2
UMAP_1
15
10
5
0
−5
−10
−10
−5
0
5
10
100
75
50
25

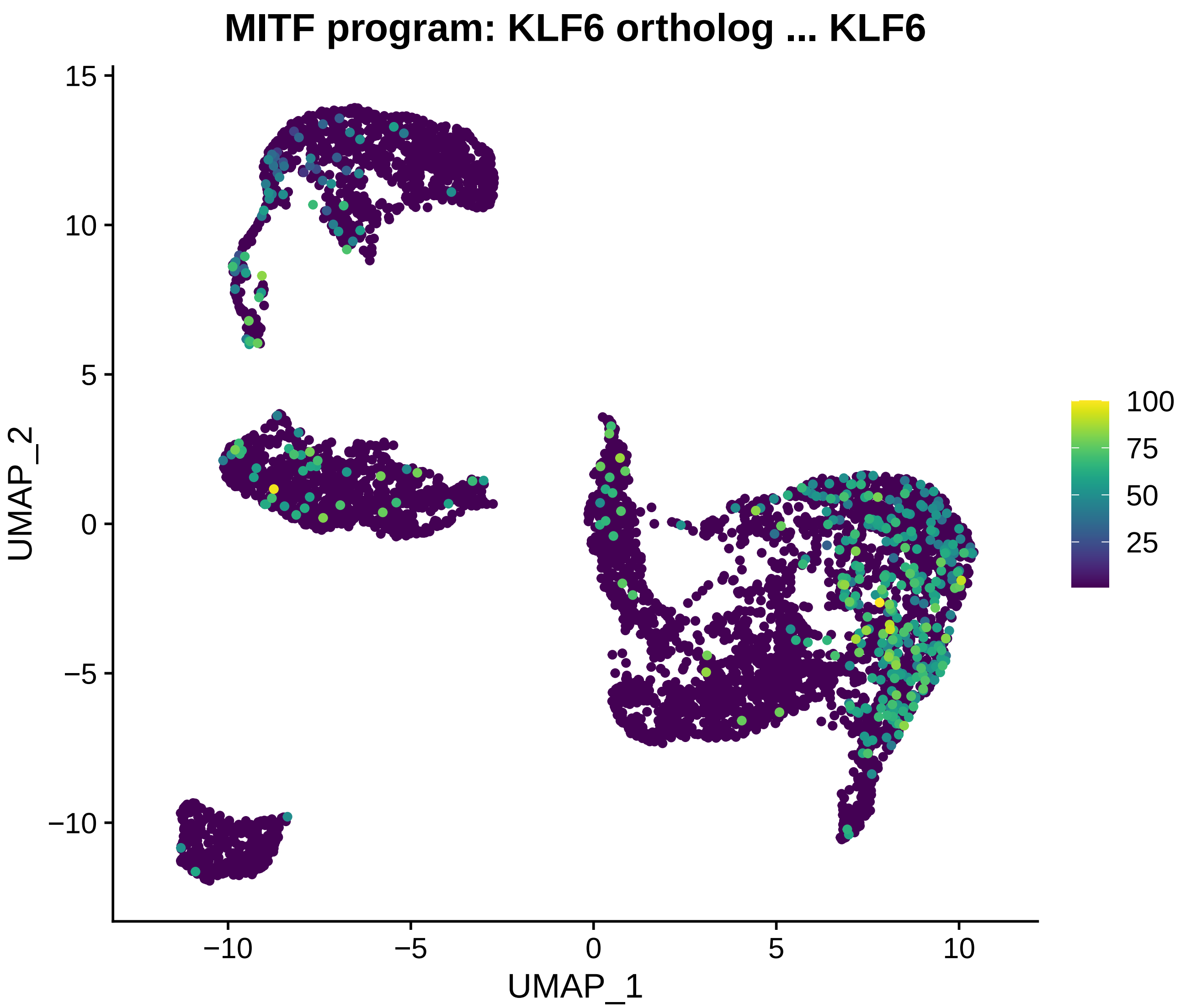
MITF program: KLF6 ortholog ... KLF6
15
10
5
0
−5
−10
UMAP_2
−10
−5
0
5
10
UMAP_1
100
75
50
25

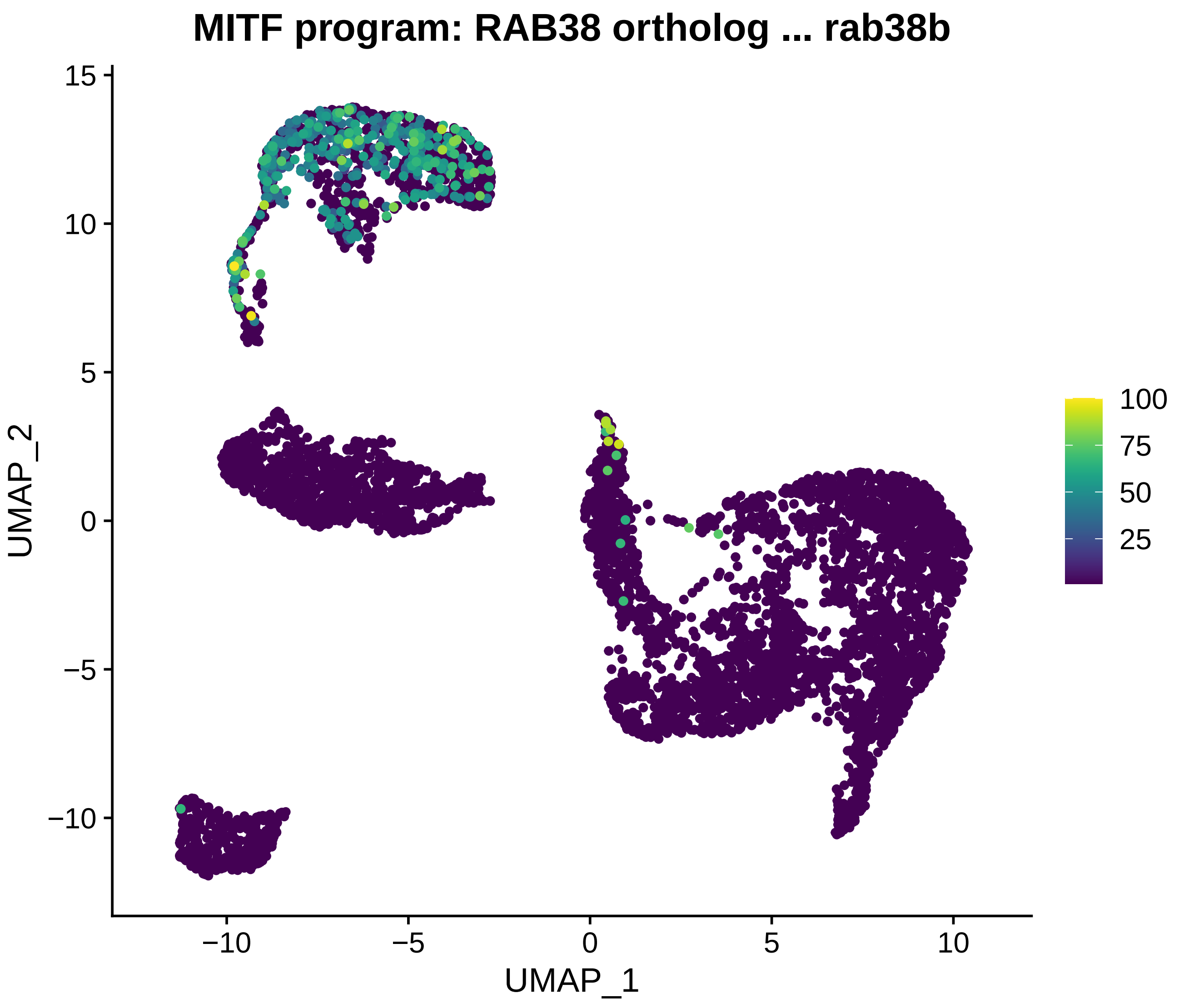
MITF program: RAB38 ortholog ... rab38b
UMAP_1
UMAP_2
−10
−5
0
5
10
15
100
75
50
25

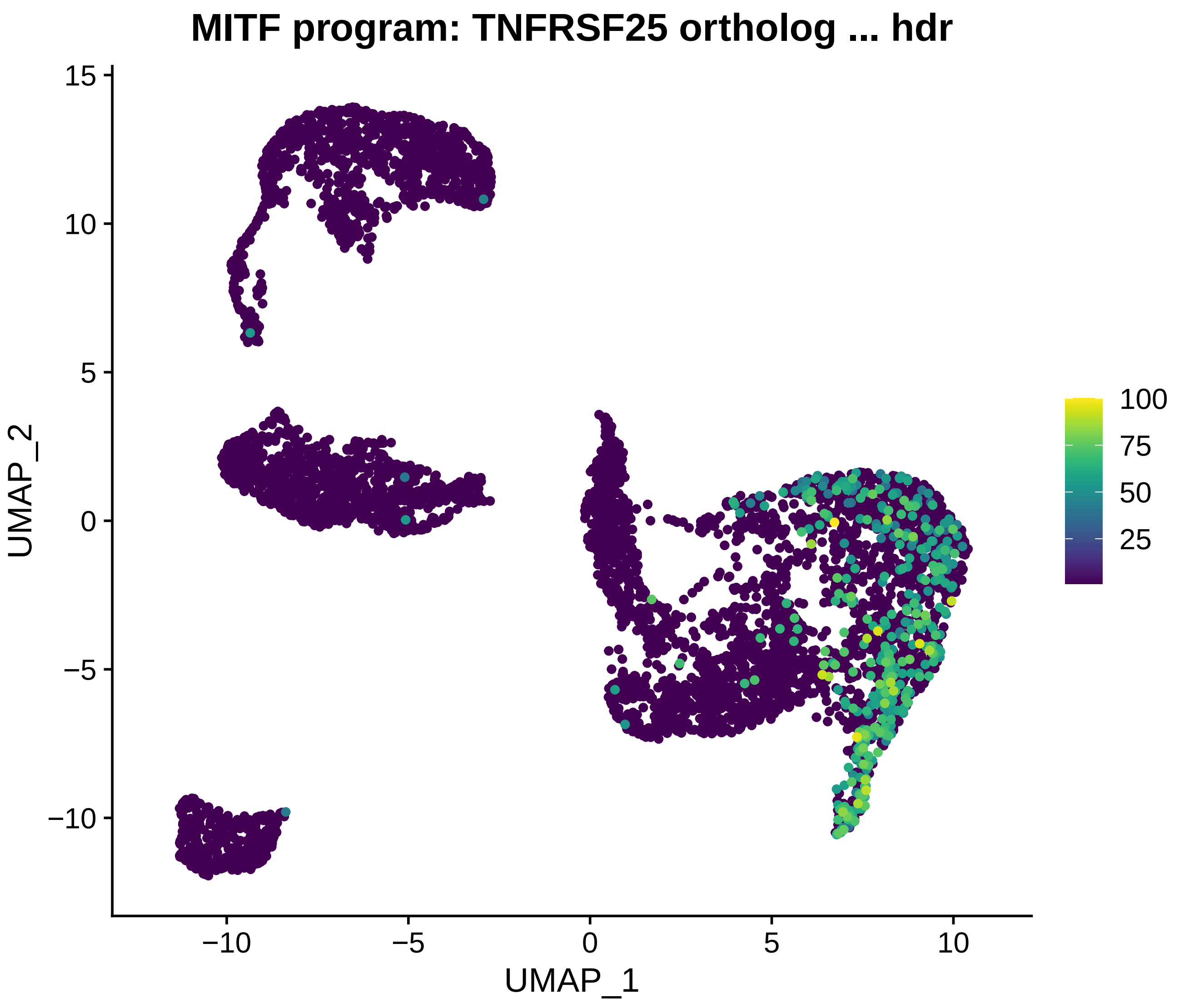

MITF program: TNFRSF25 ortholog ... hdr
UMAP_2
UMAP_1
15
10
5
0
−5
−10
−10
−5
0
5
10
100
75
50
25

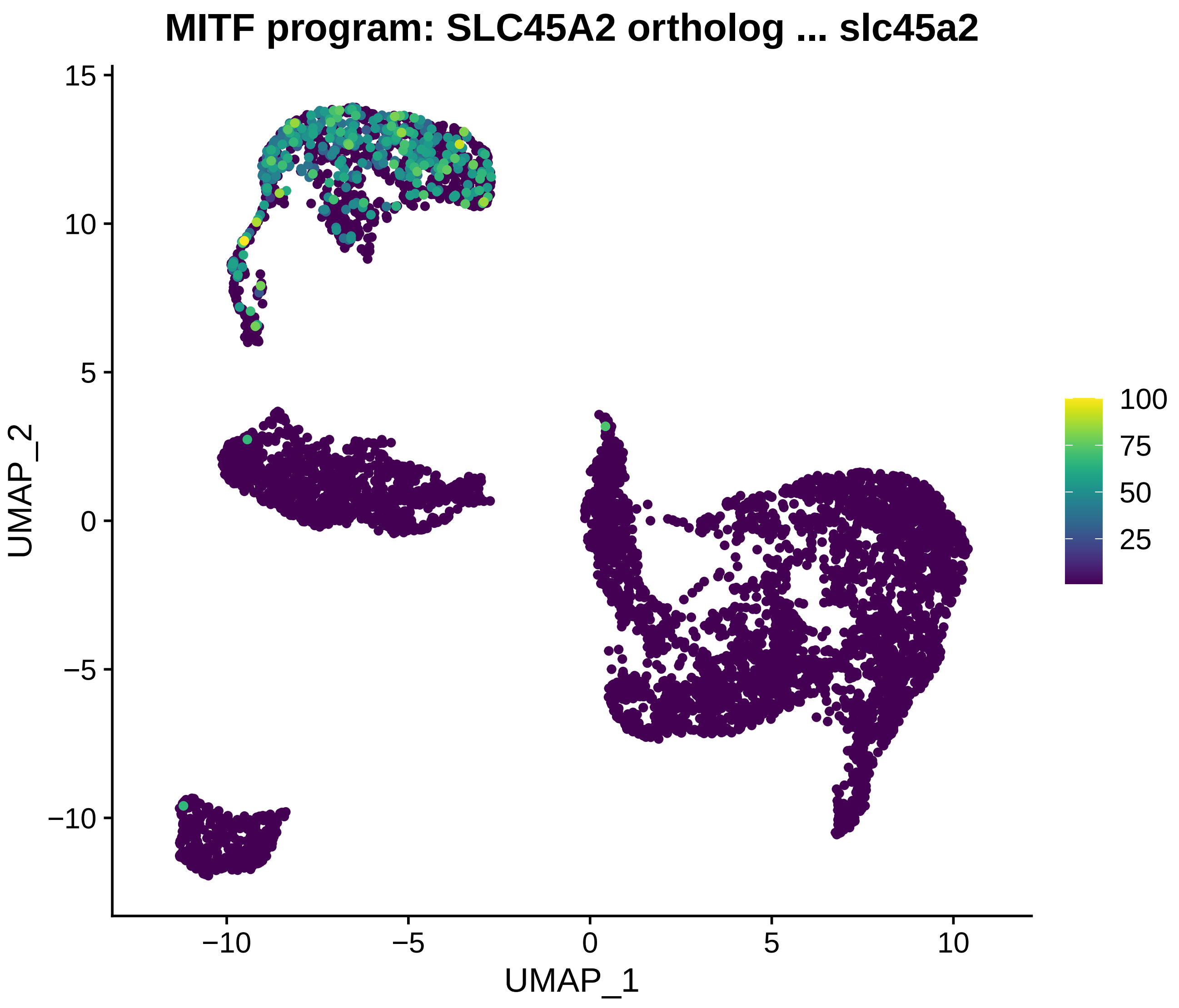
MITF program: SLC45A2 ortholog ... slc45a2
UMAP_2
UMAP_1
15
10
5
0
−5
−10
−10
−5
0
5
10
100
75
50
25

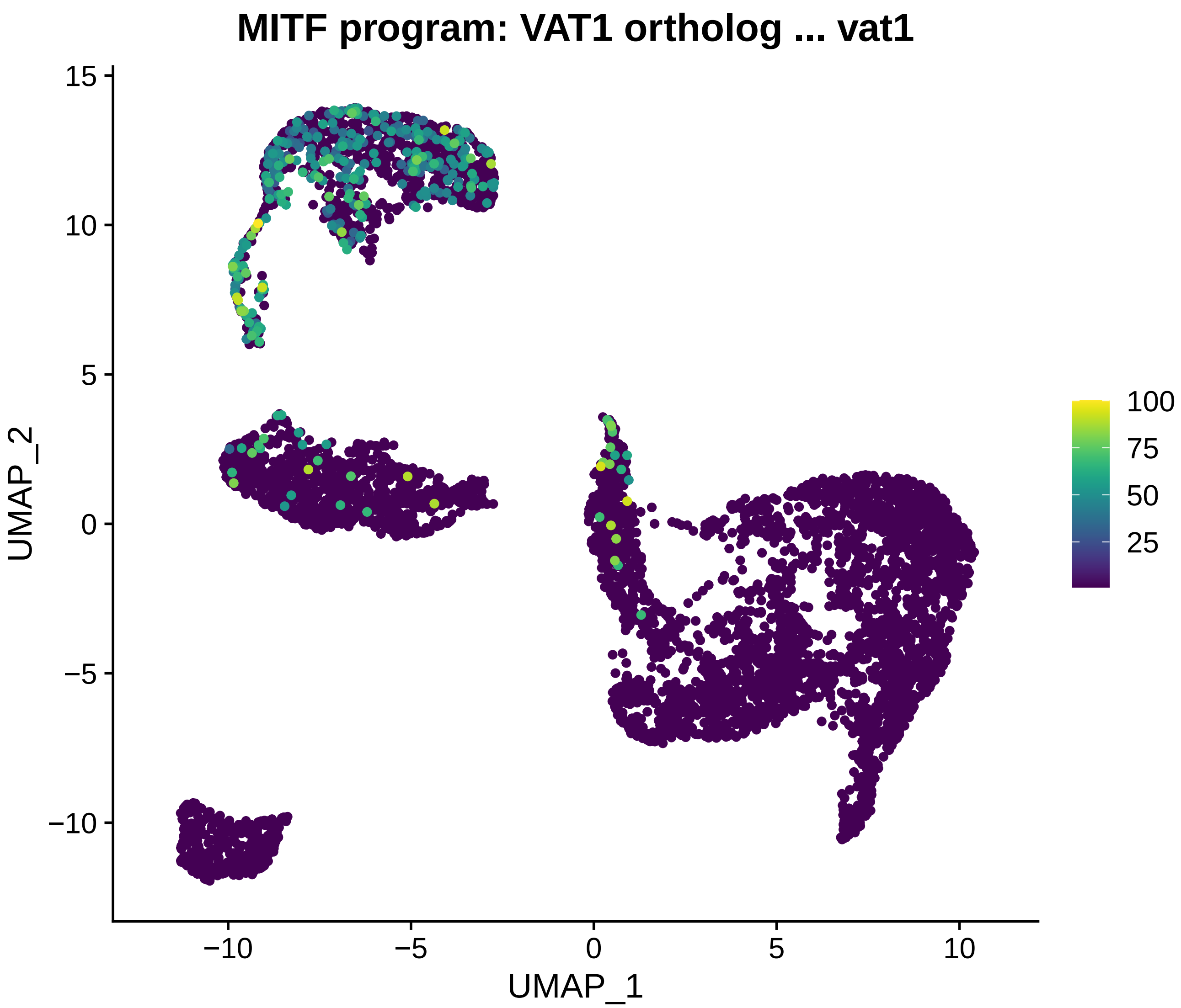
MITF program: VAT1 ortholog ... vat1
UMAP_2
UMAP_1
15
10
5
0
−5
−10
−10
−5
0
5
10
100
75
50
25

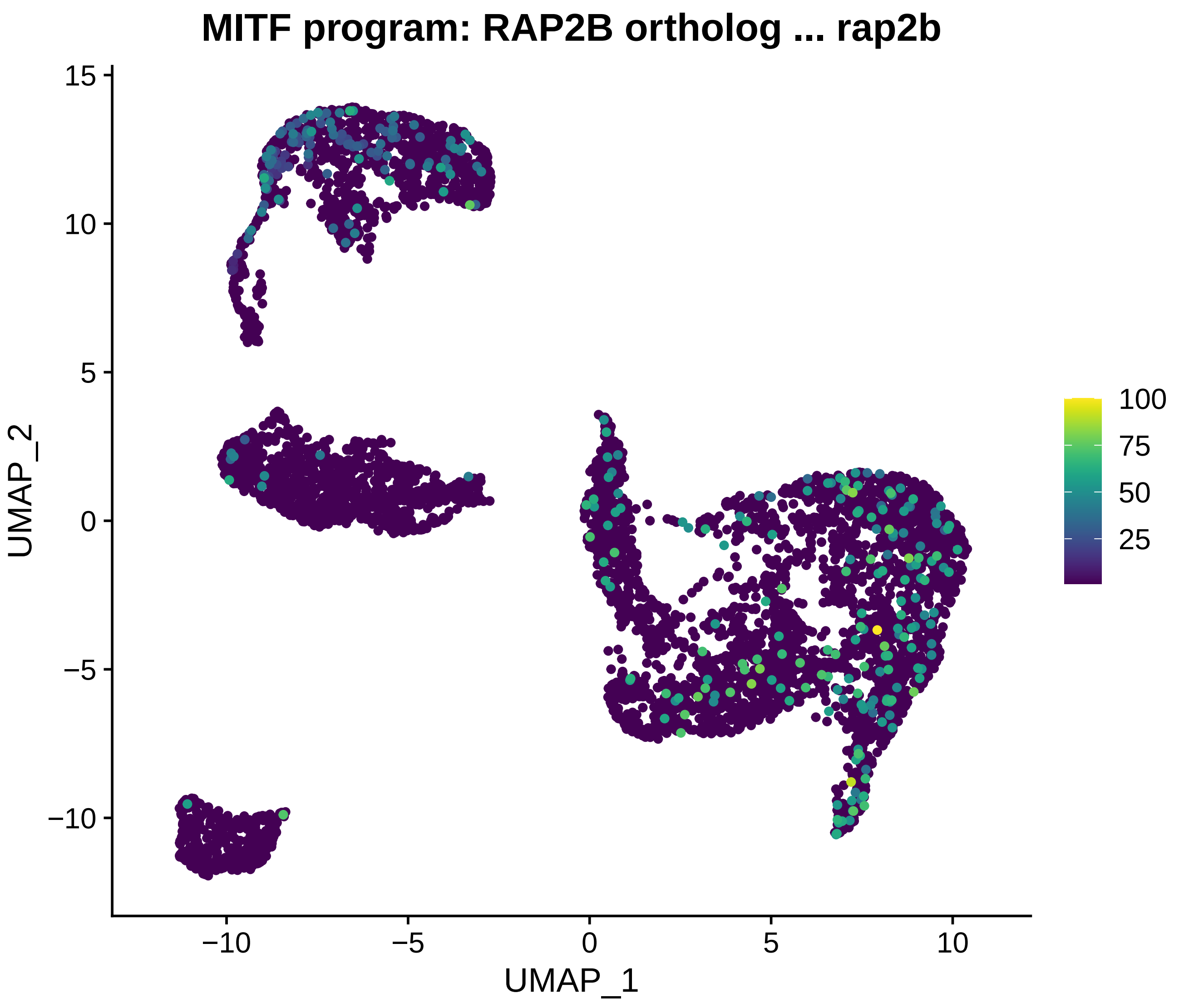

MITF program: RAP2B ortholog ... rap2b
UMAP_2
UMAP_1
15
10
5
0
−5
−10
−10
−5
0
5
10
100
75
50
25

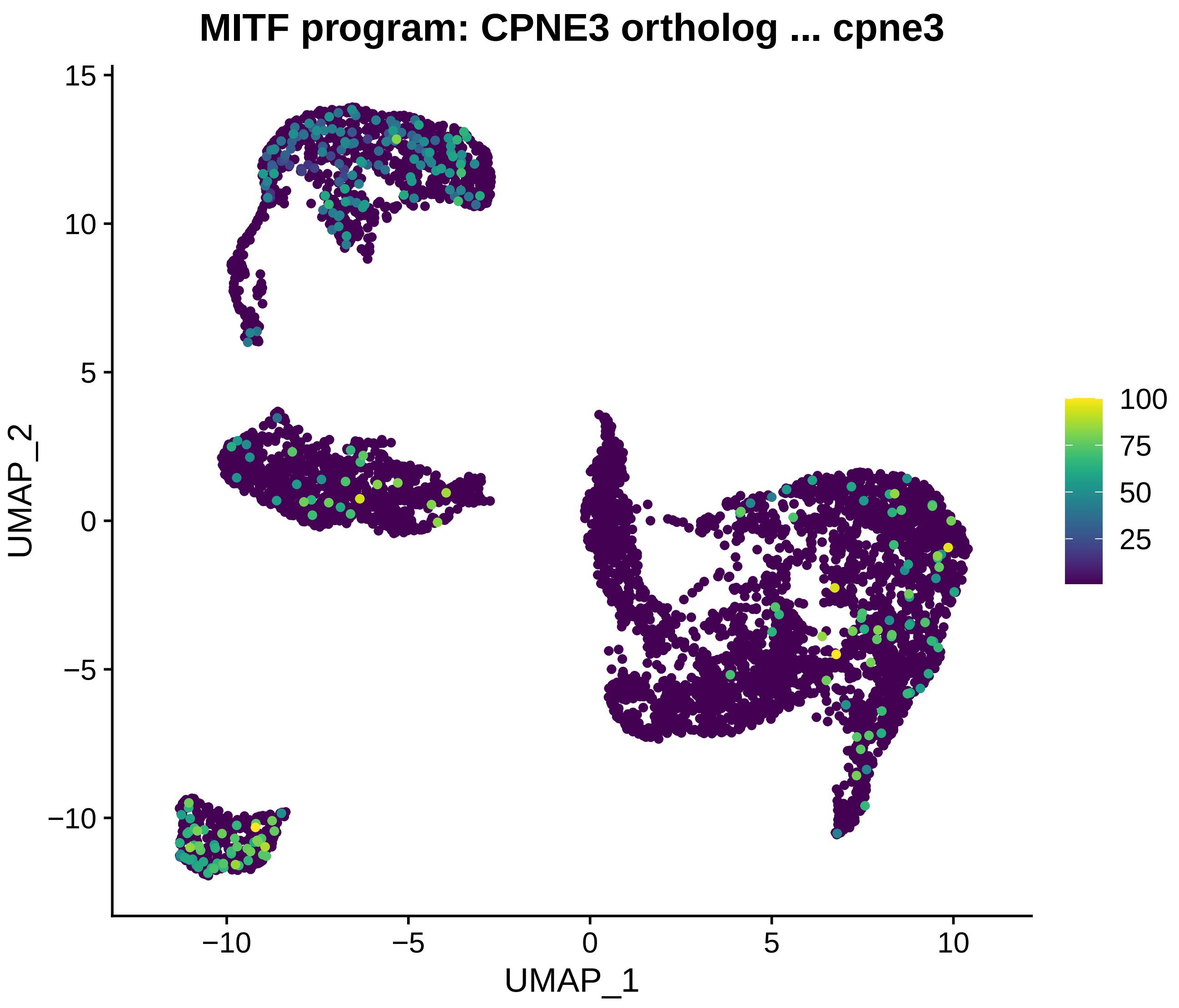

MITF program: CPNE3 ortholog ... cpne3
UMAP_2
UMAP_1
15
10
5
0
−5
−10
−10
−5
0
5
10
100
75
50
25

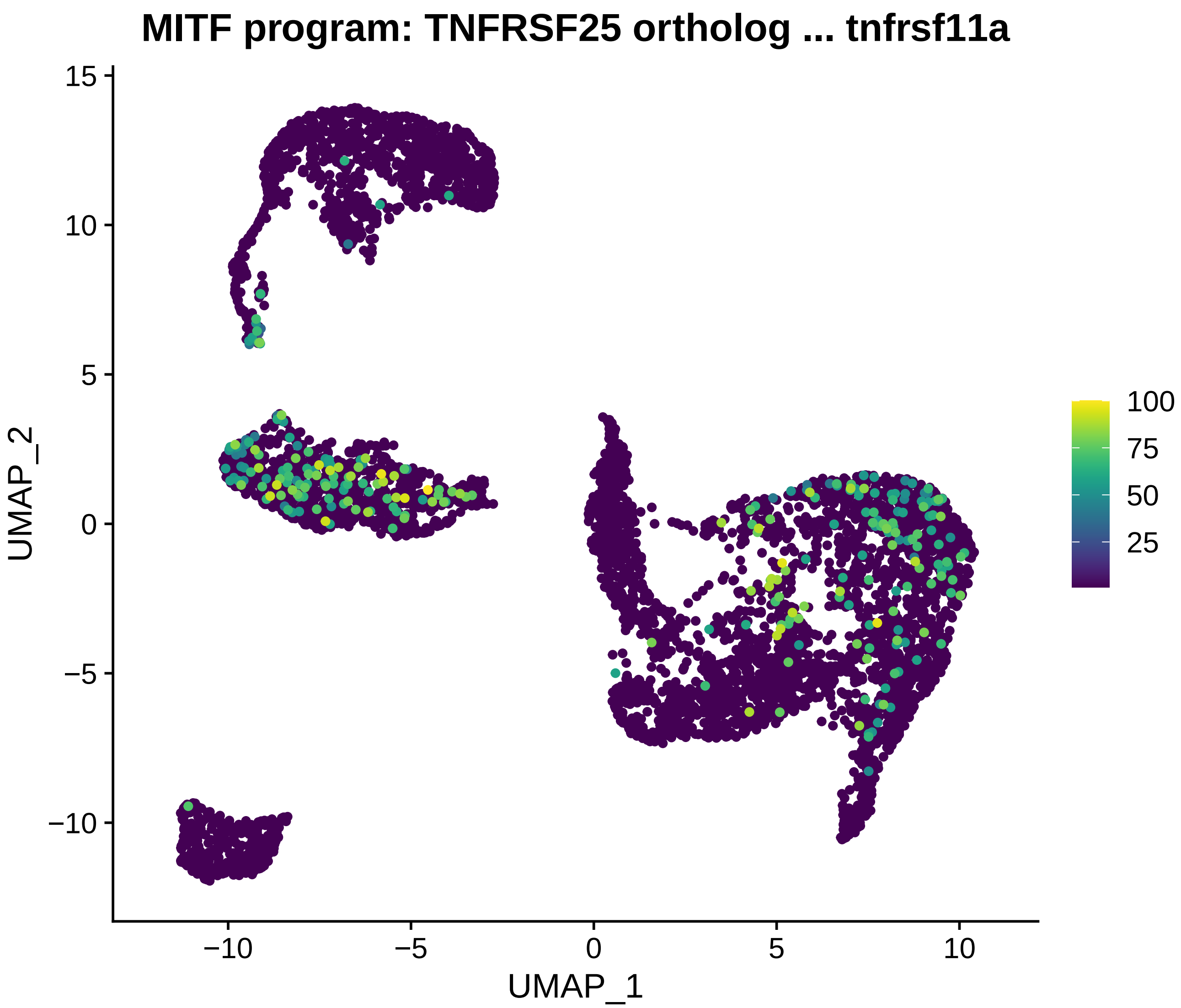
MITF program: TNFRSF25 ortholog ... tnfrsf11a
UMAP_2
UMAP_1
15
10
5
0
−5
−10
−10
−5
0
5
10
100
75
50
25

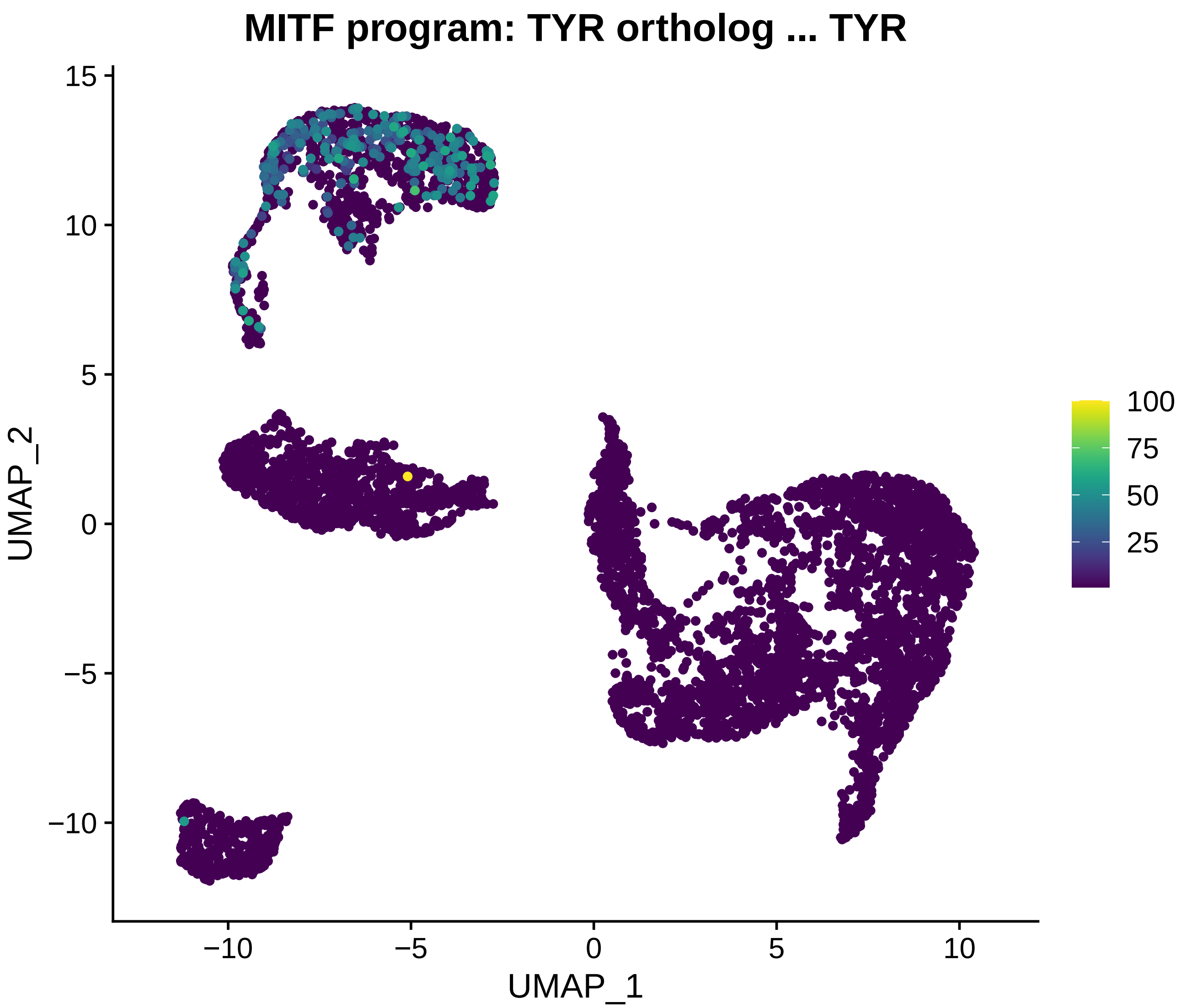

MITF program: TYR ortholog ... TYR
UMAP_2
UMAP_1
15
10
5
0
−5
−10
−10
−5
0
5
10
100
75
50
25

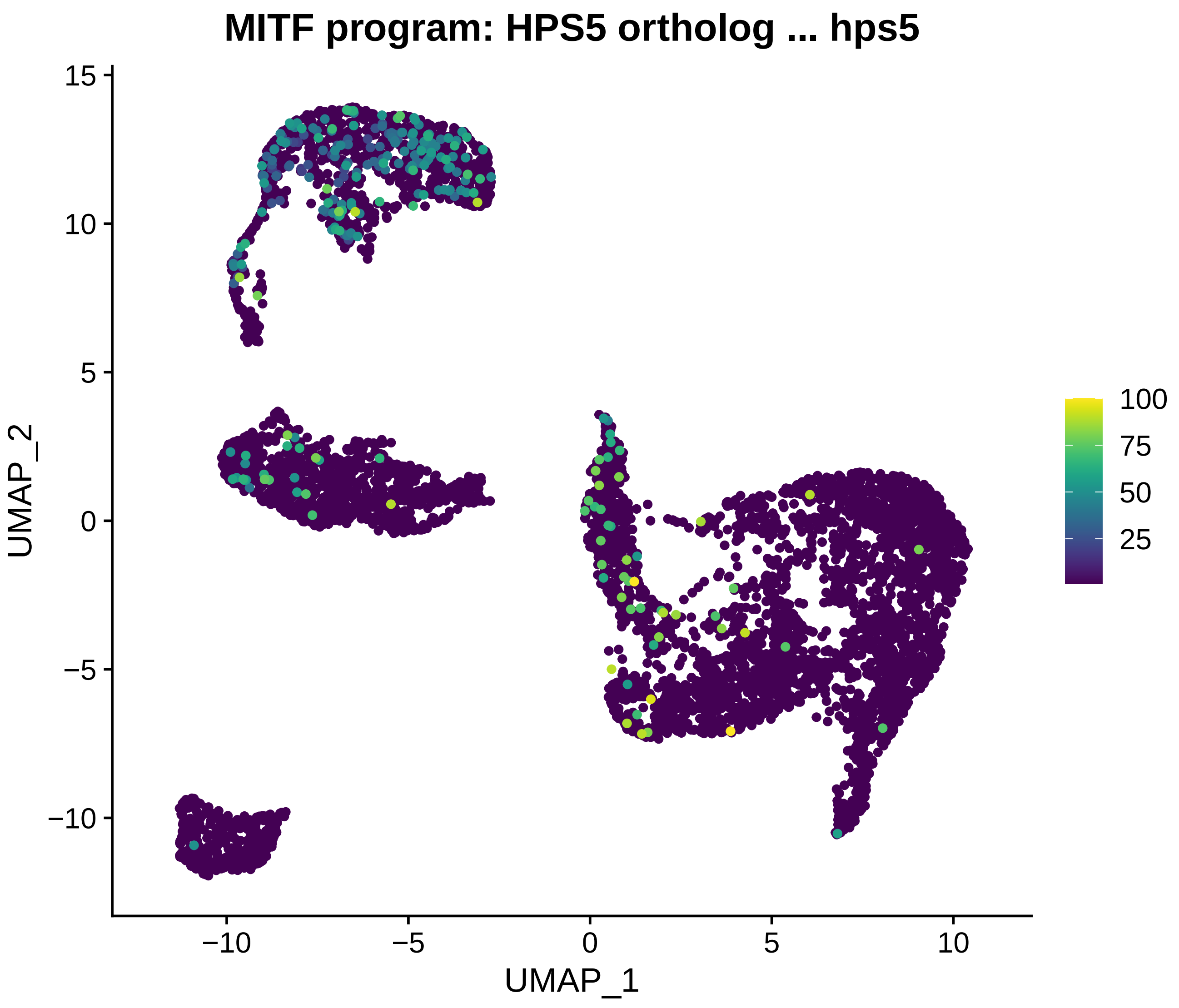
MITF program: HPS5 ortholog ... hps5
UMAP_2
UMAP_1
15
10
5
0
−5
−10
−10
−5
0
5
10
100
75
50
25

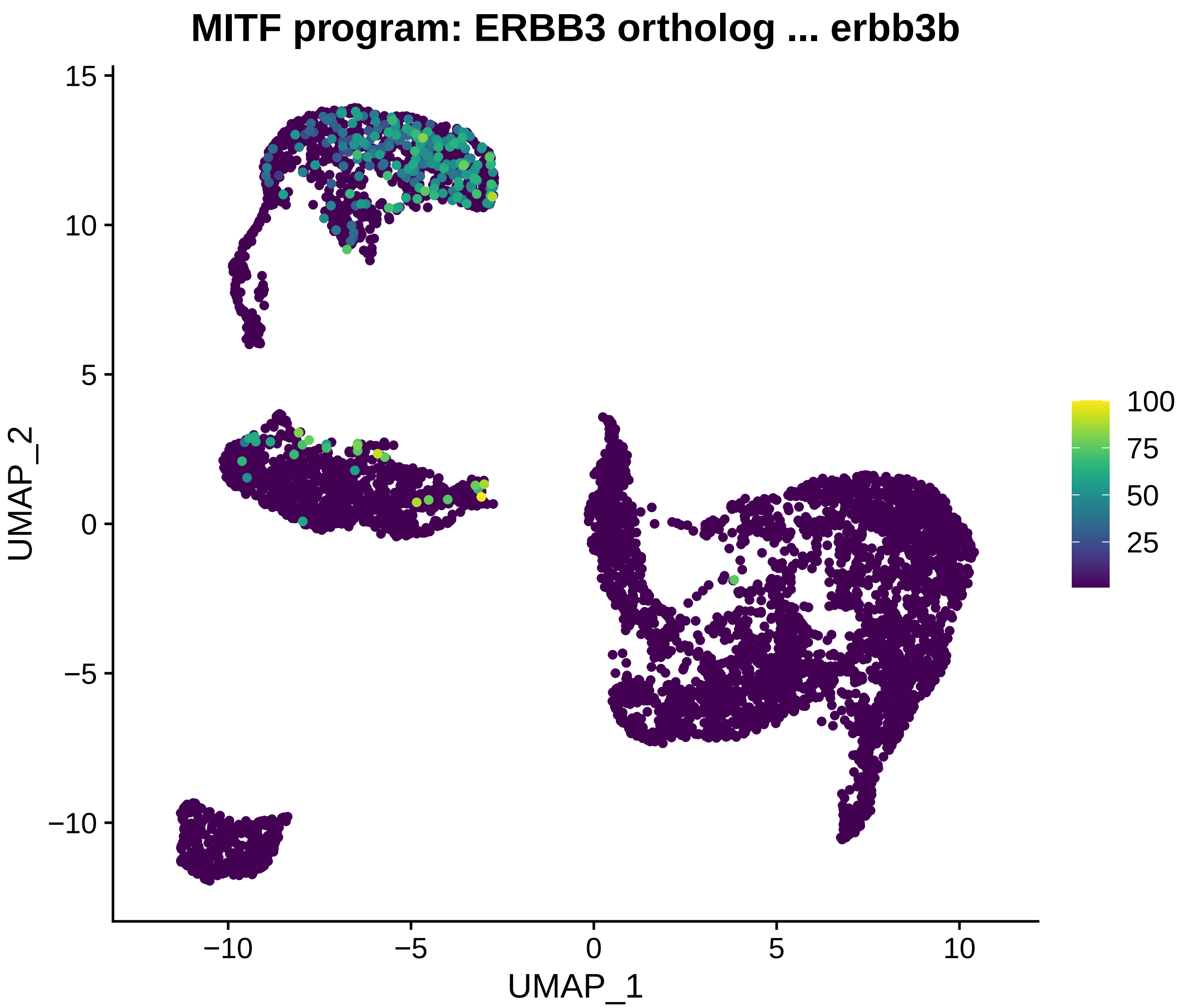

MITF program: ERBB3 ortholog ... erbb3b
UMAP_2
UMAP_1
15
10
5
0
−5
−10
−10
−5
0
5
10
100
75
50
25

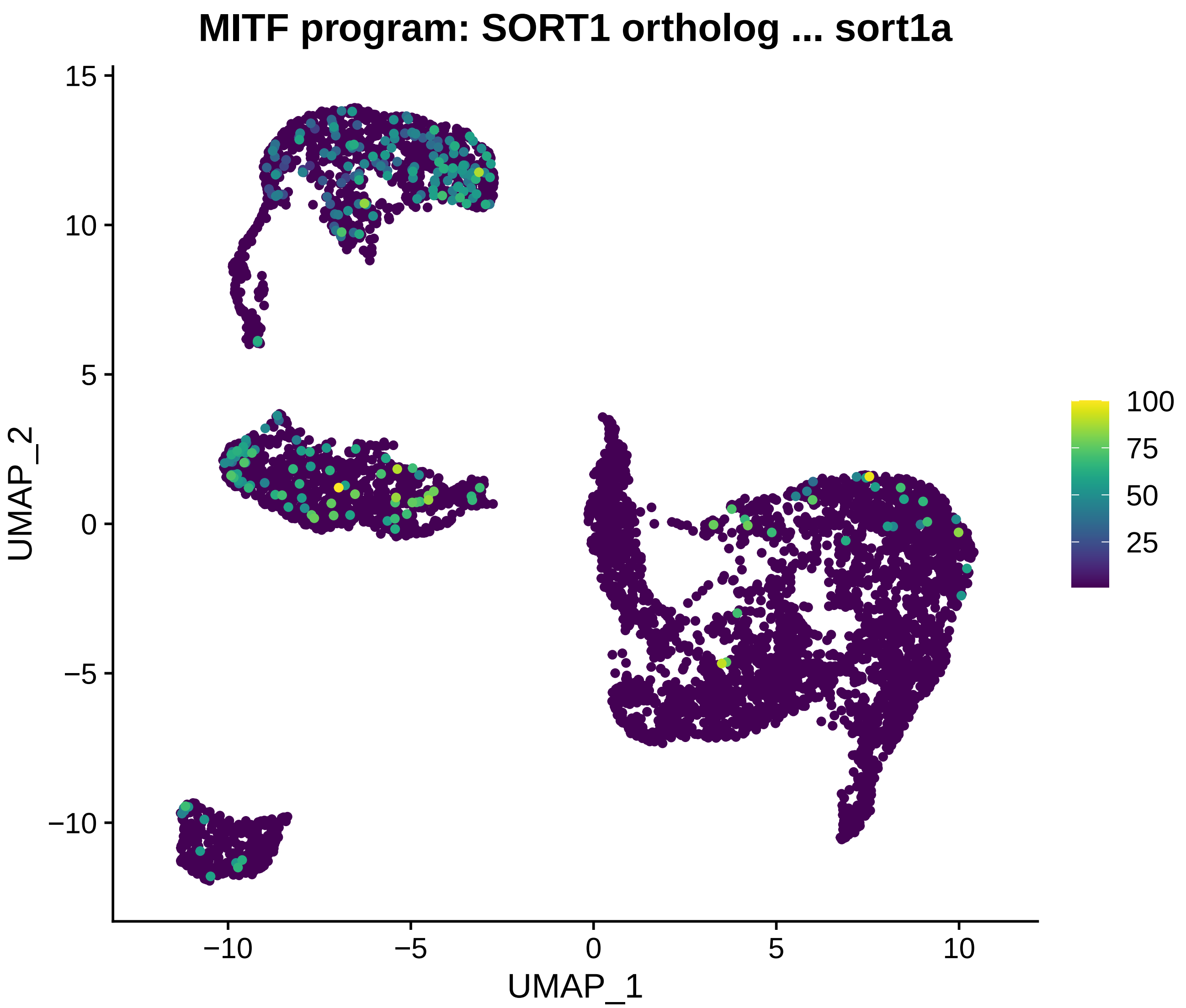
MITF program: SORT1 ortholog ... sort1a
15
10
5
0
−5
−10
UMAP_2
−10
−5
0
5
10
UMAP_1
100
75
50
25

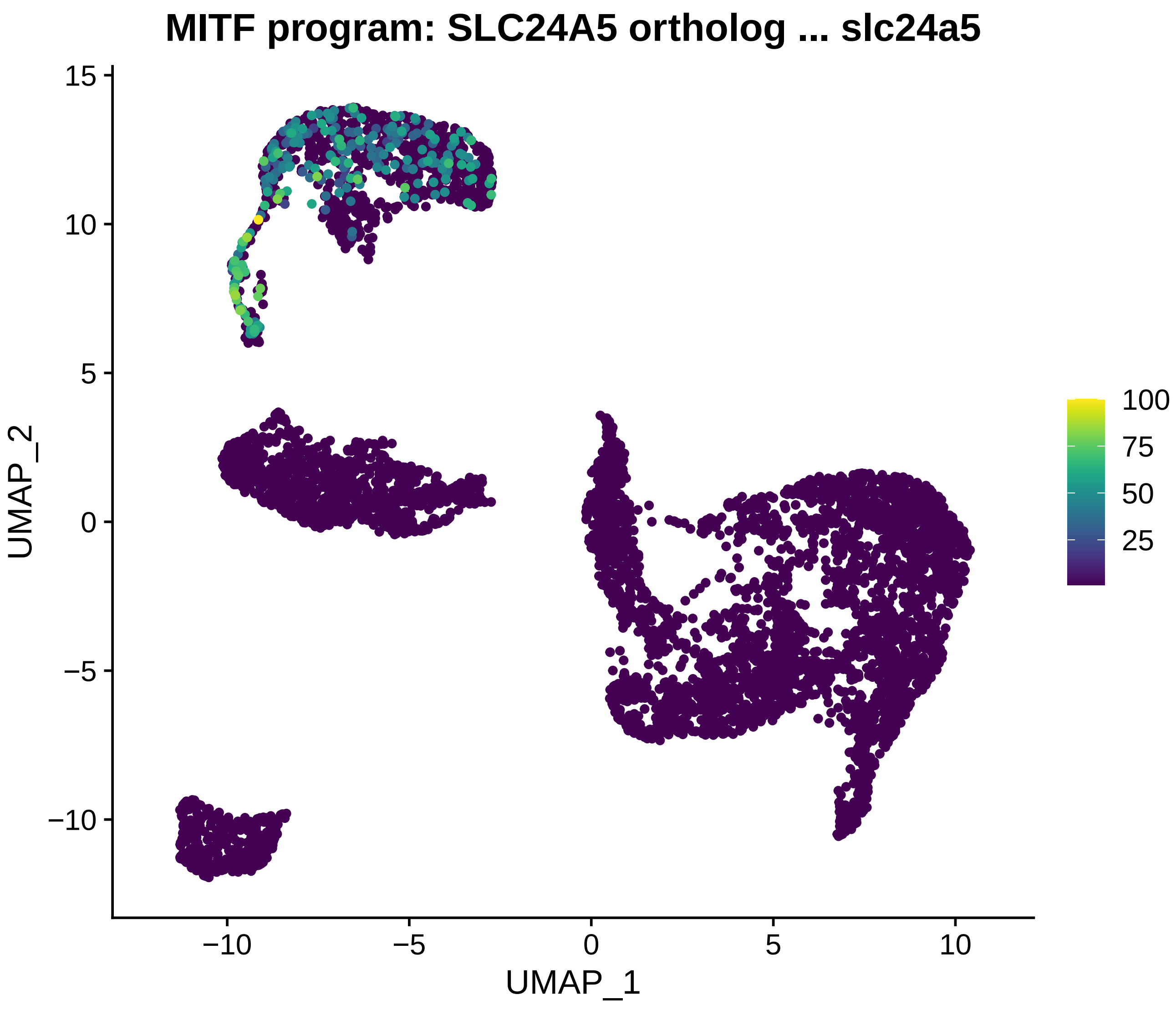
MITF program: SLC24A5 ortholog ... slc24a5
UMAP_2
UMAP_1
15
10
5
0
−5
−10
−10
−5
0
5
10
100
75
50
25

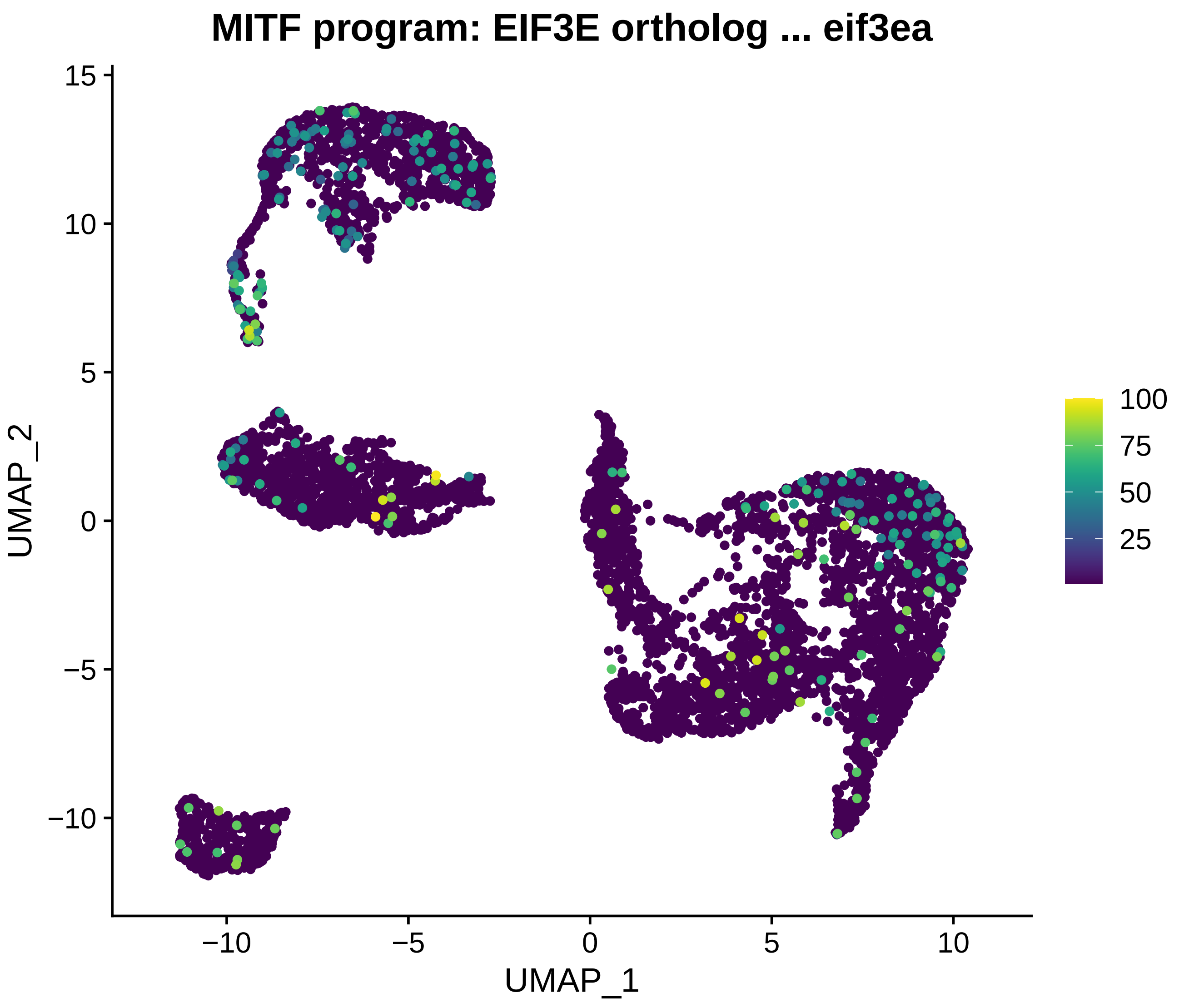
MITF program: EIF3E ortholog ... eif3ea
UMAP_2
UMAP_1
15
10
5
0
−5
−10
−10
−5
0
5
10
100
75
50
25

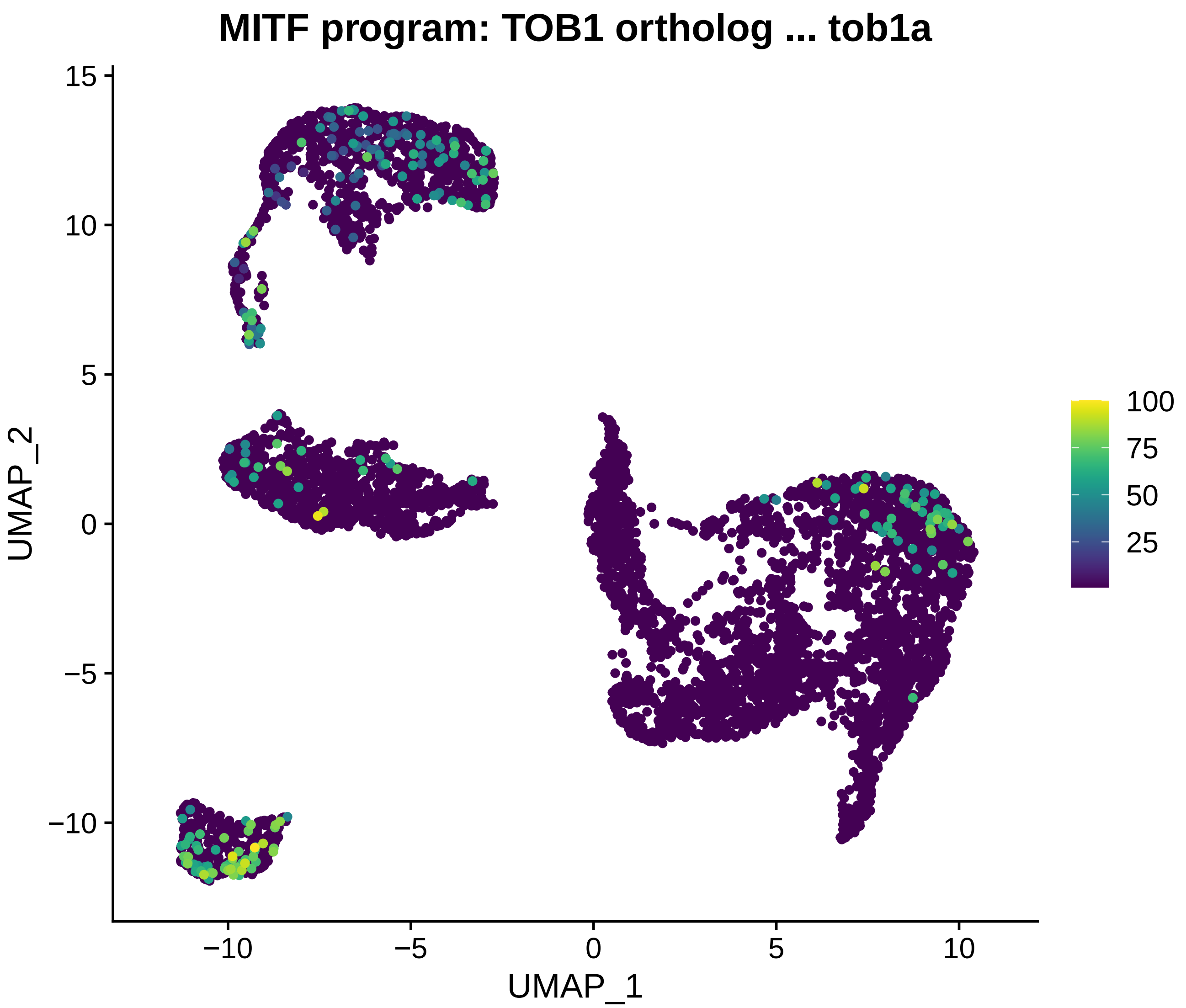
MITF program: TOB1 ortholog ... tob1a
UMAP_2
UMAP_1
15
10
5
0
−5
−10
−10
−5
0
5
10
100
75
50
25

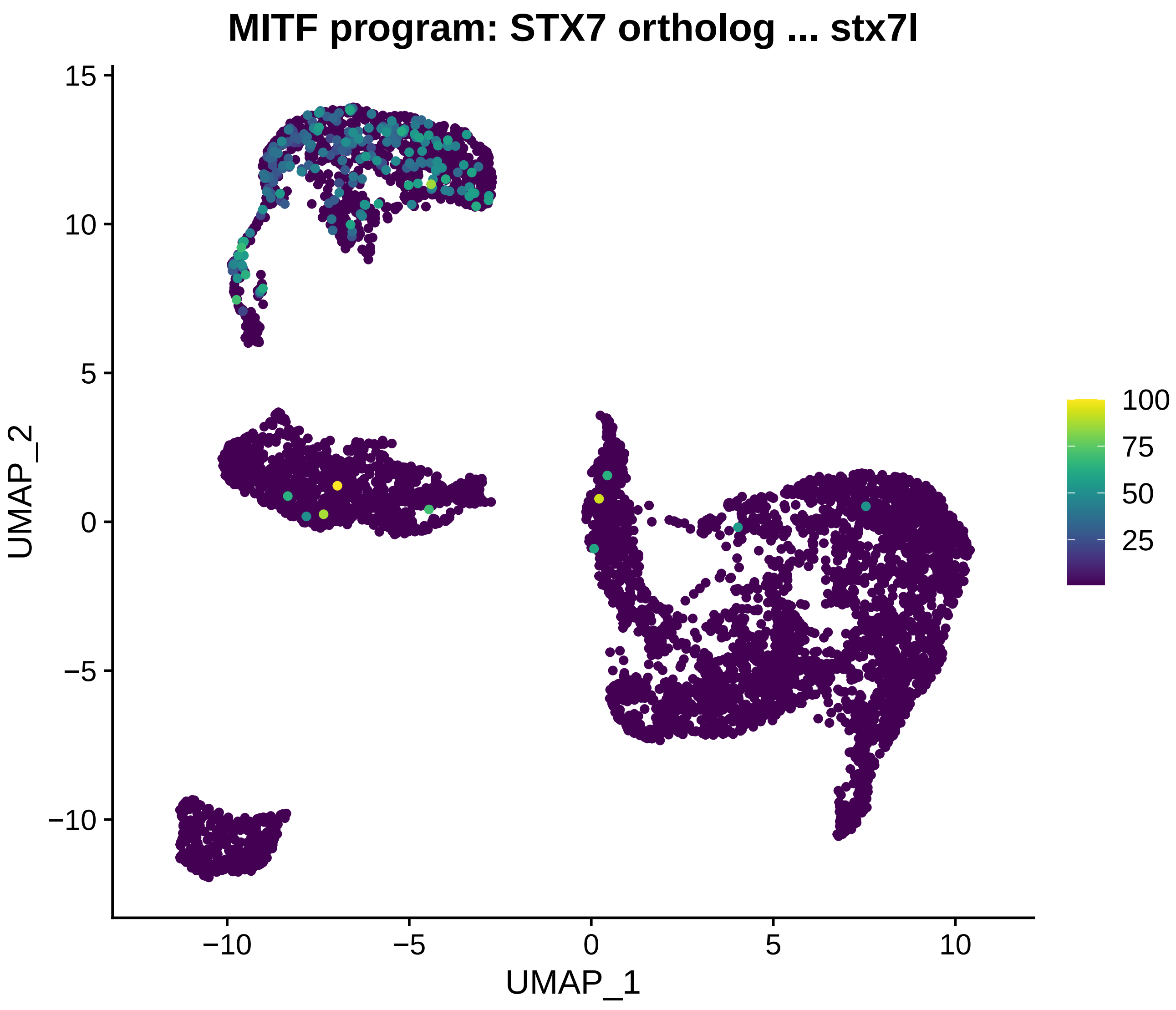

MITF program: STX7 ortholog ... stx7l
UMAP_2
UMAP_1
15
10
5
0
−5
−10
−10
−5
0
5
10
100
75
50
25

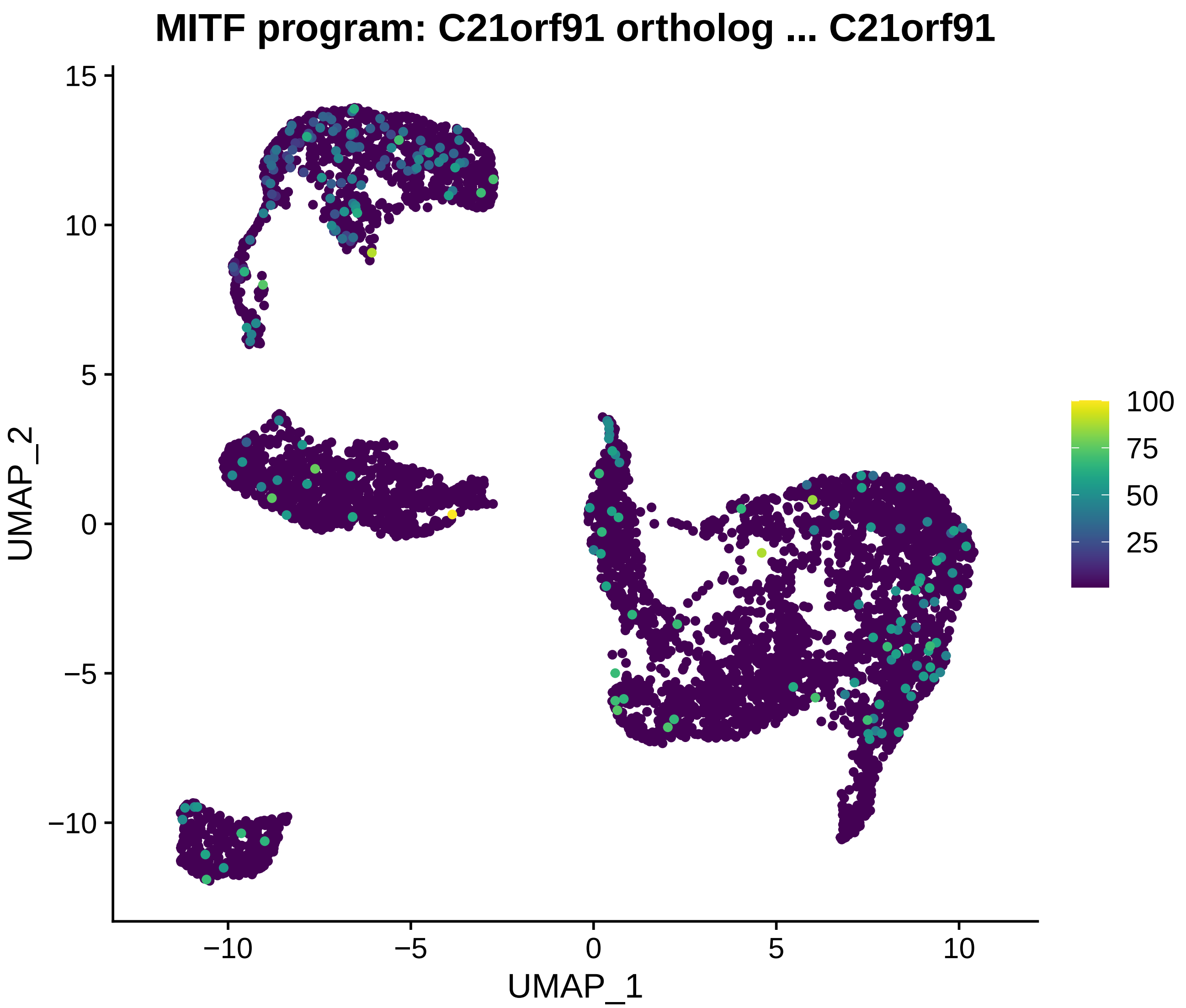

MITF program: C21orf91 ortholog ... C21orf91
UMAP_2
UMAP_1
15
10
5
0
−5
−10
−10
−5
0
5
10
100
75
50
25

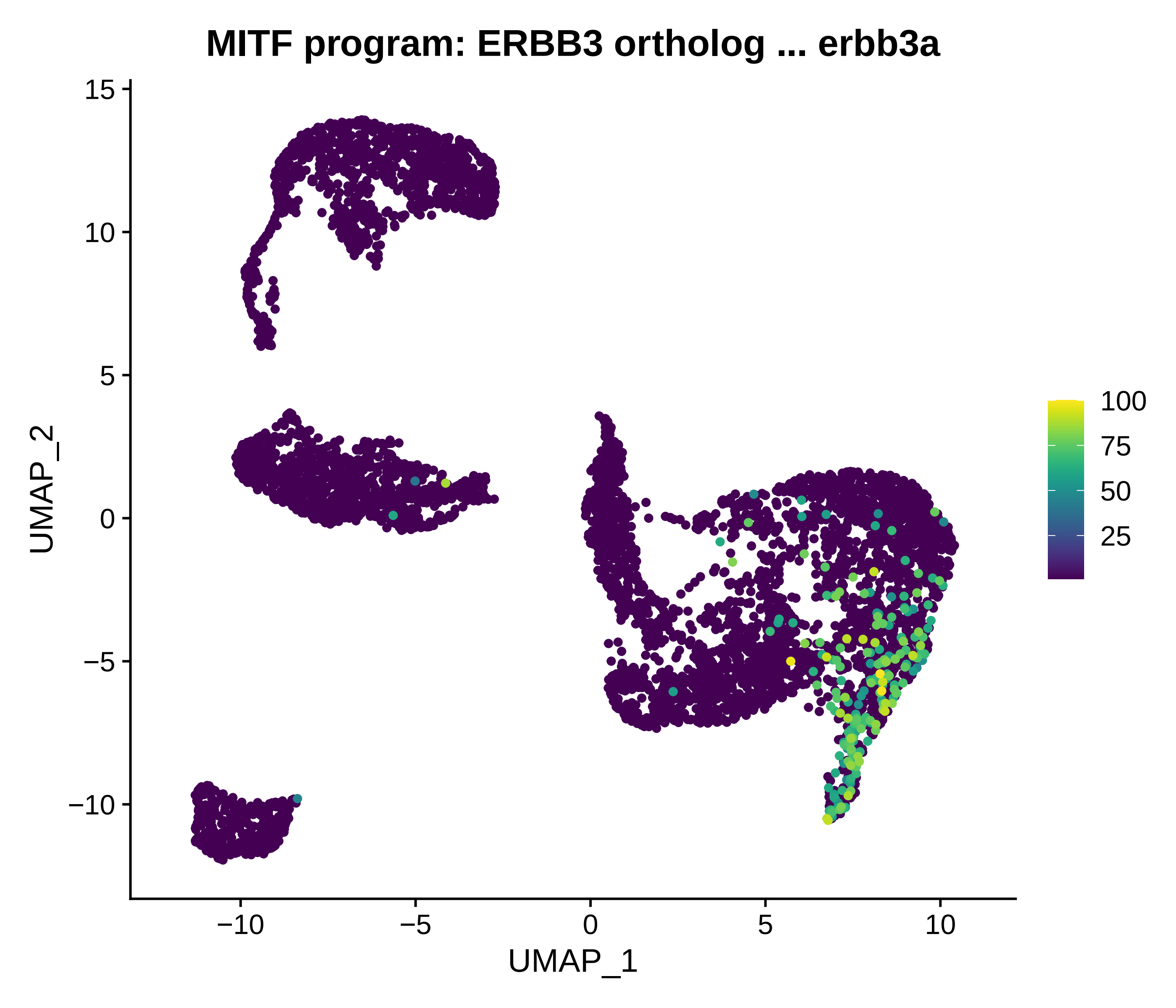
MITF program: ERBB3 ortholog ... erbb3a
UMAP_1
UMAP_2
−10
−5
0
5
10
15
100
75
50
25

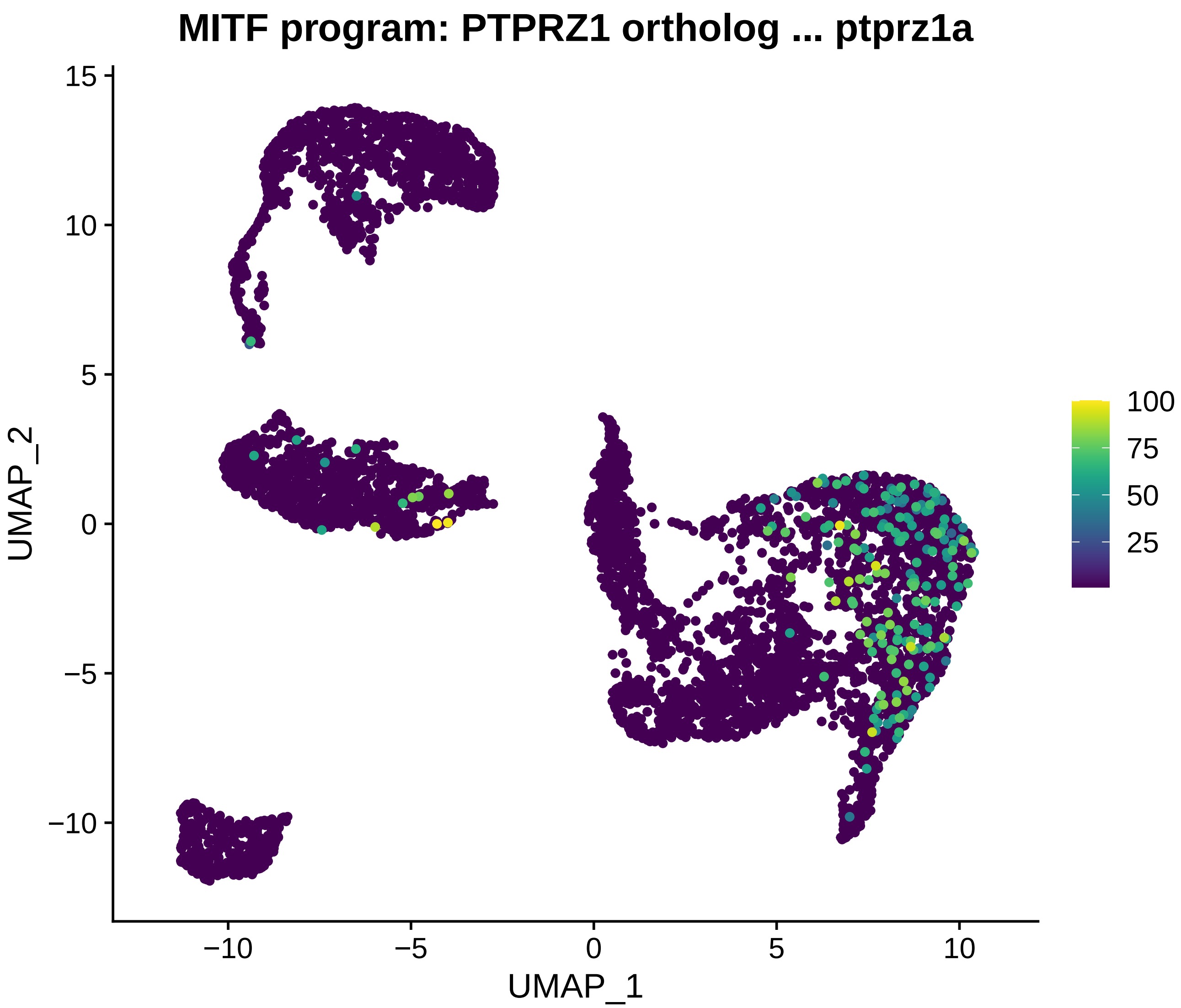
MITF program: PTPRZ1 ortholog ... ptprz1a
UMAP_2
UMAP_1
15
10
5
0
−5
−10
−10
−5
0
5
10
100
75
50
25

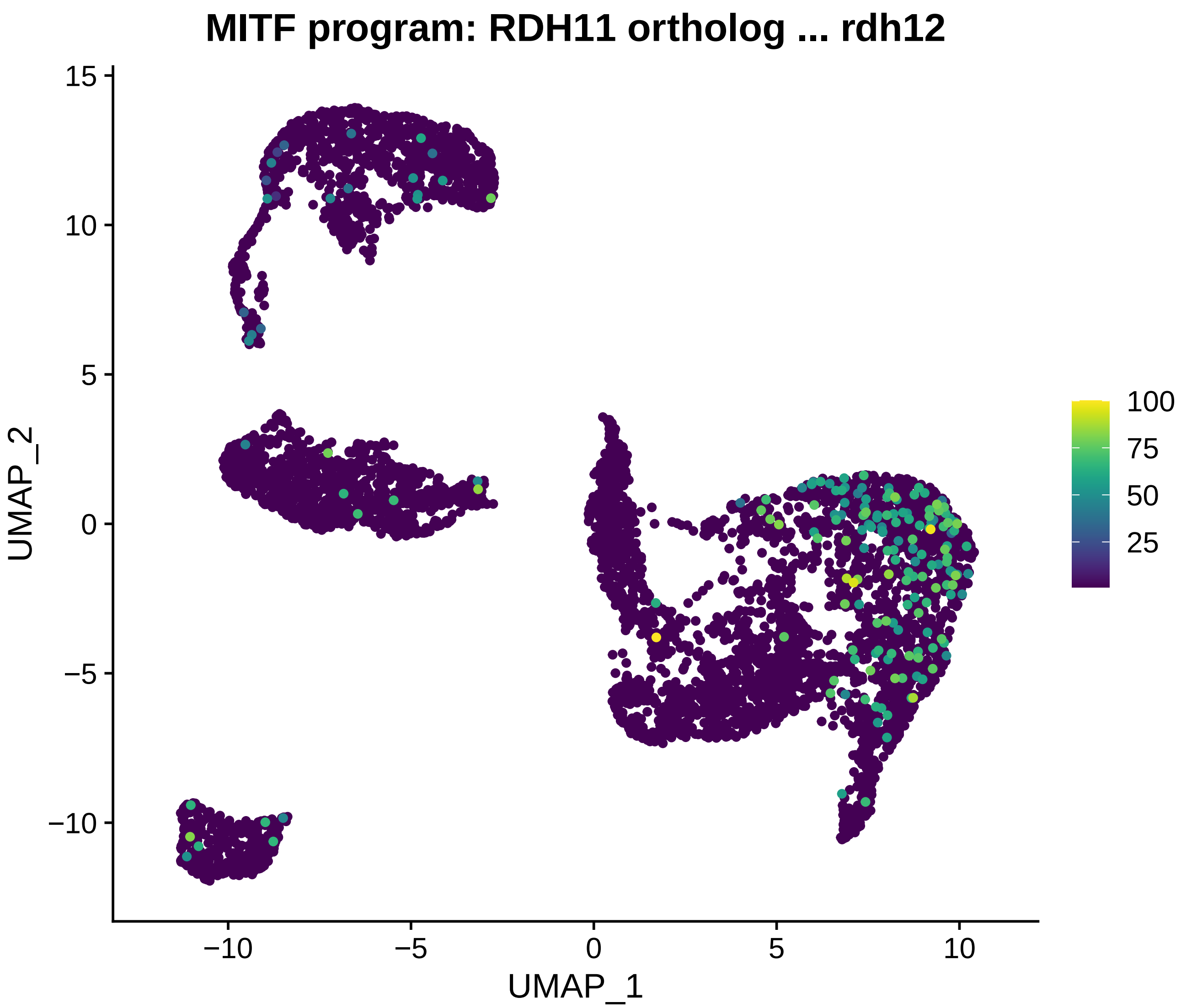
MITF program: RDH11 ortholog ... rdh12
UMAP_2
UMAP_1
15
10
5
0
−5
−10
−10
−5
0
5
10
100
75
50
25

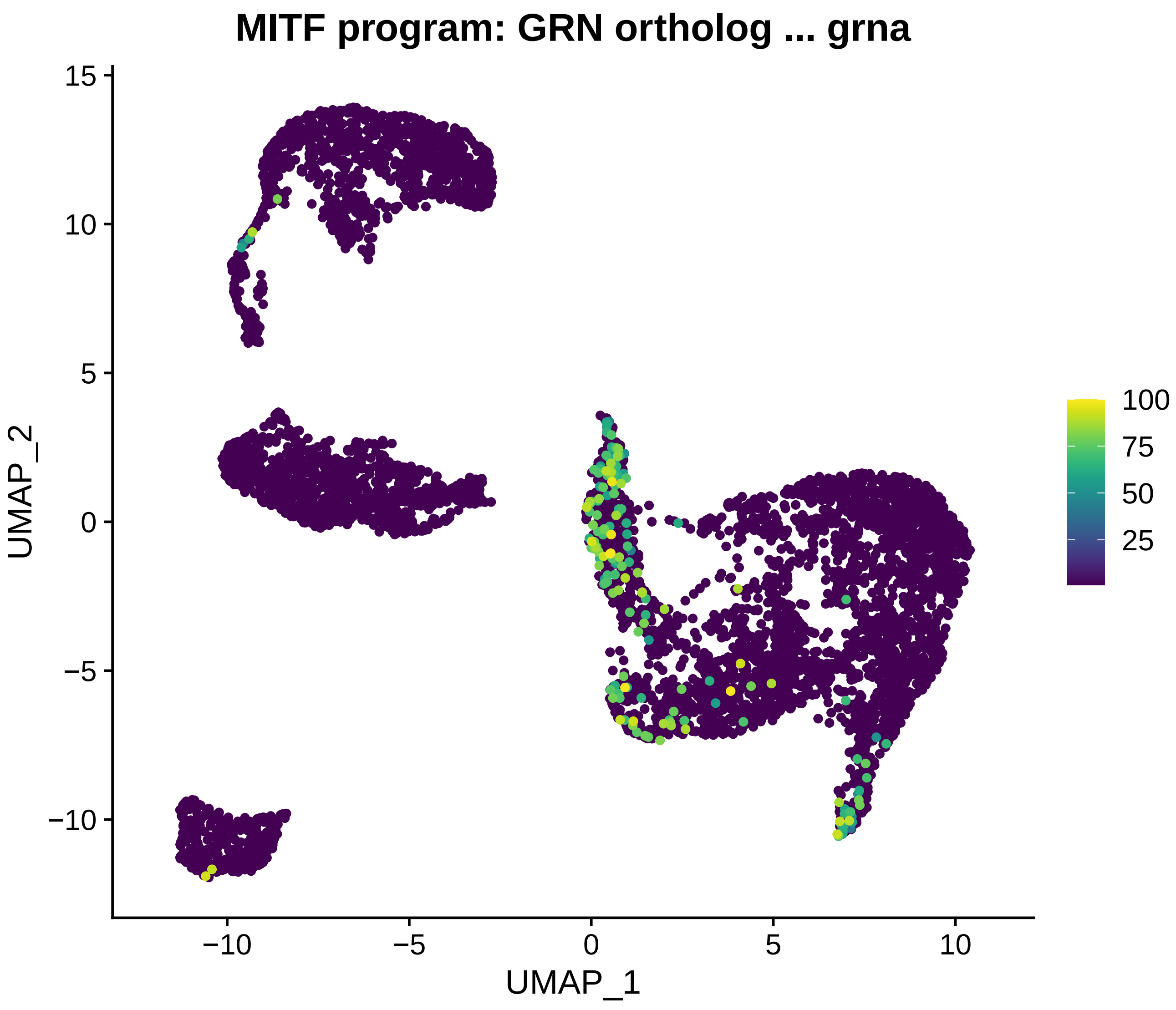

MITF program: GRN ortholog ... grna
UMAP_2
UMAP_1
15
10
5
0
−5
−10
−10
−5
0
5
10
100
75
50
25

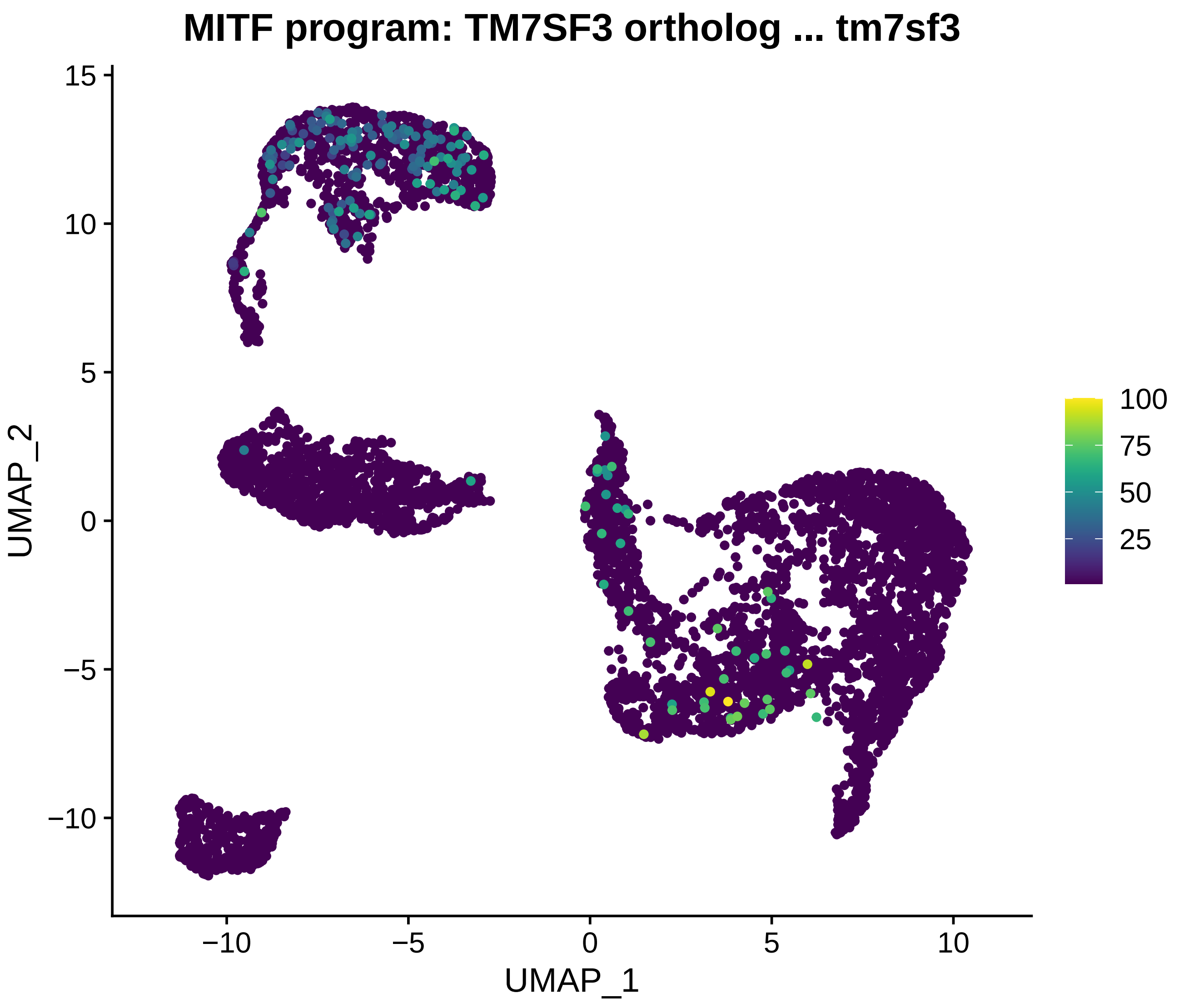

MITF program: TM7SF3 ortholog ... tm7sf3
UMAP_2
UMAP_1
15
10
5
0
−5
−10
−10
−5
0
5
10
100
75
50
25

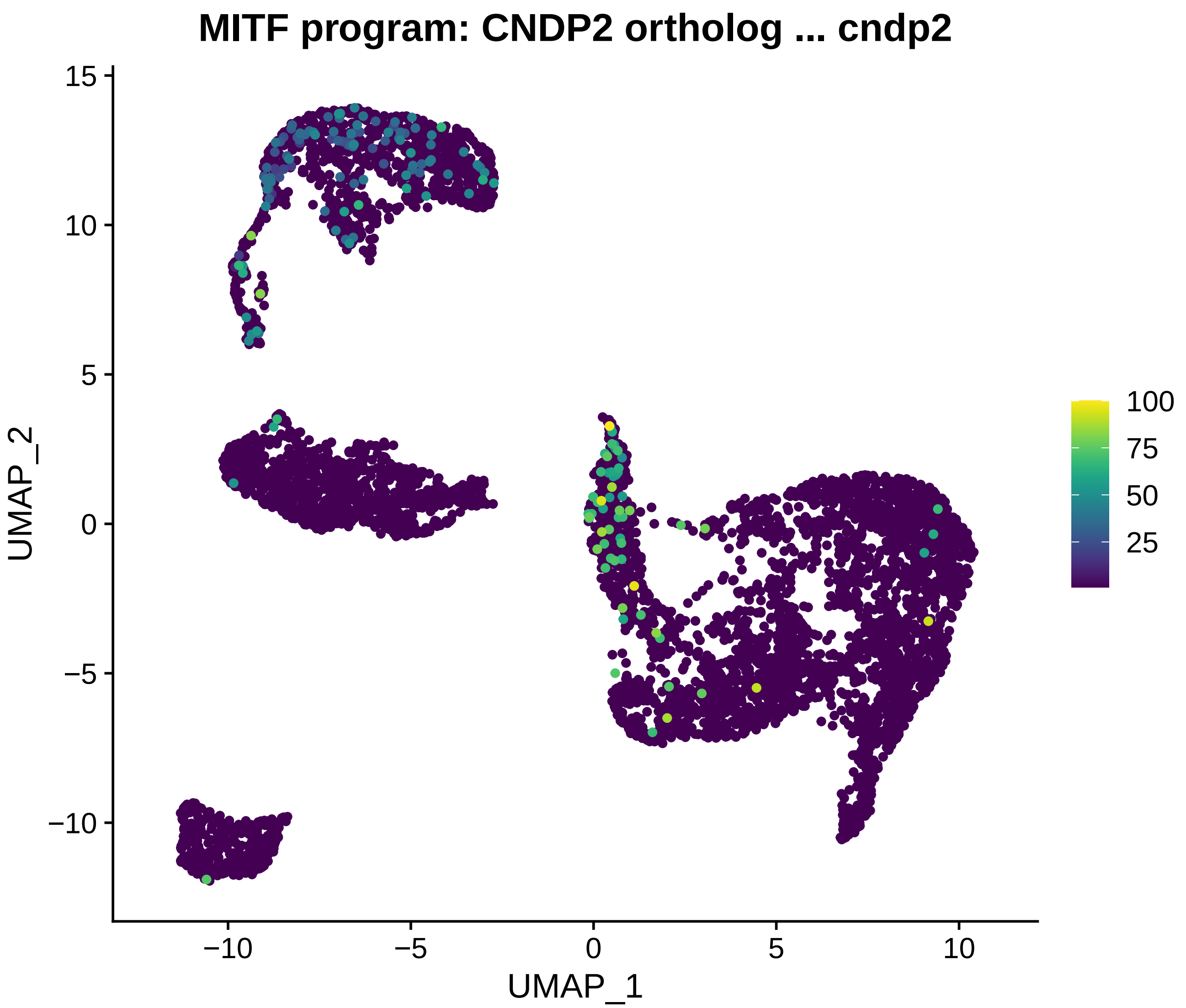

MITF program: CNDP2 ortholog ... cndp2
UMAP_2
UMAP_1
15
10
5
0
−5
−10
−10
−5
0
5
10
100
75
50
25

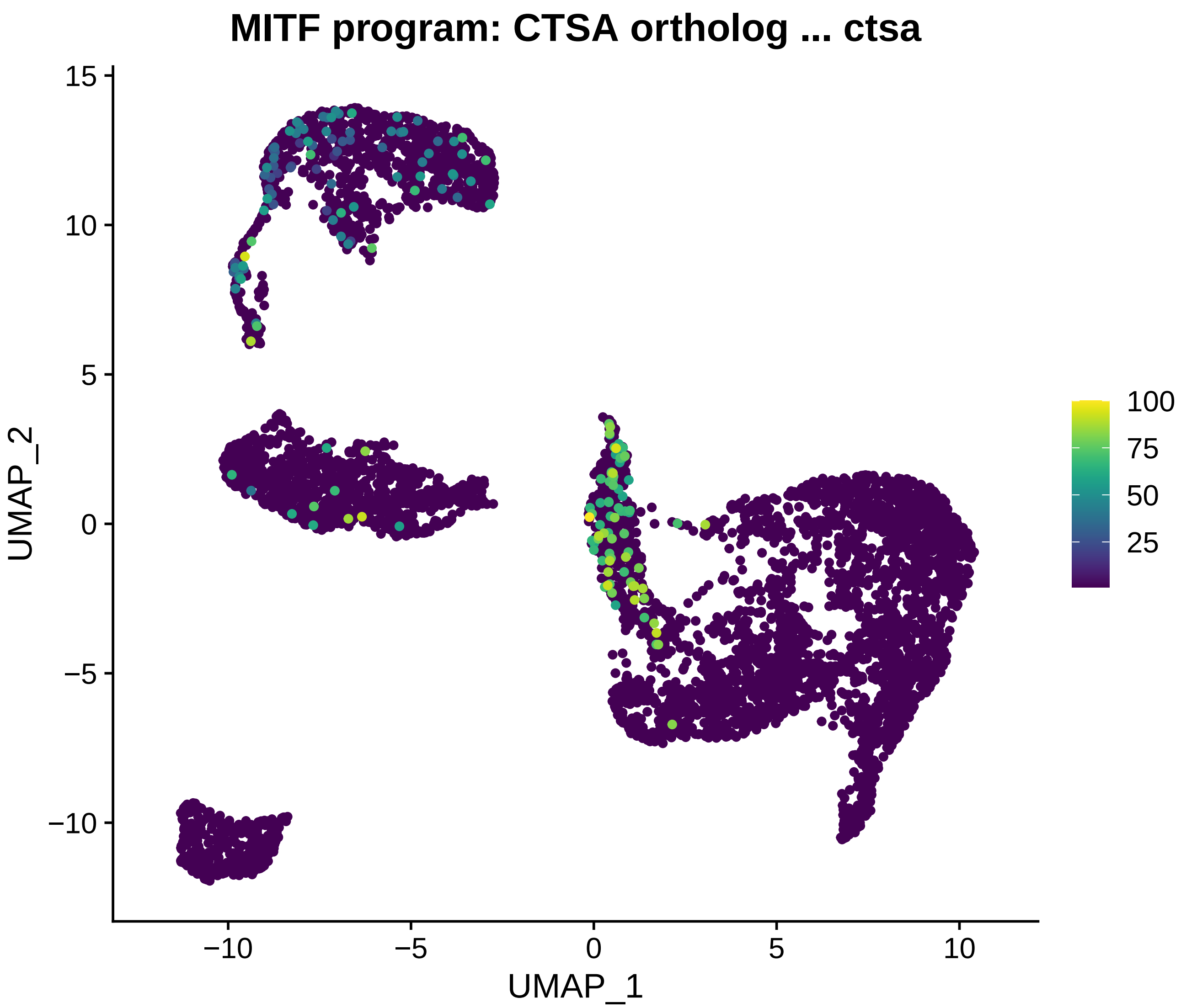
MITF program: CTSA ortholog ... ctsa
UMAP_2
UMAP_1
15
10
5
0
−5
−10
−10
−5
0
5
10
100
75
50
25

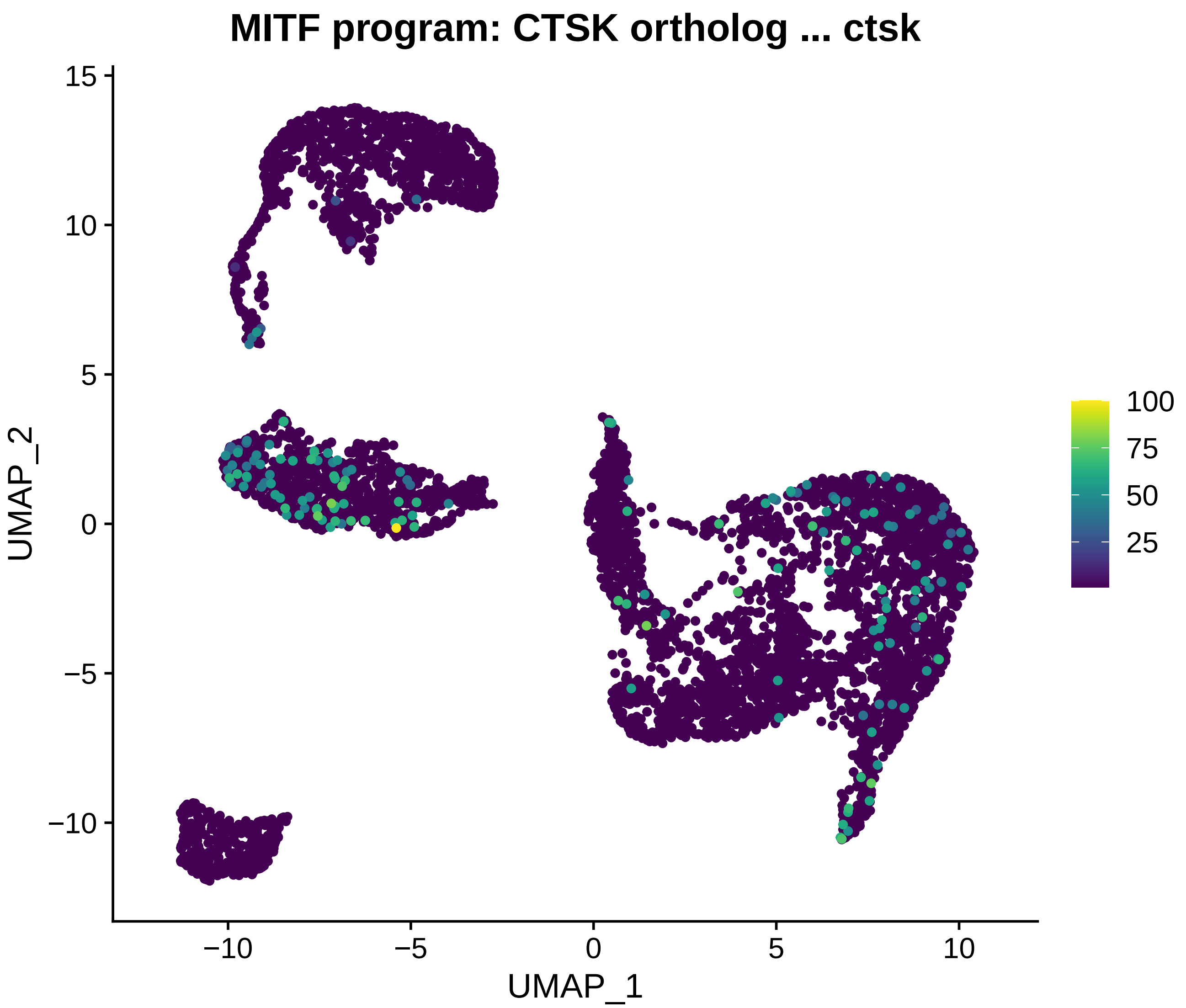
MITF program: CTSK ortholog ... ctsk
UMAP_2
UMAP_1
15
10
5
0
−5
−10
−10
−5
0
5
10
100
75
50
25

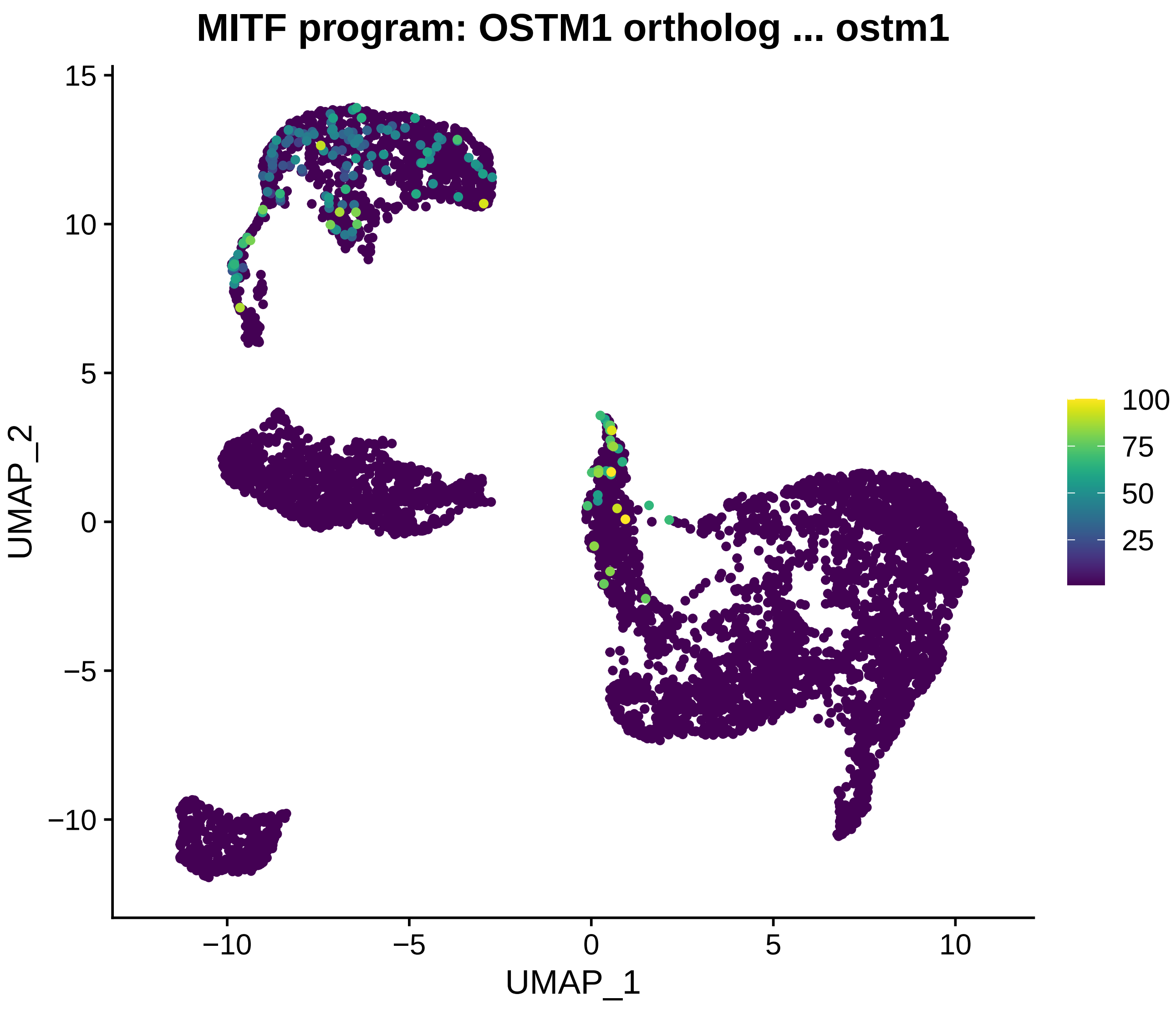

MITF program: OSTM1 ortholog ... ostm1
UMAP_2
UMAP_1
15
10
5
0
−5
−10
−10
−5
0
5
10
100
75
50
25

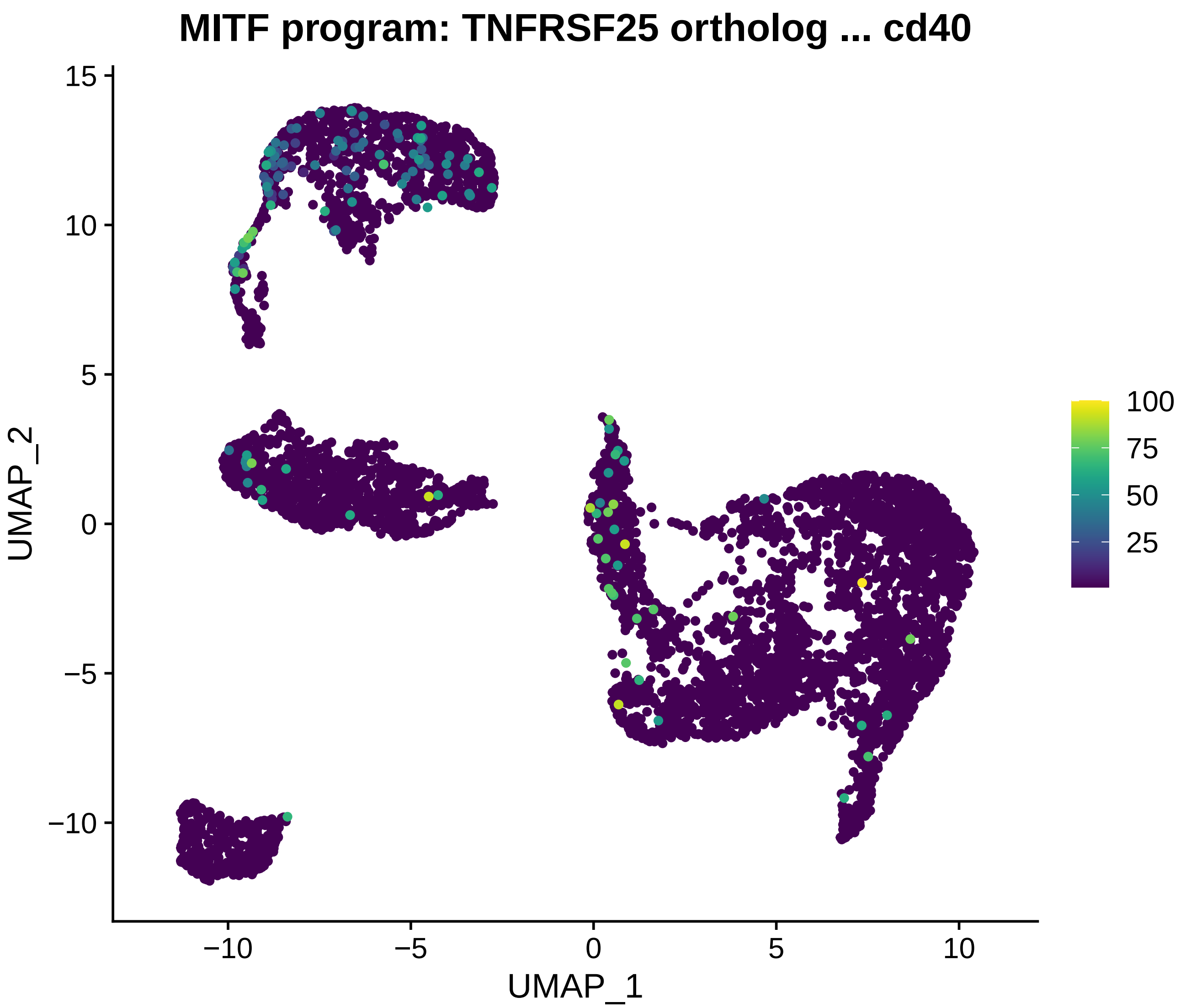
MITF program: TNFRSF25 ortholog ... cd40
UMAP_2
UMAP_1
15
10
5
0
−5
−10
−10
−5
0
5
10
100
75
50
25

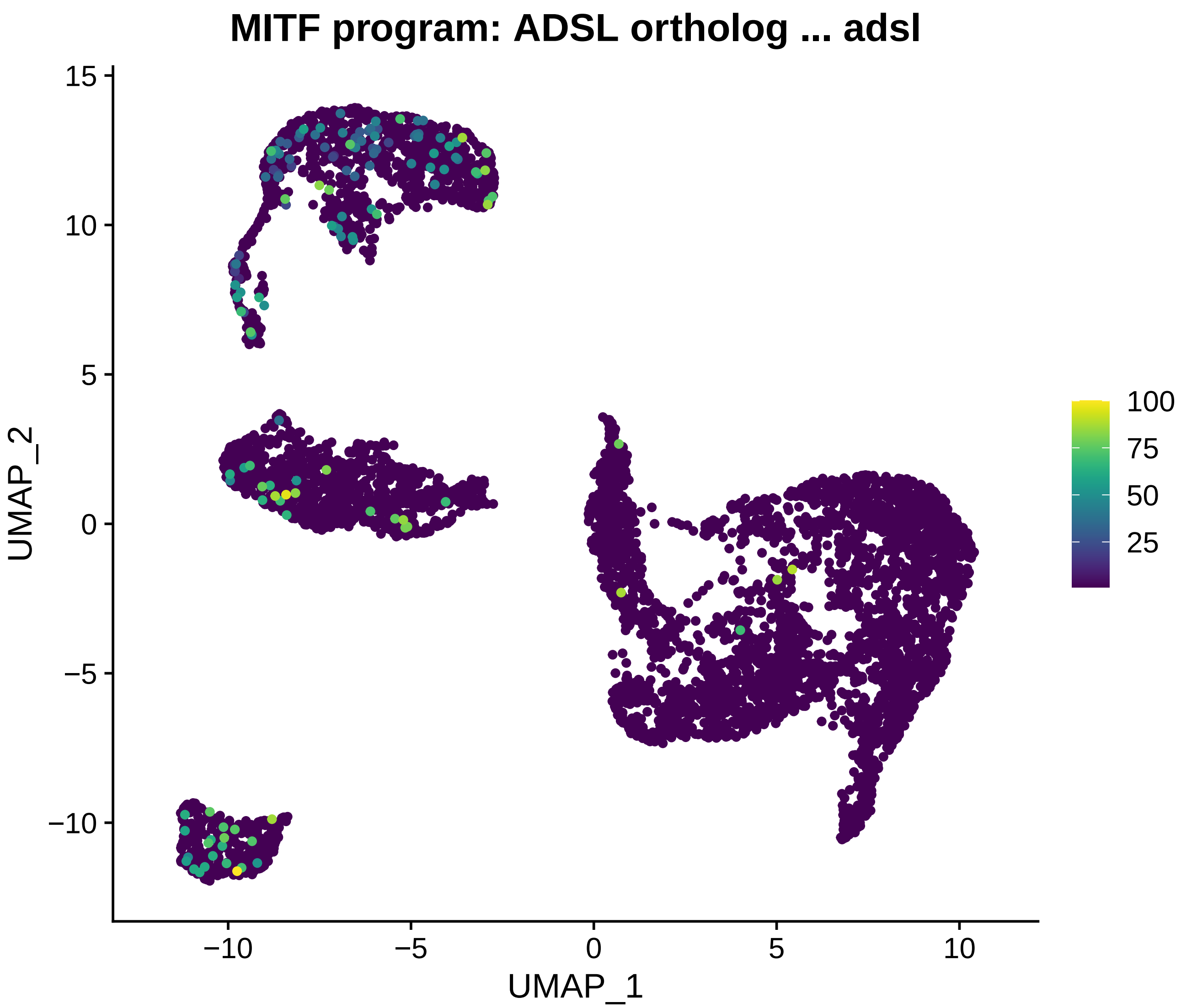

MITF program: ADSL ortholog ... adsl
UMAP_2
UMAP_1
15
10
5
0
−5
−10
−10
−5
0
5
10
100
75
50
25

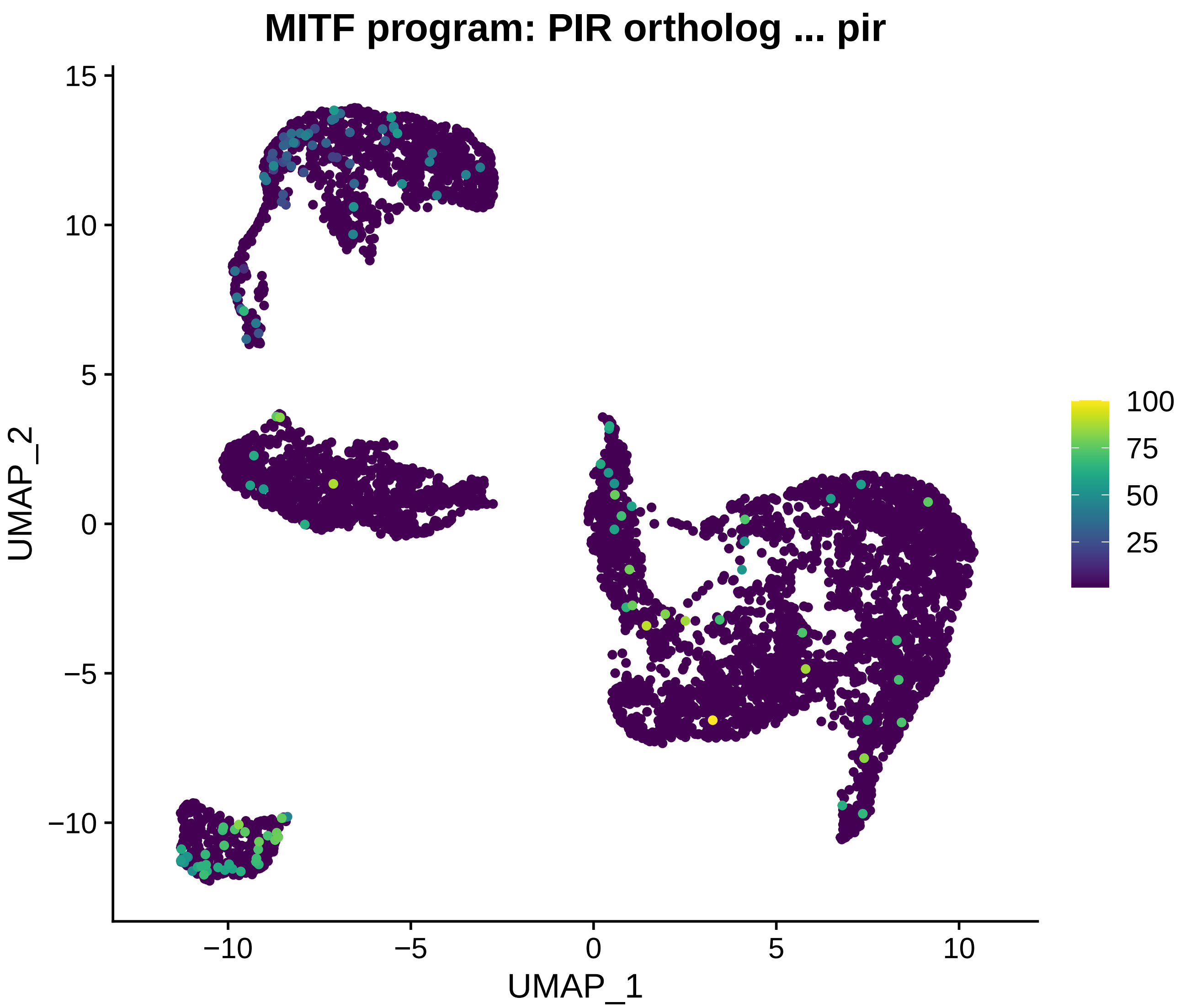
MITF program: PIR ortholog ... pir
UMAP_2
UMAP_1
15
10
5
0
−5
−10
−10
−5
0
5
10
100
75
50
25

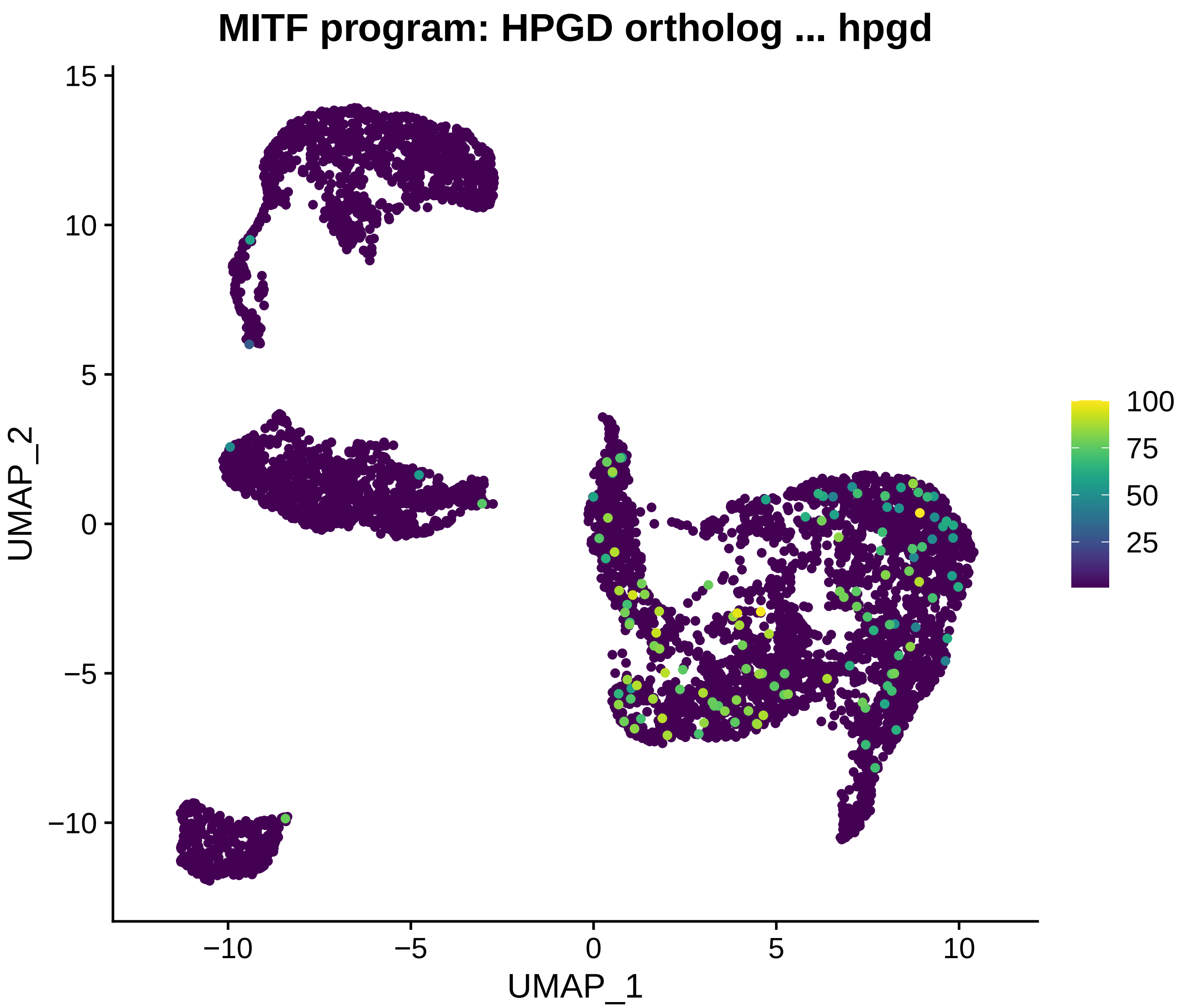

MITF program: HPGD ortholog ... hpgd
UMAP_2
UMAP_1
15
10
5
0
−5
−10
−10
−5
0
5
10
100
75
50
25

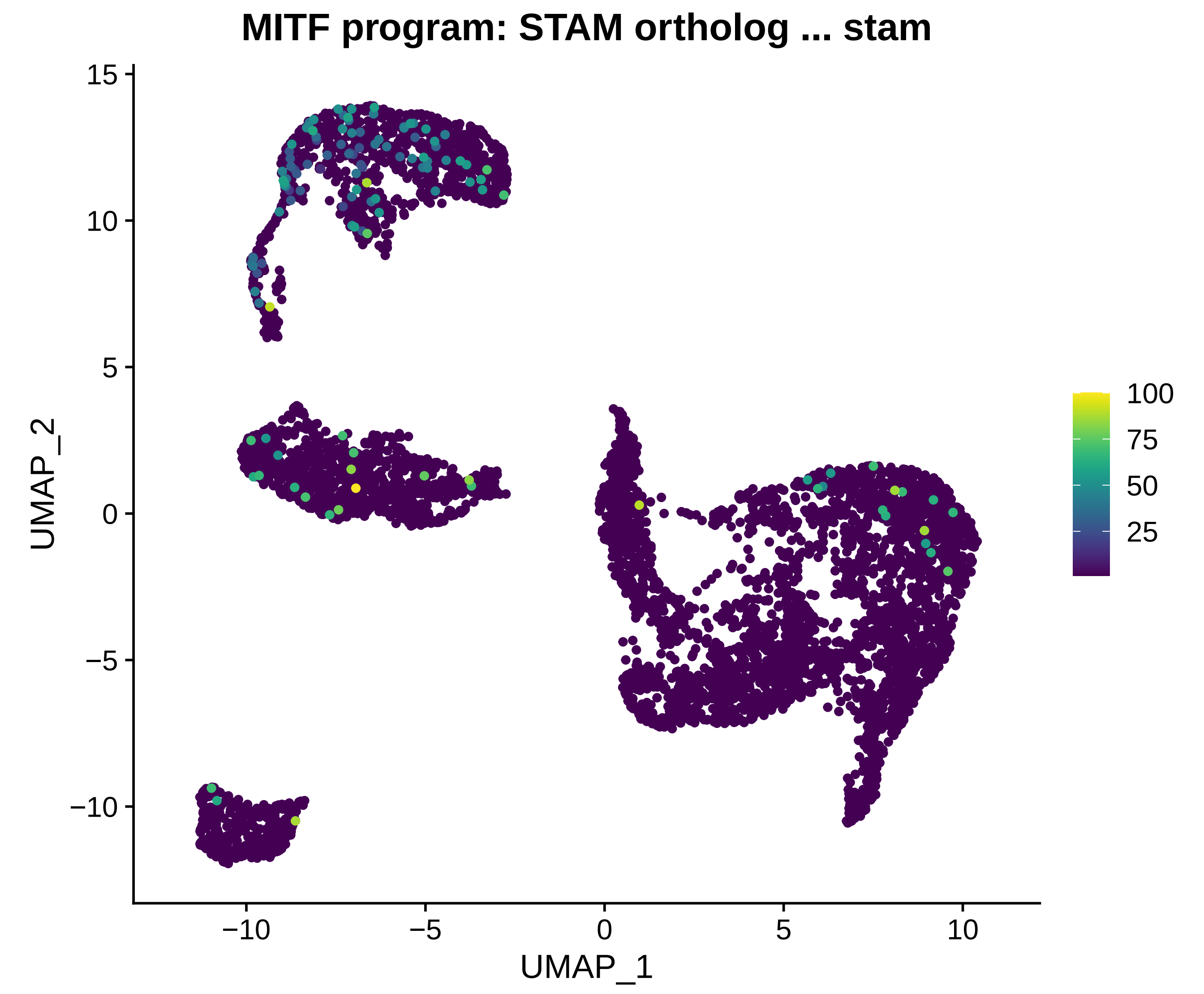

MITF program: STAM ortholog ... stam
UMAP_2
UMAP_1
15
10
5
0
−5
−10
−10
−5
0
5
10
100
75
50
25

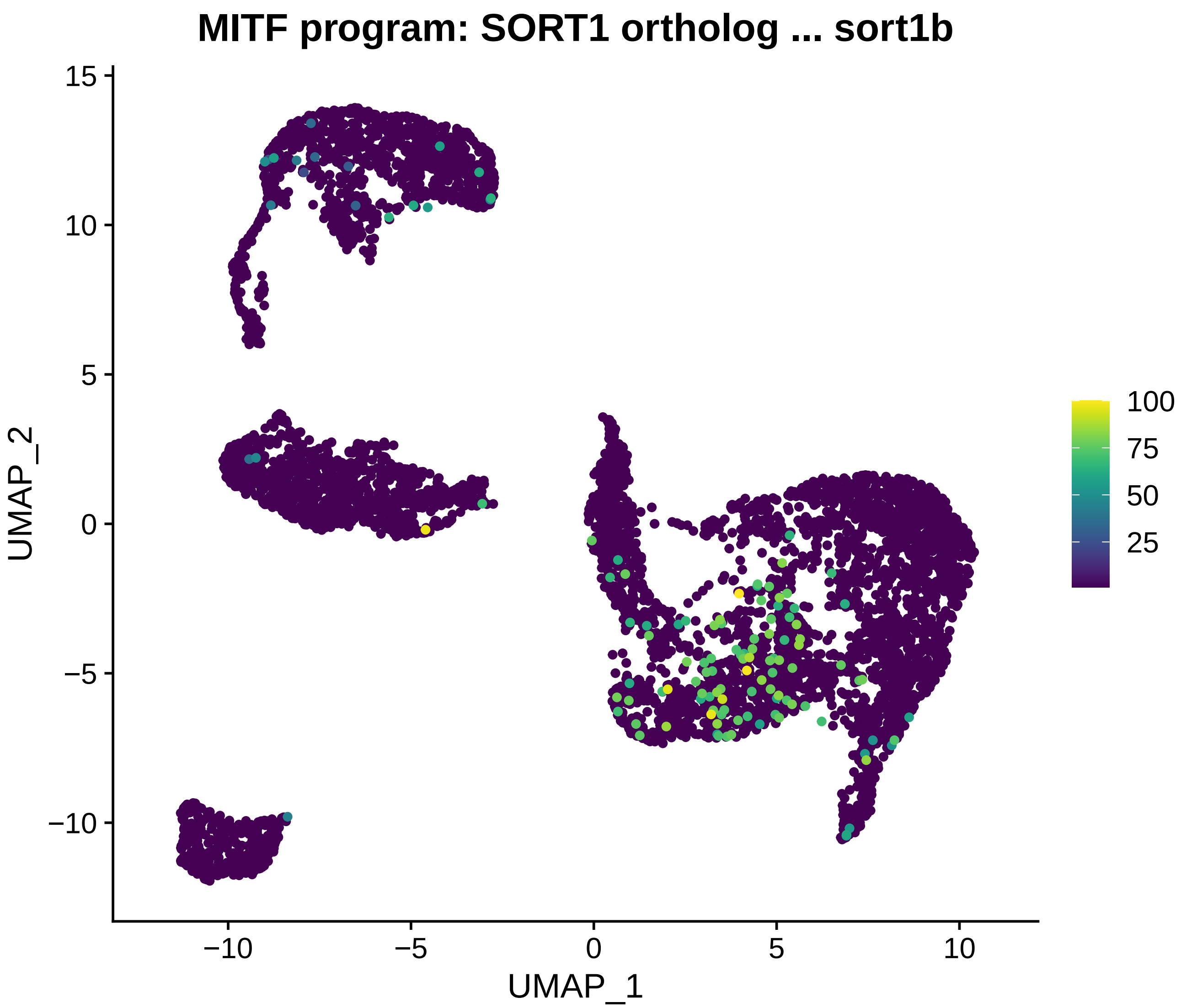
MITF program: SORT1 ortholog ... sort1b
UMAP_1
UMAP_2
−10
−5
0
5
10
15
100
75
50
25

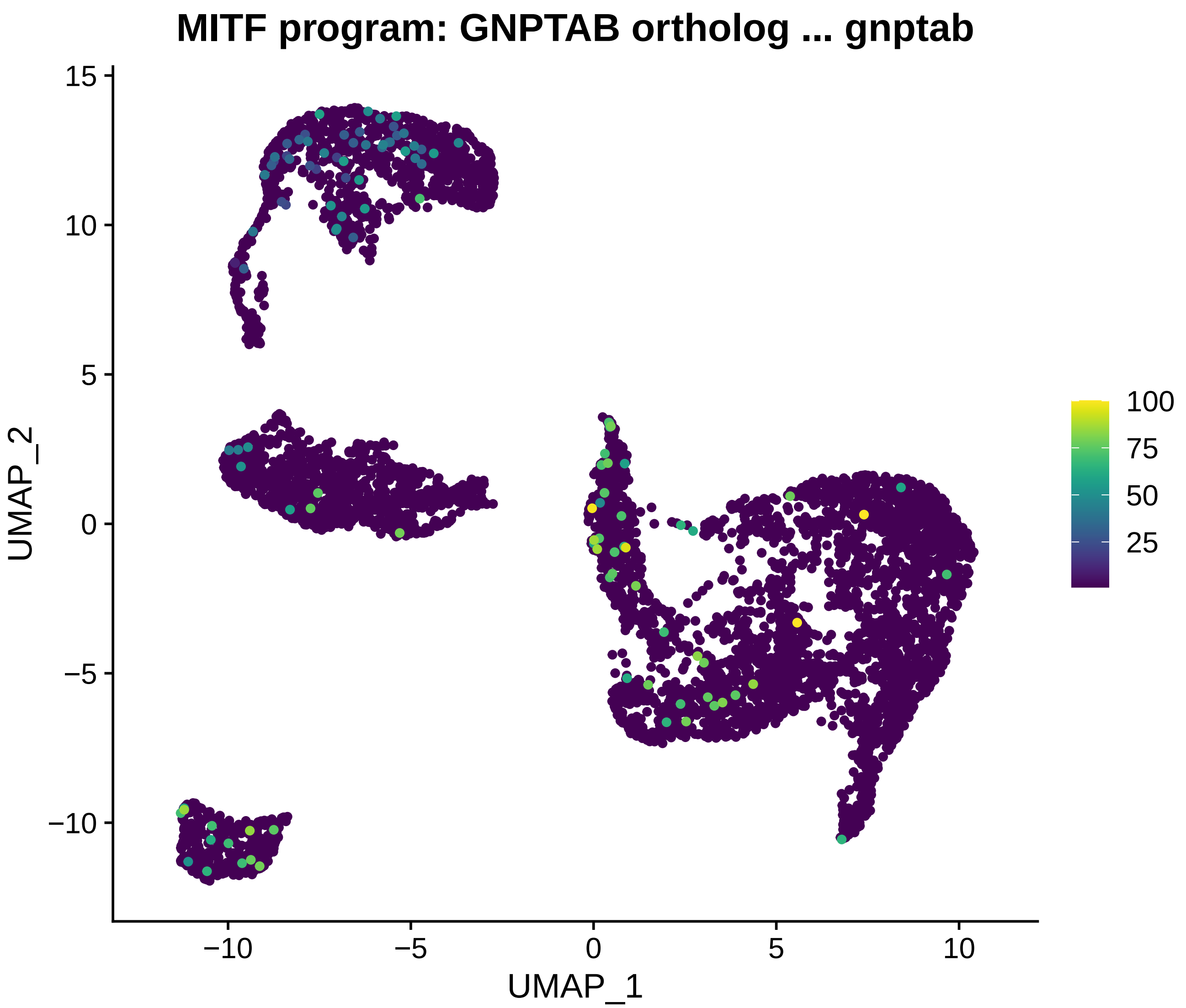

MITF program: GNPTAB ortholog ... gnptab
UMAP_2
UMAP_1
15
10
5
0
−5
−10
−10
−5
0
5
10
100
75
50
25

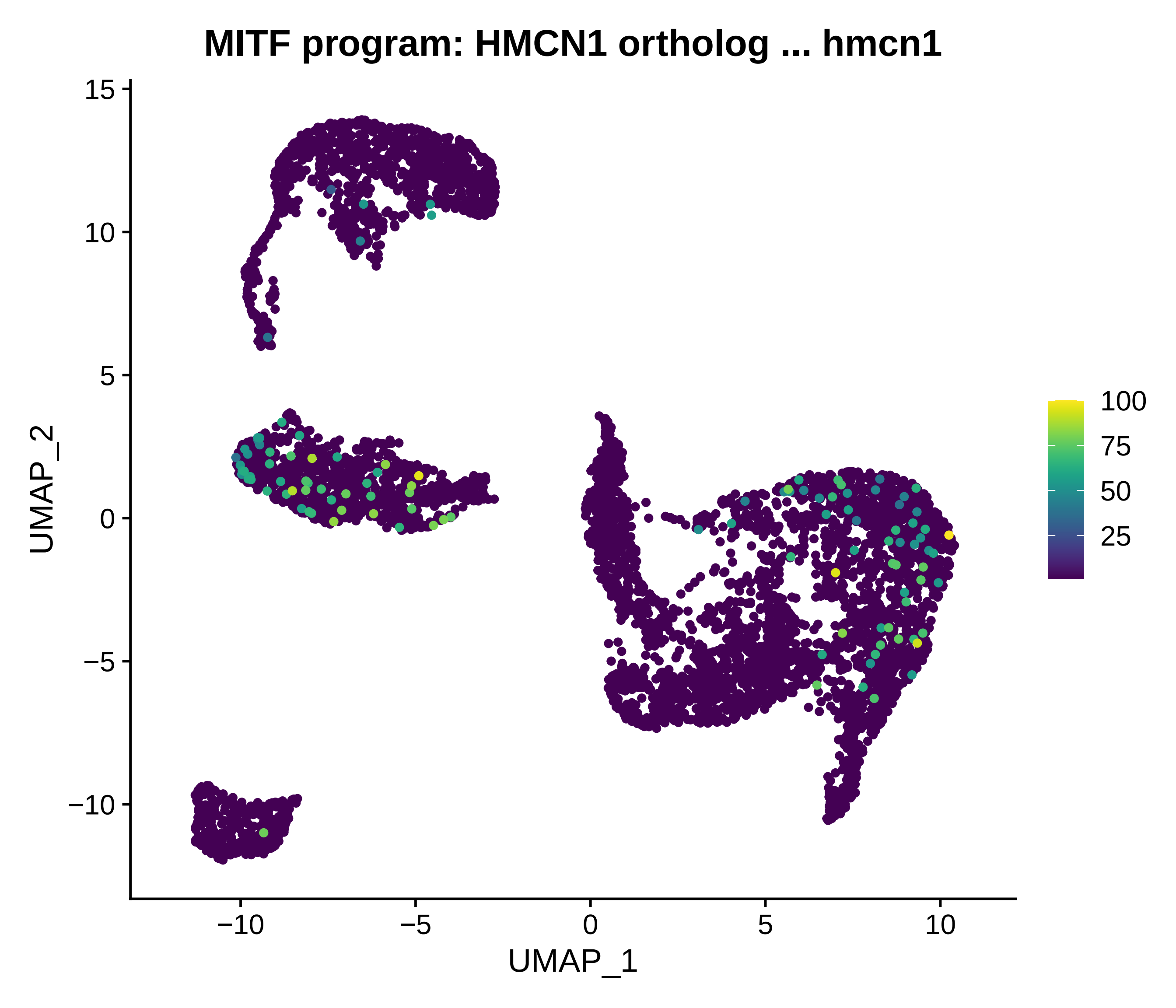
MITF program: HMCN1 ortholog ... hmcn1
15
10
5
0
−5
−10
UMAP_2
−10
−5
0
5
10
UMAP_1
100
75
50
25

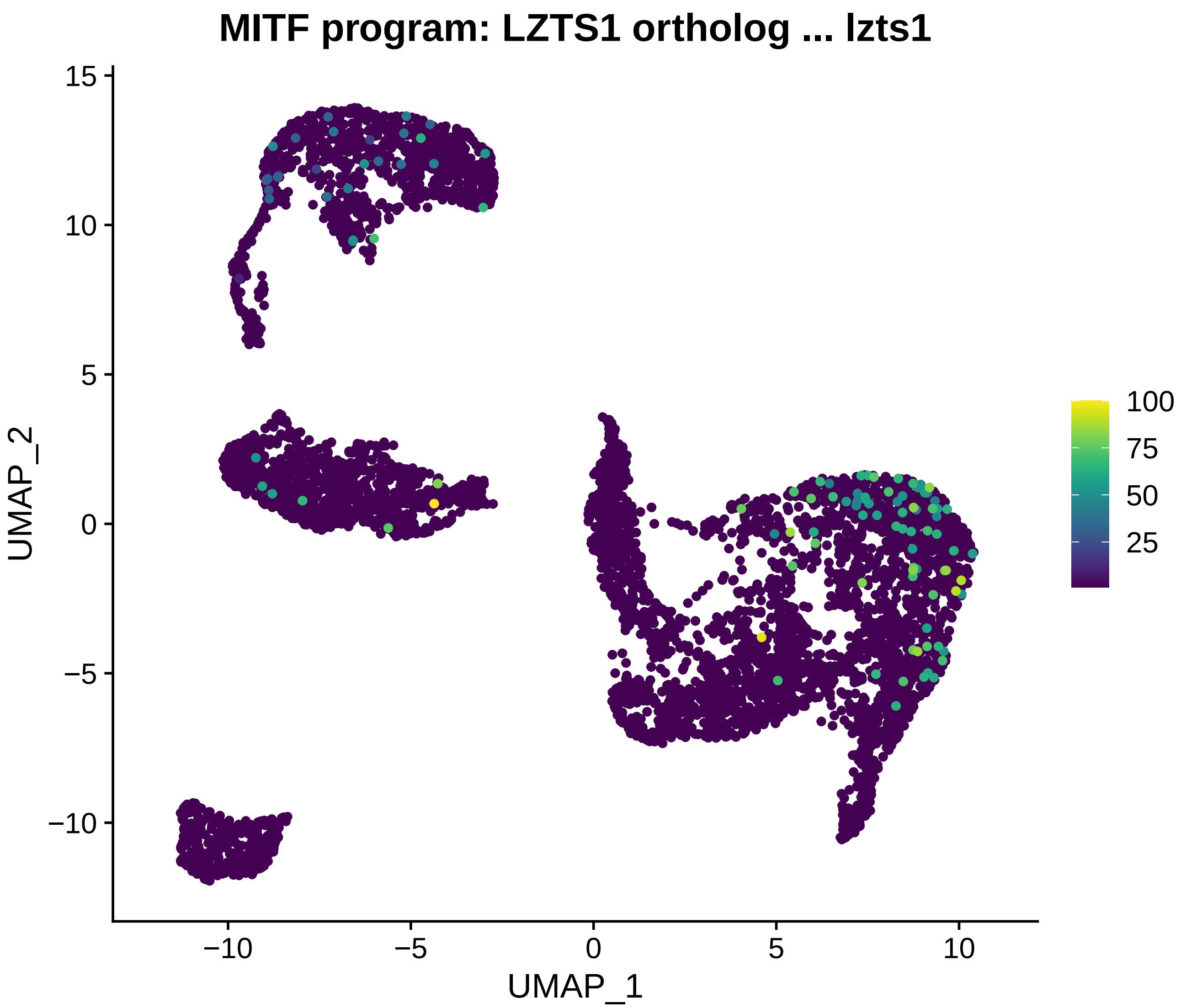

MITF program: LZTS1 ortholog ... lzts1
15
10
5
0
−5
−10
UMAP_2
−10
−5
0
5
10
UMAP_1
100
75
50
25

### MITF program: TYR ortholog ... tyr

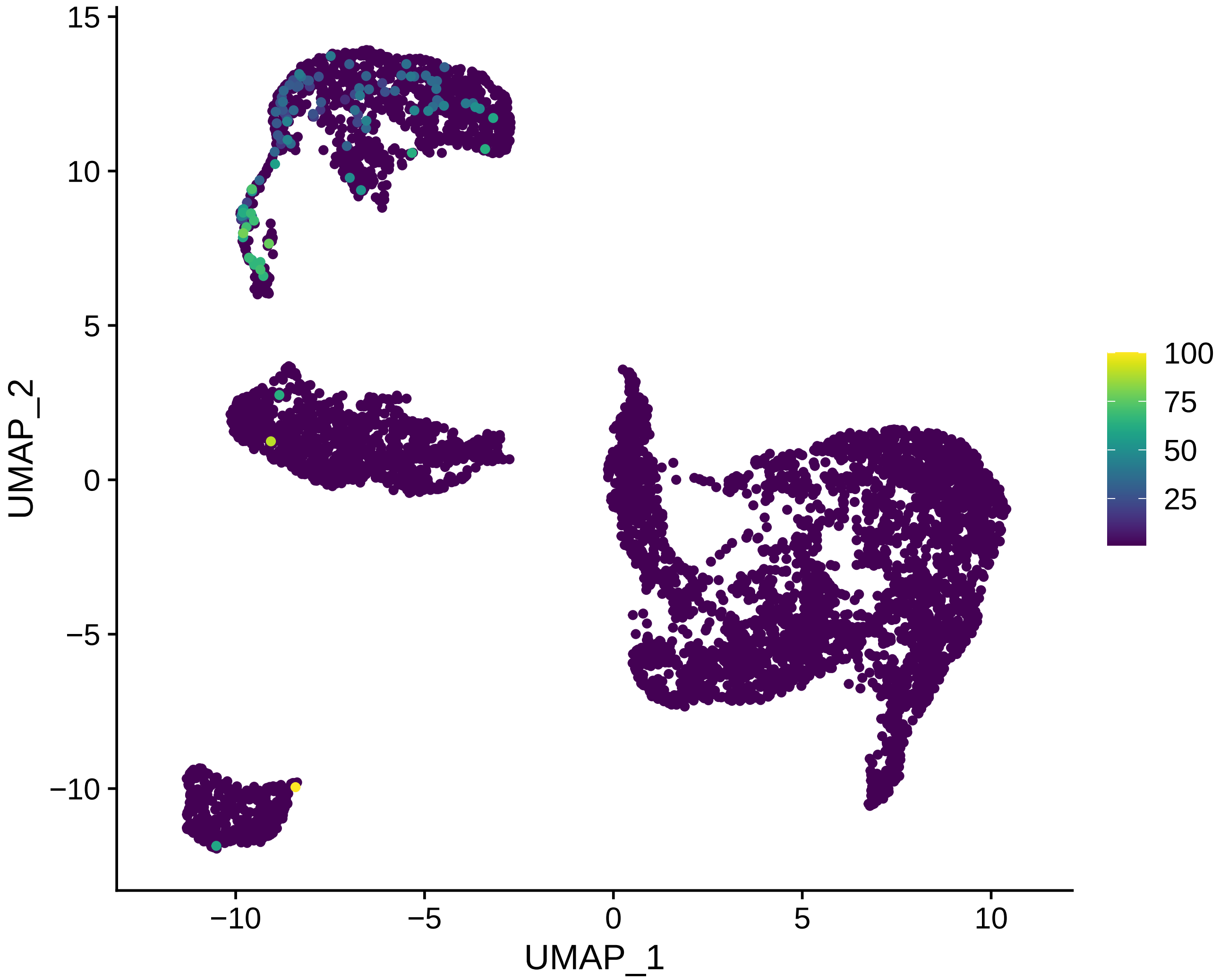

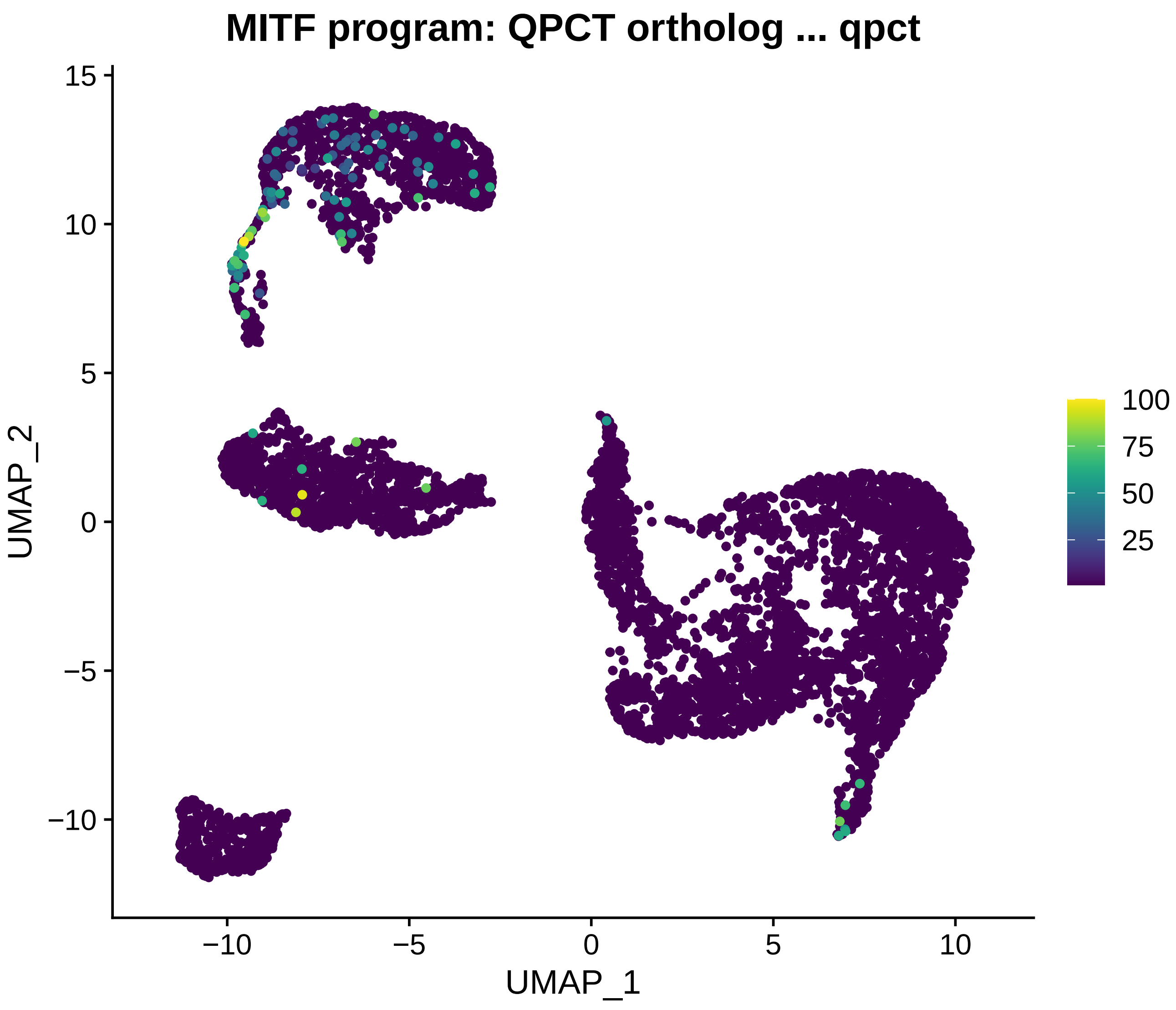
MITF program: QPCT ortholog ... qpct
UMAP_2
UMAP_1
15
10
5
0
−5
−10
−10
−5
0
5
10
100
75
50
25

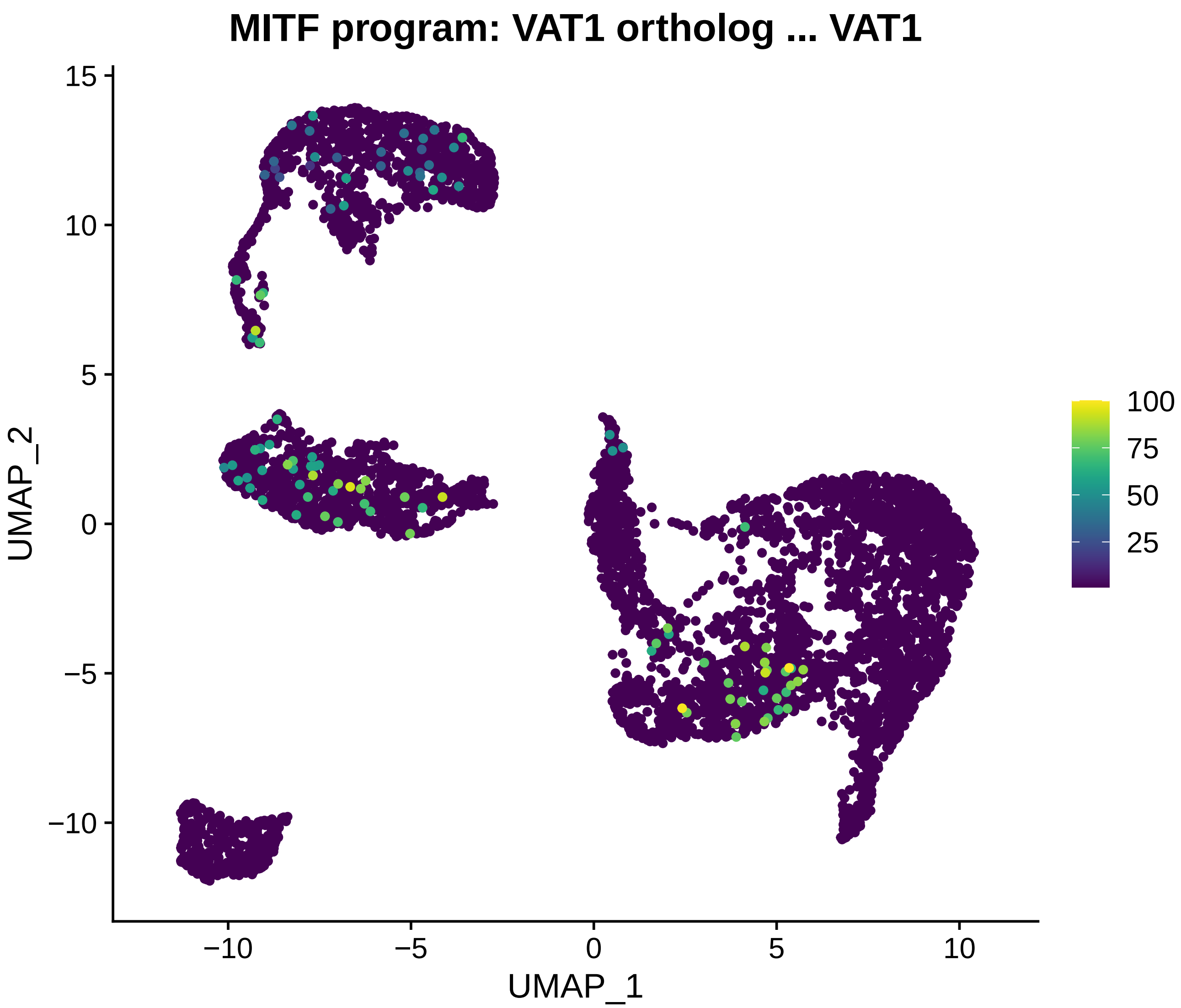
MITF program: VAT1 ortholog ... VAT1
UMAP_2
UMAP_1
15
10
5
0
−5
−10
−10
−5
0
5
10
100
75
50
25

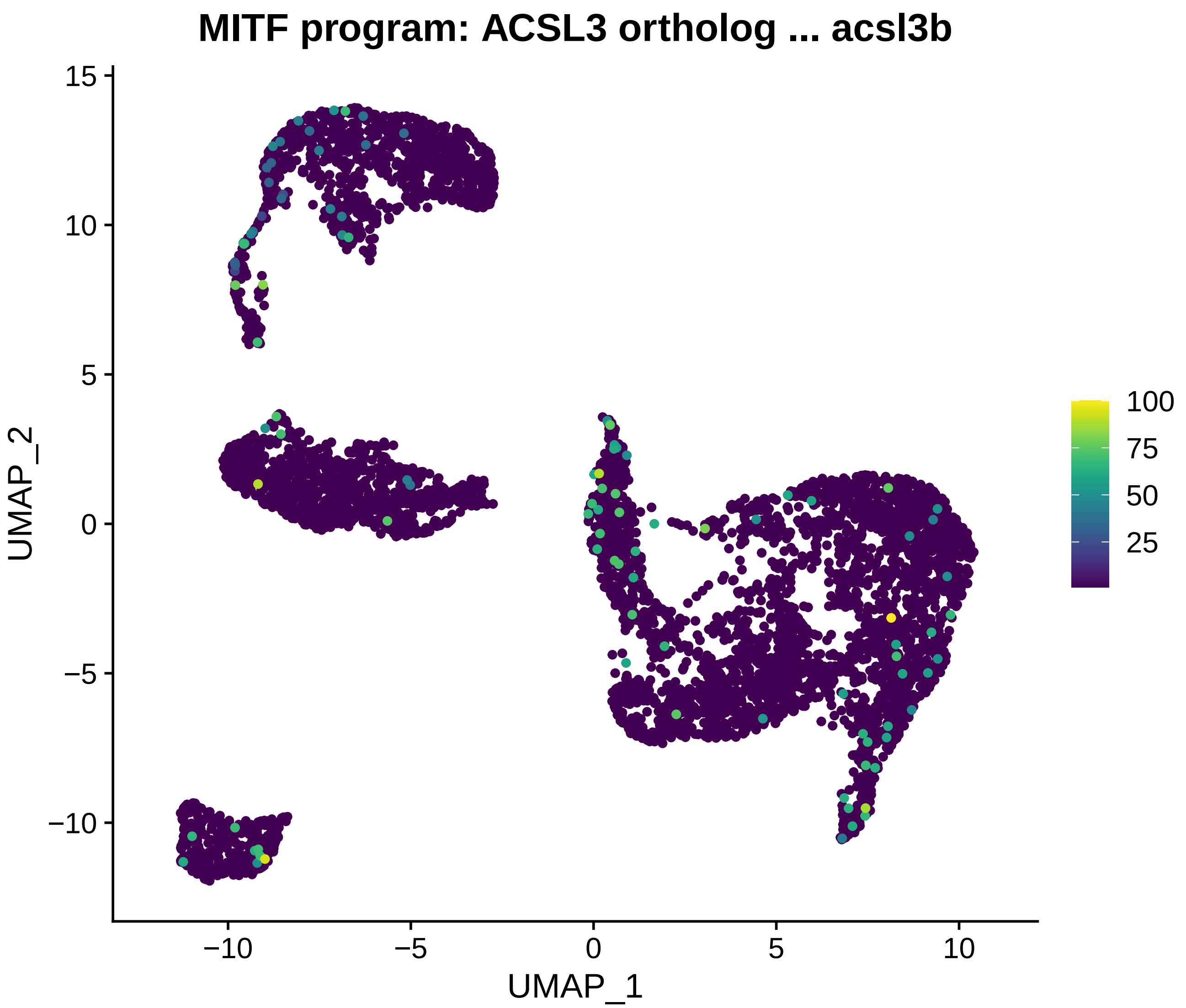
MITF program: ACSL3 ortholog ... acsl3b
UMAP_2
UMAP_1
15
10
5
0
−5
−10
−10
−5
0
5
10
100
75
50
25

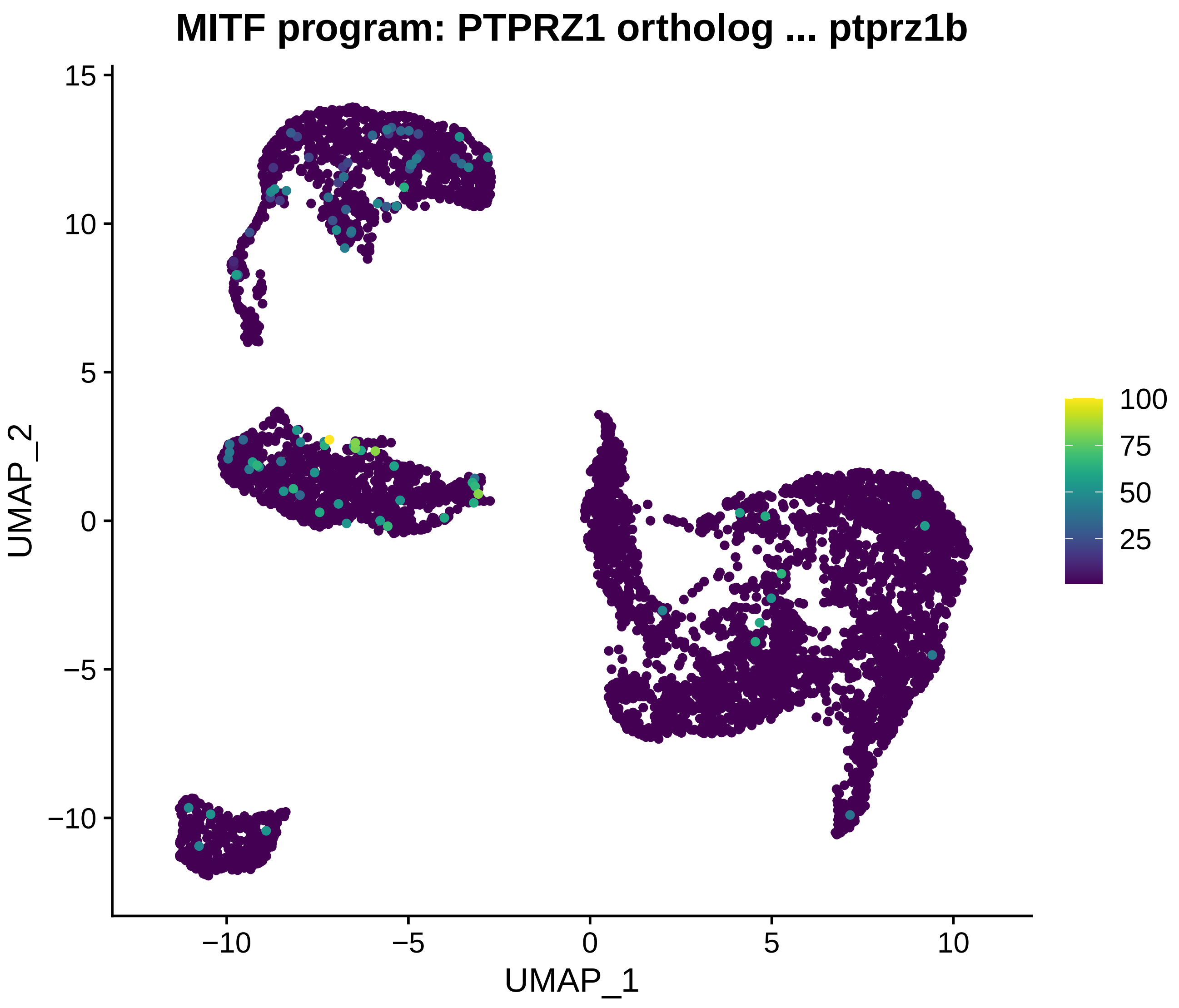
MITF program: PTPRZ1 ortholog ... ptprz1b
UMAP_2
UMAP_1
15
10
5
0
−5
−10
−10
−5
0
5
10
100
75
50
25

**MITF program: MAGED4 ortholog ... ndnl2**

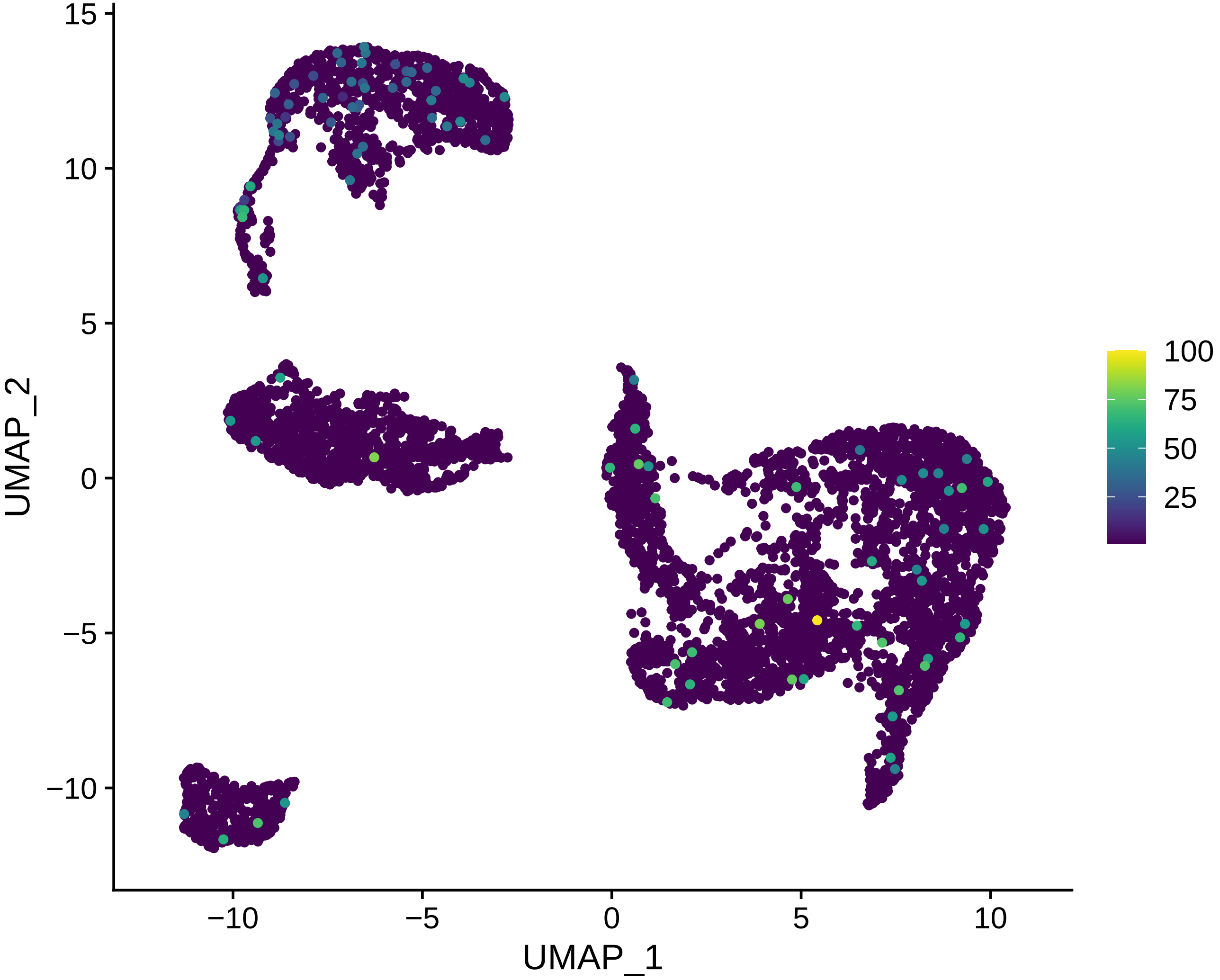

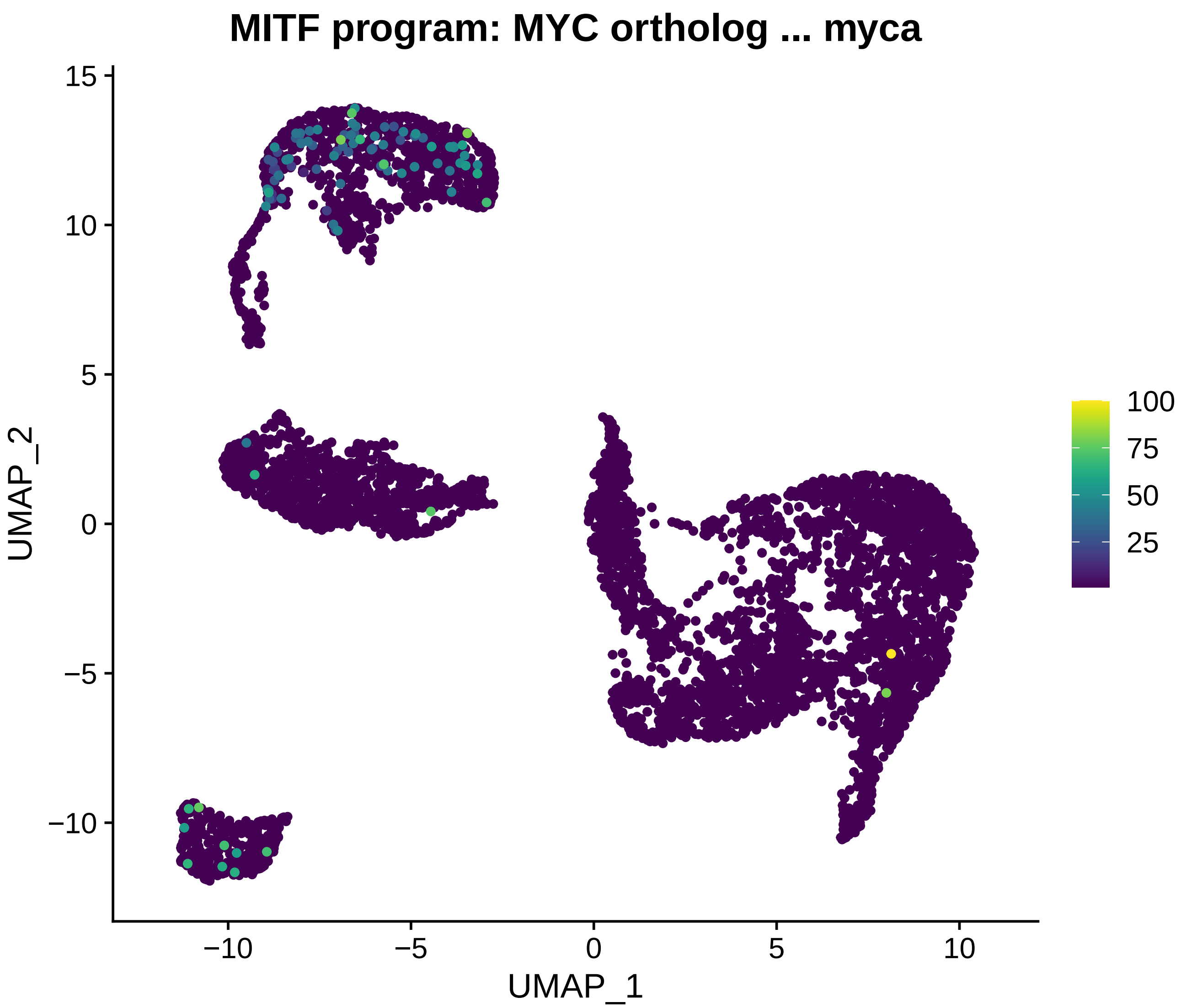
MITF program: MYC ortholog ... myca
UMAP_2
UMAP_1
15
10
5
0
−5
−10
−10
−5
0
5
10
100
75
50
25

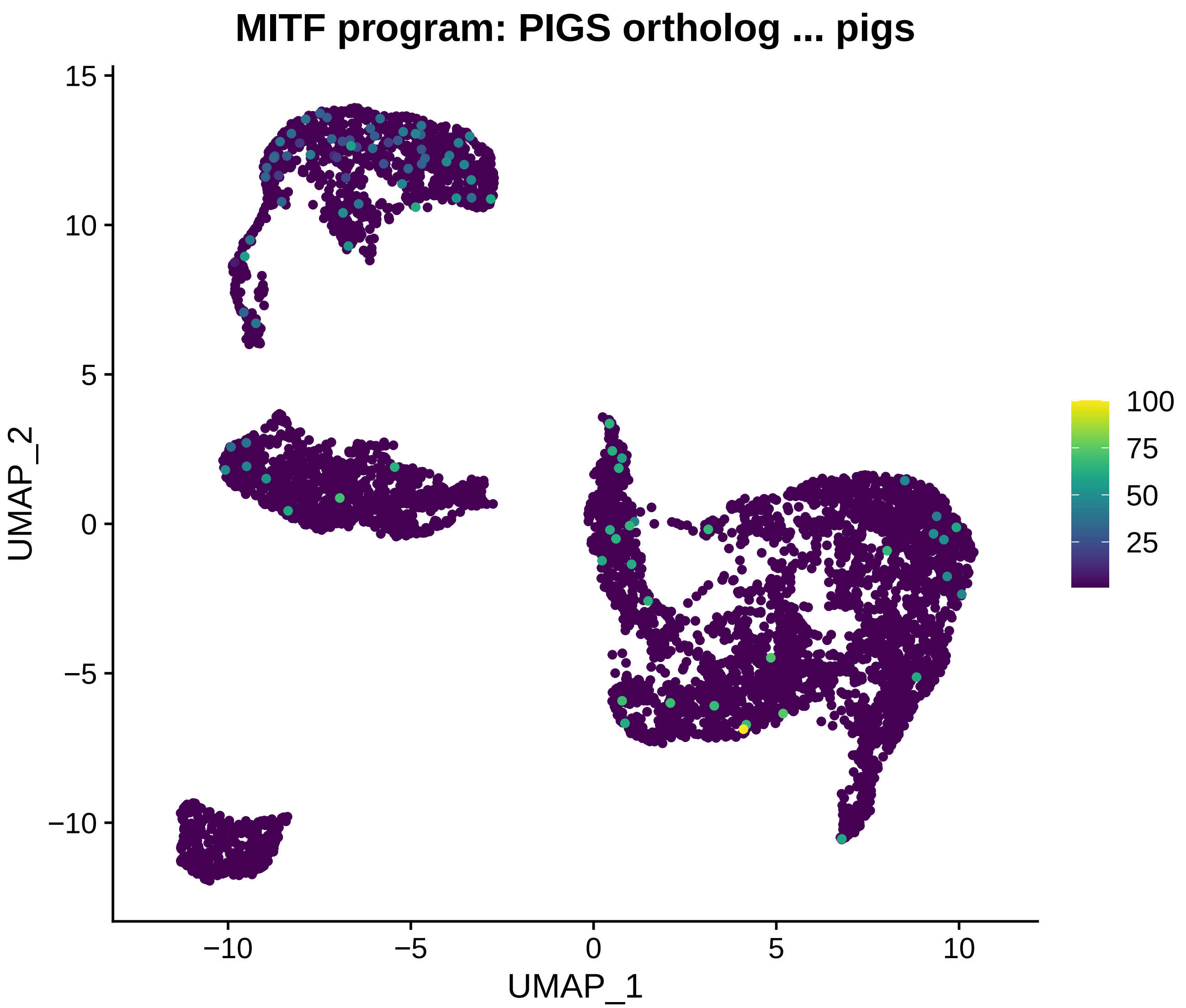
MITF program: PIGS ortholog ... pigs
UMAP_2
UMAP_1
15
10
5
0
−5
−10
−10
−5
0
5
10
100
75
50
25

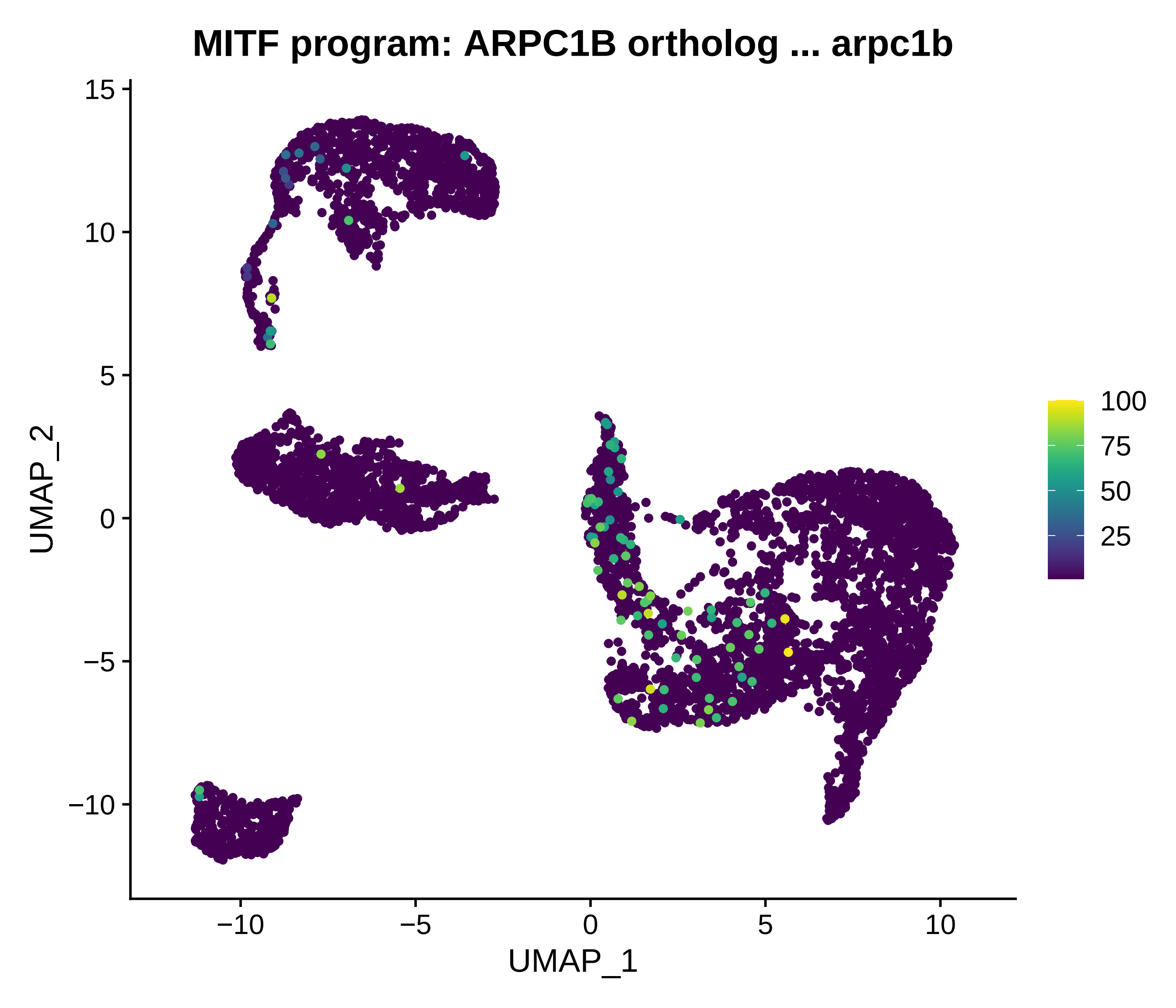
MITF program: ARPC1B ortholog ... arpc1b
UMAP_2
UMAP_1
15
10
5
0
−5
−10
−10
−5
0
5
10
100
75
50
25

### MITF program: APOE ortholog ... apoeb

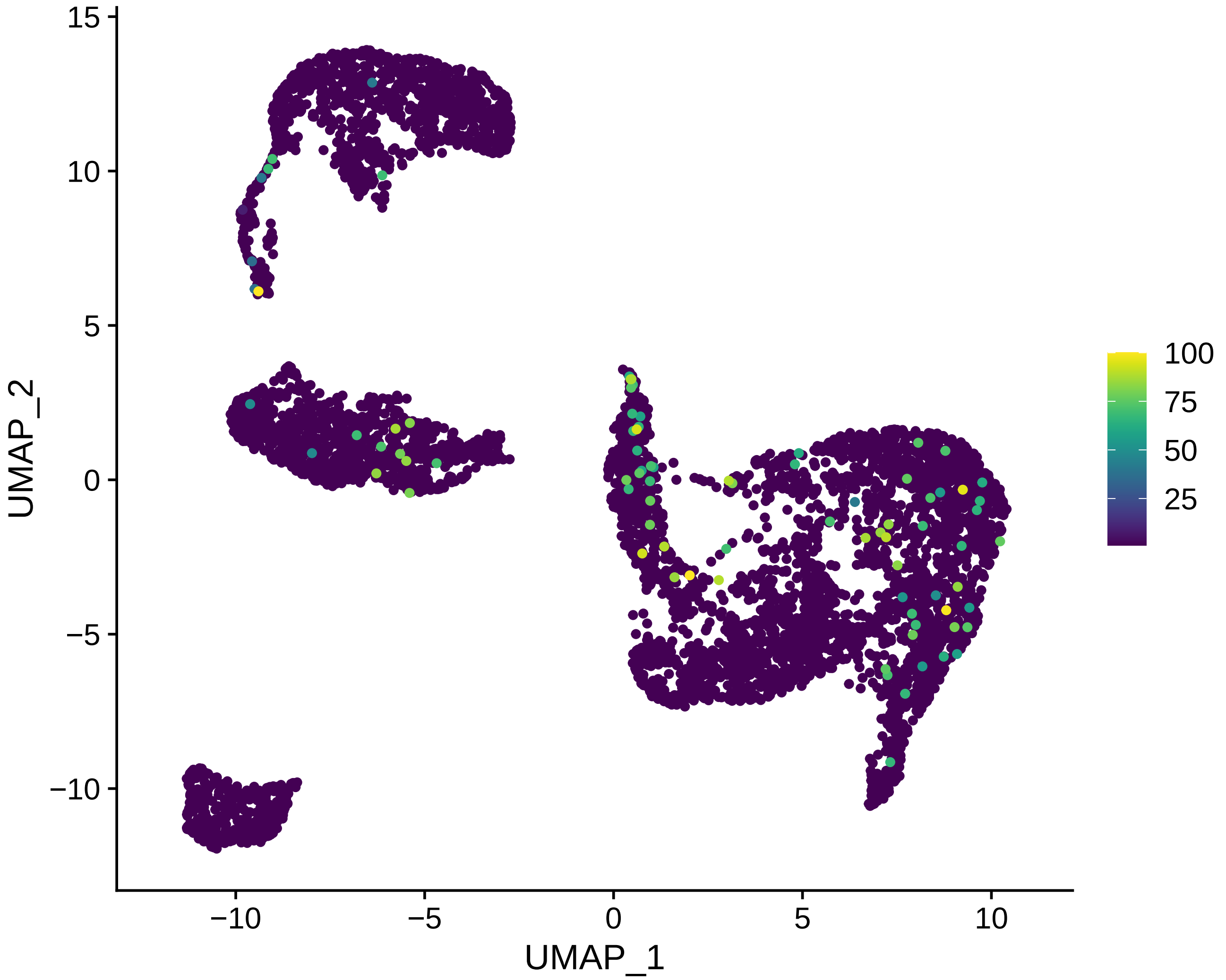

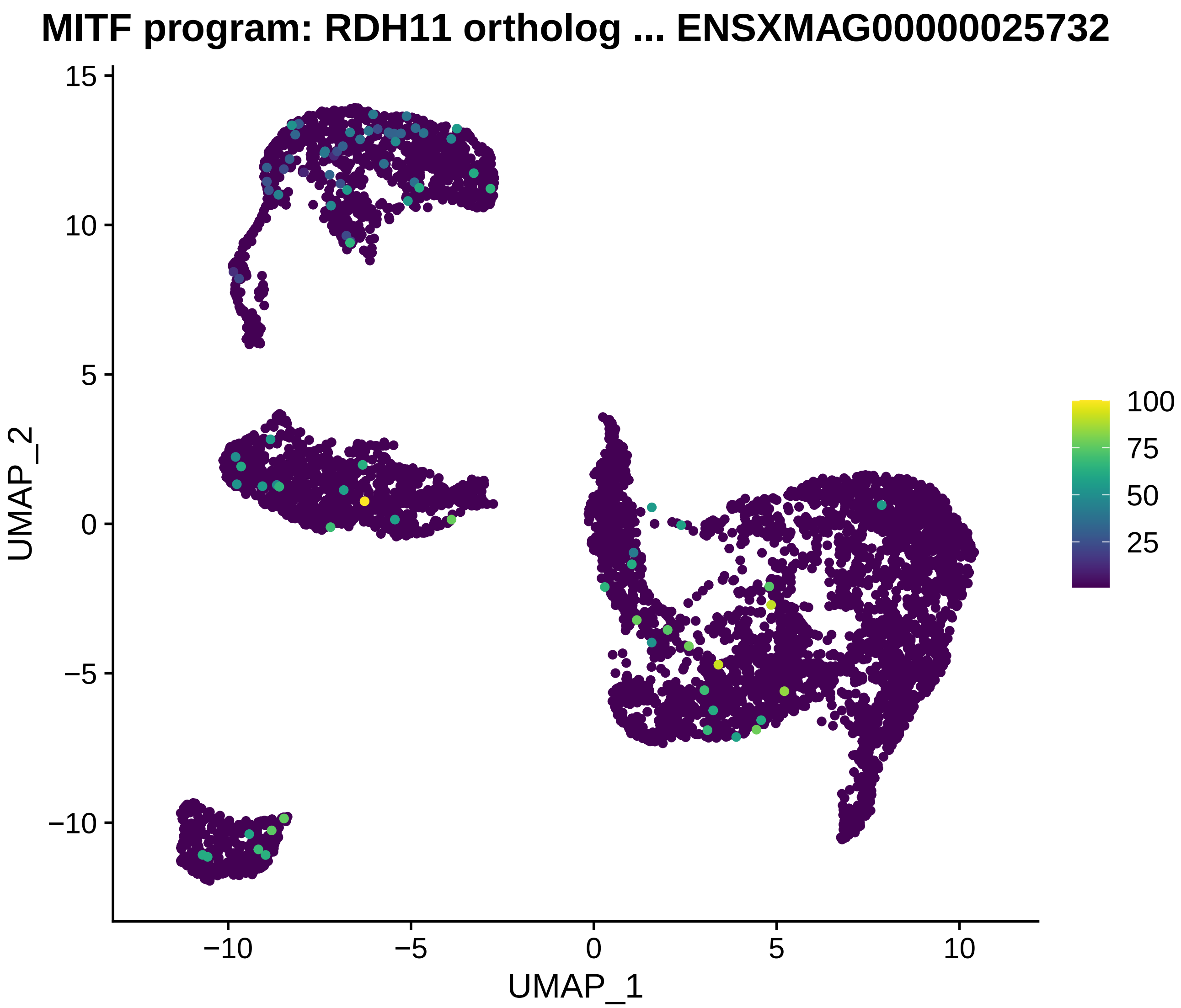

MITF program: RDH11 ortholog ... ENSXMAG00000025732
UMAP_2
UMAP_1
15
10
5
0
−5
−10
−10
−5
0
5
10
100
75
50
25

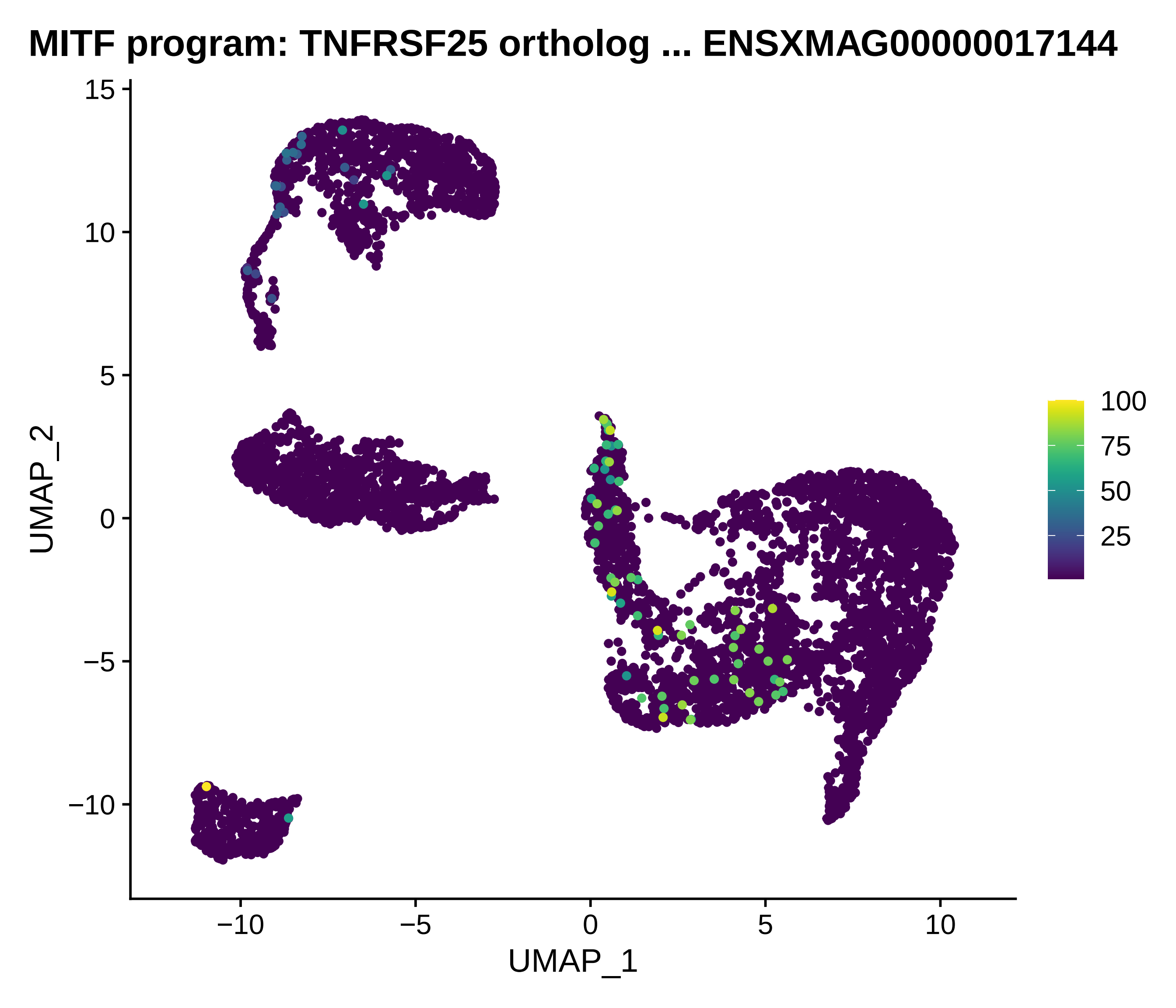
MITF program: TNFRSF25 ortholog ... ENSXMAG00000017144
UMAP_2
UMAP_1
15
10
5
0
−5
−10
−10
−5
0
5
10
100
75
50
25

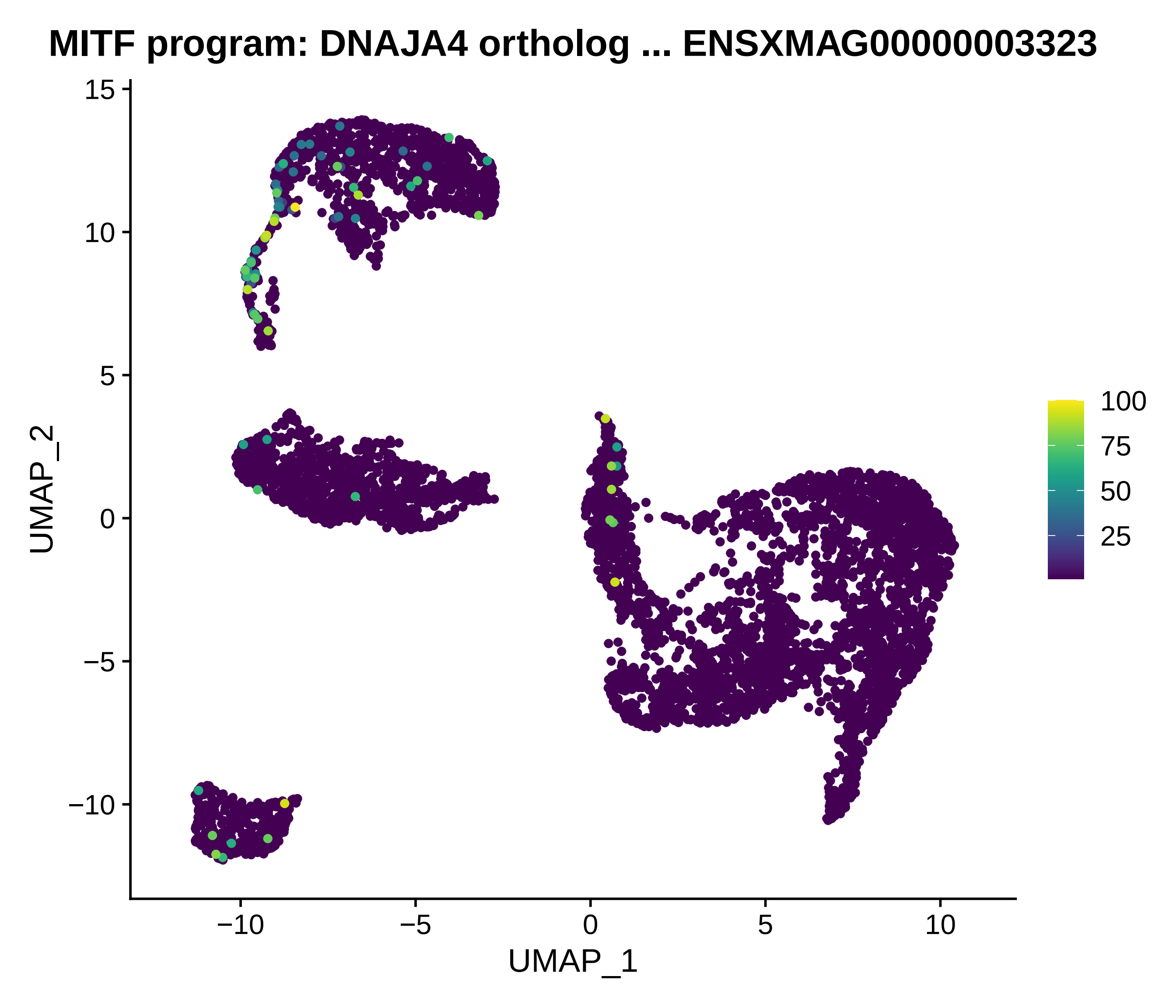
MITF program: DNAJA4 ortholog ... ENSXMAG00000003323
UMAP_2
UMAP_1
15
10
5
0
−5
−10
−10
−5
0
5
10
100
75
50
25

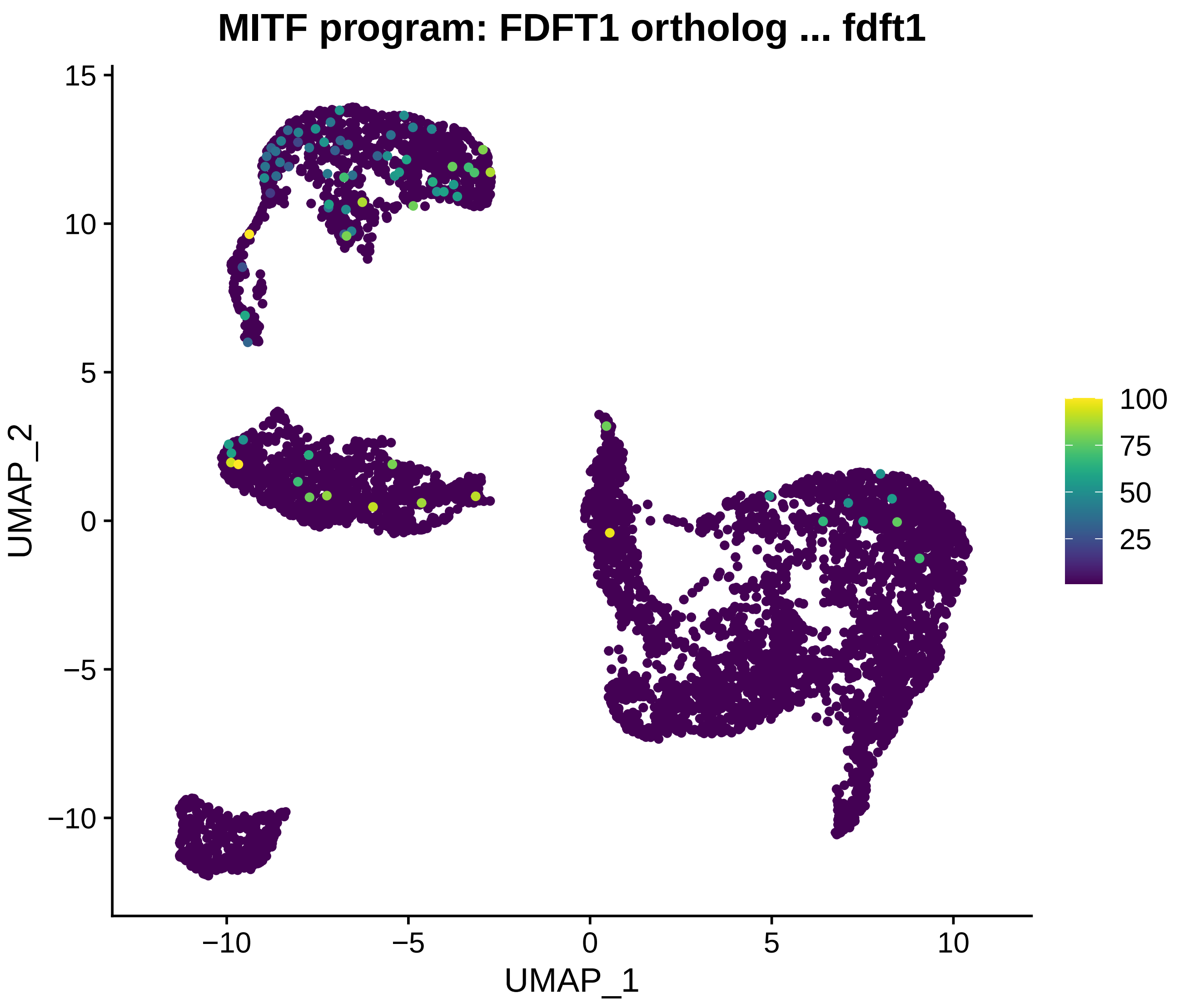
MITF program: FDFT1 ortholog ... fdft1
UMAP_2
UMAP_1
15
10
5
0
−5
−10
−10
−5
0
5
10
100
75
50
25

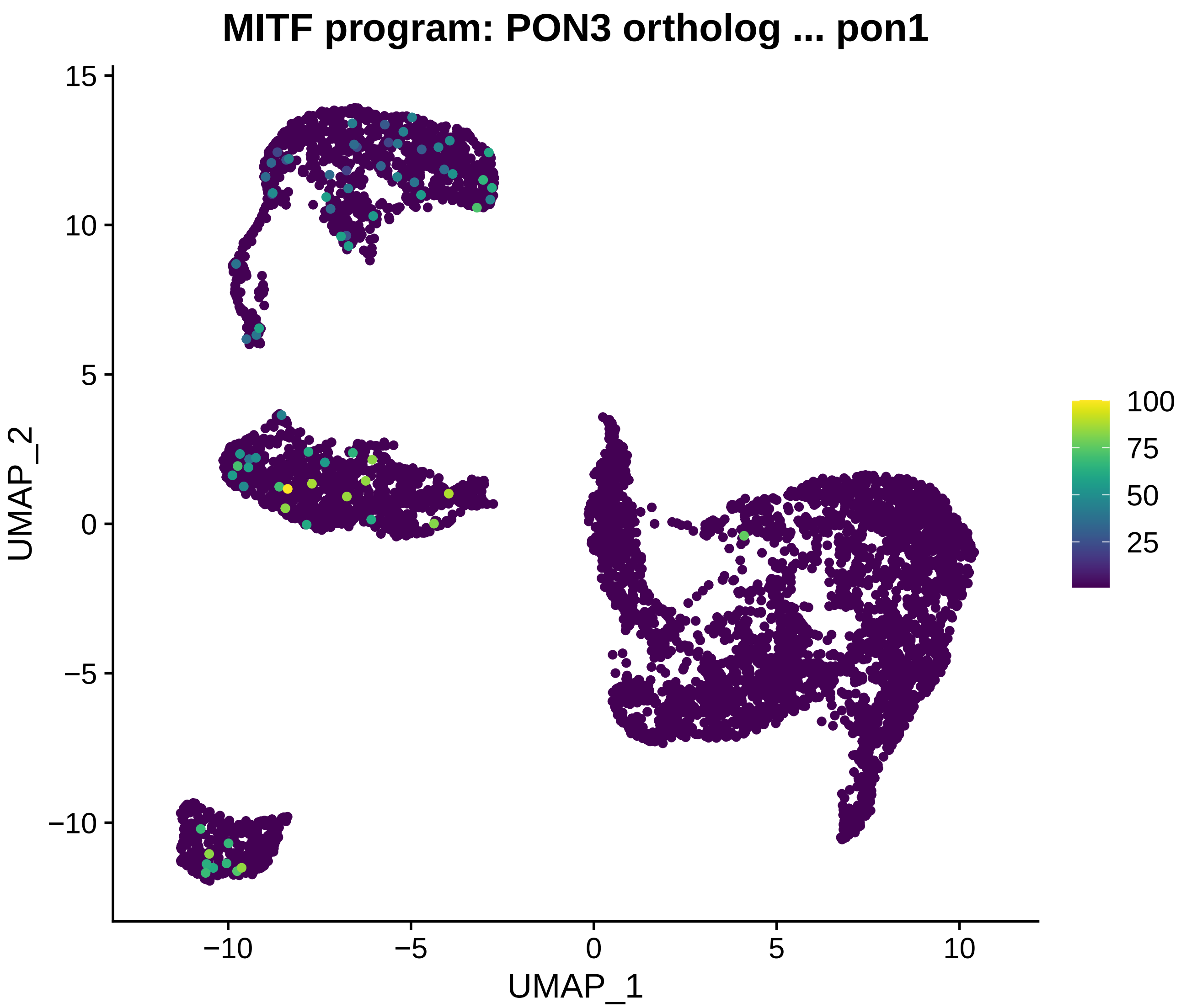
MITF program: PON3 ortholog ... pon1
UMAP_1
UMAP_2
−10
−5
0
5
10
15
100
75
50
25

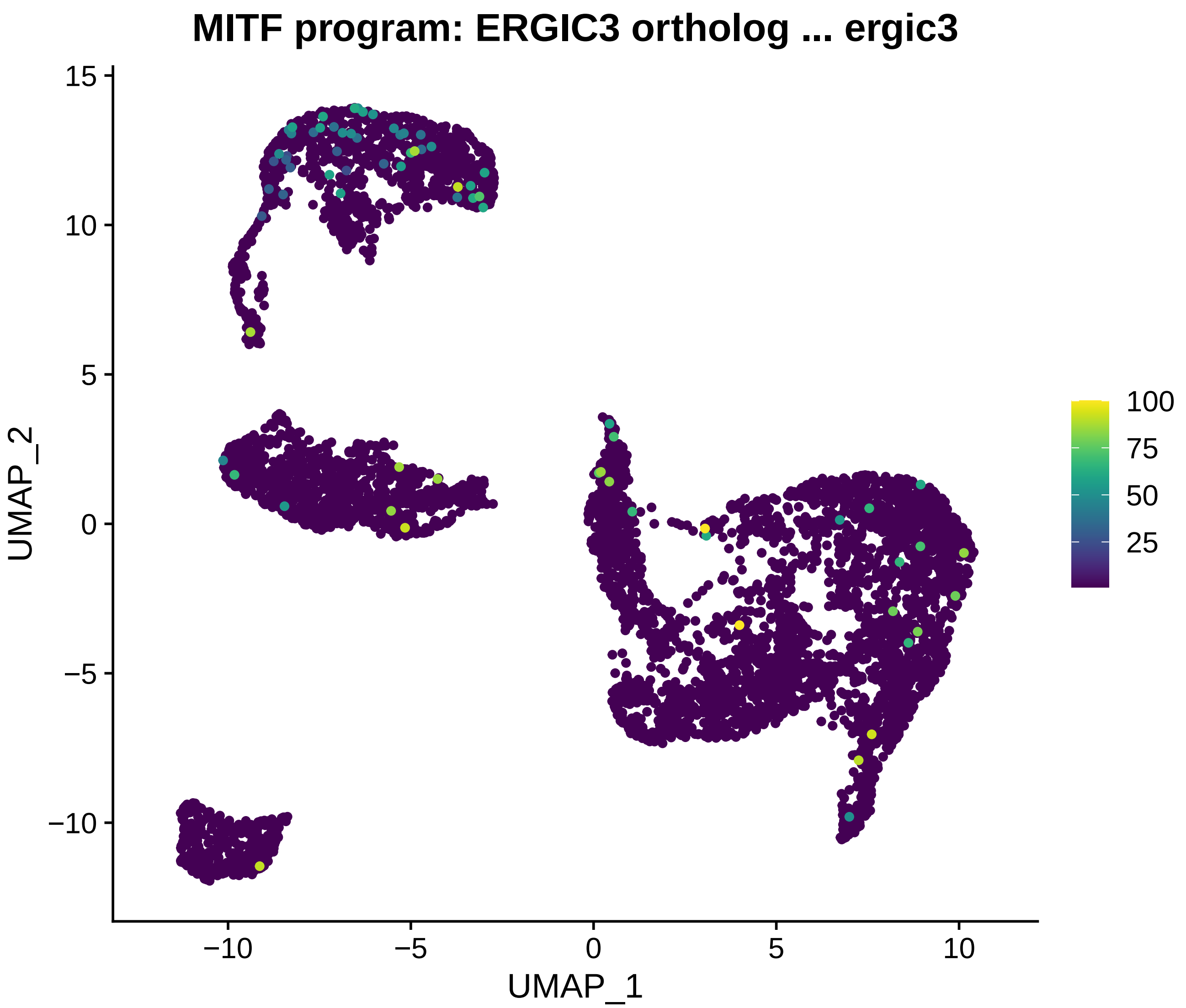

MITF program: ERGIC3 ortholog ... ergic3
UMAP_2
UMAP_1
15
10
5
0
−5
−10
−10
−5
0
5
10
100
75
50
25

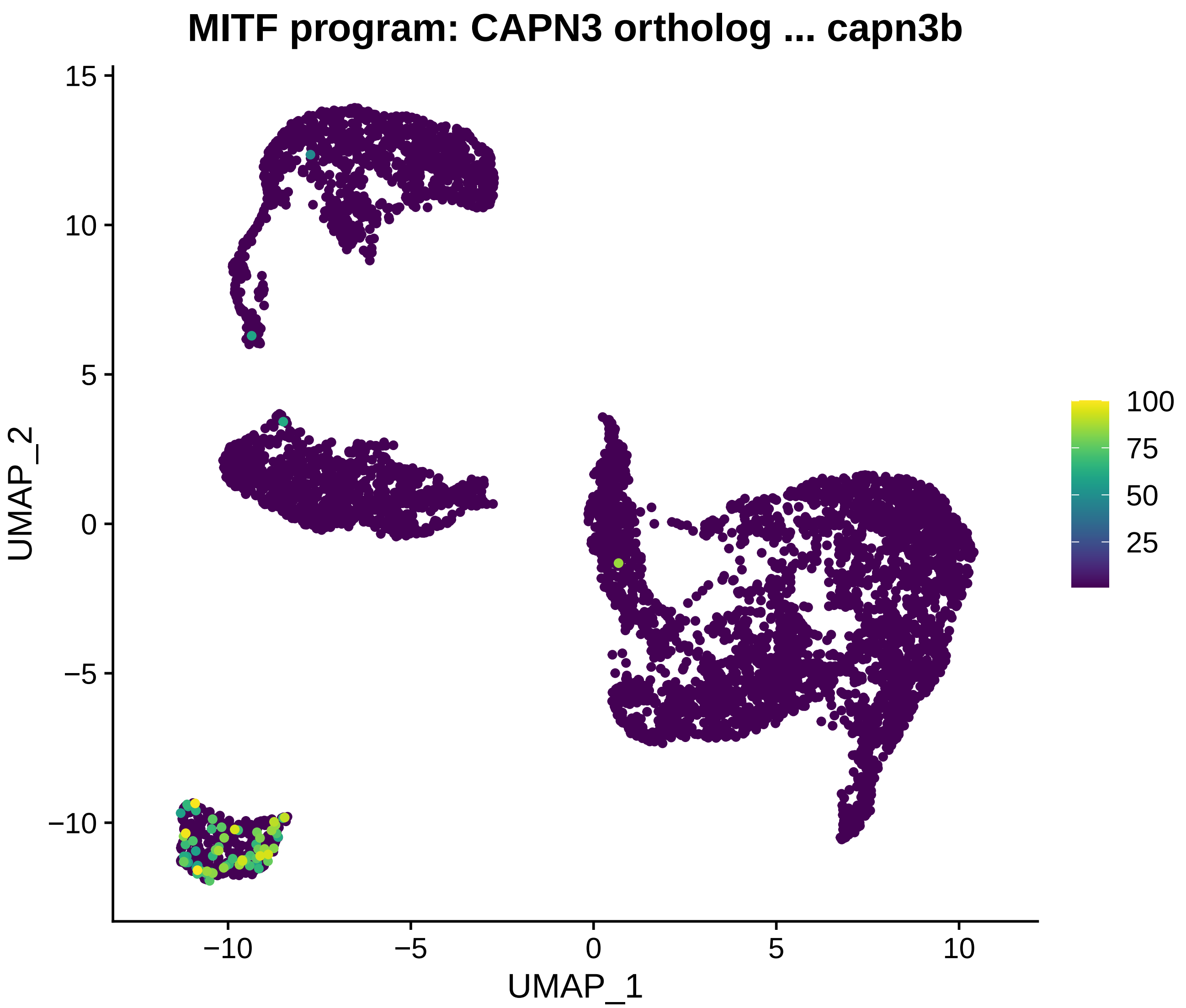

MITF program: CAPN3 ortholog ... capn3b
UMAP_2
UMAP_1
15
10
5
0
−5
−10
−10
−5
0
5
10
100
75
50
25

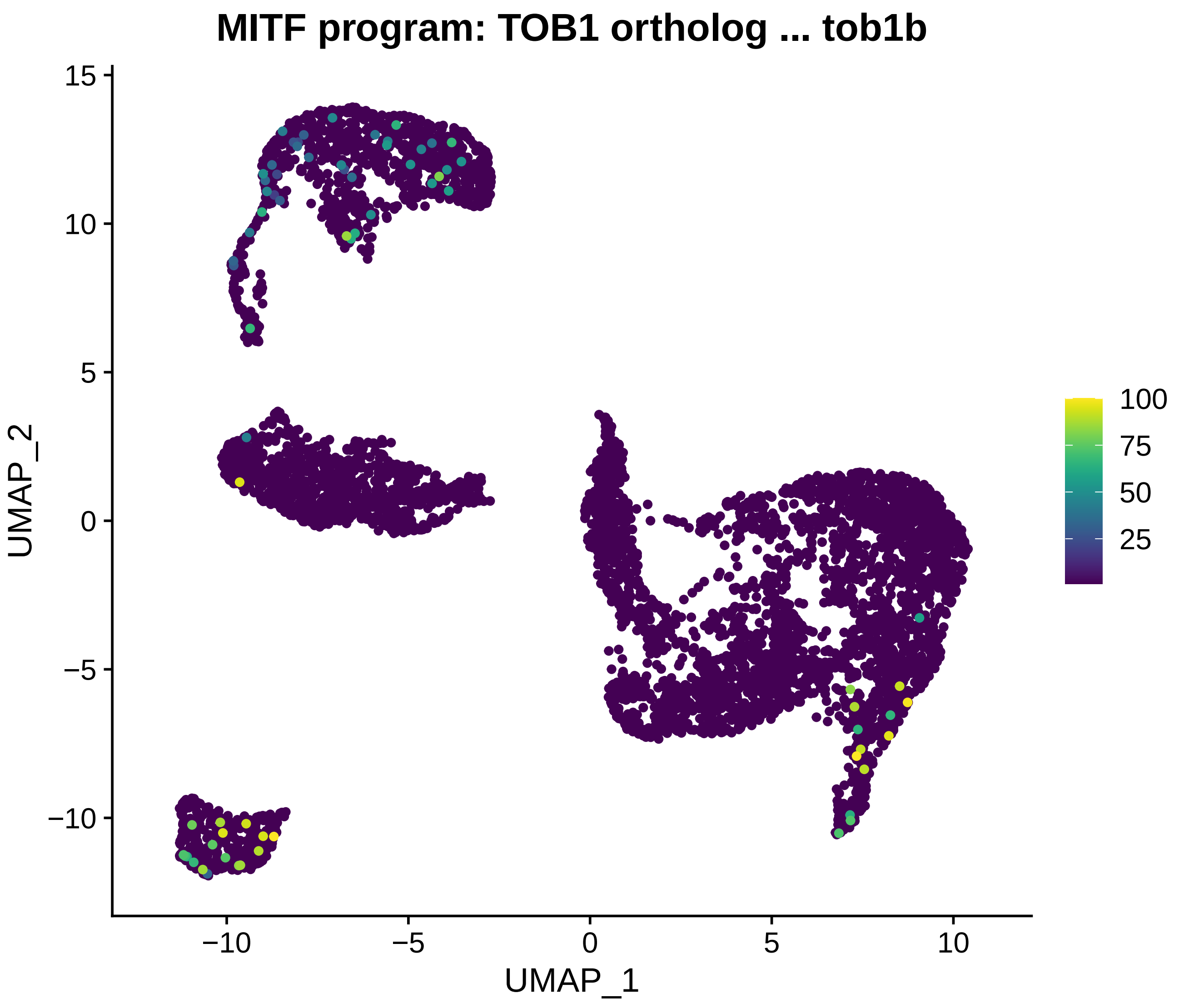
MITF program: TOB1 ortholog ... tob1b
UMAP_2
UMAP_1
15
10
5
0
−5
−10
−10
−5
0
5
10
100
75
50
25

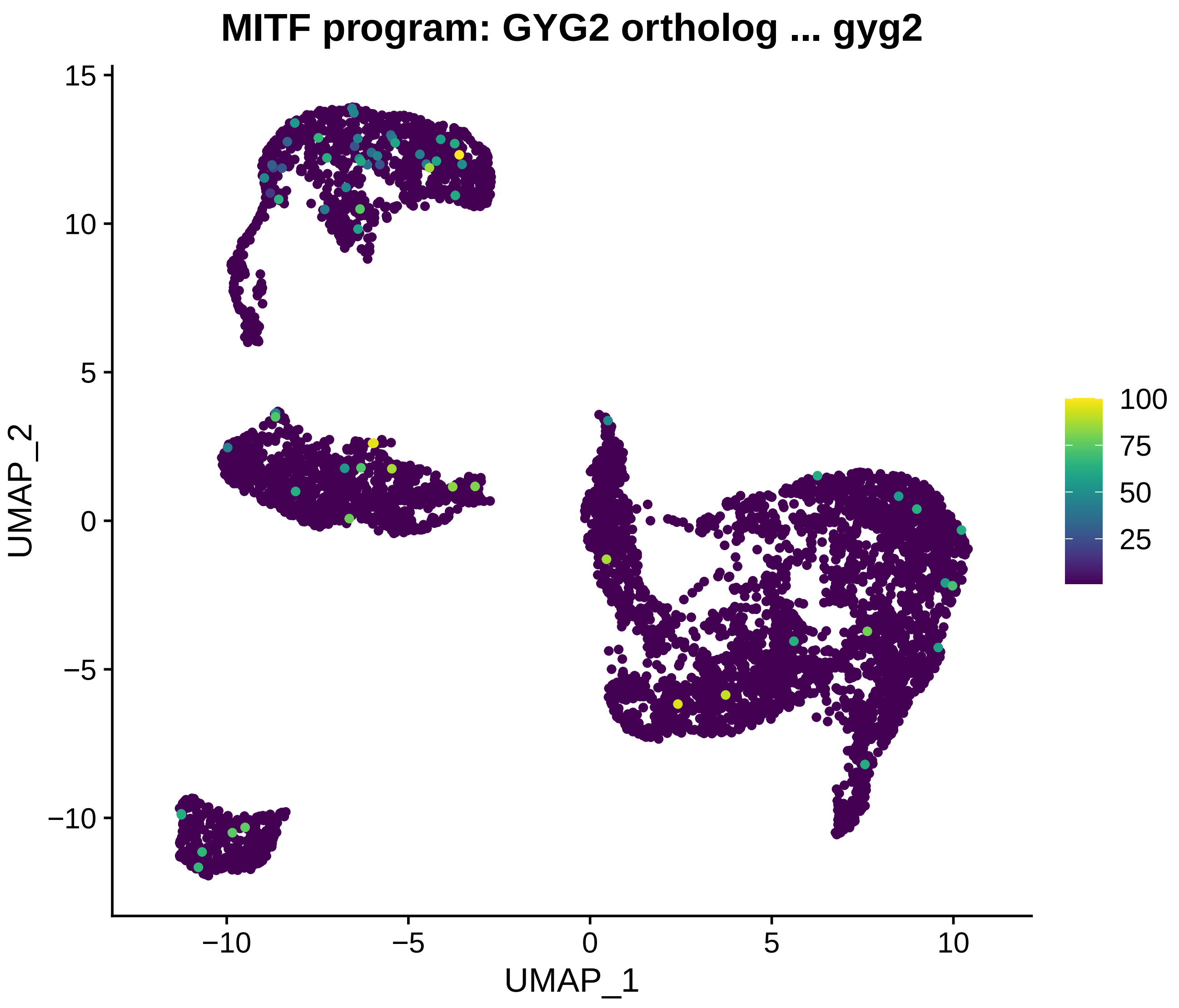
MITF program: GYG2 ortholog ... gyg2
UMAP_2
UMAP_1
15
10
5
0
−5
−10
−10
−5
0
5
10
100
75
50
25

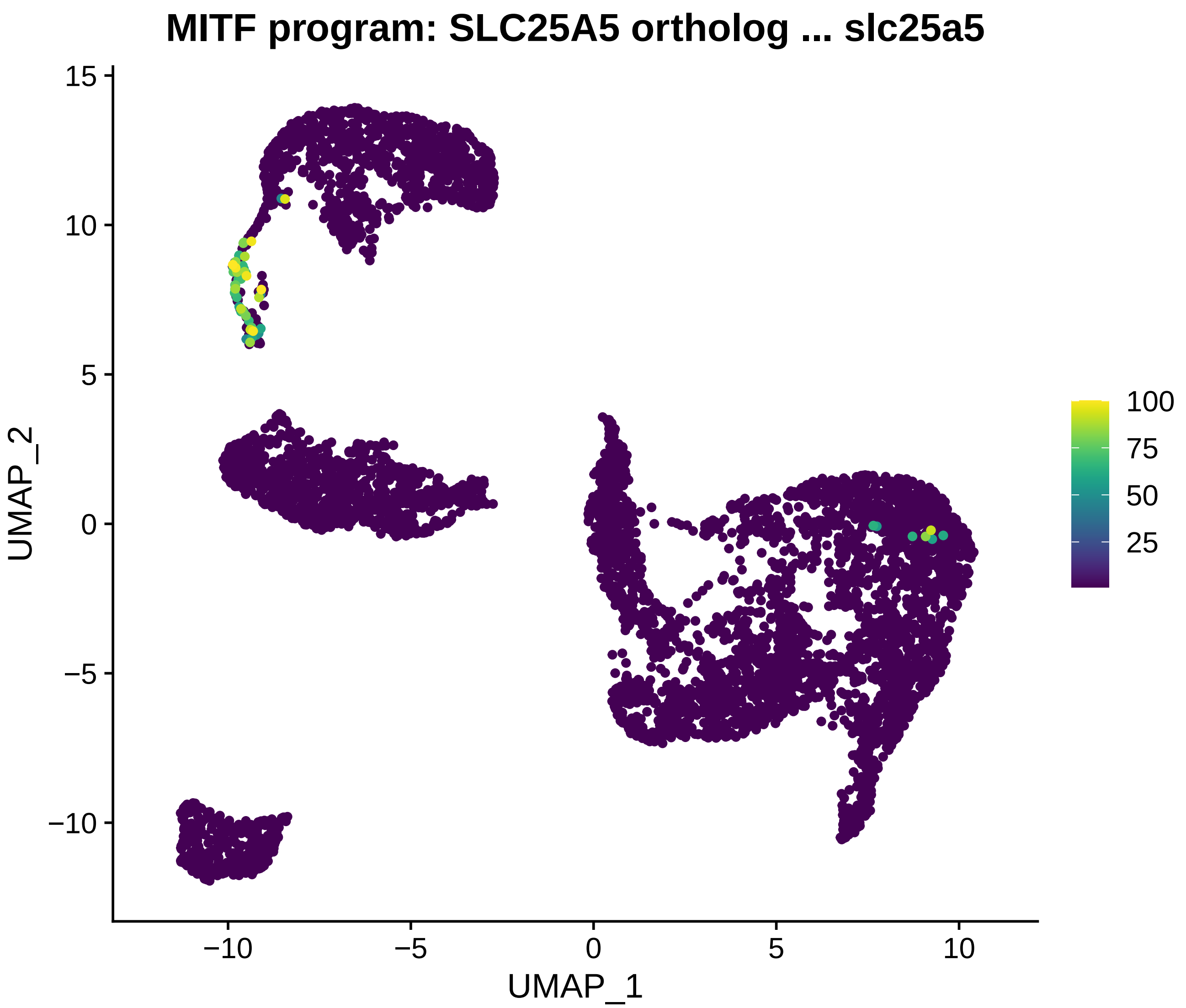
MITF program: SLC25A5 ortholog ... slc25a5
UMAP_2
UMAP_1
15
10
5
0
−5
−10
−10
−5
0
5
10
100
75
50
25

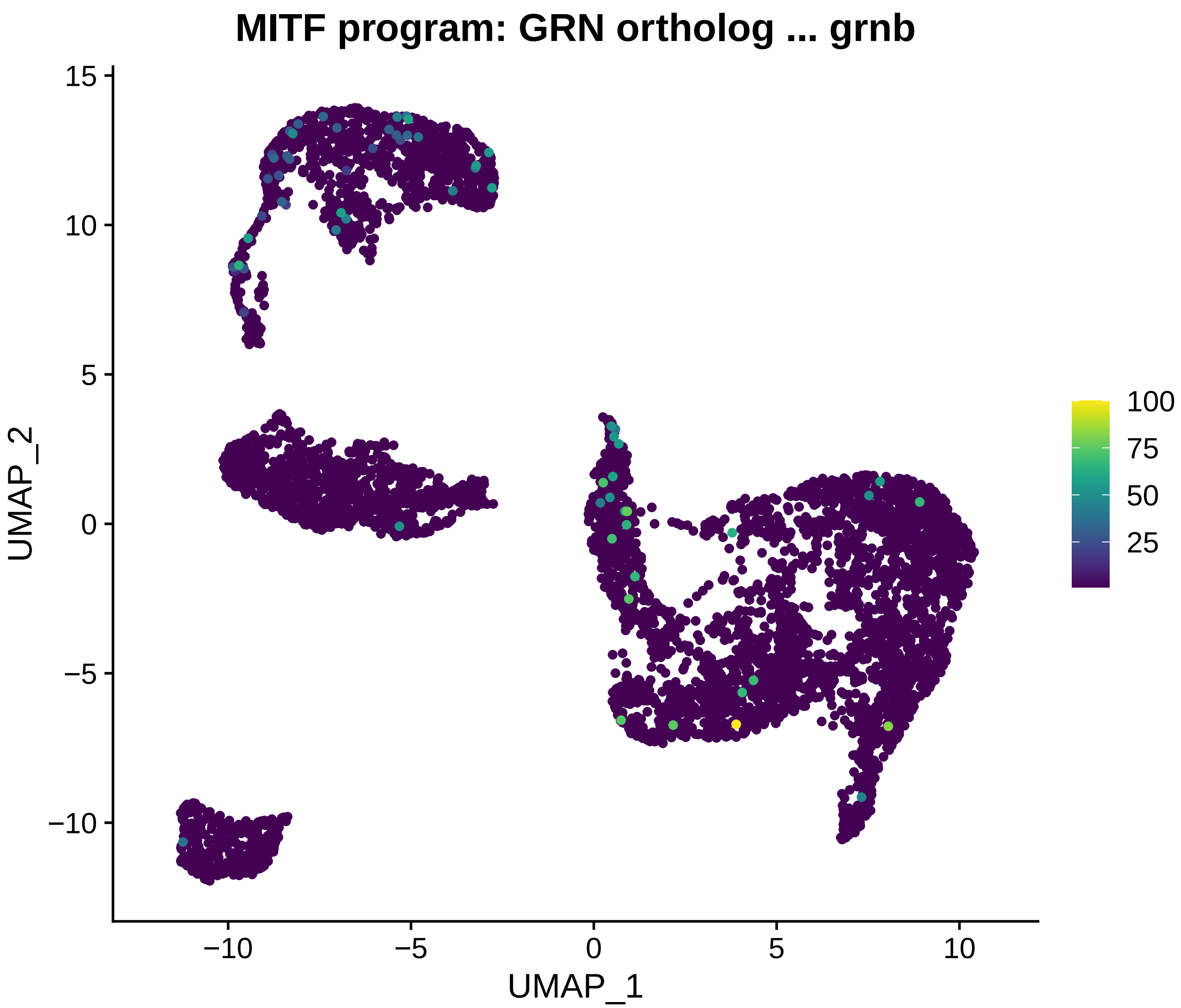

MITF program: GRN ortholog ... grnb
UMAP_2
UMAP_1
15
10
5
0
−5
−10
−10
−5
0
5
10
100
75
50
25

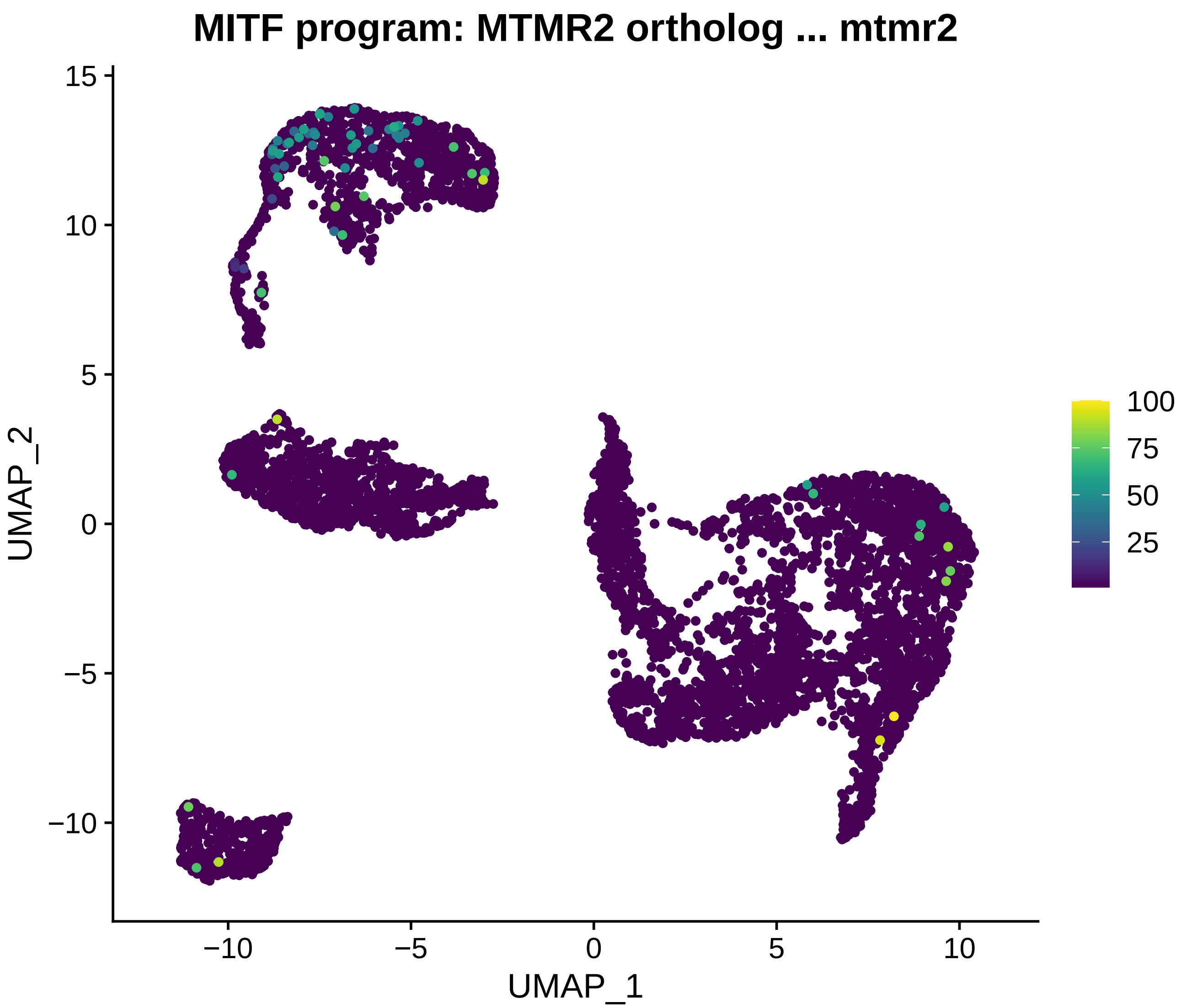
MITF program: MTMR2 ortholog ... mtmr2
UMAP_2
UMAP_1
15
10
5
0
−5
−10
−10
−5
0
5
10
100
75
50
25

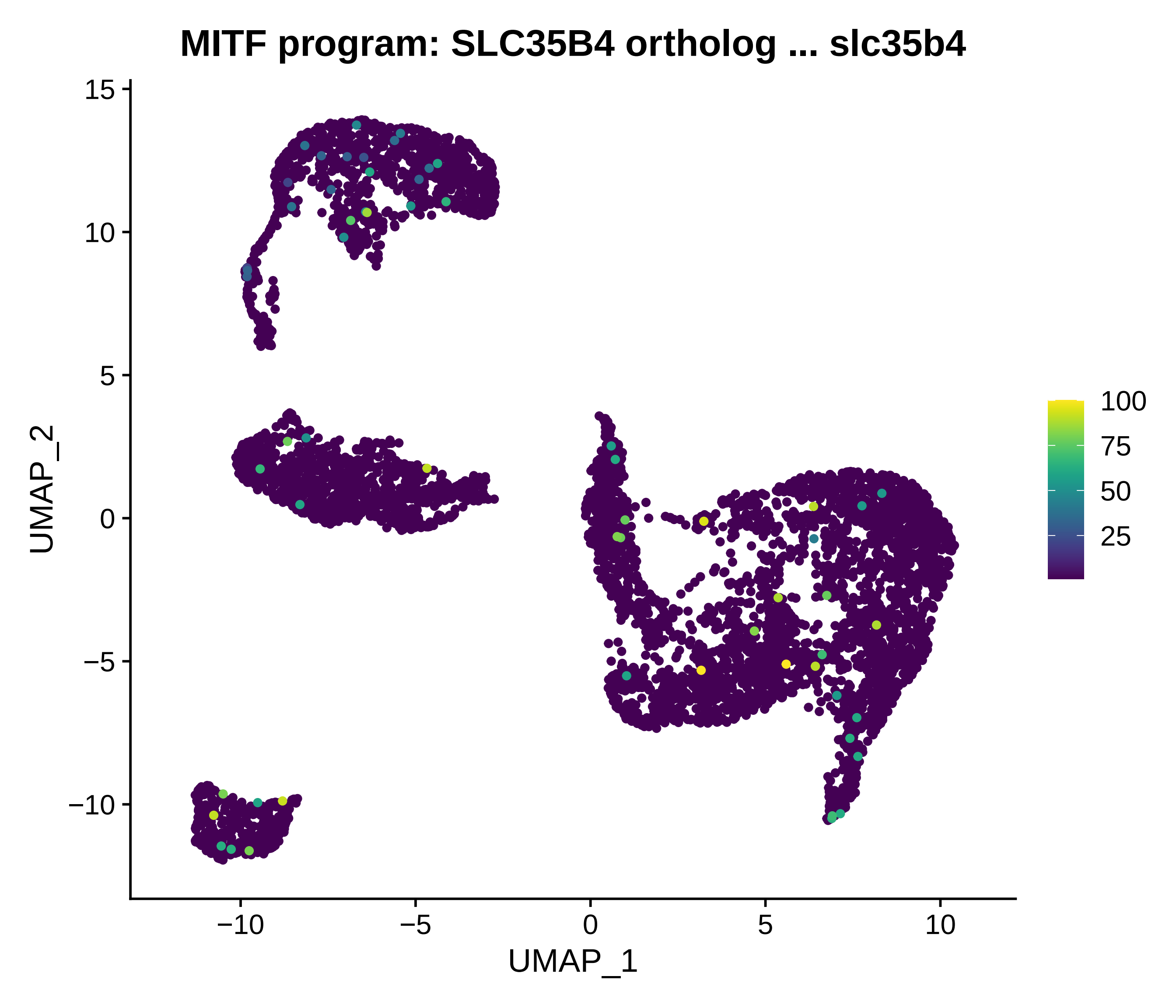
MITF program: SLC35B4 ortholog ... slc35b4
UMAP_2
UMAP_1
15
10
5
0
−5
−10
−10
−5
0
5
10
100
75
50
25

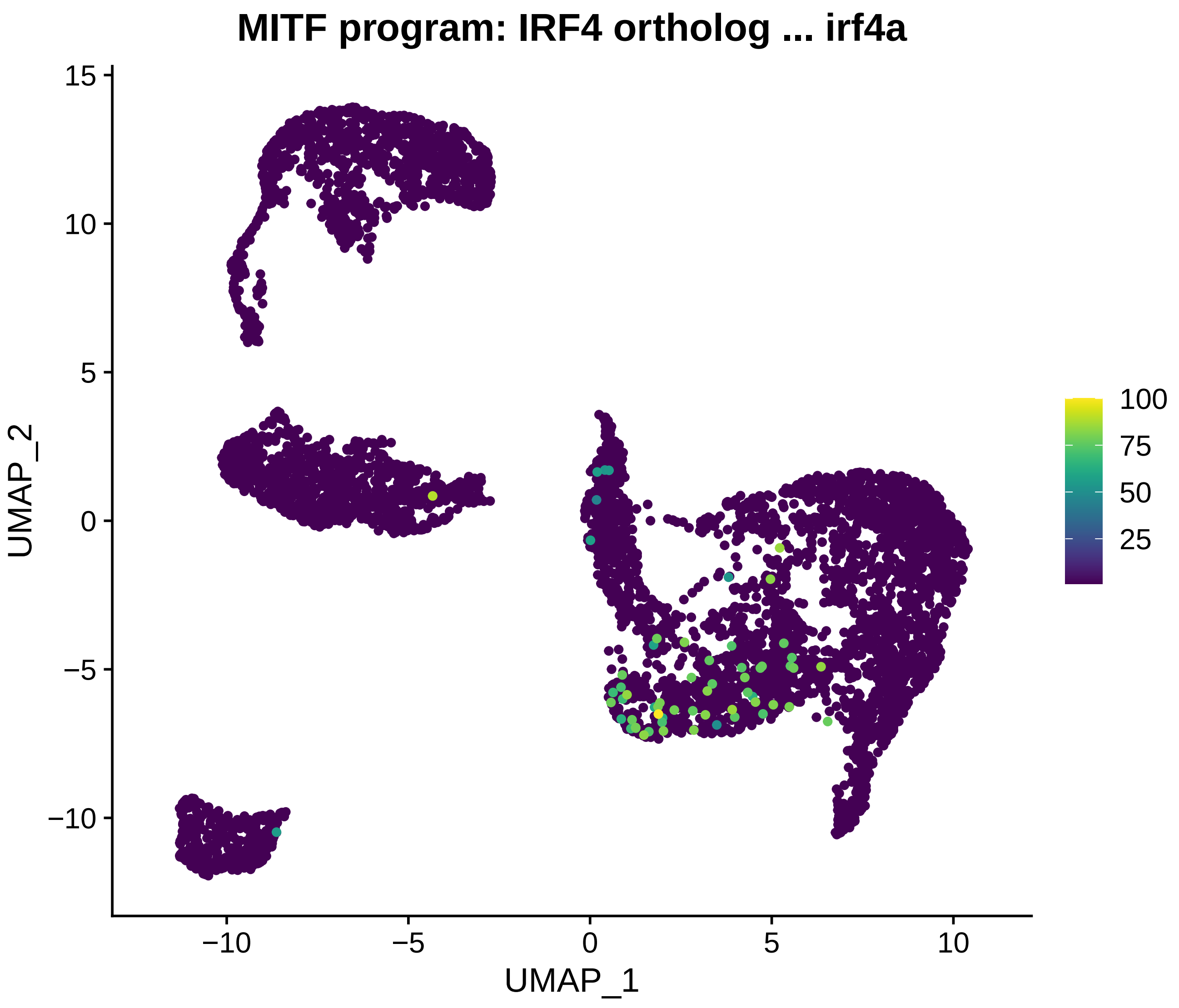
MITF program: IRF4 ortholog ... irf4a
UMAP_2
UMAP_1
15
10
5
0
−5
−10
−10
−5
0
5
10
100
75
50
25

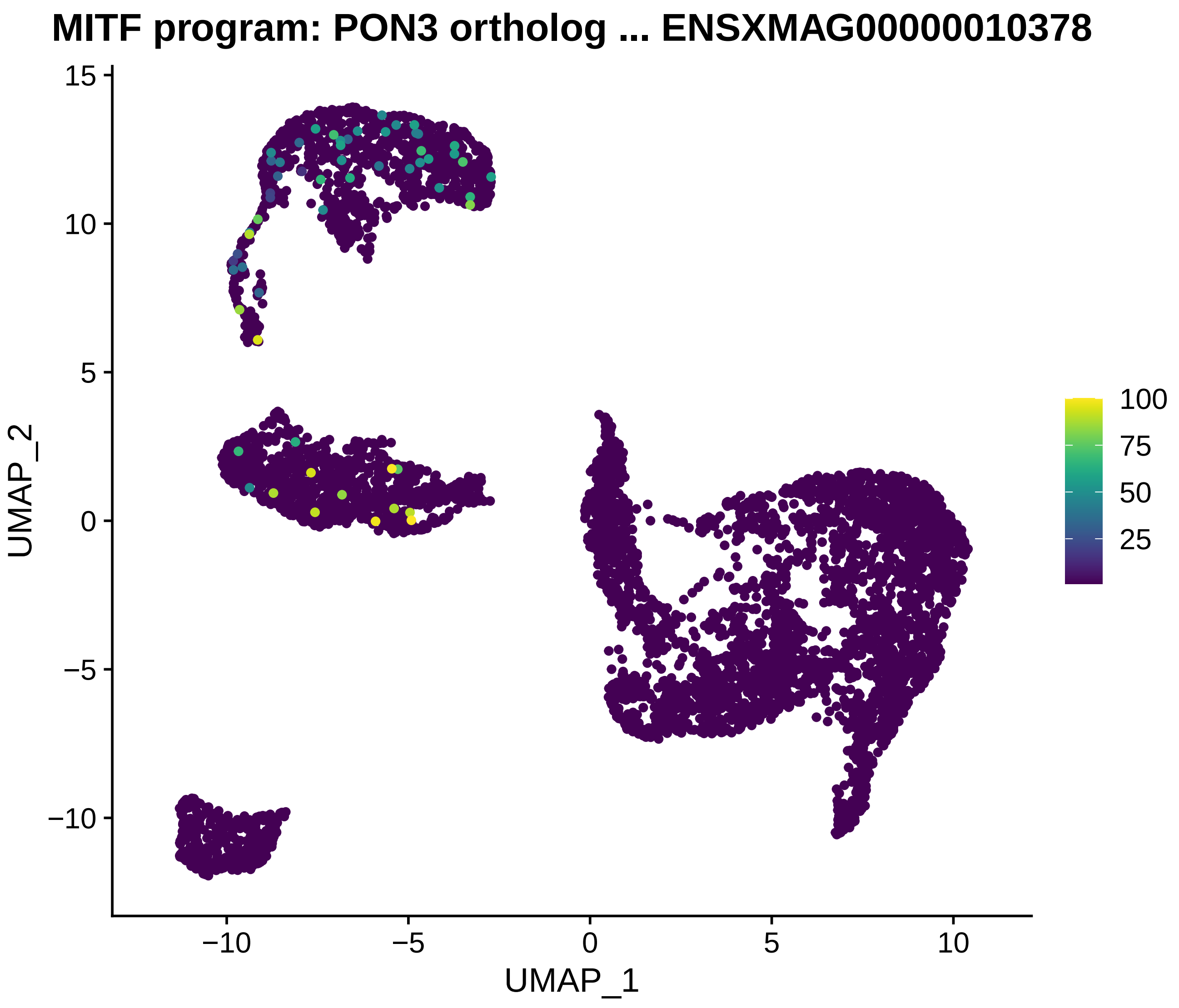
MITF program: PON3 ortholog ... ENSXMAG00000010378
UMAP_2
UMAP_1
15
10
5
0
−5
−10
−10
−5
0
5
10
100
75
50
25

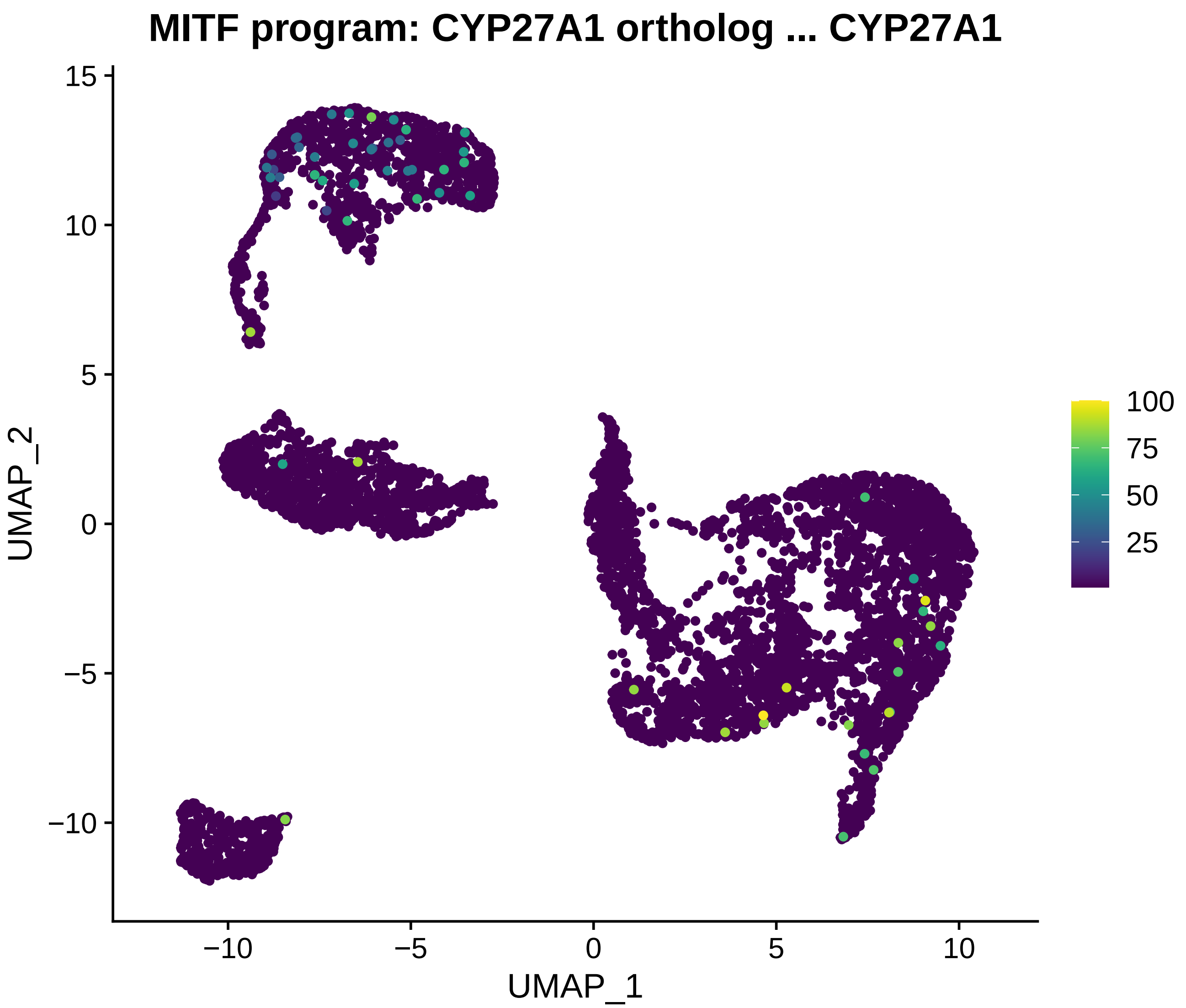
MITF program: CYP27A1 ortholog ... CYP27A1
UMAP_2
UMAP_1
15
10
5
0
−5
−10
−10
−5
0
5
10
100
75
50
25

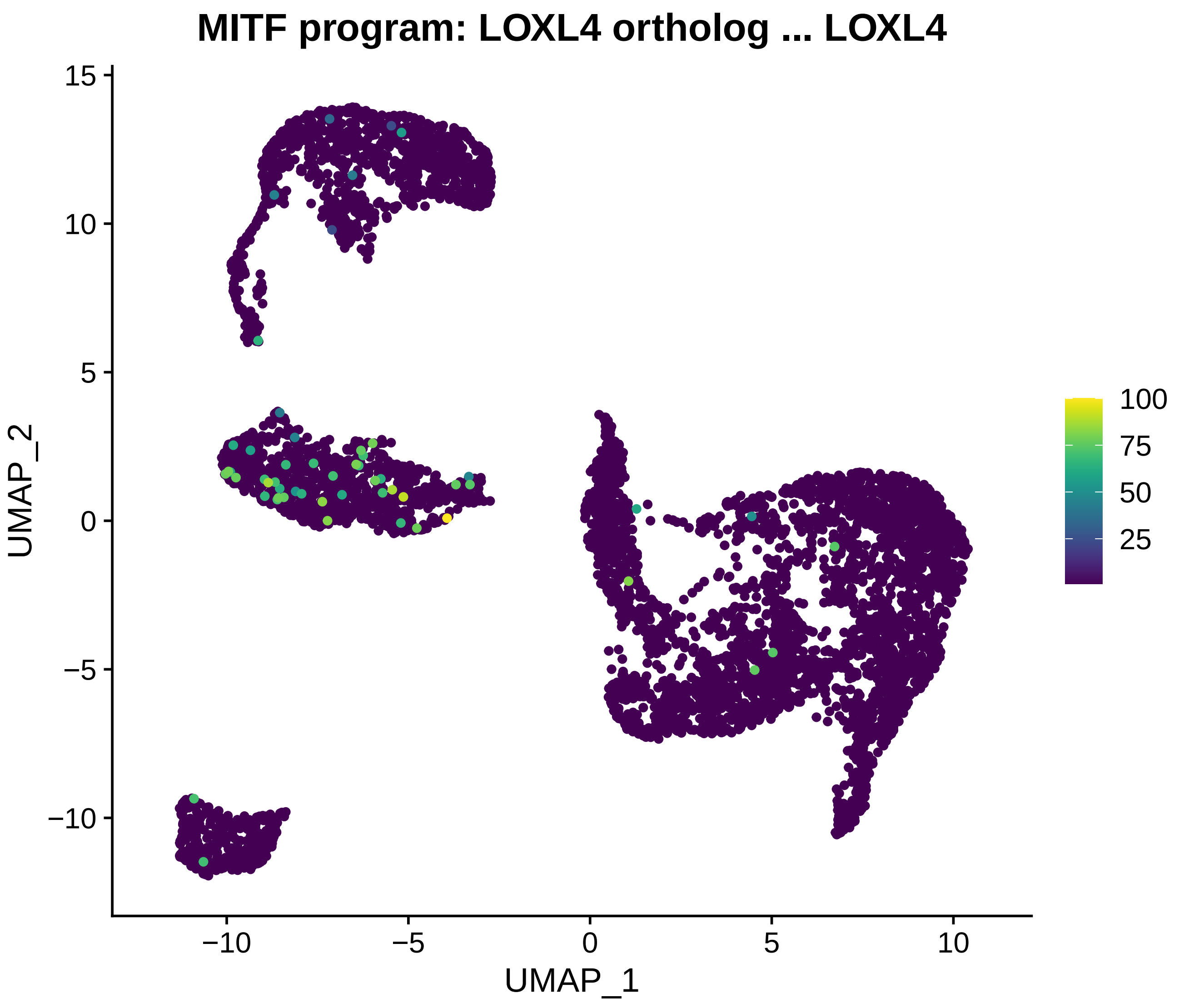
MITF program: LOXL4 ortholog ... LOXL4
UMAP_2
UMAP_1
15
10
5
0
−5
−10
−10
−5
0
5
10
100
75
50
25

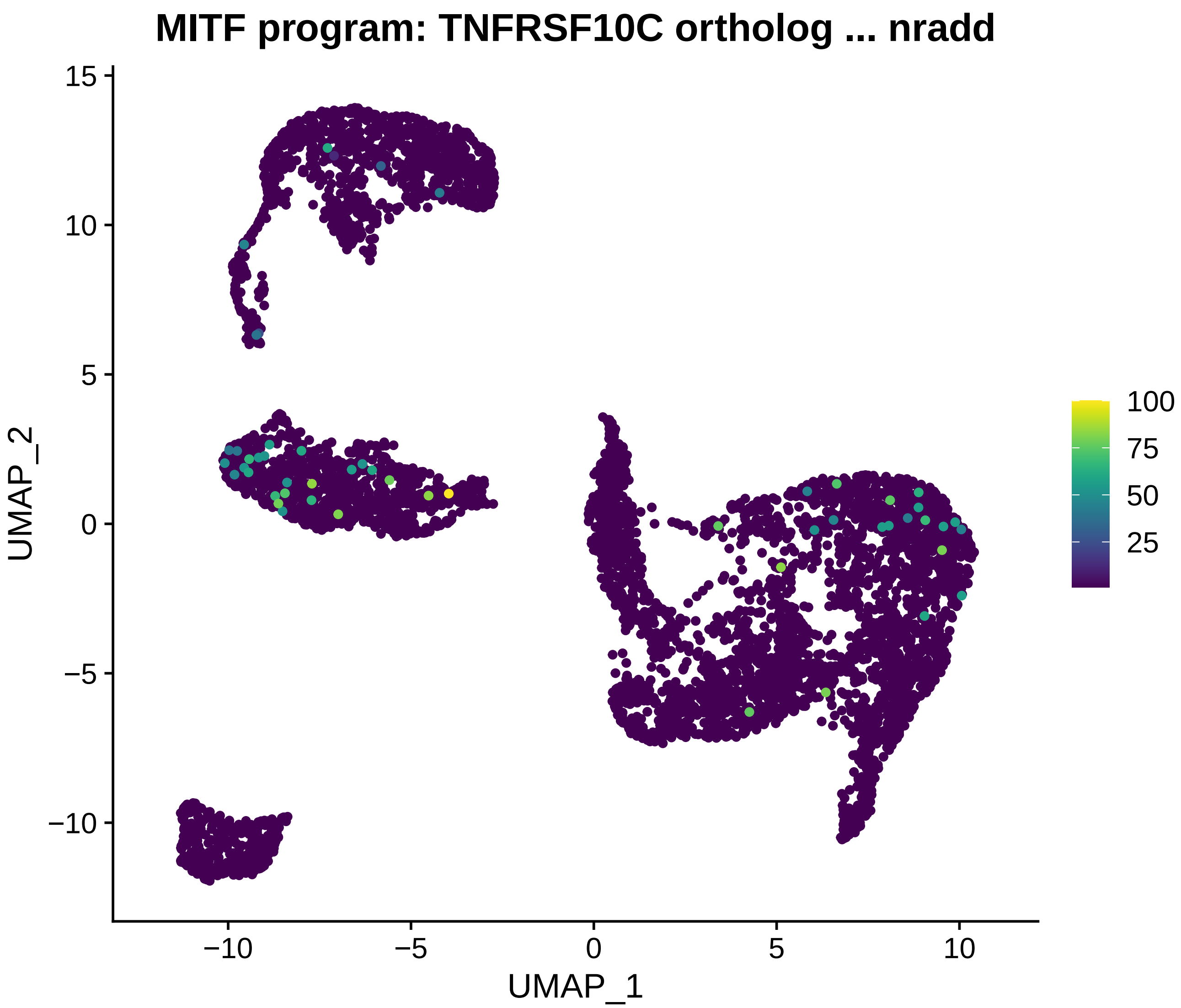
MITF program: TNFRSF10C ortholog ... nradd
UMAP_1
UMAP_2
−10
−5
0
5
10
15
100
75
50
25

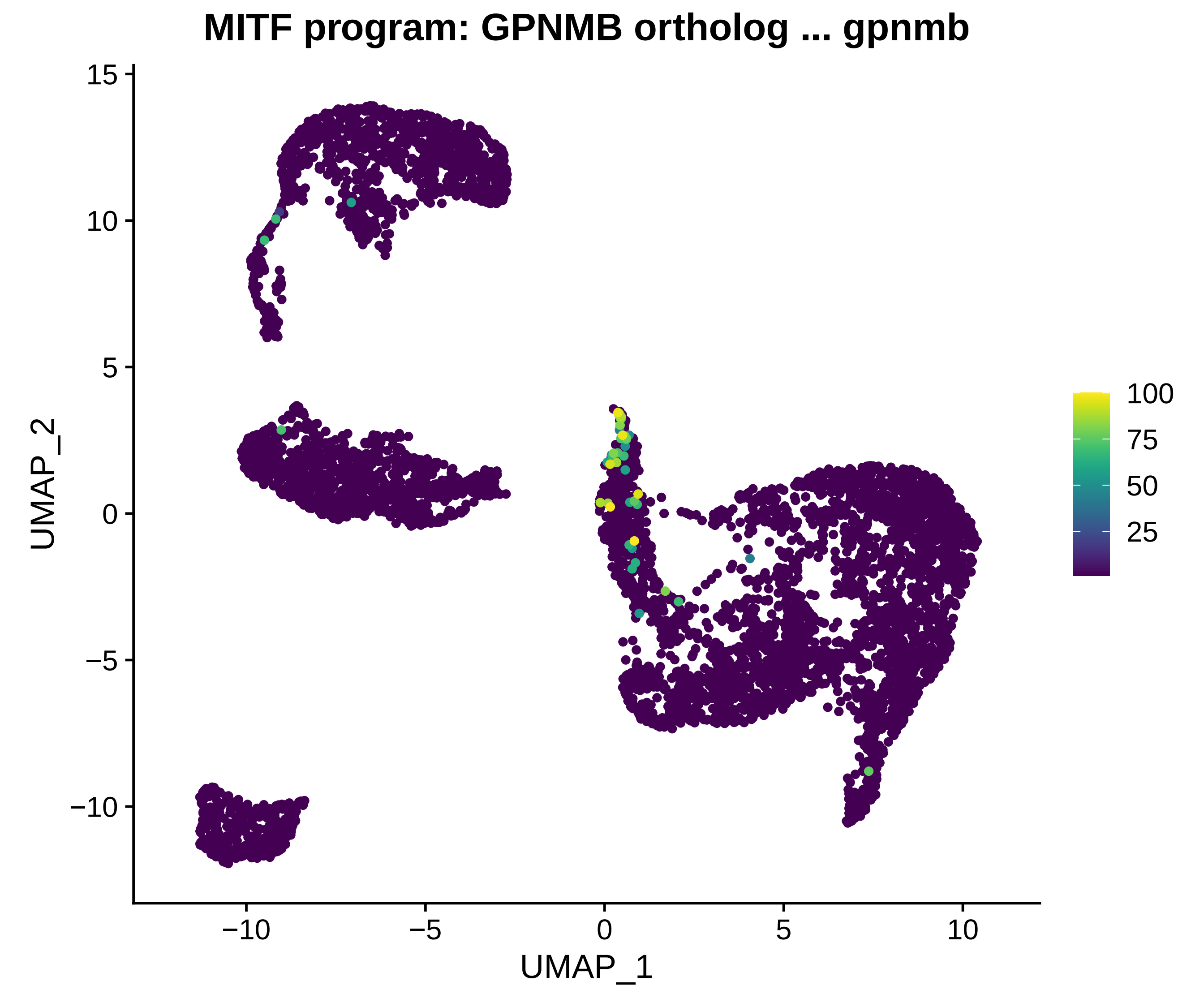
MITF program: GPNMB ortholog ... gpnmb
UMAP_2
UMAP_1
15
10
5
0
−5
−10
−10
−5
0
5
10
100
75
50
25

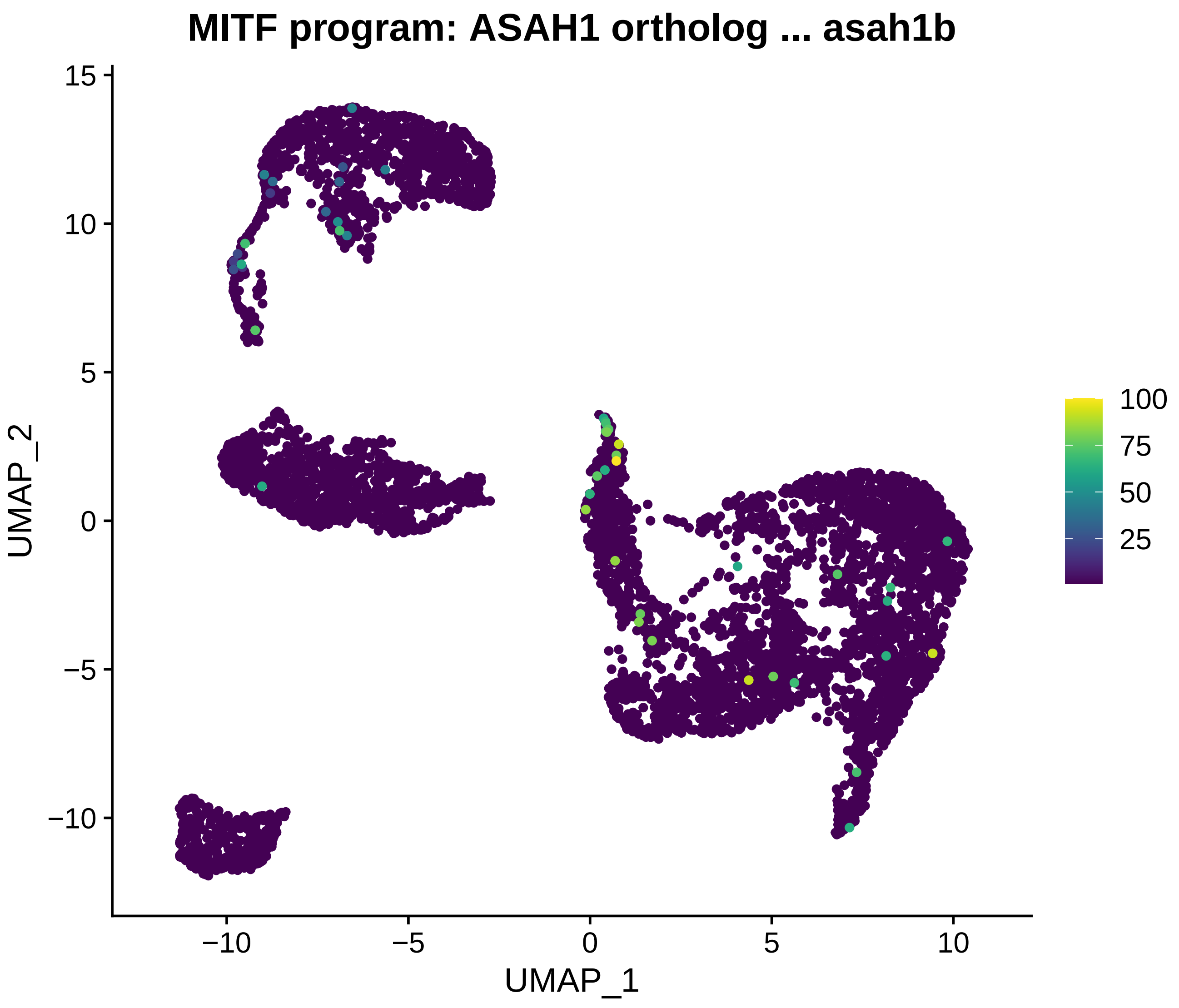
MITF program: ASAH1 ortholog ... asah1b
UMAP_2
UMAP_1
15
10
5
0
−5
−10
−10
−5
0
5
10
100
75
50
25

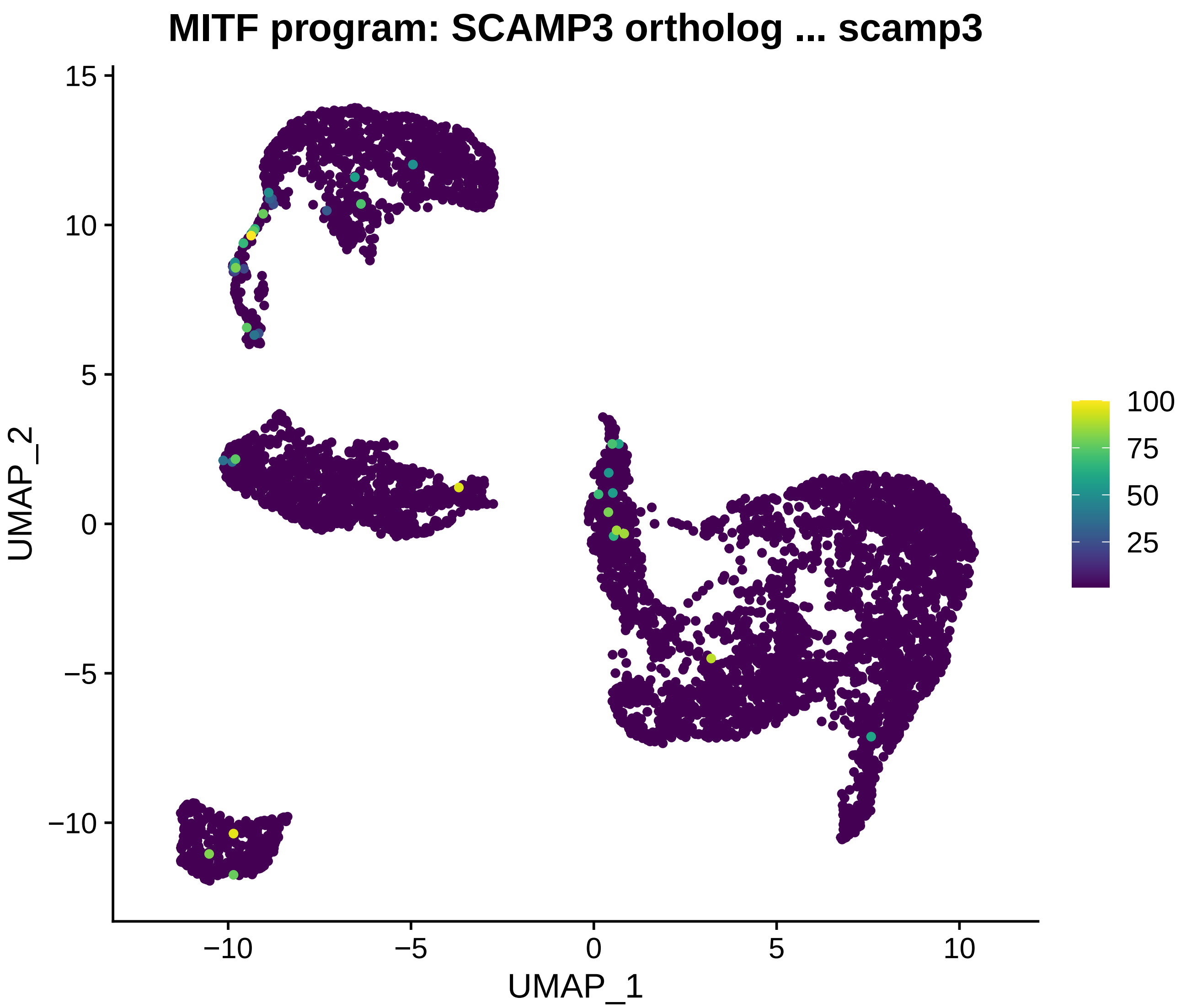
MITF program: SCAMP3 ortholog ... scamp3
UMAP_2
UMAP_1
15
10
5
0
−5
−10
−10
−5
0
5
10
100
75
50
25

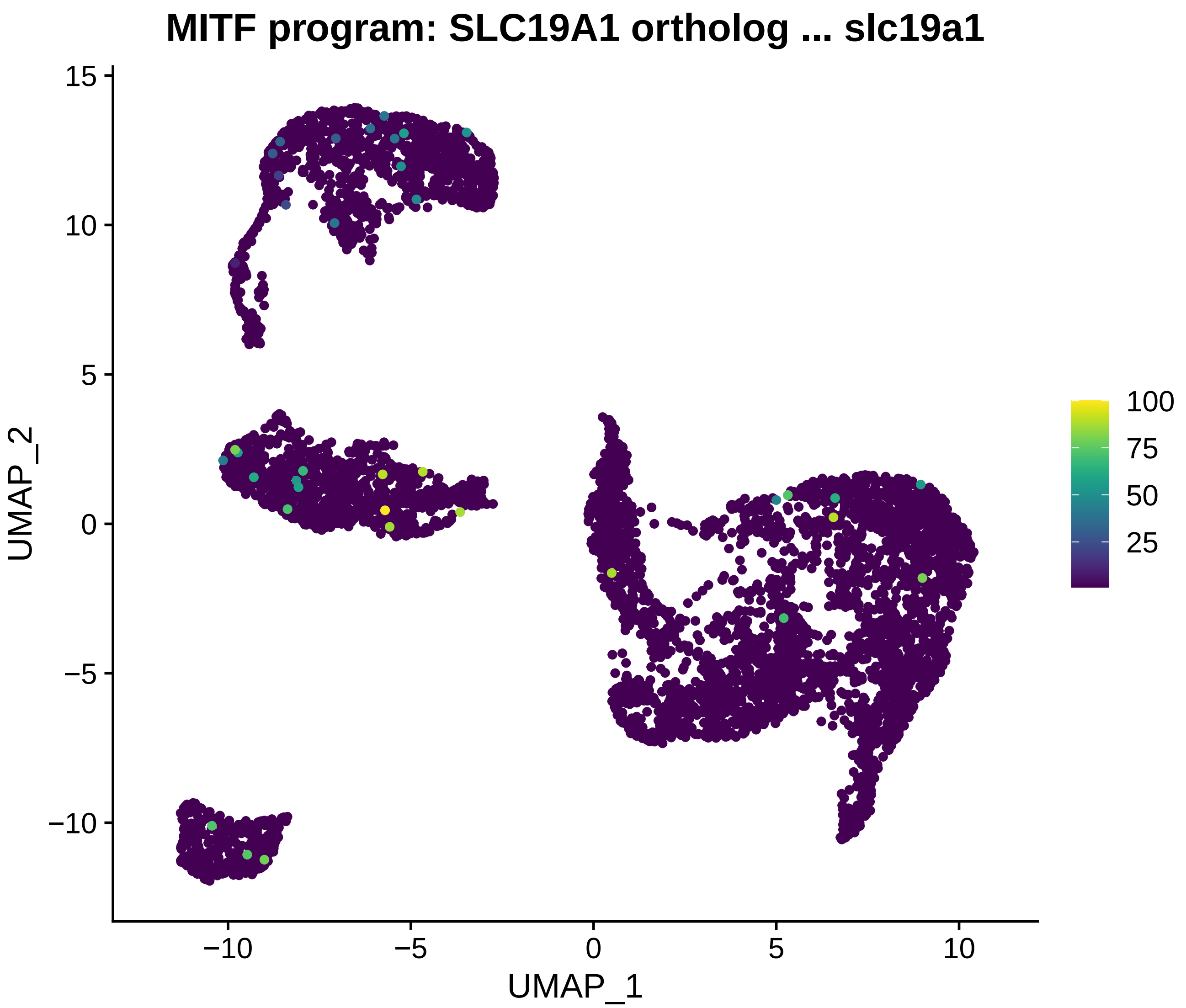
MITF program: SLC19A1 ortholog ... slc19a1
UMAP_1
UMAP_2
15
10
5
0
−5
−10
100
75
50
25

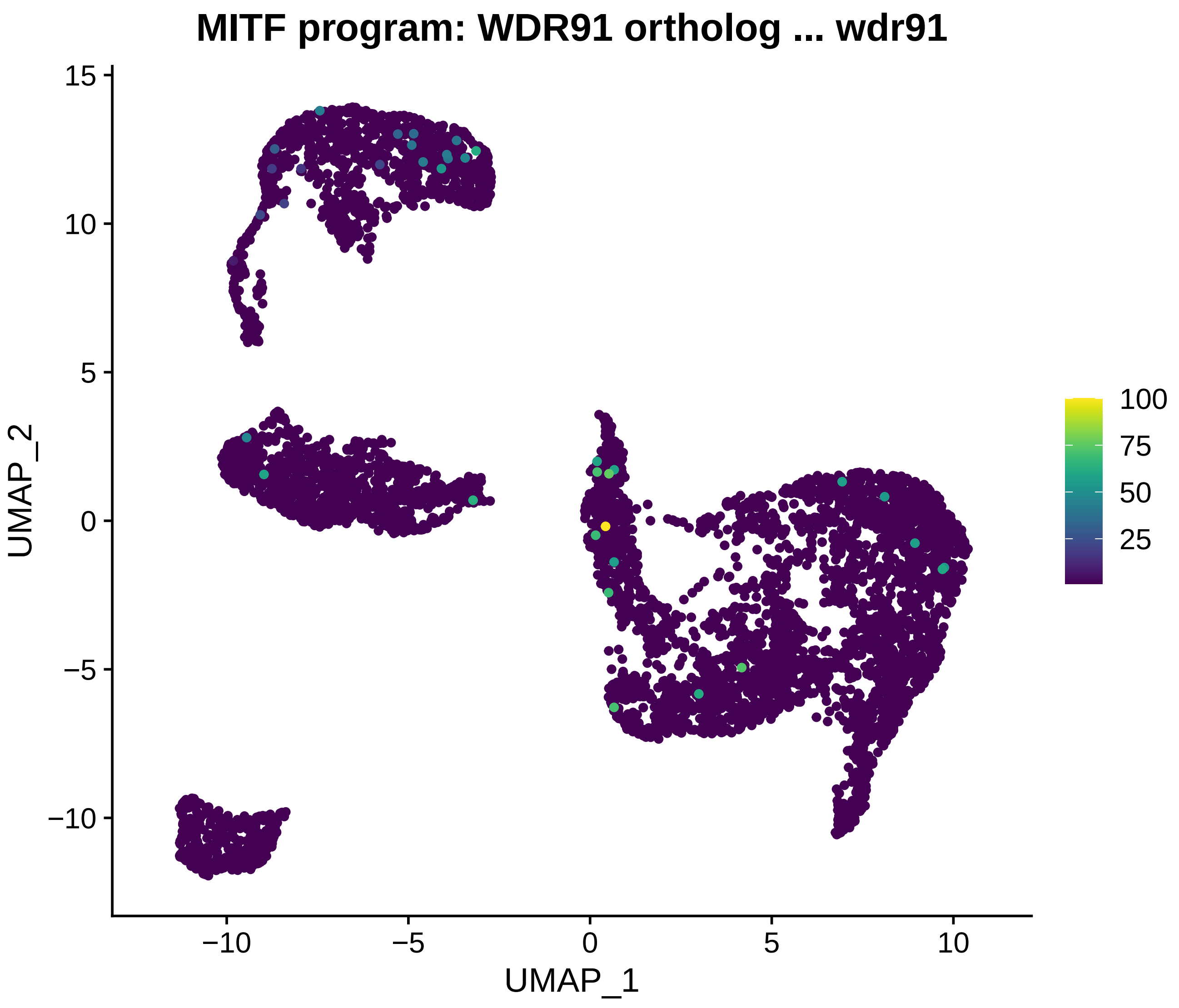
MITF program: WDR91 ortholog ... wdr91
UMAP_2
UMAP_1
15
10
5
0
−5
−10
−10
−5
0
5
10
100
75
50
25

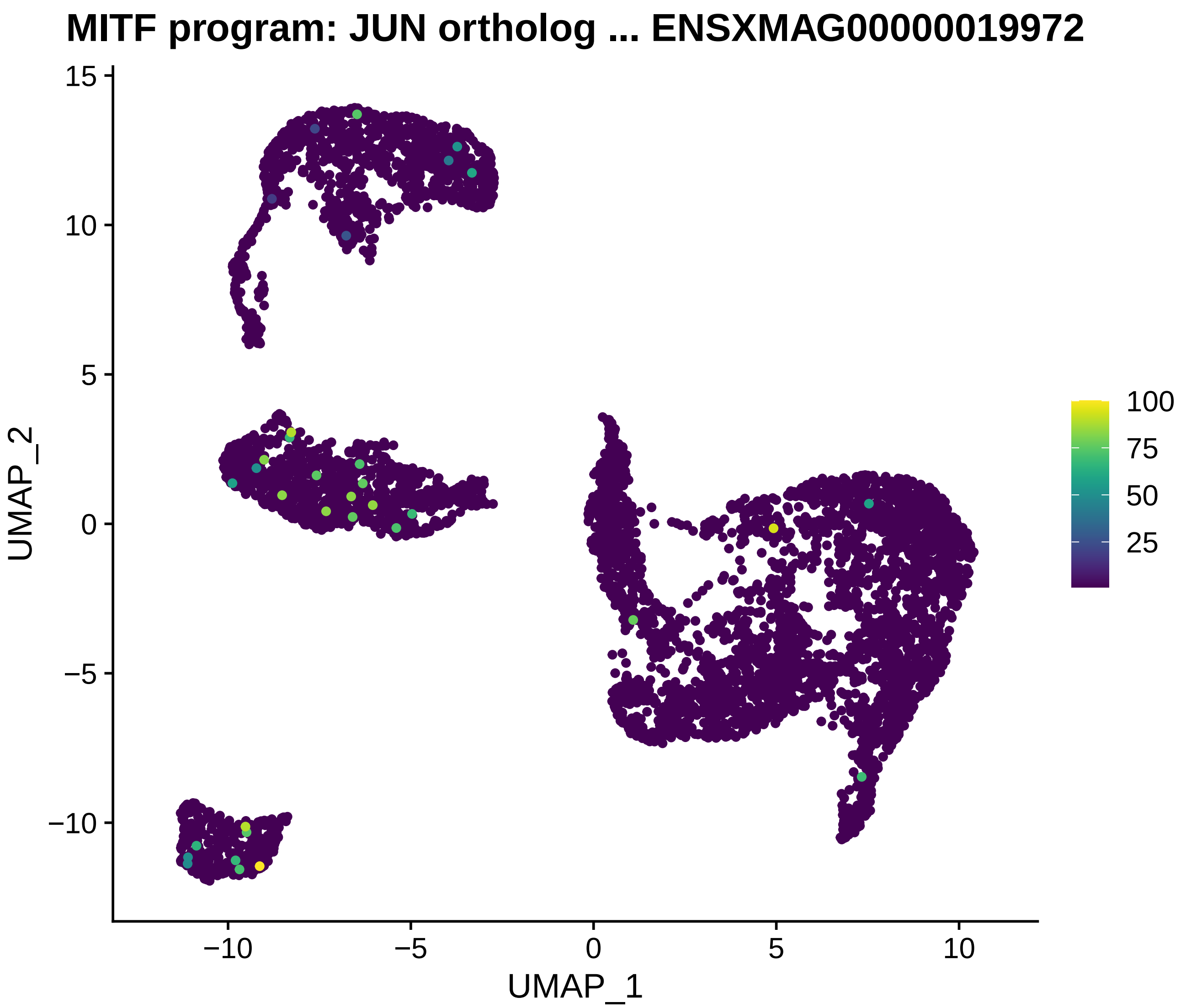
MITF program: JUN ortholog ... ENSXMAG00000019972
UMAP_2
UMAP_1
15
10
5
0
−5
−10
−10
−5
0
5
10
100
75
50
25

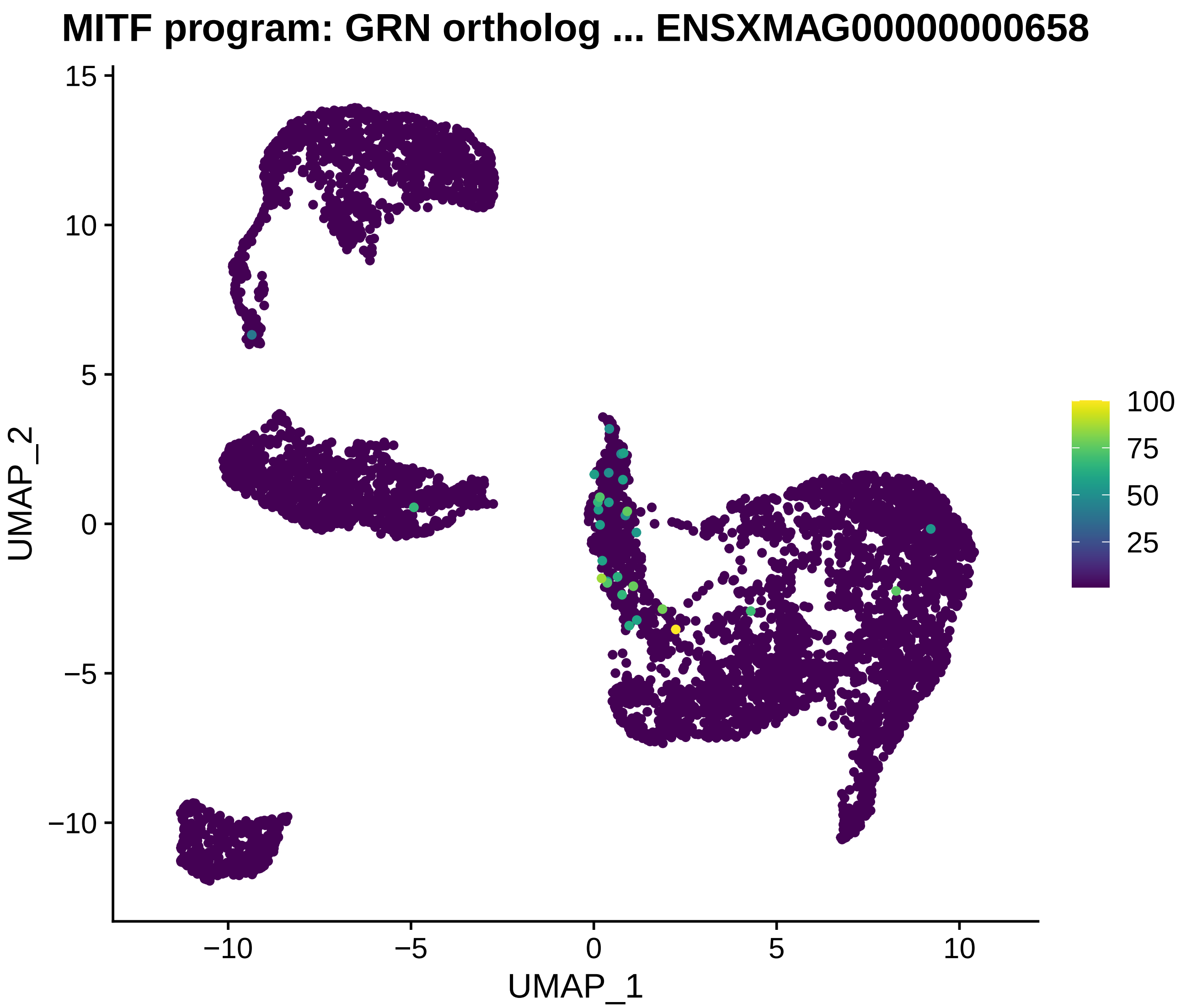
MITF program: GRN ortholog ... ENSXMAG00000000658
UMAP_2
UMAP_1
15
10
5
0
−5
−10
−10
−5
0
5
10
100
75
50
25

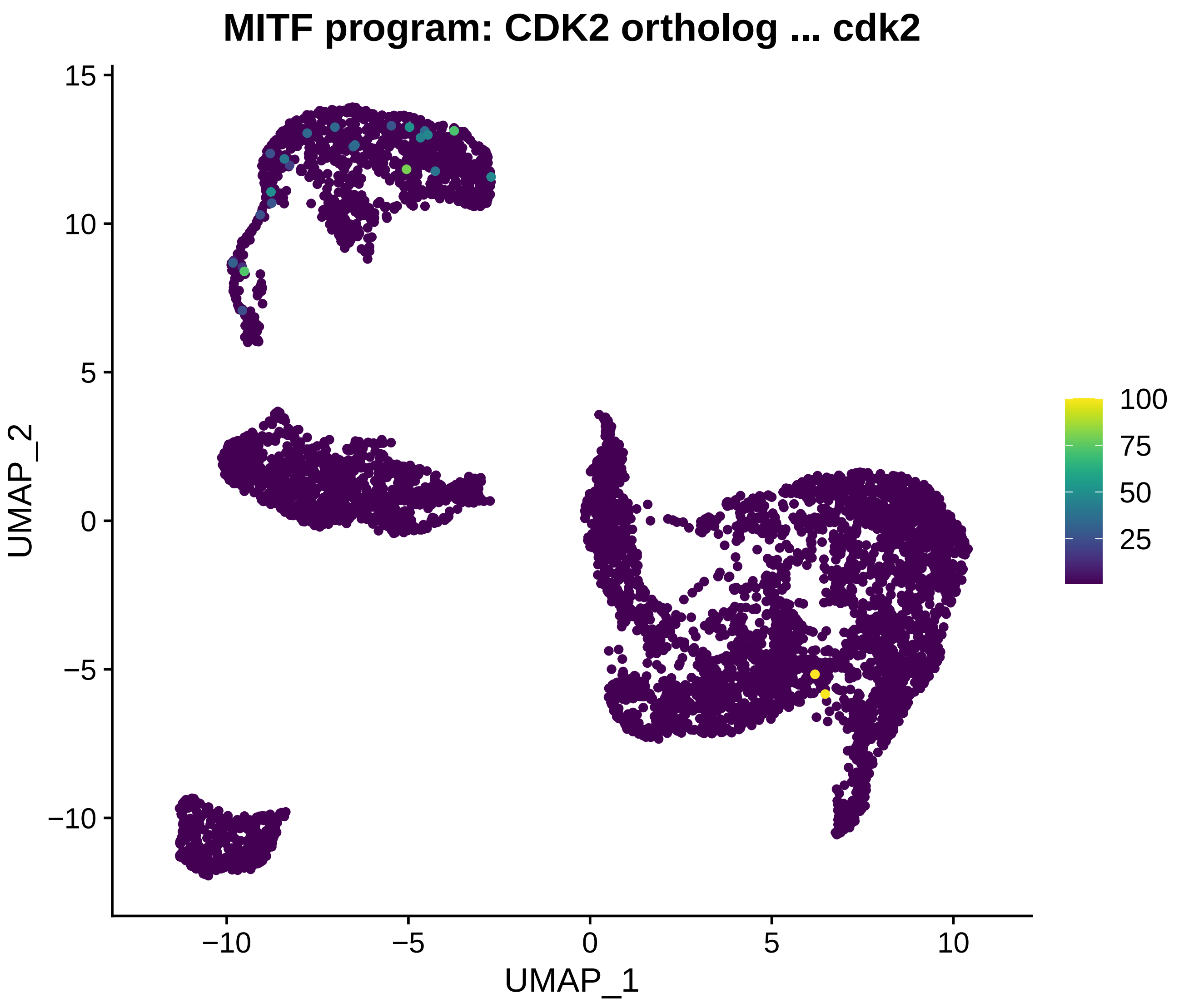
MITF program: CDK2 ortholog ... cdk2
UMAP_2
UMAP_1
15
10
5
0
−5
−10
−10
−5
0
5
10
100
75
50
25

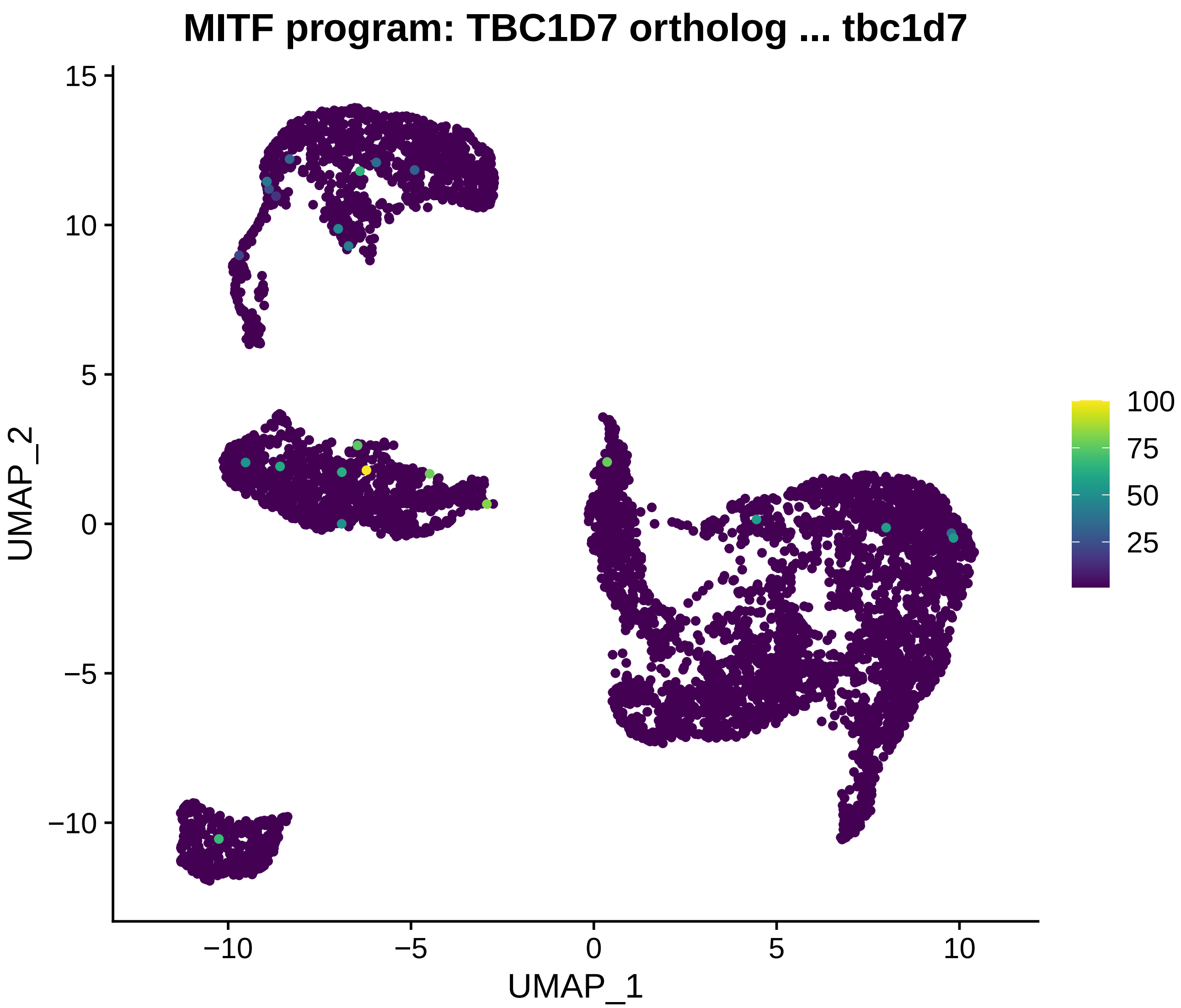
MITF program: TBC1D7 ortholog ... tbc1d7
UMAP_2
UMAP_1
15
10
5
0
−5
−10
−10
−5
0
5
10
100
75
50
25

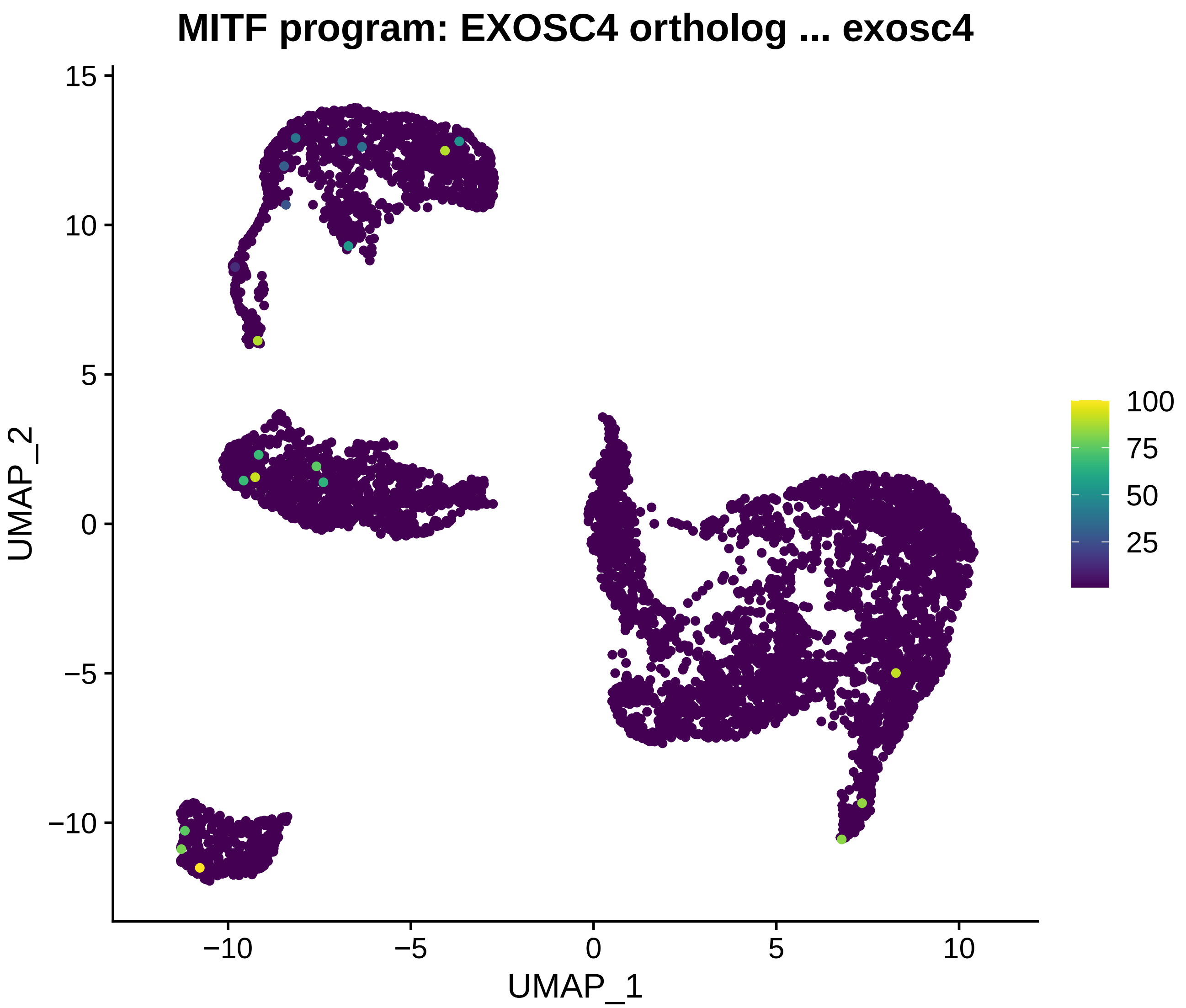
MITF program: EXOSC4 ortholog ... exosc4
UMAP_2
UMAP_1
15
10
5
0
−5
−10
−10
−5
0
5
10
100
75
50
25

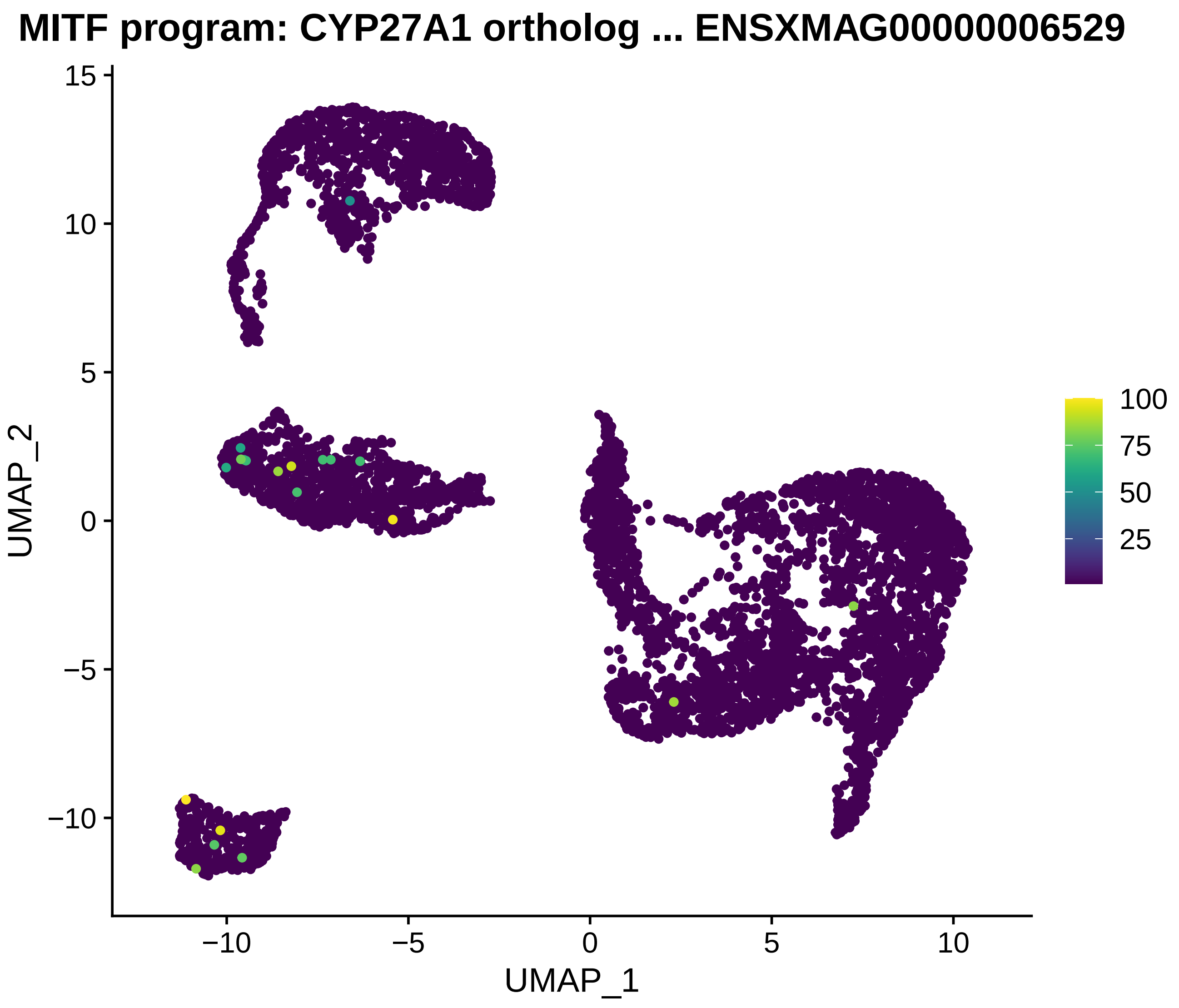
MITF program: CYP27A1 ortholog ... ENSXMAG00000006529
UMAP_2
UMAP_1
15
10
5
0
−5
−10
−10
−5
0
5
10
100
75
50
25

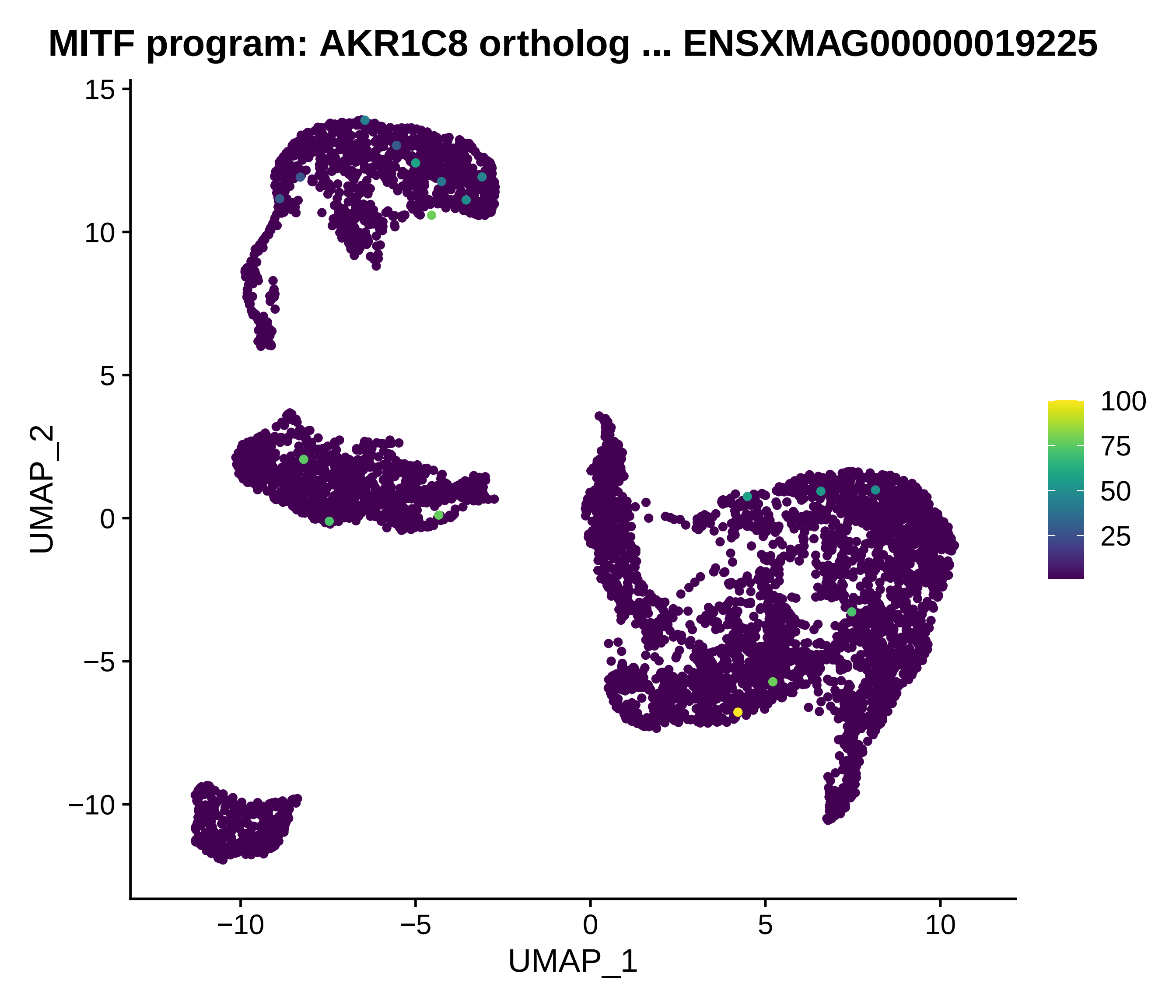
MITF program: AKR1C8 ortholog ... ENSXMAG00000019225
UMAP_2
UMAP_1
15
10
5
0
−5
−10
−10
−5
0
5
10
100
75
50
25

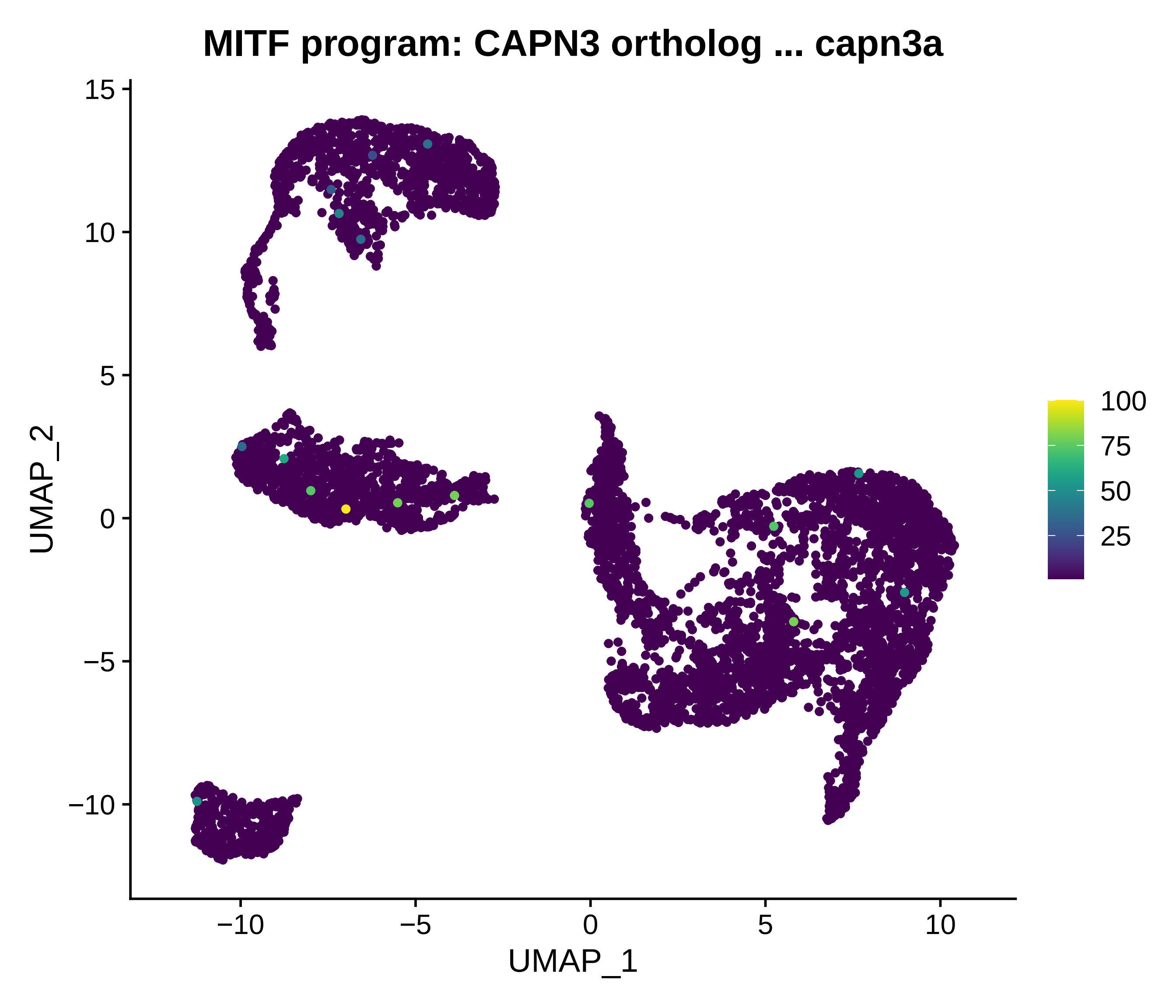

MITF program: CAPN3 ortholog ... capn3a
UMAP_2
UMAP_1
15
10
5
0
−5
−10
−10
−5
0
5
10
100
75
50
25

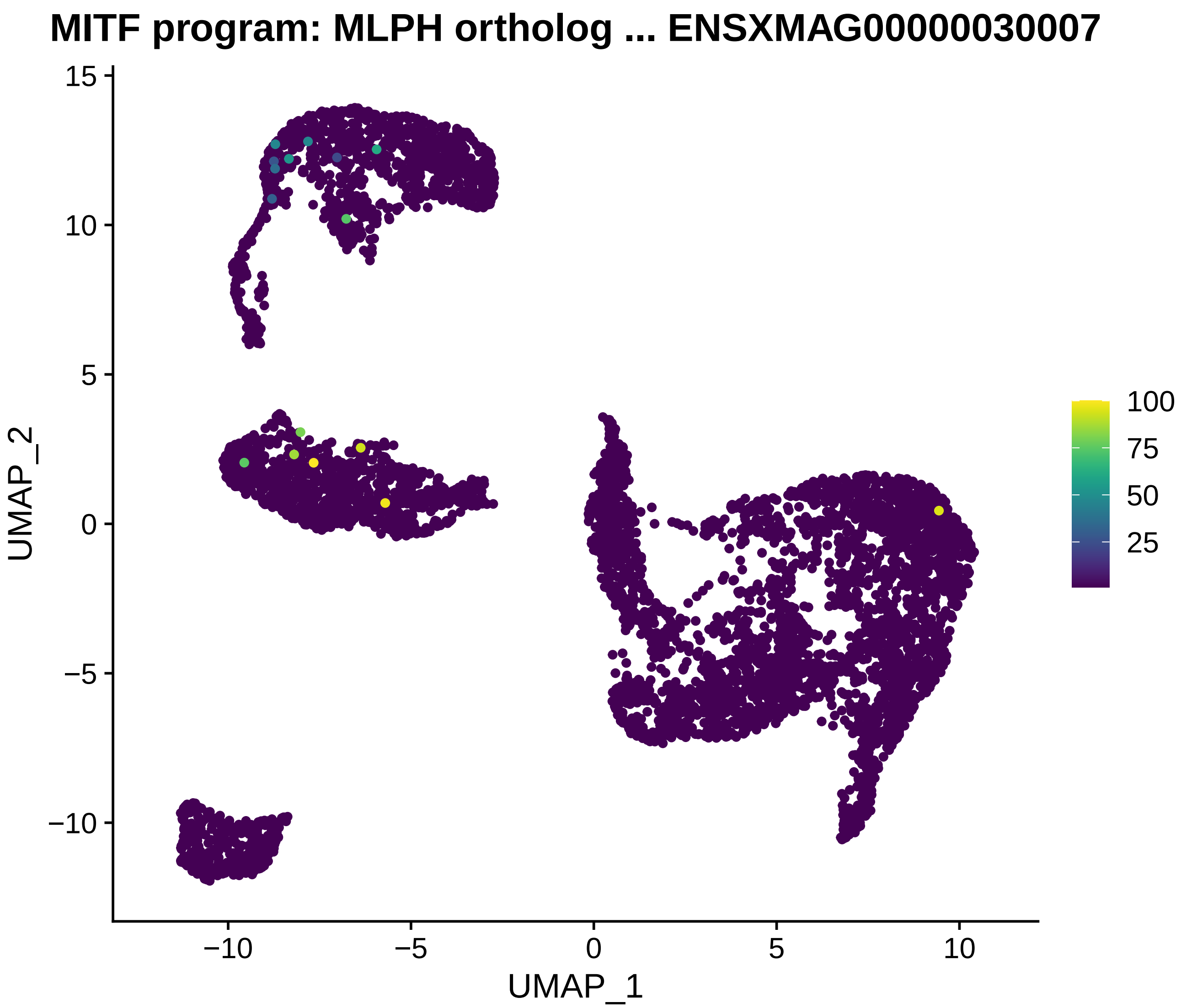
MITF program: MLPH ortholog ... ENSXMAG00000030007
UMAP_2
UMAP_1
15
10
5
0
−5
−10
−10
−5
0
5
10
100
75
50
25

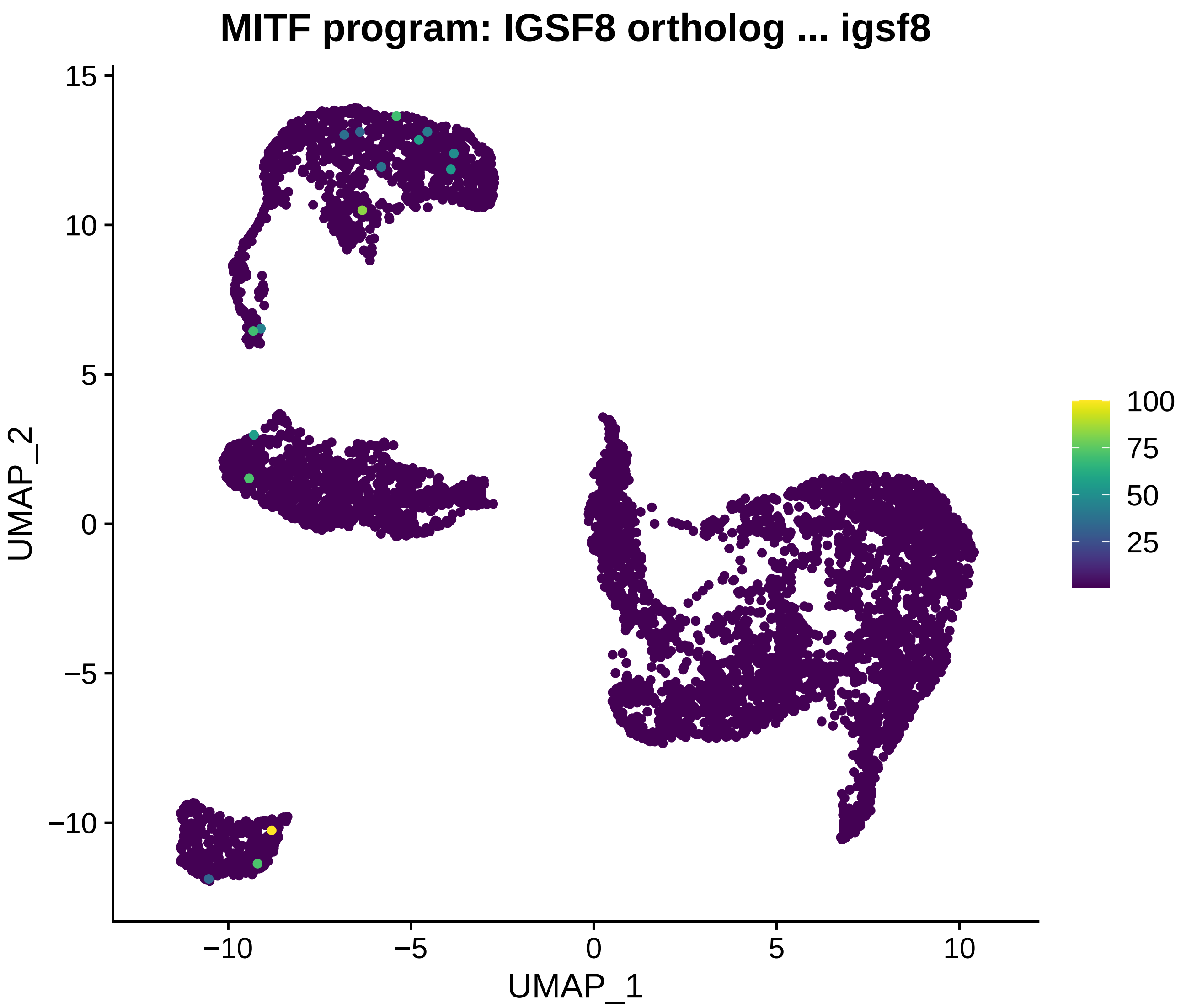
MITF program: IGSF8 ortholog ... igsf8
UMAP_2
UMAP_1
15
10
5
0
−5
−10
−10
−5
0
5
10
100
75
50
25

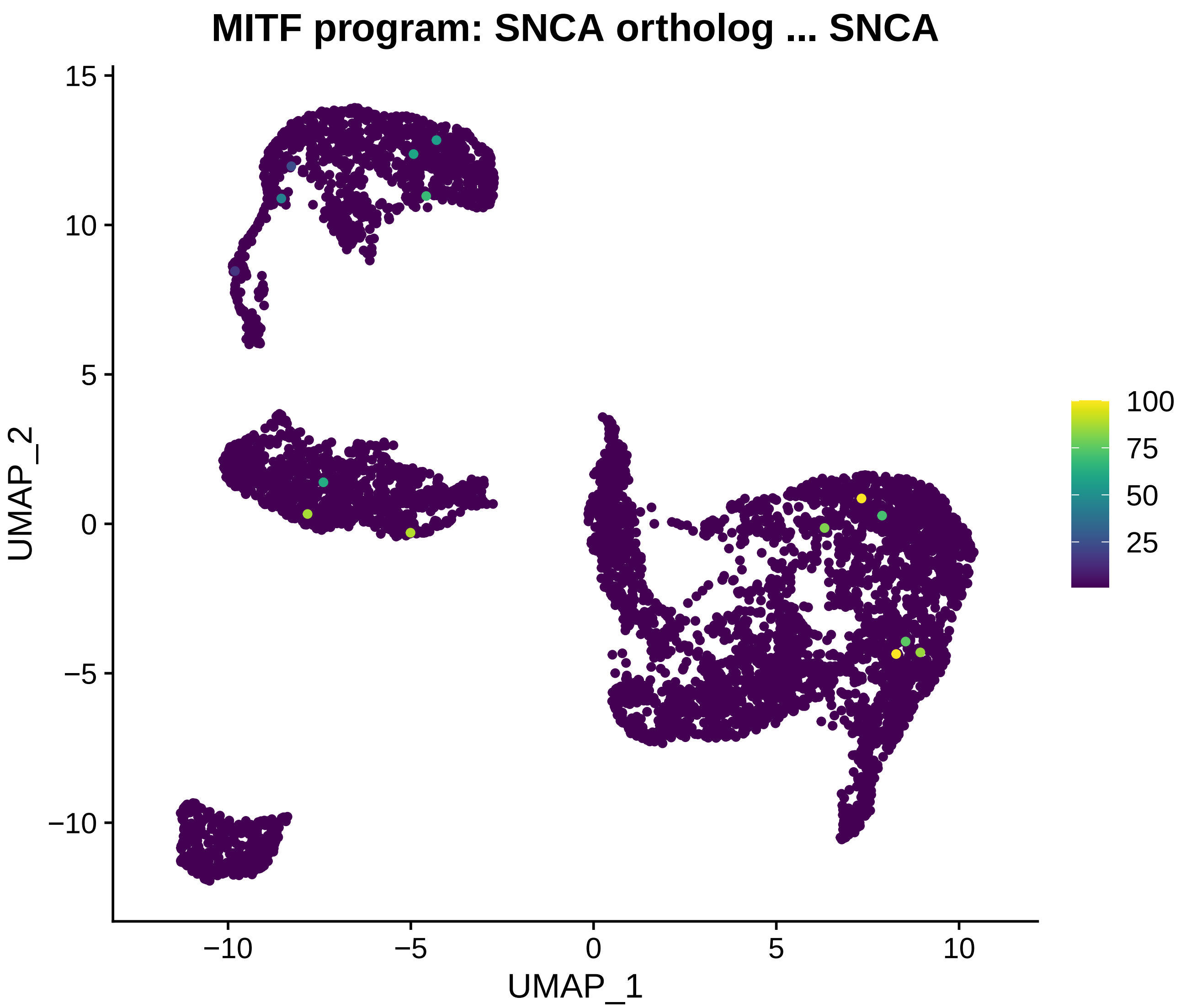
MITF program: SNCA ortholog ... SNCA
UMAP_2
UMAP_1
15
10
5
0
−5
−10
−10
−5
0
5
10
100
75
50
25

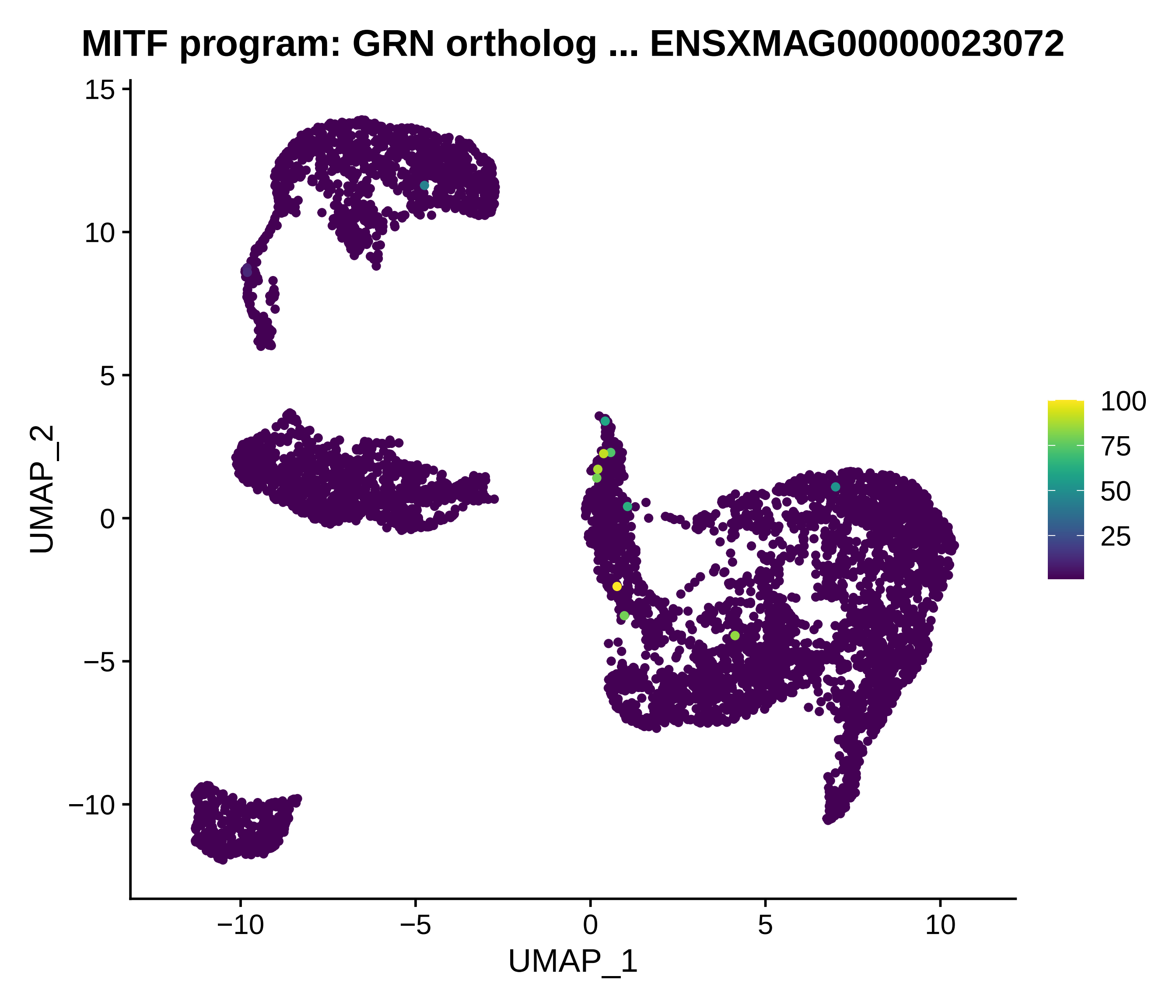
MITF program: GRN ortholog ... ENSXMAG00000023072
UMAP_2
UMAP_1
15
10
5
0
−5
−10
−10
−5
0
5
10
100
75
50
25

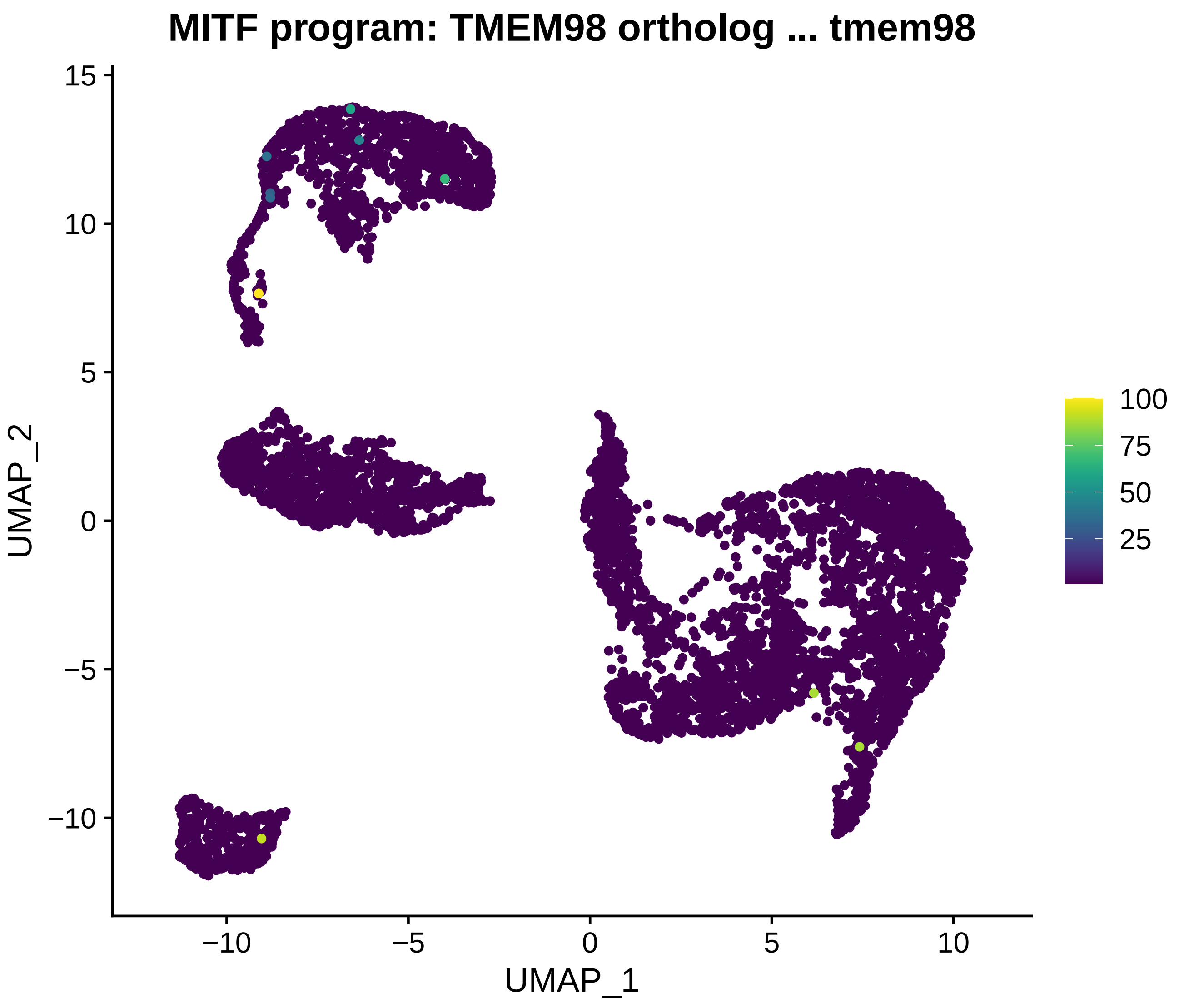
MITF program: TMEM98 ortholog ... tmem98
UMAP_2
UMAP_1
15
10
5
0
−5
−10
−10
−5
0
5
10
100
75
50
25
